# Bayesian updating for self-assessment explains social dominance and winner-loser effects

**DOI:** 10.1101/2023.05.10.540269

**Authors:** Ammon Perkes, Kate Laskowski

## Abstract

In animal contests, winners of previous contests often keep winning and losers keep losing. This coupling of previous experiences to future success, referred to as the winner-loser effect, plays a key role in stabilizing the resulting dominance hierarchies. Despite their importance, the cognitive mechanisms through which these effects occur are unknown. Identifying the mechanisms behind winner-loser effects requires identifying plausible models and generating predictions that can be used to test these alternative hypotheses. Winner-loser effects are often accompanied by a change in the aggressiveness of experienced individuals, which suggests individuals may be adjusting their self-assessment of their abilities after each contest. This updating of a prior estimate can be effectively described by Bayesian updating, and here we implement an agent-based model with continuous Bayesian updating to explore whether this is a plausible explanation of winner-loser effects. We first show that Bayesian updating reproduces known empirical results of typical dominance interactions. We then provide a series of testable predictions that can be used in future empirical work to distinguish Bayesian updating from simpler mechanisms. Our work demonstrates the utility of Bayesian updating as a mechanism to explain and ultimately predict changes in behaviour after salient social experiences.

## Introduction

The winner effect is a widely observed phenomenon where success in one contest leads to increased probability of success in subsequent contests, while the loser effect describes how defeat often leads to more defeats (Chase et al., 1994; reviewed in Hsu et al., 2006). Because of the impact of prior experiences on future success, winner-loser effects can have an important role in the formation of dominance hierarchies (reviewed in Tibbetts et al., 2022). Although it is established that an individual’s position within their dominance hierarchy will have important consequences for their success (Dewsbury, 1988; Simons et al., 2022; Snyder-Mackler et al., 2020), predicting any given individual’s position in social networks remains a challenge (Chase et al., 2002, 2022; Landau, 1951), in part because we lack a full understanding of the behavioural and cognitive mechanisms used by individuals to navigate repeated social interactions (Chase et al., 2002; Tibbetts et al., 2022). Identifying the mechanism behind winner-loser effects and how they function in the formation of social hierarchies could provide powerful insight into what determines individual dominance status and social network structure, but this requires first identifying the predictions of specific mechanisms so that they can be tested empirically.

The existence of a winner or loser effect implies a mechanistic link between past experiences and future contest outcomes. Assuming winning a contest is the result of some combination of intrinsic ability and individual behaviour, there are three mechanisms that could explain winner-loser effects: either (a) winning/losing could modify an individual’s intrinsic ability; (b) contests outcomes could change individual behaviour directly by modifying the individual rules governing that behaviour; or (c) contests outcomes could modify behaviour indirectly via an upstream internal state variable. This state variable could then change behaviour via static rules governing behaviour. While a change in internal state (specifically self-assessment) is commonly cited as the explanation for winner-loser effects (reviewed in Rutte et al., 2006), most existing models of winner-loser effects have modelled self-assessment only implicitly, using one of the first two approaches.

The first wave of models exploring winner-loser effects (e.g., Bonabeau et al., 1996; Dugatkin, 1997; Hemelrijk, 2000), termed Type I models by Mesterton-Gibbons (2016), focused on whether changes in the probability of winning could explain the nature of the resultant dominance hierarchies. Social dominance is complex, but empirical evidence shows that hierarchies are generally stable and contain more linear dominance relationships than would be predicted by chance (Jackson & Winnegrad, 1988; Tibbetts et al., 2022). In Type I models, the existence of winner-loser effects is assumed, and the focus was to explore the *consequences* of winner-loser effects for social hierarchies, specifically their role in promoting linearity, but the underlying *mechanisms* of winner-loser effects often remain something of a black box. These models tend to model the change in outcome directly, either as a linear scaling of resource holding potential (RHP) (Bonabeau et al., 1996; Dugatkin, 1997) (Hickey & Davidsen, 2019; Kura et al., 2016), or as a function of the difference between individual and opponent RHP (Hemelrijk, 2000; Hock & Huber, 2006). While these models explore the predictions of winner-loser effects on dominance hierarchies, they remain structurally agnostic as to whether the change in RHP reflects a change in behaviour, intrinsic ability, or both.

A second class of models, termed Type II models, (Mesterton-Gibbons et al., 2016), test how winner-loser effects can evolve, i.e., whether there are evolutionarily stable strategies that give rise to winner-loser effects. Because these models are interested in the evolution of behaviour, they generally distinguish behaviour from intrinsic ability. In some cases, agents’ estimates of that ability is only implicit (Leimar, 2021; Van Doorn et al., 2003), but two models (Fawcett & Johnstone, 2010; Mesterton-Gibbons, 1999) treat winner-loser effects as an explicit change in self-assessment. To our knowledge, these are the only two models in which winner-loser effects result from an explicit change in self-assessment, but in both cases, self-assessment is incidental to their respective questions, so they limit agents to two or three possible states (big/small, or naïve/post-win/post-loss). While these evolutionary models parse behaviour from individual ability to provide plausible mechanisms that can predict individual behaviour, because of the simplifying assumptions required for evolutionary analysis these mechanisms may only apply to systems where there are discrete alterative tactics (e.g. hawk-dove).

Additional models are needed which identify the predictions of continuous changes in self-assessment and effort, as well as explore how these predictions vary based on the underlying mechanisms of adjusting self-assessment.

If we assume winner-loser effects are driven by modifying uncertain estimates of individual ability, Bayesian updating is a clearly relevant approach. In brief, Bayesian updating entails using Bayes theorem to modify an existing estimate of the state of the world using newly acquired information (McNamara et al., 2006), thereby calculating the precise probability of some given state (e.g., being an individual of size, *x*). Bayesian updating itself thus operates entirely on individual perception, making it ideally suited to describe assessment-based winner/loser effects, and because the relevant features (e.g., size distribution, probability of winning a contest) are generally modelled explicitly, researchers can input relevant knowledge of these distributions during model specification. Because Bayesian updating calculates the precise conditional probability of some event, it provides a theoretical best-case scenario, and existing theory and empirical research indicate that many animals at least approximate Bayesian processes when making decisions in other contexts, for example during foraging or mate choice (Luttbeg, 1996; McNamara et al., 2006; Okasha, 2013; Olsson, 2006; Valone, 2006). Despite being so well-suited to model post-contest changes in self-assessment (Whitehouse 1997), to our knowledge, only one model (Fawcett & Johnstone, 2010) has incorporated Bayesian updating when modelling winner-loser effects, and only in the context of binary ability (Big/Small) and effort (Hawk/Dove) described above. Natural contests are generally decided by the relative size, ability, and effort of competitors—all continuous traits— but no model has used Bayesian updating for continuous abilities and outcomes. Bayesian modelling can be challenging to implement (see McNamara and Leimar 2020), particularly as the complexity of the social information increases, but it is likely that the predictions of a continuous model would differ from a binary scenario, and understanding these specific predictions of changing self-assessments via Bayesian updating could provide important insight into the actual mechanisms underlying the behaviour of various systems. Given the potential power of Bayesian updating as a hypothesis to predict and explain winner-loser effects, it is important to identify the specific predictions of a Bayesian model and understand how we might distinguish Bayesian updating from alternative mechanisms.

Here, we present an agent-based model of winner-loser effects—and dominance formation generally—as a process of Bayesian updating for continuous self-assessment, in which individual agents with self-assessed contest ability compete to win social contests, using Bayesian updating to modify their self-assessments. The purposes of this model are 1) to explore whether Bayesian updating is a plausible explanation of winner-loser effects by comparing the features and limitations of Bayesian updating to empirical observations; and 2) to identify testable predictions that could assess empirically whether animal behaviour is consistent with Bayesian updating, and ideally distinguish it from alternative mechanisms of modifying self-assessments. We show that Bayesian updating can reconcile disparate empirical observations of the attributes of winner-loser effects, while laying out clear, testable predictions that can be used in future experiments, to better understand the mechanisms which drive winner-loser effects specifically and individual dominance status and social structure generally.

### General approach

We were motivated by a scenario where animals, lacking ready access to mirrors, cannot directly observe their own size (used here as a proxy for any intrinsic trait that drives contest outcomes), but are able to observe opponent size (Arnott & Elwood, 2009), and that animals are motivated to win contests without over-investing (Maynard Smith & Parker, 1976; Parker, 1974). For illustration purposes, in figure 1 we depict the agents in our model as a tank of fish of varying sizes, but there is nothing specific about this model to fish, and readers should imagine the individual agents representing any species where conspecific competition occurs and there is some uncertainty about relative contest ability (e.g., rats, baboons, humans).

**Figure 1.**
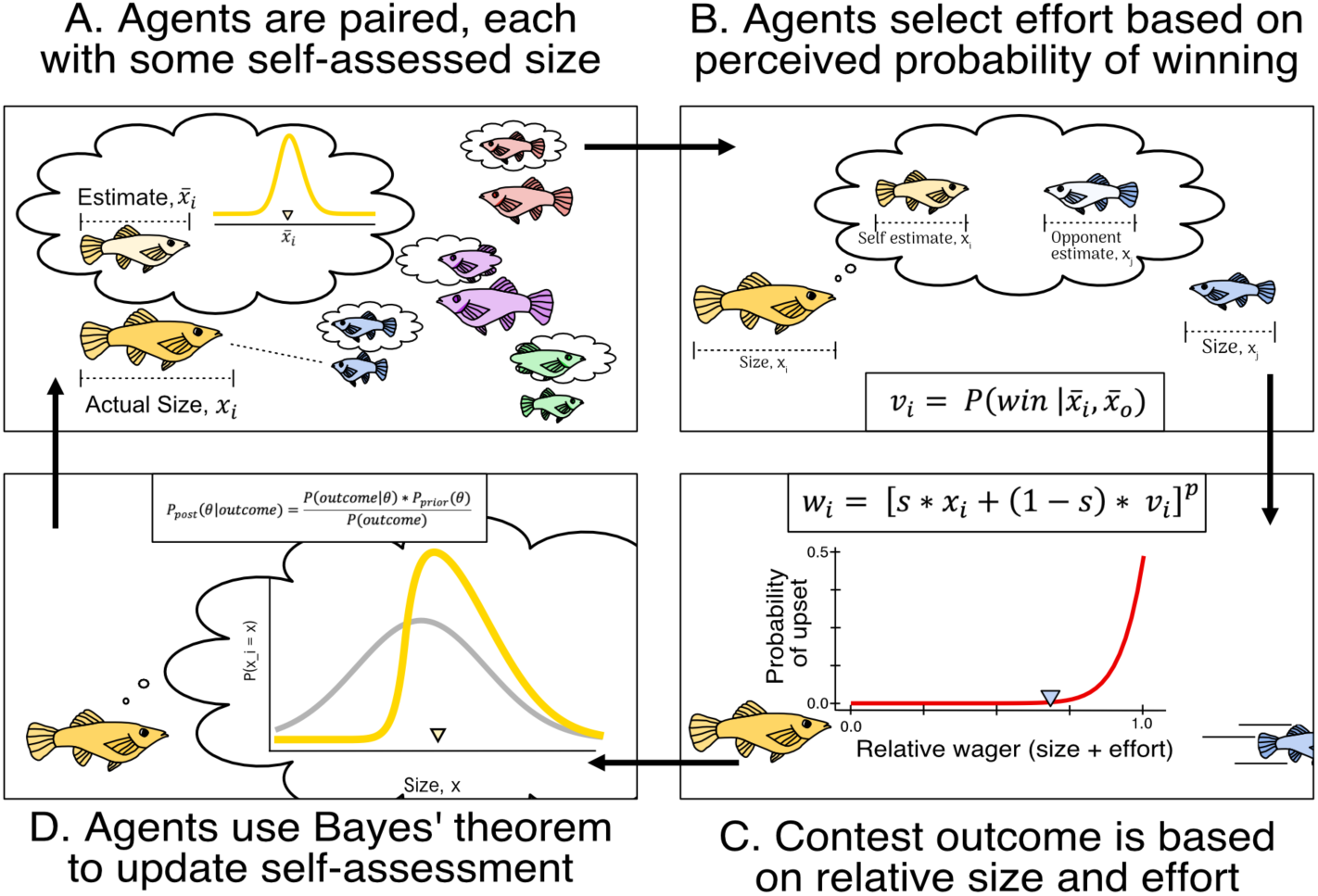
Overview of the model **A**, individuals with fixed *size*, representing intrinsic ability (which is unknown to self), and an estimated self-assessment are paired. **B**, each individual estimates their own and opponent’s size to determine their effort, based on their estimated probability of winning. **C (bottom right)**, the size and effort of each agent determines their relative wagers. The relative wager of the smaller individual determines the probability of winning. **D (bottom left)**, based on this probabilistic outcome, agents update their self-assessment of their size, multiplying their prior assessment by the calculated likelihood of the observed outcome.

**Table 1.** Overview of parameters and their default values.

| <u>Symbol</u> | <u>Default</u> | <u>Range</u> | <u>Explanation</u> |
| --- | --- | --- | --- |
| $x$<br>$: (x, x_i, x_o, \bar{x}_i)$ | Random | $1 \leq x \leq 100$ | Size of an individual. $x_i$ represents the size for individual $i$, while $x_o$ is the size of opponent. $\bar{x}$ is the max-likelihood estimate of size |
| $\mu_x$ | 50 | $1 \leq x \leq 100$ | Mean of the population from which individuals are drawn |
| $\sigma_x$ | 10 | $0 \leq \sigma_x < \infty$ | Standard deviation of population from which individuals are drawn. At $\sigma = \infty$, uniform distribution is used. |
| $v : (v, v_i, v_o)$ | None | $0 \leq v \leq 1$ | Effort, the amount (e.g., of time) agents are willing to invest in a contest. |
| $w : (w, w_i, w_o)$ | None | $0 \leq w \leq 1$ | Wager, or contest performance, a combination of the agent's size and effort. |
| $s$ | 0.7 | $0 \leq s < 1$ | Relative importance of size vs. effort when calculating the weighted sum. |
| $p$ | 6.3 | $0 \leq l < \infty$ | Relative wager modifier, controlling *predictability*, such as $p$ increases, the probability of an upset decreases |
| $\sigma_a$ | 20 | $0 \leq \sigma_a < \infty$ | *self-assessment error*, the accuracy of agents' starting size estimate |
| $\sigma_c$ | 3.1 | $0 \leq \sigma_c < \infty$ | *opponent-assessment error*, the accuracy when guessing opponent size |
| $P(\theta)$ | - | - | Used to denote the prior, the estimated probability distribution of size |
| $n$ | 5 | $2 \leq n < 10$ | The number of individuals in a group, when placed in round-robin style tournaments. |

Under these conditions, we would expect individuals to update their self-assessment following a contest in order to better optimize contest effort. Specifically, we construct a scenario where animals compete for dominance in paired contests where individuals win by outlasting their opponents. This is similar to the war of attrition described by Maynard Smith (1974) which is well suited to describe animal contests where effort and/or persistence determines the outcome, as are common in nature (Enquist et al., 1990; Koops & Grant, 1993; Leimar et al., 1991; Marden & Waage, 1990; Mesterton-Gibbons et al., 1996), but note that this is not quite a game in the mathematical sense, as there are no explicit costs or benefits to winning/losing, and our model is not evolutionary: we specify investment strategies and other aspects of the model based on known features of empirical systems.

Our model consists of four steps, described graphically in figure 1. First, we generate agents with random (normally distributed) intrinsic contest ability and imperfect naïve self-assessments of that ability (figure 1A). For illustration purposes, we will refer to the intrinsic ability as “*size*”, although this could describe any combination of intrinsic traits. Second, these agents are paired with opponents and allowed to interact. In each contest, individuals determine the maximum effort they are willing to invest, which is based on their estimated probability of winning, using their existing self-assessment of their size and their assessment of opponent size (figure 1B). Third, contest outcomes are determined probabilistically based on the relative size and effort of each participant (figure 1C, bottom-right). Finally, following the contest, each agent updates their self-assessment based on their prior estimate of size and their estimated likelihood of the observed outcome (figure 1D, bottom-left). We simulate these contests either in controlled one-on-one contests or small groups where all individuals meet via randomized round-robin pairings. For these group simulations, groups are closed, without immigration or emigration, and are composed of (*n* = 5 agents), lacking individual recognition or social eavesdropping.

### Generation of agents

In order to model the modification of self-assessment, we generate agents with some intrinsic contest ability, called *size*, either specified (e.g., when generating size-matched agents) or drawn from a truncated normal distribution between 1 and 100 (arbitrary units), such that

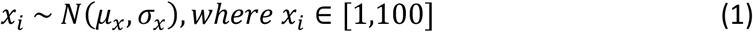

Each agent starts with a point estimate, *x*^-^_l_, of their self-assessment of size, itself drawn from a truncated normal distribution centred around their actual size:

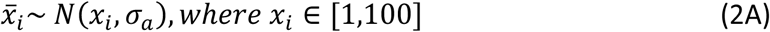

where σ_*a*_ reflects individual *self-assessment error*, i.e., the initial precision with which they can estimate their own size. Because we were interested in scenarios where naïve self-assessment is difficult, for all main results, we set the naïve self-assessment error quite high, σ_*a*_ = 20, but as with all parameters, we vary this value to observe the impact on our predictions (see Appendix A for notes on parameter selection and Appendix B for sensitivity analyses).

Individuals then calculate a starting prior as the truncated normal distribution centred on their estimate, with standard deviation, σ_θ_ = σ_a_,

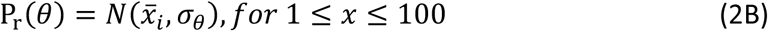

For our model, we will use the term “estimate” to refer specifically to the point value, *x*^-^, which is the maximum-likelihood estimate of the probability distribution of size, while “self-assessment” typically refers to the full distribution.

### Investment strategy for effort

Although the purpose of our model is to explore updating mechanisms, the impact of individual self-assessment is wholly dependent on the rules individuals use to determine behaviour, so we must first establish how agents determine their *effort*, *v*_*i*_, for a contest. In our model, *effort* represents any degree of variable investment in that contest (e.g., the amount of time/energy an animal will expend before yielding). Empirical observations (Hsu et al., 2008) and theory (Enquist & Leimar, 1983; Maynard Smith & Parker, 1976) suggest that the time/energy an individual is willing to invest should be proportional to their perceived potential to win that contest (particularly if it were to continue to escalate), so we designed out investment strategy to match this. To determine *effort* (i.e., maximum investment), each agent first assesses the size of their opponent, by drawing their estimate of opponent size, *x*^-^_o_, from a truncated normal distribution centred on their opponent’s true size,

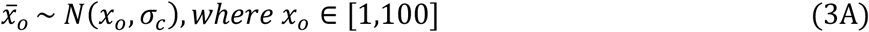

Here the standard deviation, σ_*c*_, represents opponent assessment error. For all main results, σ_*c*_ = 3.1. Agents then set their effort, *v*_*i*_, based on their estimated probability of winning, such that

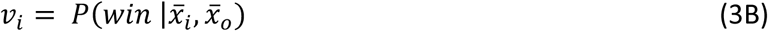

where effort, *v*_*i*_, is equal to the conditional probability of winning given an agent’s estimates about their own and their opponent’s sizes and assuming both agents will equally invest 0.5 effort. Of course, we might expect individuals to better anticipate both their own and their opponent’s effort, and there is a rich literature investigating the evolution of optimal strategies (Maynard Smith, 1974; McNamara & Leimar, 2020), including for winner effects (Leimar, 2021; Mesterton-Gibbons et al., 2016), but establishing a full evolutionary model was beyond the scope of this paper. As our model is primarily focused on how self-assessment changes *following* a contest under Bayesian updating, our specified effort strategy served as a plausible simplifying assumption for our purposes here, although future work should expand this model to observe whether an evolved effort strategy changes the predictions.

Having established their assumptions of own and opponent’s size, agents can calculate their effort, *v*_*i*_, being equal to their perceived probability of winning, via equations (4*A-B*) below. Note that for practical purposes, assessment in our model is instantaneous and conducted prior to the contest, but this also captures scenarios where the assessment occurs over the course of the contest, as is the case in many natural systems (Arnott & Elwood, 2008). Obviously, our approach will not precisely match every animal system, but it models a broad range of systems where animals are uncertain of opponent ability and behaviour, including systems in which size and/or effort are continuous, which may not have been captured by existing models of winner-loser effects.

### Determining contest outcome

Once individuals determine their maximum effort, the winner is decided probabilistically, based on the relative size and effort of each agent, combined to define each agent’s *wager*, *w*_*i*_:

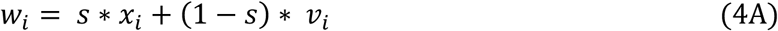

The wager thus scales with individual effort and their size, with *s* allowing us to control the relative importance of *size* and *effort*.

In most natural systems, the contest ends when one individual yields (becoming the loser). For this reason, we calculate the outcome probability from the perspective of the lower-wagering individual, termed the underdog. We thus calculated the probability of an upset (i.e., a win by the underdog) with

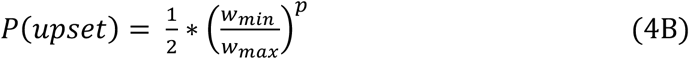

where *p* is a bias parameter, *predictability*, controlling how close the two wagers need to be for there to be a meaningful probability of upset, such that as *p* increases to infinity, the probability of an upset goes to 0 unless the two wagers are equal. Once *P*(*upset*) is established, we use a random number generator to simulate whether an upset occurs.

This approach for settling contests allows us to vary the relative contributions of size, effort, and stochasticity while observing the (simulated) behaviour of individual agents and groups. For all main results, we set the values of these parameters at s = 0.7, *p* = 6.3, which corresponds to a system where intrinsic ability is the most important factor in determining contest outcome, and larger/better individuals generally win, but sufficient effort and/or luck drives occasional upsets.

### Updating self-assessment post contest

In Part 2 of the results, we compare Bayesian updating with a simpler strategy (linear updating) and a null hypothesis (static estimates, i.e., no updating). Each is described in detail below.

These three strategies do not capture the entire range of possible updating mechanisms, but linear updating is a common approach in existing models of winner-loser effects (Bonabeau et al., 1996; Dugatkin, 1997; Kura et al., 2016), and serves as an instructive contrast with Bayesian updating, allowing us to distinguish what predictions are specific to Bayesian updating and what is simply a consequence of changes in self-assessment.

### Static Estimate (No updating)

The simplest possible approach to maintain a self-assessment is to never vary from your initial self-assessment of size (i.e. intrinsic ability). We model this scenario, keeping each agent’s initial assessment of their size fixed throughout the simulation, regardless of contest outcomes.

Under this “no updating” approach, the accuracy of individual estimates depends on their initial awareness, σ_*a*_.

### Linear Updating

A common way to implement simple self-assessment updating, first implemented by Dugatkin (1997), is to either increase or decrease the agent’s estimate of their size, using some scalar, *k*, multiplied by its prior estimate.

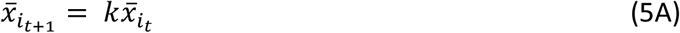

To make this compatible with our model construction, we set bounds on the maximum and minimum possible size (*X*_*min*_ = 1, *X*_*max*_ = 100), such that an agent’s new estimate, *x*^-^_*it*+1_, is a function of their previous estimate, shifted by some factor, where

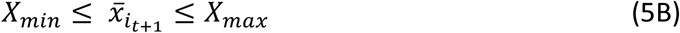

We also set *k* to vary dynamically as a function of distance from the max/min size, where

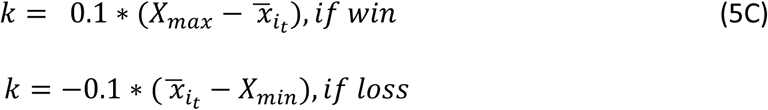

This dynamic shift more closely matches Bayesian updating and prevents estimates from escaping their bounds. Since this function only acts on the point-estimate (defined by the maximum likelihood estimation of the prior distribution), after updating this max likelihood estimate, we generate a new, truncated normal distribution, centred around that estimate, based on their *self-assessment error*, σ_*a*_, as in equation (2*B*). This approach provides a simple heuristic that, like Bayesian updating, allows for weighted updating of self-assessment, but does not change an estimate’s confidence and does not calculating a likelihood function.

### Bayesian updating

In contrast with simpler heuristics, Bayesian updating can in theory calculate the true probability of some value or event, based on known/estimated parameters about the state of the world and the conditional probability of outcomes. Bayesian updating is thus defined by a prior assessment and a likelihood function. In our case, the prior is the agent’s current self-assessment of its own size, modelled as a discrete probability distribution. The likelihood function is the probability of the observed outcome (i.e., winning a given contest), conditioned on being some assumed size, calculated across all possible sizes. Then the posterior distribution, i.e., the new probability distribution of an agent’s size estimate, θ, is calculated by Bayes’ formula

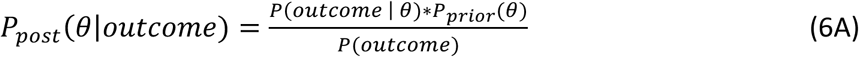

We discuss here the case of a winning outcome, but the formulation is similar following a loss. For each possible size in our discrete size range, the posterior probability of being a given size is

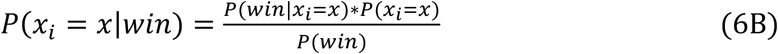

where, *P*(*x*_*i*_ = *x*) is the estimated prior probability of being a given size, while the likelihood

*P*(*win*|*x*_*i*_ = *x*) is the probability of an individual of size *x* winning the previous contest, which is calculated according to the probability of upset from equation (4*B*). *P*(*win*) is calculated as the sum of *P*(*win*|*x*_*i*_ = *x*) ∗ *P*(*x*_*i*_ = *x*) across all sizes, x,

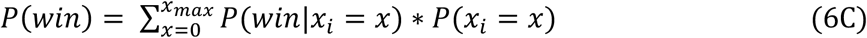

After calculating the posterior estimate, agents update their point estimate of size, *x*^-^_*i*_

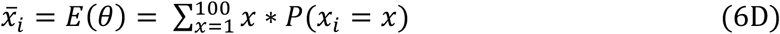

Where *E* is the maximum likelihood estimate of the size distribution. This value is then used to compute agent effort in the next contest.

### Parameter space

Our possible parameter space comprises the contest outcome parameters (*s*, *l*), reflecting the influence of *size vs effort*, and *luck,* respectively; and the assessment parameters (σ_*a*_, σ_*c*_), representing *self-assessment error* and *opponent-assessment error*. For all main figures, we use the values *s* = 0.7; *l* = 6.3; σ_*a*_ = 20; σ_*c*_ = 3.1, the first two were chosen to approximate empirical observations that upsets are rare, and contests are roughly predicted by size (figure A1) (Beacham, 1988; Bierbach et al., 2012), and the latter two to explore a scenario where that opponent assessment is somewhat error prone and naïve self-assessment is poor. We explore the full parameter space in Appendix B.

### Model Analysis

The goal of our model is to answer two questions. 1. Does Bayesian updating for self-assessment produce winner-loser effects and dominance hierarchies that match the behaviour of empirical systems? 2. How can we distinguish, empirically, Bayesian updating from alternative models of dominance establishment and winner-loser effects? To answer these questions, we run simulations of model behavioural experiments while extracting behavioural metrics that would be tractable in and relevant to an empirical context, such as contest intensity. These experiments, and the behavioural metrics we measure, are discussed in their relevant results sections.

## Data availability

We implemented this model in python. All code is available in a supplemental file.

## Ethical Note

No animal subjects were used in this research.

## Results

### Part 1: Matching Empirical Observations

Before addressing how we might test for Bayesian updating, we first compare the predictions of this model to known empirical features of winner-loser effects. Here we focus on three well-documented phenomena: 1. Winner-loser effects exist. 2. The relative strength of winner-loser effects varies across species. 3. More recent contests tend to be more impactful. We also (4) explore how Bayesian updating might account for the observed differences in the duration of winner-loser effects across systems, and (5) the extent to which Bayesian updating promotes stable linear hierarchies. These observations need not be unique to Bayesian updating, indeed there are several simple models that are consistent with these empirical observations, but they are common observations of empirical contest behaviour so it is important to test whether a Bayesian model can also recreate these effects, and this provides insight into how Bayesian updating functions in the context of winner-loser effects.

### 1.1 Bayesian updating predicts the existence of winner and loser effects

We first test whether agents using Bayesian updating show winner-loser effects. To do so, we model a simulated experiment (designed to emulate standard empirical approaches) by forcing either a win or loss in a contest between a focal agent and a “treatment” opponent of known size (in this case, the same size as the focal individual). We then measure winner-loser effects by pairing these focal winners or losers against “assay” opponents (a second size-matched agent). We repeat this process for 1000 focal winners and losers. We can then quantify the winner (loser) effect as the proportion of focal agents which win (lost) against their assay opponent. We would expect 50% of focal agents to win (lose) in the absence of winner (loser) effects.

As shown in figure 2A, we find that agents using Bayesian updating for self-assessment exhibit obvious winner-loser effects, in that previous winners are more likely to win subsequent contests, while previous losers are more likely to lose. These winner and loser effects are broadly observed across the range of parameter values wherever size and effort combine to determine contest outcome, the only exception being where naïve self-assessment was perfect (figure B1).

**Figure 2.**
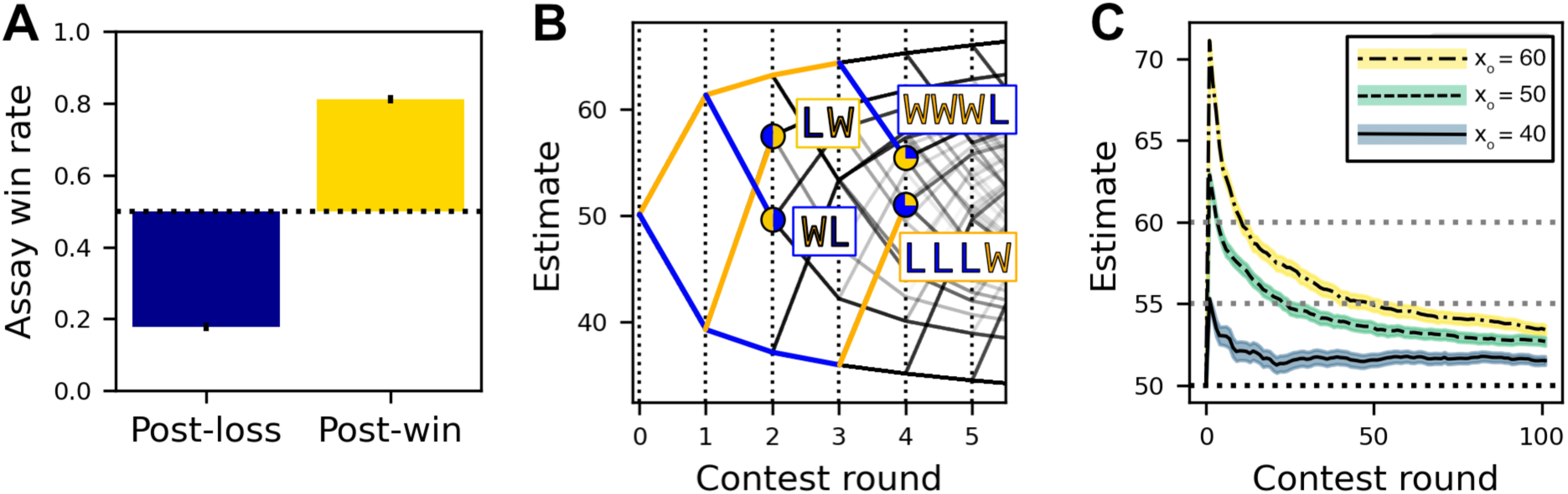
**A,** the proportion of focal agents (out of n=1000) winning against a naïve size-matched assay opponent, after a treatment win/loss. **B,** agent estimates over the course of repeated contests against naive, size-matched opponents, with the branching paths showing the estimates following a win or loss. The boxes note specific trajectories highlighting the recency effect, which diminishes with increased experience. **C.** Focal agents’ estimates change following a staged win against agents of varying size. The shift in the focal agent’s estimate following a win is greater when their opponent is larger.

### 1.2 Bayesian updating can generate variable biases in the strength of winner-loser effects

Given our model’s construction that winning increases an individual’s estimate, estimate determines effort, and effort determines the probability of winning, the fact that Bayesian updating produces winner-loser effects is perhaps not surprising. However, the details of how winner-loser effects function under Bayesian updating are not trivial: We find that, as implemented here, Bayesian updating produces a loser effect that is marginally stronger (on average) than the winner effect, but this depends on the parameters chosen (figure B2). It has long been assumed that loser effects tended to be stronger than winner effects (Hsu et al., 2006), but a recent meta-analysis shows high variation (with some species being winner-biased and others loser-biased) but no general effect. Our model is broadly consistent with this finding, as in our case, the relative strength of winner versus loser effects is highly sensitive to the investment strategy agents use to determine their effort in each contest. Under the simple “proportional-effort” strategy we use for the main results (equation 3*B*), we find a very slight loser-effect bias (figure B2), driven by the fact that winning a contest is determined by the relative size of the smaller individual, and for any given shift, the relative decrease in the estimated size difference is greater than for an increase in estimate. To illustrate this, consider an individual of size 50, that either increases or decreases their estimate by 25 units, and then faces another individual of size 50. Decreasing their estimate results in perceiving themselves as 50% of their opponent (25/50), while increasing their estimate results in perceiving their opponent being 67% of their size (50/75). This effect of proportionality built into agent’s investment strategy drives the slight loser-effect bias in the model described above, but under alternative investment strategies which we explore in the Appendix (equation B1) we can produce agents with stronger biases towards winner or loser effects (figure B3), even with equal changes in self-assessment. The fact that loser-effect bias is mediated by, and highly dependent on, the specific effort strategy used by a given system could account for the high degree of variation in the relative strengths of the winner- and loser-effects observed across systems, based on their specific evolutionary history.

### 1.3 Bayesian updating produces a behaviour-mediated recency bias in winner-loser effects

Although winner-loser effects vary across species, there is consistent evidence for a recency bias (Benincasa et al., 2023; Hsu & Wolf, 1999), in which the outcomes of more recent contests have a stronger effect on behaviour than do the results of earlier contests. To explore this, we emulate empirical tests of the winner-loser effect, by simulating contests between naive, size-matched agents and then pair these focal winners and losers against new, naïve, size-matched opponents, repeating this process over 6 rounds of contests. We control the outcome of each contest to generate new branching paths with both winners and losers. To limit noise, we centre all individual priors on their true size, and agents’ opponent-assessment is set to match opponent size exactly. Here, we measure winner-loser effects directly as the agents’ own size estimates.

In figure 2B, we see that our model recreates winner-loser effects with an experience-dependent recency bias. In particular, “recent losers”, (WL), who first won a contest and then lost a second contest, update their size estimate to be lower than “recent winners”, (LW), who initially lost and then won. This is particularly interesting, since Bayesian updating is known to be “order invariant” in that the order of identical events should not impact the final estimate. However, in this case the events are not identical: the recent winners (LW) win while investing less in their second contest than the recent losers (i.e., initial winners, WL). Because estimate updating occurs at every step, shifting the amount of effort, these events might be better described as LW^-^, WL^+^, since later wins and losses occur under different circumstances which make them more impactful. The strength of this effect depends on the specific parameters used (see figure B4), but under all parameters, recent winners have posterior estimates that are greater than or equal to those of recent losers. Like the other phenomenon described in this section, this recency effect is of course not *unique* to Bayesian updating, but it confirms that Bayesian updating is consistent with empirical observations.

### 1.4 Under Bayesian updating, experimental methodology drives variation in the persistence of winner-loser effects

In empirical observations of dominance contests, winner-loser effects are persistent in some cases (Lan & Hsu, 2011; Laskowski et al., 2016) while in others they are short-lived (Chase et al., 1994) or even absent (reviewed in Hsu et al. 2006). If we assume Bayesian updating, what mechanic could produce these discrepancies? To assess this, we again simulate a forced-win experimental approach, this time allowing a simulated focal agent of size *x*_o_ = 50 to win a staged contest against a treatment opponent of variable size: either smaller, size-matched or larger than the focal agent (*x*_o_ = [40,50, or 60]). We then measure the effects of the contest outcome directly by extracting the estimate of the agent’s own self-assessment. Following this initial treatment contest, focal agents are allowed to interact with additional naïve individuals so that we could observe subsequent changes in their estimates over time.

We found that the duration of the effect depends on size of the initial shift, which is a function of the unexpectedness of the outcome—in this case the opponent’s size—with wins against larger opponents generating larger shifts in agent estimates (figure 2C). This initial increase in self-assessment attenuates rapidly with additional experience, although it does not fully return to baseline, with the mean estimates of winners (for all opponent sizes) persisting significantly above the true individual size (x=50). The extent to which the change in self-assessment—and the associated winner effects—persist above a given threshold depends somewhat on the nature of these post-fight experiences (figure B7), but even after 100 contests, the mean self-assessments of these initial winners are higher than their actual size, and those who won against bigger opponents have significantly higher self-assessments than those who won over smaller ones.

Although under Bayesian updating, winner-estimates remain higher indefinitely, whether such a change in self-assessment could be detected empirically is a function of the power of the experiment used to assay winner-loser effects. If a small shift in estimate (blue line, figure 2C) does not lead to visible changes in behaviour, the winner-effect may appear to be temporary or even non-existent. Social experience following the “treatment” contest, and variation in the size and/or effort of the assay opponent would add additional noise that may further mask small differences in focal-individual estimates. In a simulated experiment, moderate sample sizes with some intervening social experience were generally insufficient to detect winner-loser effects at the behavioural level (i.e., a deviation from the expected 50% win (loss) rate against a size-matched opponent), even though these effects persisted in individual estimates (figure B6)

Indeed it has been shown that in empirical contexts, the experimental methods used influences whether, and for how long, winner-loser effects are observed (Chase et al., 1994; Huang et al., 2011). Given that, within our model, detecting persistent winner-loser effects requires far greater statistical power than is often feasible, it is not surprising that many studies have found winner-loser effects to be short-lived, even if we assume that the winner-loser effect can be perfectly described by Bayesian updating.

### 1.5 Bayesian updating increases the efficiency of size-based dominance hierarchies

So far, we have only assessed winner-loser effects at the level of individuals, but these effects generally function within group hierarchies. It is thus important to model and test how Bayesian updating is predicted to impact winner-effects within social networks, since the marginal increases in effort predicted under experimental contexts may not translate to observable changes within dynamic dominance hierarchies. To do this, we establish groups of n=5 individuals which are paired in randomized rounds such that every individual faces every other individual and recorded the structure and intensity of dominance contests.

The transition to low intensity contests is a standard prediction of established social groups (Jackson & Winnegrad, 1988), as individuals should seek to avoid serious injury from escalated social contests, especially when contests are frequent, as is the case in many group-living species. Although individuals in our model are forced to “interact”, agents choose how much effort to invest, between 0 and 1, allowing them to effectively choose whether to engage in a contest. Future implementations could more fully model the costs and benefits of opponent selection, but here we simply assume individuals are willing to invest proportional to their perceived probability of winning, and ask whether increasing the accuracy of individual estimates decreases the intensity of contests, as well as producing dominance hierarchies that are more linear and stable than we would expect without updating,

We find that our simulated agents using Bayesian updating rapidly reduce the intensity of contests (figure 3A). This occurs in our model as the smaller agents begin to yield quickly to larger opponents, resulting in lower mean effort across contests. This shift is attributable to the rapid decrease in estimate error, calculated using equation (B2), as shown in figure 3B. This increase in accuracy, and the associated decrease in intensity, is widely observed across our parameter space (see figures B7 and B8). For comparison, we repeated these simulated groups using a fixed estimate (figure 3) or a linear estimate (Supplemental), and Bayesian updating results in more accurate estimates which, in turn, result in the rapid reduction in high-intensity contests.

**Figure 3.**
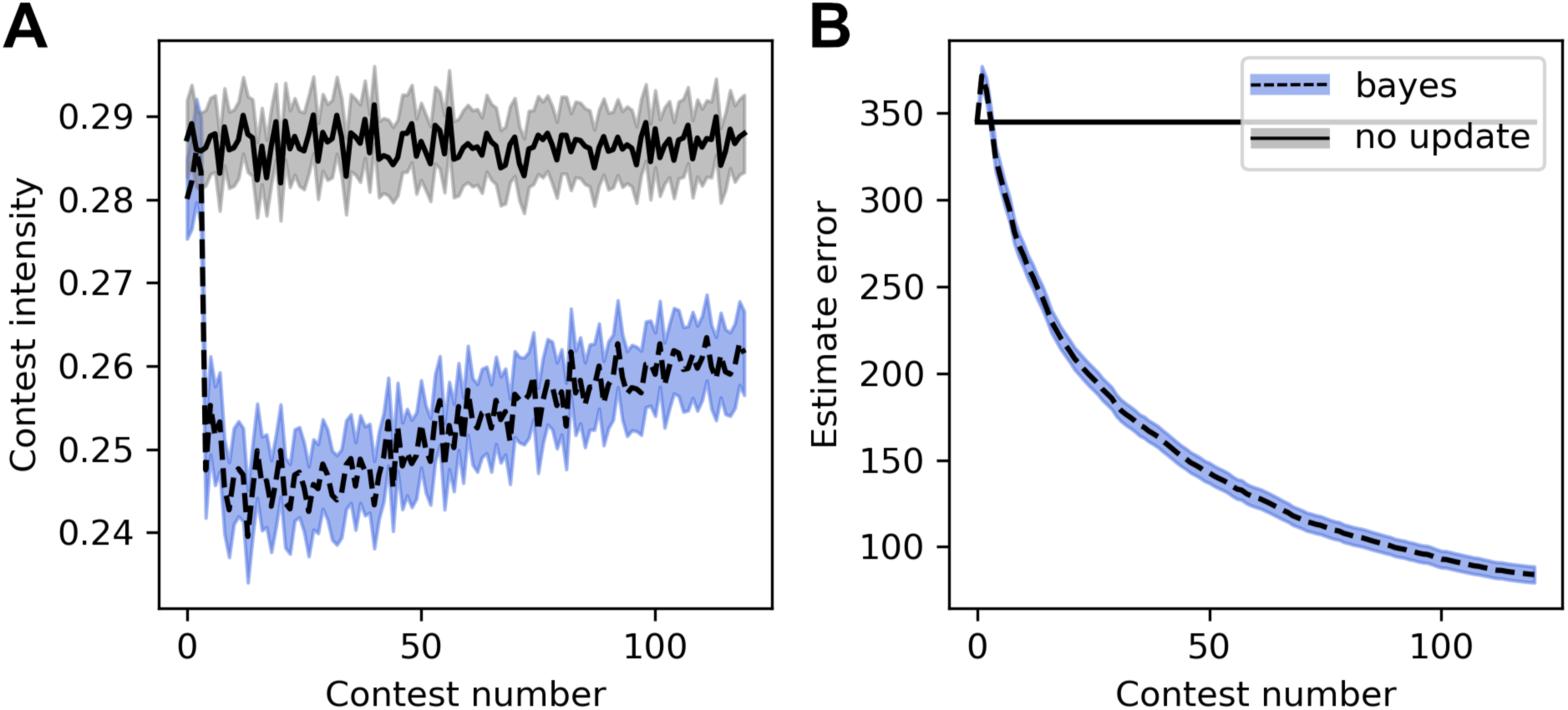
**A,** contest intensity, measured as the effort of the losing individual, decreases with repeated contests as individuals become more confident in their assessments (blue line); however, when agents cannot update, intensity remains high (grey line). **B,** agent estimate error decreases over time with Bayesian updating (blue line), but not when updating is prevented (grey line). The initial spike in error shows the initial winner/loser effects, which cause agents to temporarily over/underestimate their size. Contests over 6 randomized rounds of paired contests between all individuals in a group (n=5 individuals). Lines and shading show the mean result, +/- 1 SEM, averaged across 1,000 simulations.

This improved accuracy also increases the stability and linearity of networks over time (see figures B10 and B11), but these networks do not “self-organize” into stable, linear hierarchies in the absence of intrinsic differences in size. This makes sense, since Bayesian updating results in accurate assessments, so where intrinsic differences are small, individuals accurately assess that they have a high potential to win a contest and invest accordingly. If we modify our model to allow for feedback on agent size (equation B3), networks do organize into linear hierarchies, and in this context, Bayesian updating causes networks to self-organize more quickly and to a greater extent than with feedback on size alone (figure B12*E*). In short, the increased accuracy of Bayesian updating has a stabilizing effect on dominance hierarchies, beyond what can occur from inaccurate self-assessment or changes in intrinsic ability alone.

### Part 2. Distinguishing Bayesian updating from other mechanisms

We have shown that our Bayesian model is successful at recreating many empirical observations of animal contests. We now explore which testable predictions distinguish Bayesian updating from other potential mechanisms of self-assessment, identifying three potential updating mechanisms: Bayesian, linear updating, or static estimates (no updating). In particular, we focus on the two defining features of Bayesian updating: the likelihood-based updating and the probabilistic prior. We investigate each of these two aspects separately, with simulated experiments using empirically tractable approaches to describe potential experiments in animal systems. Their contrasting predictions are summarized in table B2.

### 2.1 The certainty effect is characteristic of Bayesian updating

Because Bayesian updating encodes all previous experiences as a probability distribution, it is able to capture not just the individual estimate but the statistical certainty of that estimate. As such, the statistical certainty gained from increased experience should distinguish Bayesian updating from linear updating, which only maintains the estimate itself. To test for this certainty effect, we simulate focal agents—using either Bayesian or linear updating—engaging in between 0 and 50 contests against naïve opponents, each contest alternating between a win against a smaller opponent and a loss against a larger opponent, thereby producing individuals with similar point-estimates but with different numbers of pre-treatment experiences. We then assess the strength of the winner-loser effect as we would empirically, by pairing the focal individual against a size-matched treatment opponent in which we force a win/loss, and then recording whether these focal winners and losers win against a subsequent naïve size-matched assay opponent. This allows us to measure the extent to which certainty gained through experience attenuates the winner-loser effect of the penultimate, “treatment contest”. As expected, in agents using Bayesian updating (figure 4A, blue line), experienced agents with more prior contests show weaker winner effects than do naïve agents (i.e., for whom the number of fights prior to assay is 0). In comparison, in agents using linear updating (figure 4A, green line), winner effects remain strong regardless of previous experience. Carefully controlling the prior experience is important, because with random opponents, the estimates of linear updating agents tend to diverge over time to either extremely large or extremely small estimates, which can limit their susceptibility to winner-loser effects (figure B13). Thus, depending on the experimental paradigm, this “certainty effect” may not distinguish Bayesian updating from simpler mechanisms, but it remains a core feature of Bayesian updating (across a range of parameter values, as shown in figure B14), and failing to find that winner-loser effects attenuate with increased experience would provide strong evidence against Bayesian updating for continuous self-assessment as a mechanism.

**Figure 4.**
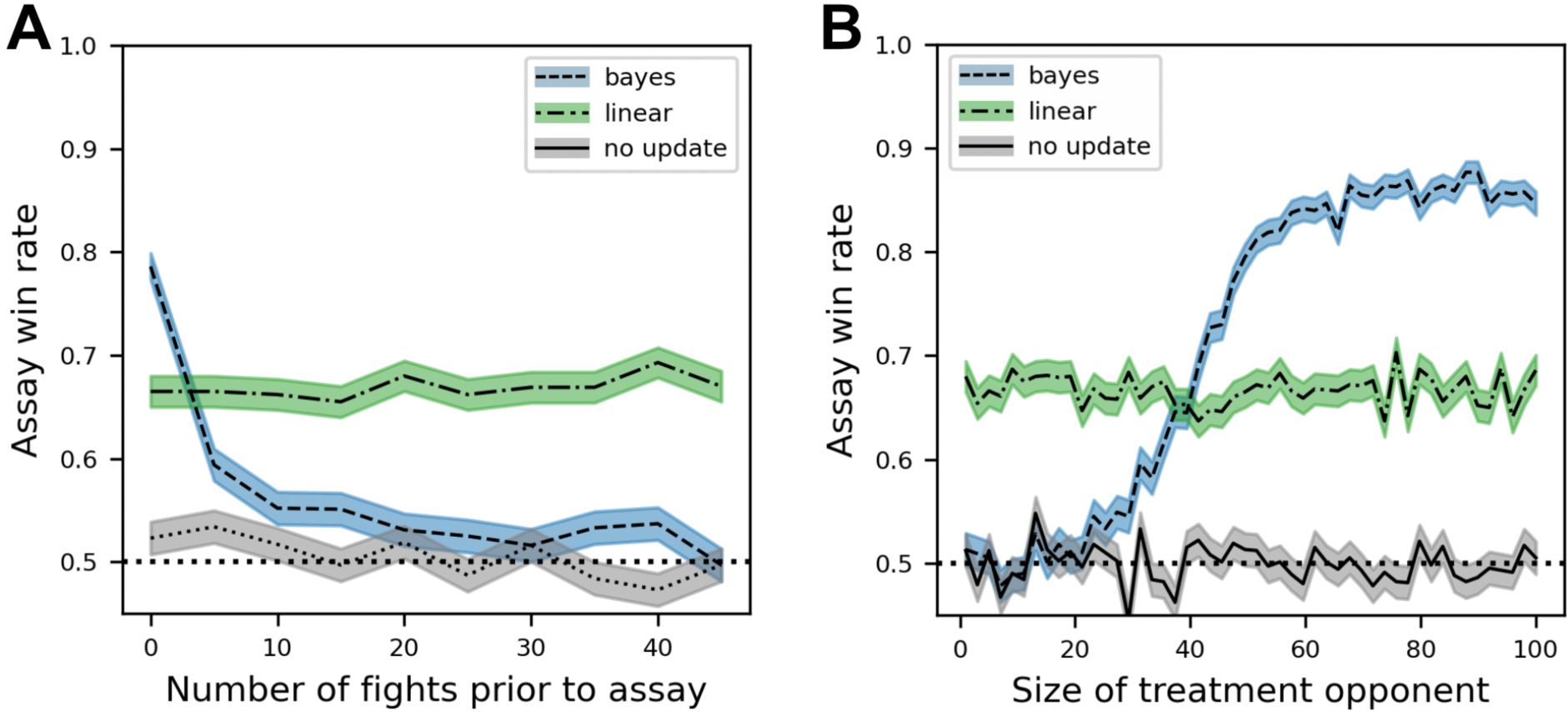
Bayesian updating is characterized by a certainty effect and a discrepancy effect. **A,** under Bayesian updating (but not linear updating), the strength of the winner effect (the proportion of agents winning a size-matched contest after a staged win) depends on the certainty of an agent’s self-assessment and is a function of the number of contests prior to the forced win. **B,** similarly, the strength of the winner effect depends on the discrepancy between expectation and outcome (i.e., the treatment opponent size) under Bayesian updating, but not linear updating.

### 2.2 The discrepancy effect can distinguish Bayesian updating from simpler models

In addition to the probabilistic prior, Bayesian updating is defined by the likelihood function, which calculates the probability of any given outcome based on various possible conditions. As such, we should expect winner-loser effects to vary based on the extent to which the outcome of the contest is surprising. As was already shown in figure 2C, agents using Bayesian updating show stronger winner effects when their initial opponent is larger, and this effect can be observed across parameters wherever size and effort both affect contest outcome (figure B16). Importantly, this “discrepancy effect” is not seen for agents using a linear shift (figure 4B).

Note that while this specific demonstration relies on Bayesian updating having access to opponent size, which is not a factor for linear updating, the general observations that more surprising outcomes drive more dramatic shifts in the posterior estimate is a fundamental feature of Bayesian updating (Courville et al., 2006). As such, this pattern should occur for any observable variable that influences the predictability of outcomes, although what that variable is depends on the specifics of the study system. Furthermore, this test of the discrepancy effect, like the certainty effect described above, cannot distinguish a model that is Bayesian, *per* se, from one that is merely Bayes-like, and linear updating could be easily modified to incorporate the size of the opponent, or the number of past experiences, in order to approximate discrepancy or certainty effect, respectively. But as mentioned above, the lack of any such effect would exclude Bayesian updating as a likely mechanism for modifying self-assessment.

## Discussion

Here, we show that Bayesian updating for continuous self-assessment can explain the existence of winner and loser effects, resulting in contest behaviour and dominance hierarchies that are broadly consistent with those observed within social systems in nature. Our model shows that changes in self-assessment alone, whether through linear updating or Bayesian updating, can be sufficient to drive winner-loser effects (measured as changes in contest outcomes), without any change in actual ability or opponent behaviour. This prediction is consistent with previous self-assessment based models (Fawcett & Johnstone, 2010; Mesterton-Gibbons, 1999), and empirical observations of the winner effect occurring independent of changes to individual ability (Hsu & Wolf, 2001). In addition to being broadly consistent with empirical observations, our model generates testable predictions which distinguish Bayesian updating from the simpler, non-Bayesian linear model. These predictions are designed to lead directly to future empirical tests, allowing us and others to assess the mechanisms behind winner-loser effects. Even if these tests do not conclusively establish the underlying mechanism, they could identify features of winner-loser effects (e.g., does the size of the opponent determine the strength of winner-loser effects?) that have not been previously described.

Like the Type 1 models which explored the consequences of winner effects, our model predicts that Bayesian updating (compared to maintaining naïve estimates) results in increasing the stability, linearity, and efficiency of dominance hierarchies over time (Balph, 1979; Senar et al., 1990), but in our model, this only occurs when there exist some intrinsic differences in ability (figure B12f). Unlike models where winning directly alters contest ability (Bonabeau et al., 1996; Dugatkin, 1997; Hemelrijk, 2000), our model with Bayesian updating alone did not generate linear networks from identically sized individuals (figure B12d). We can extend our model to include direct feedback whereby winning a contest does increase intrinsic ability, and when we do, stratification of initially identical individuals does occur (figure B12b). In this context, Bayesian updating increases the observed linearity of simulated social networks (figure B12e) compared to fixed estimates. Based on these results, we would expect the observed stratification of dominance hierarchies (Tibbetts et al., 2022) to require some real intrinsic differences in individuals, which either existed prior to the contest or which come about because of contests (e.g., as winners gain better access to resources) (Bonabeau et al., 1999).

Accurate self-assessment updating would then act as secondary mechanism which allows individuals to match their effort to their ability, thereby facilitating the stability, linearity, and efficiency of these dominance hierarches.

If we accept that Bayesian updating is a plausible mechanism, how might we test for it? As we have shown, Bayesian updating can be distinguished from a linear updating model by testing for the effects of experience-based certainty and the discrepancy of contest outcomes, which are natural consequences of the probabilistic prior and likelihood functions, respectively. As a note, researchers sometimes use “Bayesian” to refer to individuals that begin a task with informed assumptions. We explore this alternative aspect of Bayesian learning in the supplement (figure B17-19), but we consider the certainty effect and the discrepancy effect characteristic features of any Bayes-like response. While failing either of these two tests would provide strong evidence against Bayesian updating being behind winner-loser effects, these effects are not unique to Bayesian updating, and it will be important to directly compare continuous Bayesian updating to the predictions of other plausible mechanisms.

There are two models in particular whose constructions produce similar predictions to ours. The first is the binary Bayesian model from Fawcett and Johnstone (2010); the second is the actor-critic reinforcement learning model of Leimar (2021, extended in Leimar and Bshary (2022)). In both cases, the most obvious difference between these two models and ours is the extent to which effort and individual ability can vary. Our model focuses on continuous changes in self-assessment and effort. In contrast, Fawcett and Johnstone binarize both effort (Hawk/Dove) and size (Strong/Weak), whereas Leimar allows for continuous ability but keeps contest effort binary (Hawk/Dove). Fully implementing these models for a direct comparison is beyond the scope of our work here, but we briefly explore some of the relevant predictions of these models and how their explanatory mechanisms differ from our model (summarized in supplemental table B2).

In Fawcett and Johnstone (2010), reducing aggression and ability to binary values results in highly stratified estimates, with individuals often becoming “stuck” playing Dove, after which they are unable to gain more information. In our model, because there are many scenarios where individuals win even with very low investment, unlucky individuals tend to recover from initial losses (figure B12c). In natural systems, subordinate individuals often engage with dominants (Guiaşu & Dunham, 1997; Hotta et al., 2021; Rowell, 1974), which could provide an opportunity for recovery, whereby stronger subordinates eventually supplant weaker dominant individuals (Favre et al., 2008; Samuels et al., 1987). In this vein, Fawcett and Johnstone find that their “juvenile” individuals (comparable to naïve individual in our model), tend to be hyper aggressive, because of the benefit of correctly identifying themselves as large early on and being able to invest accordingly. Empirically, it does not appear that juveniles are consistently more “aggressive” (Groves 1978; Bernstein et al. 1983; but see Baxter and Dukas 2017; Fortunato and Earley 2023), although juveniles individuals famously engage in play-fighting (Thompson, 1998; Thor & Holloway, 1984), and as mentioned subordinate individuals tend to seek out agonistic encounters (Guiaşu & Dunham, 1997; Rowell, 1974), so it does appear that naïve and lower ranking individuals highly value information and may engage in contests based on this. In our model there is no explicit reward for seeking information (or for winning contests for that matter). If we expanded our model to incentivise information gathering, we might expect individuals to “over-invest” in potentially informative contests in order to gain valuable information, seeking out information rich encounters despite the cost of more intense losses.

Whereas Fawcett and Johnstone provide a powerful implementation of Bayesian updating within a highly specific context, Leimar (2021) demonstrates a fully realized simulation of social dominance with a fundamentally different learning mechanism than Bayesian updating, i.e., actor-critic reinforcement learning. Like Fawcett and Johnstone, Leimar finds that individuals evolve to be highly aggressive, and interestingly this aggression drives the observed loser-effect bias. This is consistent with our model, in which shifting the effort function to be more “hawkish” results in far stronger loser-effects (figure B4), although the specific mechanisms are different: in Leimar’s model, this shift is specified by their policy gradient factor being inversely proportional to their level of aggression, while in ours, it is a consequence of how an agent’s effort function translates a shift in their estimate to behaviour (figure B20). This comparison is emblematic of these two approaches generally, in which distinct underlying mechanisms (Bayesian updating vs. Actor-critic reinforcement learning) arrive at similar outcomes, which would require careful modelling and very careful experimentation to distinguish.

Indeed, Bayesian updating and actor-critic learning (and the field of reinforcement learning generally) are closely related—since the former specifies the formula to calculate the probability of some expected outcome—and so any effective reinforcement learning paradigm should at least approximate the “correct” behaviour that would be calculated by Bayesian updating. There is a rich field of literature on the application of reinforcement learning to behaviour (Dayan & Daw, 2008; Niv, 2009), how it relates to Bayesian updating (Courville et al., 2006; Kang et al., 2024) and (to a much lesser extent) how reinforcement learning relates to the winner-loser effect specifically (Leimar, 2021). While they are closely related, Bayesian updating functions slightly differently from standard reinforcement learning models (Le Pelley, 2004; Pearce & Hall, 1980; Rescorla & Wagner, 1972). This is because unlike most models of reinforcement learning, Bayesian updating usually establishes a representative model of reality (Courville et al., 2006; Vlassis et al., 2012). For complex systems, establishing this model can quickly become intractable (McNamara & Leimar, 2020), but assuming it is possible, Bayesian updating should provide the most accurate possible solution. This is not to say reinforcement learning is less effective, or even significantly less accurate, as prediction error approaches have been shown to closely approximate Bayesian updating (Kolter & Ng, 2009; Poupart et al., 2006). Given their similarity it seems unlikely that there are large, fundamental differences between the two mechanisms, at least in the general sense. There is some theoretical work that has been able to parse the specific predictions of Bayes updating from other forms of reinforcement learning (Courville et al., 2006; Kumaran et al., 2016), but in our view, the more relevant differences are in the practical application of these models. By abstracting the latent features of reality into the behavioural rule and updating mechanisms rather than having to explicitly infer them, reinforcement learning can be much faster to simulate and analytically more tractable. In contrast, Bayesian updating requires making decisions about a range of internal and environmental state variables in order for the prior and likelihood functions to be meaningful. Empiricists often have access to this information and need to identify testable predictions or perform power analyses. In these contexts, the computational run time of Bayesian updating is less of an issue, while the ability to directly input known environmental variables is highly useful.

While model usefulness is a practical question, Bayesian updating is also a hypothesis for the real underlying biological process, and it is useful to briefly discuss how our predictions of Bayesian updating relate to the current understanding of the biological bases of winner-loser effects. First, as mentioned, Bayes theorem is theoretically the most accurate-possible strategy for dealing with uncertain information and thus represents the target which evolved neurophysiological mechanisms should approximate, whether or not they explicitly infer the underlying variables of interest (Higginson et al., 2018). For example, it is known that the endocrine system is tightly linked with winner-loser effects (Fuxjager, Montgomery, et al., 2011; Fuxjager, Oyegbile, et al., 2011; Zhou et al., 2018). Even without cognitive “learning”, we might expect endocrine mechanisms to reflect the predictions of Bayesian updating with larger shifts in the levels of circulating testosterone and other hormones following a dramatic win or loss, as well encoding learning as the establishment of a new, stable-state (e.g., high circulating testosterone or changes in receptor densities). To our knowledge, the role of contest intensity on endocrine state remain untested and future empirical work could address this.

Although hormones have been broadly implicated in winner-loser effects, winner-loser effects do not appear to be mediated solely by endocrine state (Fortunato & Earley, 2023; Rutte et al., 2006); for many species they are likely mediated at least in part by processes in the brain, where Bayesian updating could be implemented as the explicit mechanism of learning. The neural correlates of winner-loser effects are even less well defined, but if we assume winner-loser effects are driven by Bayesian learning, we might expect expression in the brain following a contest to include genes involved in plasticity and learning, with brains adopting a new, stable-state reflecting the change in self-assessment. We would also expect priors and likelihood functions to be encoded in the brain (Ashwood et al., 2020; Colombo & Seriès, 2012;

Pouget et al., 2013), although identifying where and how these distributions are encoded would be a significant challenge; it seems more plausible that careful theory may be able to identify a method to distinguish Bayesian updating based on behavioural correlates. Regardless of the specific approach, fully identifying the mechanism behind winner-loser effects will require very careful theory paired with careful measures of behaviour and/or the underlying neurophysiological processes involved.

## Conclusion

Given the inherent difficulty of inferring internal processes, and the limitations of any single study, what should empirical and theoretical biologists take from this work? First, Bayesian updating provides a plausible and intuitive framework for winner-loser effects—in which winner-loser effects follow naturally from individual attempts to accurately respond to an uncertain reality—and future experiments can test the predictions we have laid out here to identify novel features of winner-loser effects and assess whether they are consistent with Bayesian updating. More broadly, winner-loser effects should be thought of as a combination of multiple pathways, which interact to link prior experience to future outcomes. In this manuscript, we have explored a few potential mechanisms of individual assessment, updating mechanisms, contest strategies, and how contests are settled, but there are likely additional mechanisms like social eavesdropping (Earley & Dugatkin, 2002; Tibbetts et al., 2020), motivation (O’Connor et al., 2015), short-(Zhou et al., 2018) and long-term (Gherardi, 2006) feedback on intrinsic ability, and individual variation in behaviour (Laskowski et al., 2022) which influence winner-loser effects and dominance contests generally. All of this can be daunting, but experimentation and theory can isolate individual effects to generate at least some diagnostic behavioural predictions, providing models which can predict individual behaviour as well as powerful insights into the underlying behavioural mechanisms.

## Appendices

This supplement is organized to match the structure of the main paper, and is divided into three appendices:

Appendix A provides greater detail and rational regarding the methods;

Appendix B, with subsections matching the results in the main body, explores the parameter space and the sensitivity of the results, while providing greater insight into the relevant mechanisms; and

Appendix C compiles all equations and parameters used for easy reference.

## Appendix A: Extended Methods

As mentioned in the main text, in order to model winner-loser effects, we had to make a series of decisions about the mechanisms by which individuals interact. First, the rule(s) agents followed to determine the maximum effort to invest in a contest (“investment strategy”). We then had to decide how the effort invested interacts with intrinsic ability (size) to determine contest performance (“wager function”). Next, we determined our “contest function”, describing how contests are settled based on the wagers of each individual agent. Finally, we implemented Bayesian updating as the mechanism for modifying self-assessment from one contest to the next. Appendix A provides additional details on the nature of each these assumptions, in order, and our justification for making them.

### The investment strategy was chosen for simplicity and to be proportional to odds of winning

Theory and observations suggest that the amount of time/energy animals are willing to invest in a contest should vary with their perceived probability of winning. Empirical work suggests that strategies in contests vary both across (Arnott & Elwood, 2008, 2009) and within species (Briffa et al., 2015), but in order to maintain our focus on how changes in self-assessment would impact behaviour, for all main results, we assume a very simple investment strategy, in which agents’ maximum effort (*v*_*i*_) is directly proportional to their estimated probability of winning. Because the probability of winning is a function of the relative size of the smaller individual, in equation 3*B*, the assessed probability of winning *P*(*win* |*x*^-^_*i*_, *x*^-^_o_), and thus effort (*v*_*i*_), varies depending on whether an agent estimates itself to be bigger or smaller than its opponent.

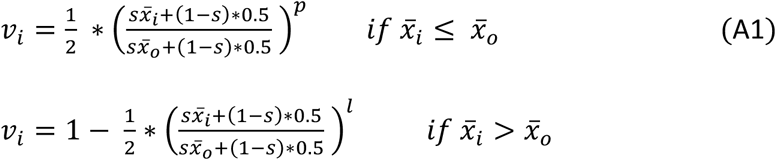

where, *s* and *p* are the simulation parameters used to account for the influence of *size* and *predictability* respectively. With this investment strategy, agents approximate the probability of winning by assuming that both they and their opponent will invest the same amount in the contest (*v* = 0.5).

### Our wager function is designed to explore the impacts of size and effort

Individual effort must interact with individual attributes to determine the effectiveness of this effort. We describe this intrinsic ability as “size”, which encapsulates all intrinsic attributes (body size, weapon size, energy resources, etc.) that impact an individual’s ability to win contests. We thus needed determine how *effort* would interact with *size* (intrinsic ability) to form an agent’s *wager* (their contest performance), and how that wager would determine contest outcome. The way to do this is a sum of these two inputs, which we implement as a weighted sum in order to parameterise the relative impacts of size and effort, as stated in the equation 4*A* from the main text:

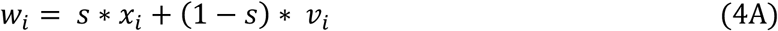

### The contest function allows us to explore the predictability of contest outcomes

For any given contest, each agent supplies this wager, with the larger wager being more likely to win. The “contest function” establishes how this probability is chosen. For example, if both wagers are equal, it is obvious that *P*(*win*) for each opponent should be 50%, but if one agent’s wager is 10% higher, should their probability of winning be 60% or 90%? This is a fundamental assumption of the model that influences agent’s effort and likelihood functions, both in shaping what an optimal strategy would be—if we were evolving the various components of our model—and directly in that the effort and likelihood function takes this calculation into account when determining effort and updating self-assessments.

We attempted to establish a contest function that emulates how contests are settled natural systems. Empirical observations suggest that there exists an S-shaped curve for contest outcome as a function of relative size, that is, contests are highly predictable when individual size is dramatically different, and unpredictable when agent size is very similar, as shown in figure A1 below, reprinted from Graham & Angiletta, 2020 (with permission from the author).

**Figure A1.**
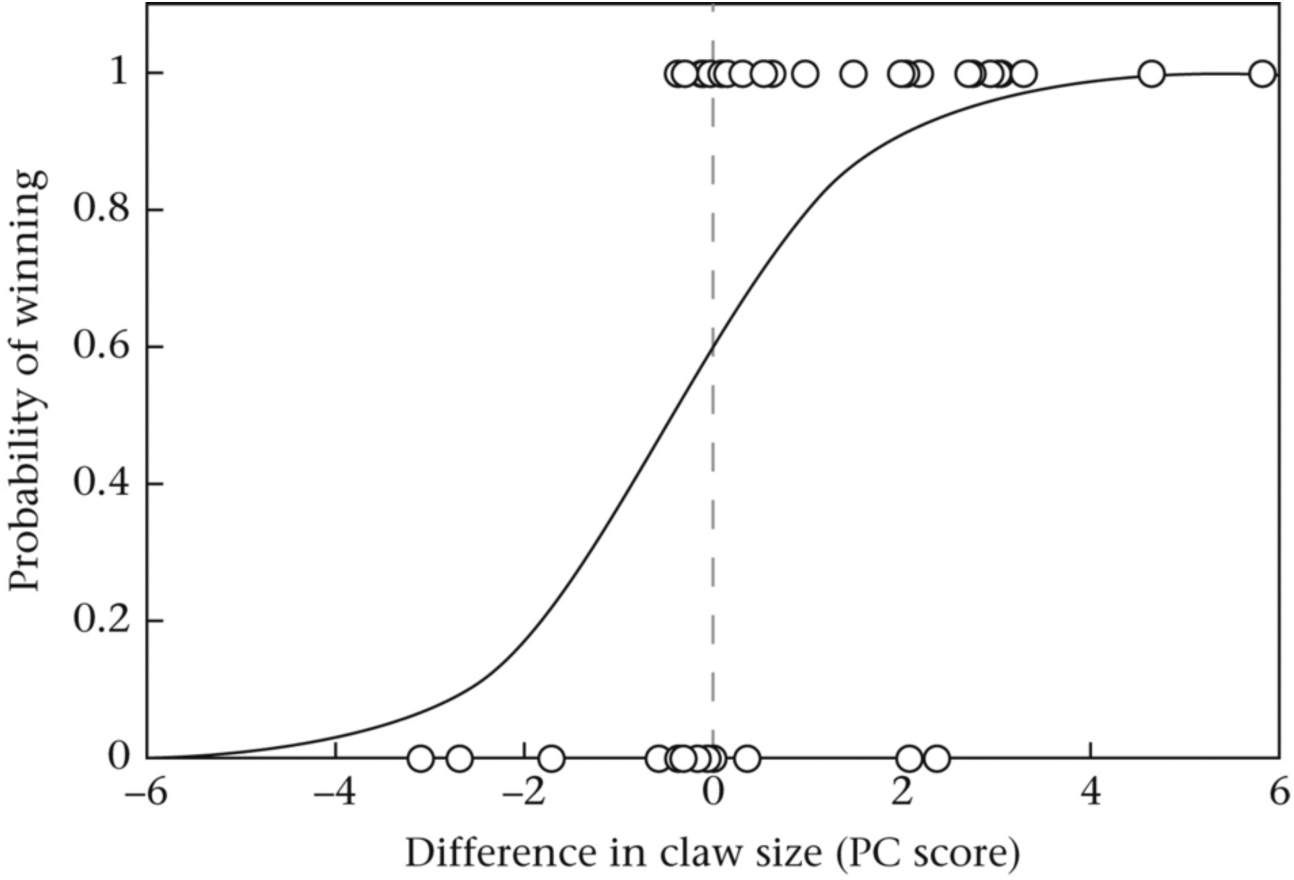
Empirical measure of the role of intrinsic attributes (claw size) in determining contest outcome, (from Graham and Angilletta 2020). In this case, overall body size was matched between participants, in order to observe the effect of claw size specifically. Note that while the sigmoid function used above is fit to the absolute difference in claw size, our model is based on relative size of the smaller individual.

Because contests are generally settled when the subordinate individual yields, it seemed appropriate to model this probability with respect to the lower-wagering agent (roughly, the left half of figure *A1*), so we assume there is some exponential function between the relative wager of the lower-wagering agent and their probability of wining, with the parameter *p* determining predictability, i.e., how convex this function should be. This predictability parameter is a way to model the level of random noise between their expected performance and actual performance in a given contest, which could come from both intrinsic and extrinsic factors.

With this relative wager calculation (equation 4*A*) the agent with greater relative size and effort (i.e., the ‘*favourite’)* usually wins, although it is important to highlight that the smaller wager is not necessarily from the smaller sized individual; if a relatively smaller agent contributed far higher effort, they could have the larger wager, *w*_*max*_, and thus be the ‘*favourite’* for the purposes of our calculations.

### Further details of Bayesian updating

Having established how contests are settled, it is relatively straight forward to calculate the likelihood function as the probability of winning for any given set of parameters and use this to perform Bayesian updating for self-assessment. We describe the basics of Bayesian updating within our model in the main results, but there are some additional details of how we calculate the likelihood function, which we describe here. Because the probability of winning is calculated as function of the smaller relative wager, 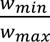, the probability of winning is equal to the probability of upset when the focal agents hypothetical wager is less than the opponent’s wager (*w*_*i*_| *x* ≤ *w*_o_), but is its inverse (i.e. probability of no upset) once the focal agent’s hypothetical wager is greater than the opponent’s:

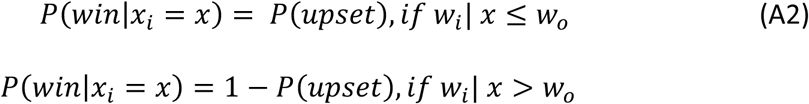

Note that this inflection point is based not on relative size but on relative wager, and agents must take into account both their own and their opponent’s effort into any calculation of the probability of winning, *P*(*win*). Each of these could be modelled as a full probability distribution, but for simplicity, we assume that after the contest, agents have perfect knowledge of the size and effort of the opponent. From a practical point of view, adding these probability distributions would increase any simulations runtime by orders of magnitude, and we would expect them to broadly serve to “flatten out” the likelihood function, limiting the confidence with which agents can update their estimates. Future models could implement uncertainty in opponent size and effort in order to observe whether this changes the predictions of Bayesian updating, or the conditions under which it might be useful.

### Overview of parameter space

There are a few technical details on how we sample across unbounded parameters. These do not have any impact on our results, but we report them here for completeness. Throughout the four mechanisms of our simulation that we describe above (“investment strategy”, “wager function”, “contest function”, and updating), there are 4 variables (*s*, *l*, σ_*a*_, σ_*c*_) which define the parameter space. For *s*, which has the range [0,1], we sample evenly across this range. For the other, unbounded parameters (e.g., σ_*a*_ ∈ [0, ∞)), we use scaling functions for ease of use in implementation.

For *p*, we use the scaling function

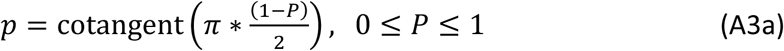

Our default *P* = 0.8, corresponds to *p* = 6.3138. For simplicity, we show the scaled, *p* values in our plots, rounded to the first decimal place, which represent sampling evenly across the range of *P* values. This function can also be used to map P to values of *p* between [0,1], generating a concave contest function, but simulations within this range all behave very similarly to *p* = 0, so we have omitted those results in order to provide better resolution on values between [0,1], which are also far more relevant to our behavioral model.

Similar to *p*, for both σ_*a*_ and σ_*c*_, we use the similar scaling function

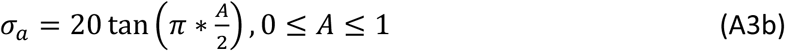

For implementation purposes, evaluating at the limits sometimes requires some conditional statement to avoid errors, for example where σ_*a*_ = ∞, we use a conditional statement to set the assessment distribution to be precisely uniform across the range of sizes.

Although we sample across the range of parameters, for the main results we needed to select a single parameter set. Our size and luck parameters (*s*, *p*), were chosen to match empirical observations as in figure A1. Our default assessment parameters assume a situation where opponent assessment is fairly accurate (σ_*c*_ = 3.1), while naïve self-assessment is very noisy (σ_*a*_ = 20). This is based on the idea that opponent size is relatively easy to observe, but inferring your own size without feedback is difficult, but this is not necessarily the case. We explore the full range of this space in the figures below, including where σ = ∞, in which assessment is a random uniform variable, and σ = 0, where assessment is perfect.

## Appendix B: Exploration of Results

### Part 1. Comparing Bayesian updating to empirically observed phenomena

#### 1.1 Bayesian updating drives winner and loser effects

Figure 2A shows that agents using Bayesian updating display winner-loser effects (i.e., changes in the probability of winning a future contest based on past experience). In our model, winner-loser effects are mediated by 3 mechanisms: 1) the change in an agent’s self-assessment, 2) the extent to which that shift leads to a change in effort, and 3) the extent to which the change in effort leads to a change in contest outcome. Here we explore how the parameter values effect each of these mechanisms to influence the final contest outcome.

Figures B1 (like B2 below) repeats the simulation shown in figure 2A, this time across the range of possible parameters. In short, focal agents are selected to either win or lose against a size matched “treatment” agent, and then tested against a second, size-matched “assay” agent in order to measure the strength of the winner effect. For simplicity, starting priors always begin centred on the true agent size, which is set to *x* = 50. We then compare the change in estimate (pre-treatment vs post-treatment), the change in effort (treatment vs assay), and the assay contest win rate.

With regards to the self-assessment, across all parameters where size contributes to contest outcome, winning increases estimated size, while losing decreases it. The only exception is where *s* = 0, in which wagers are entirely determined by effort, *p* = 0, where contests are entirely determined by luck, and σ_*a*_ = 0, in which agent’s being with 100% confidence of their correct size. Because agent effort is proportional to their perceived probability of winning, the change in effort is roughly proportional to the change in estimate.

There are additional parameter ranges where, despite a difference in agent estimate/effort, there is no observable difference in win rate, notably wherever *p* is close to 0, since this flattens the probability of winning, resulting in similar outcomes. This is not a particularly relevant condition, since it represents a system where contests are entirely determined by chance.

Winner-loser effects also disappear where *s* = 1, since a change in effort cannot impact the outcome where wagers are entirely determined by size. This latter finding suggests that we should not expect winner-loser effects (from Bayesian updating), if animals cannot use increased effort to overcome differences in intrinsic ability. In such a system, any observed winner-loser effect would need to be ascribed to a change in actual intrinsic ability.

**Figure B1.**
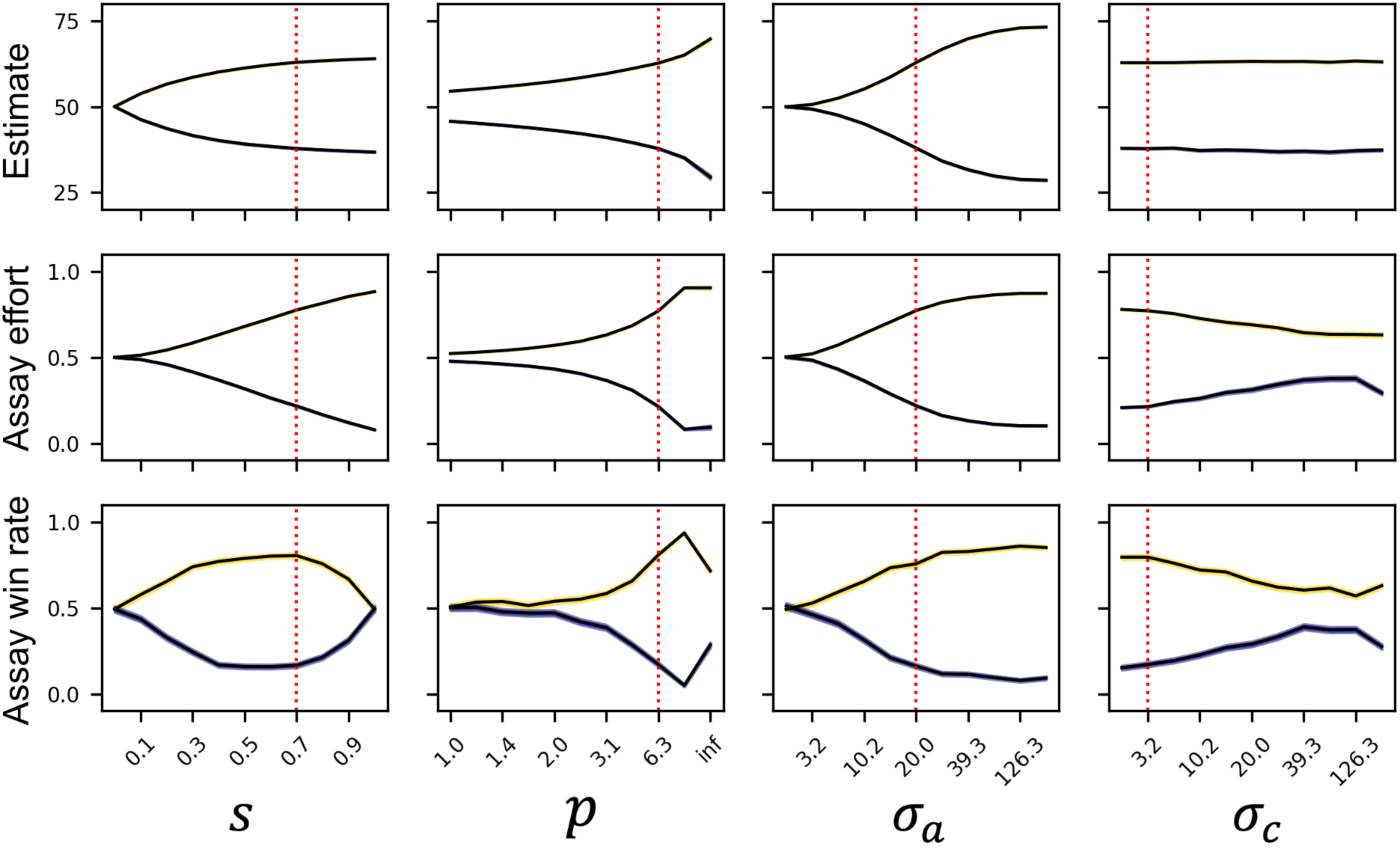
The strength of winner-loser effects varies across a range of parameters, measured as changes in agent self-assessment (top row), contest effort (middle row), and proportion of focal agents who won in assay contests (bottom row). Lines and shading represent the mean outcome from 1000 iterations, +/- 1 SEM. Gold lines show post-win, blue lines show post-loss. Although size and treatment are the same for all agents, inter-simulation variation comes from variation in initial self-assessed estimates, as well as the error in opponent-assessment, leading to variation in effort during both the treatment and assay contests.

#### 1.2 Relative strength of winner and loser effects

As with the existence of winner and loser effects, their relative strength, compared to one another, is mediated by changes in estimate, how that impacts effort, and how effort affects contest outcomes. In figure 2A we can see that there is a slight bias towards the loser effect, but in empirical studies there is variation across species with the relative strength of winner-loser effects, with most species showing a moderate bias towards the loser effect.

Overall, across most parameters, there is a consistent, slight bias in our model towards the loser effect. Comparing the change in self-assessment, the change in effort, and the change in assay win rate (figure B2), we can see that this bias towards the loser effect occurs at the step when agents are determining effort. Interestingly, during self-assessment updating there is a slight bias towards *increasing* estimates for most parameters (figure B2, *top row*). This slight winner-effect bias in updating occurs because the likelihood function is based on the relative size of the smaller agent, and so the probability of winning increases more slowly when the size of the larger individual increases. To understand why this biases estimates to be larger, it is useful to consider a step function, where *P*(*win*) = 0, wherever *x*_*i*_ < *x*_opponent_, and *P*(*win*) = 1, wherever *x*_*i*_ > *x*_*opponent*_. In this context, winning would simply set *P*(*size* = *x*) = 0, for all values bigger than the opponent, while losing would do the same for all values smaller than the opponent. In this situation, there is no directional bias, but consider the scenario where *P*(*win*) = 0.7, *where x*_*i*_ > *x*_*opponent*_, (and still 0 otherwise). Then while winners still set the posterior distribution to 0 for all sizes smaller than the opponent, for losers, there is some probability of having lost while being bigger, and so their posterior distribution is skewed towards larger sizes. While this difference is not dramatic (see figure B2, *top row*), in isolation, we would expect this to create a weak bias towards the winner effect, if contest outcome were based on the change in self-assessment alone.

In contrast to estimate updating, assuming our proportional investment strategy, effort changes more after a loss than after a win, particularly where size is important and initial self-assessment error is high (figure B2, *middle row*). This loser-effect bias seems to be based on the importance of relative size, since increasing an estimate by some fixed amount (e.g., 50 + 25 units) results in a smaller perceived relative change (50 : 75, where 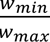 = 0.66) than decreasing your estimate by the same amount (25 : 50, 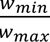 = 0.5). This effort-bias was enough to completely overcome the estimate-bias, resulting in a net bias towards the loser-effect as measured by contest outcomes. This bias is observed across most parameters tested, consistent with the general empirical bias towards loser effect, and during the assay contests, this shift in effort translates to a change in win rate (figure B2, *bottom row*).

**Figure B2.**
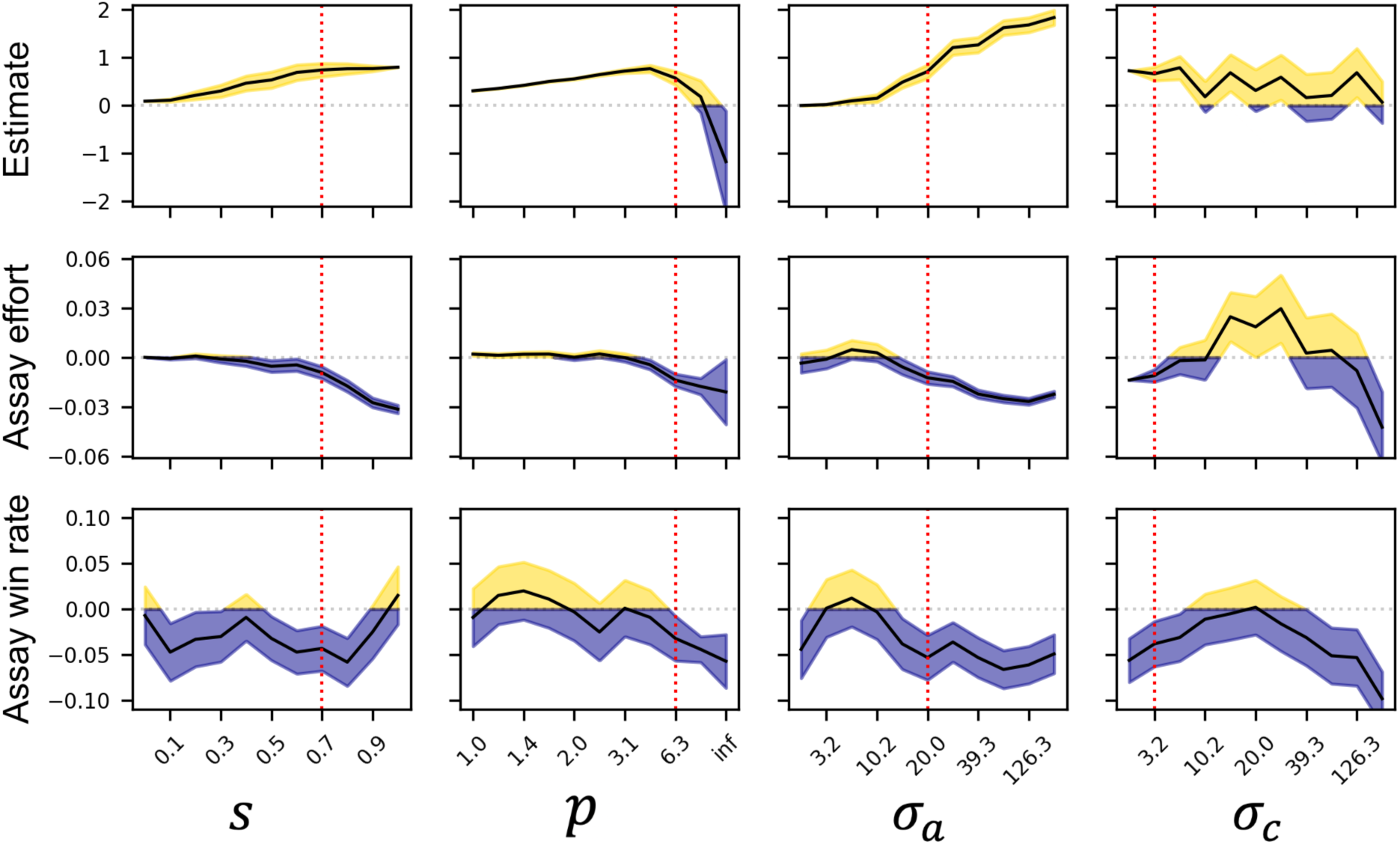
There is weak bias towards loser effects under Bayesian updating. This simulation for this plot is identical to figure B1, but here we plot the difference in the strength of the winner and loser effects (i.e., the difference in the listed measure between winners and losers). The colour here simply shows whether the outcome is biased towards winner effects (gold) or loser effects (blue).

Although in our model, agents using Bayesian updating consistently display a stronger loser effect, this bias is modest (the maximum difference observed is only around 5%). In empirical examples, winner and loser effects can be dramatically different, and in some examples only winner (or loser) effects are observed. Given that the change in self-assessment skews towards greater changes in winners, strong loser-effect bias would have to come from more dramatic changes in effort function and/or contest determination. To further explore this, we generated an alternative investment strategy whereby agents determine effort non-linearly (equation B*1*), and this strategy proves to be sufficient to generate strong winner/loser bias under Bayesian updating. In this strategy, agent effort is no longer directly proportional to the assessed probability of winning, instead agents invest a large amount if their perceived probability of winning is above some threshold, and otherwise invest little-to-nothing, essentially a continuous “hawk-dove” strategy. For simplicity, we use the hyperbolic tangent,

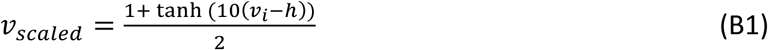

which yields a S-shaped curve in the range [0,1], as shown in figure B3*A* below. Under this investment strategy, we do observe strong loser-(or winner-) effect biases, depending on where this inflection point occurs (figure B3*B*). This demonstrates that the specific investment strategies employed by a given species are a potential source for winner/loser effect bias, even if their self-assessment is relatively unbiased. These investment strategies are likely driven by the specific costs and benefits of competition, which are different across taxa, and those seeking to make predictions for specific systems should seek to identify the relevant underlying functions for that specific context.

**Figure B3.**
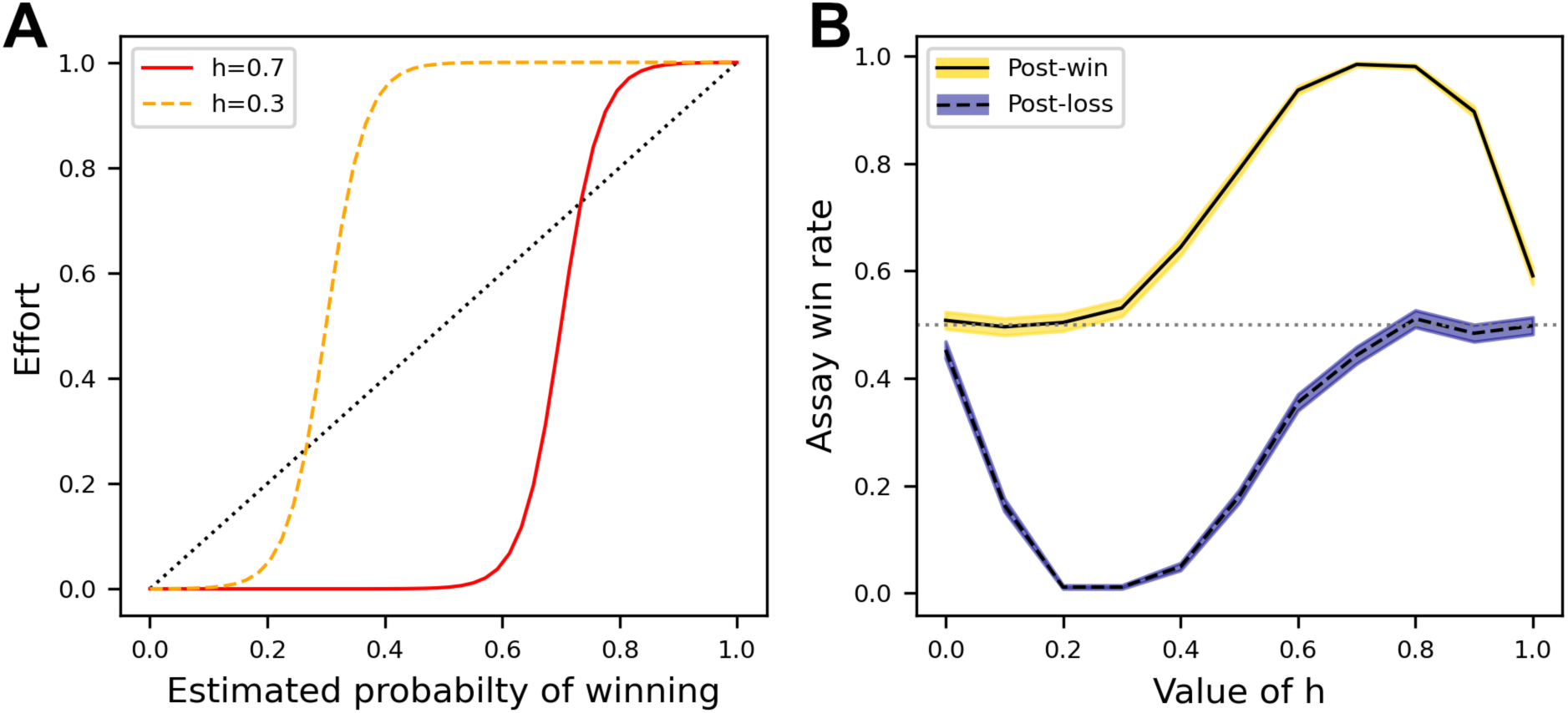
Using a hyperbolic tangent scaling function leads to strong winner/loser effect bias. **A,** the hyperbolic scaling function sets an inflection point at the parameter ℎ, which defines at what probability of winning agents invest 0.5, and more broadly, defines the point where agents transition from very-low investment to very-high investment. **B,** for low values of ℎ, there is a strong loser-effect bias, while for high values of ℎ, there is a strong winner effect bias.

#### 1.3 The recency effect occurs wherever size and effort predict outcome

As shown in figure 2B, agents performing Bayesian updating replicate the empirical observation that more recent contests are more impactful (the “recency effect”). Initially, this would seem to conflict with the fact that Bayesian updating is known to be order invariant. However, this occurs because earlier fights determine self-assessment—and therefore effort invested—in later fights. In other words, winning the first fight will cause an agent to invest more in the second fight. Because recent-winners invest more in the next contest, the likelihood of losing is lower for any assumed size, *x*, except where *x* is very small. This results in recent-winners being more susceptible to loser effects, while recent-losers are more sensitive to winner effects.

To assess the sensitivity of the recency effect, we use a simplified version of the simulation in figure 2B. Using size-matched agents, we compare two conflicting cases, recent winners (LW) and recent losers (WL), and compare their final estimate, effort, and outcome in an assay contest. In this context, there is a consistent recency bias (red cells, figure B4). This is particularly true where both size and effort play a role, and where contests are strongly biased towards the higher-wagering ‘favourite’. If *s* is either small or very large, the difference between recent winners (LW) and recent losers (WL) disappears. Similarly, where self-assessment error is very low, or opponent-assessment error is very high, this effect disappears. There are no scenarios where we observe a significant primacy effect, i.e., the first event overrules later conflicting information. Note that all of this is specific to conflicting recent information (e.g., losing and then winning). For consistent information (e.g., winning and then winning again), the most recent contest results in a less dramatic shift in estimate than the initial contest (see figure 2B).

**Figure B4.**
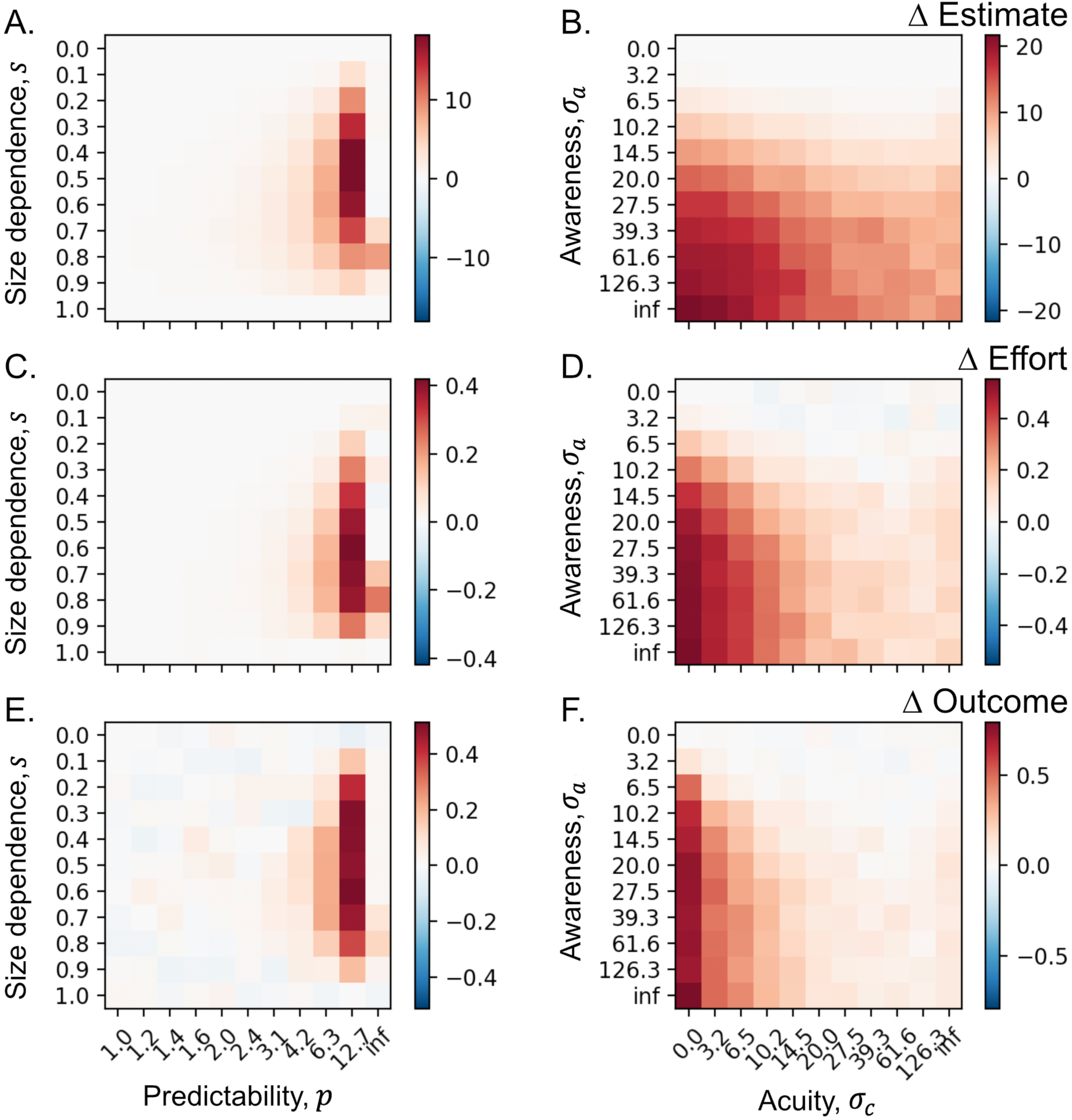
Here agents experienced two contests with conflicting information, i.e., a win then loss (WL) or a loss then win (LW). Heatmaps show the difference between these outcomes, where red corresponds to a measurable recency effect, and darker red to stronger recency effects. The top row (**A-B**) shows the difference in final estimate, showing recent winners were generally higher (red). The second row (**C-D**) shows that the same is true for invested effort. The bottom row (**E-F**) shows the results of a size-matched assay contest. Under all parameters, recent winners are more successful than or (statistically) the same as recent losers.

That said, this recency effect is not necessarily strong enough to overcome any history: with each additional experience, the difference decreases until a single recent win is not enough to supersede the sum of losses (or vice versa). Across all parameters tested, this occurs after no more than 4 prior events (e.g., 4 wins), such that WWWWL always results in higher win-rate than LLLLW.

#### 1.4 Experimental design can determine whether winner-loser effects are observed

Like the observed variation in the strength of winner-loser effects, there is variation in the persistence of winner-loser effects, or more precisely, whether winner-loser effects can be detected. As we show in figure 2C, the persistence of winner effects is a function of both the initial event and the intervening experience. Figure B5 expands this result to depict both winner and loser effects. As in figure 2C, the size of the opponent determines the scale of the initial shift in estimate, and the size of this shift affects the persistence of the winner-loser effect: the greater this initial shift, the longer it lasts (“slow asymptotic updating”, figure B5*A*). Depending on the sensitivity of assay, it may be difficult to identify changes in self-assessment due to less-dramatic treatment contests. Although small differences in self-assessment persist indefinitely, their attenuation over time may mean these differences cannot be detected experimentally.

**Figure B5.**
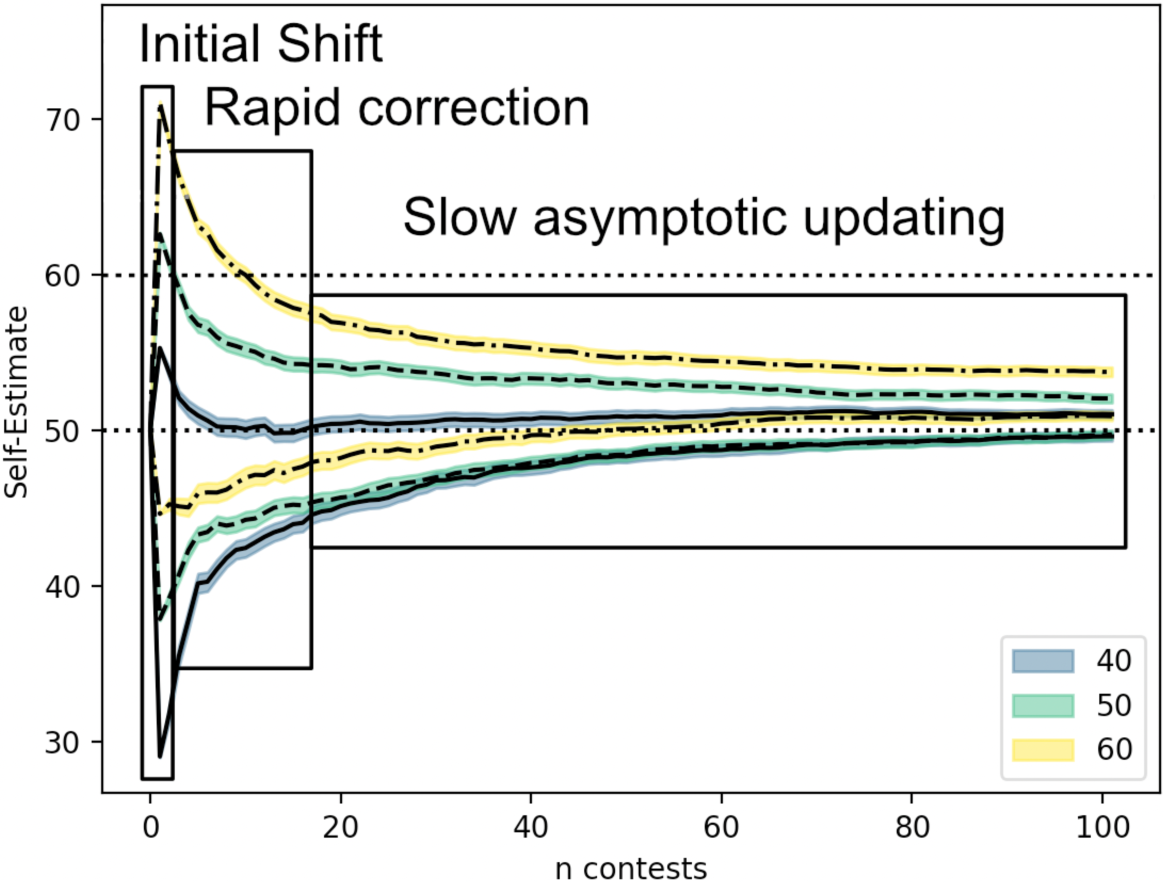
The persistence of winner and loser effects is a function of opponent size and intervening experience. Here focal agents were subject to a single treatment contest (either win or loss) followed by repeated, unbiased contests against other agents. The long-term behaviour of focal agents following a forced win/loss is characterized by an initially large shift, whose magnitude depends on the size of the opponent, followed by rapid correction (which can sometimes overshoot the correct value, as in the blue-winner line), and then a long slow process that asymptotically approaches the true value.

To demonstrate how the experimental approach can determine whether winner-loser effects are detected, we simulated an experiment aimed at detecting persistent winner-loser effects in social contexts, in which we generated paired replicate groups of 5 randomly sized individuals and allowed each group to interact for 5 rounds (such that every individual faced every other individual 5 times for a total of 20 focal interactions). We then “removed” the middle-sized focal individual from each group and forced either a win or a loss against a size-matched opponent. We “returned” the focal individual to the group and allowed them to interact for another 5 rounds. During these 5 rounds, we measured the proportion of fights the simulated focal individuals won against their respective size-adjacent group mate (i.e., the individual that was one size bigger or one that was one size smaller). Following these intervening experiences, the focal individual faced a final size-matched opponent, allowing us to test for winner-loser effects as a percentage of wins for focal winners vs focal losers. Thus, we had three assays of winner-loser effects: the mean proportion of wins in the 5 contests against the next-largest group-mate, the similar proportion of wins against the smaller group-mate, and the mean win rate for the final match against a naïve size-matched assay opponent. We repeated this simulation 100 times for different numbers of replicates, between 5 and 500 pairs of groups.

Not surprisingly, the number of replicates influenced the statistical power (figure B6*B*, measured as the probability of detecting a significant difference using a t-test comparing the assay scores of winners and losers, using a t-test). However, under these conditions, 200 replicates were required to detect a significant difference greater than 50% of the time, which in this case would require around 2000 individual animals. While size-matched tests yielded the strongest mean difference (figure B6*A*), because of increased power of observing multiple contests against groupmates, all approaches were roughly equally likely to detect a significant difference between winners and losers over time. Under these assumptions, detecting persistent winner-loser effects would require a very strong initial shift, an extremely sensitive assay (for example reducing all variation in size and effort of the various individuals), or extremely high sample size.

**Figure B6.**
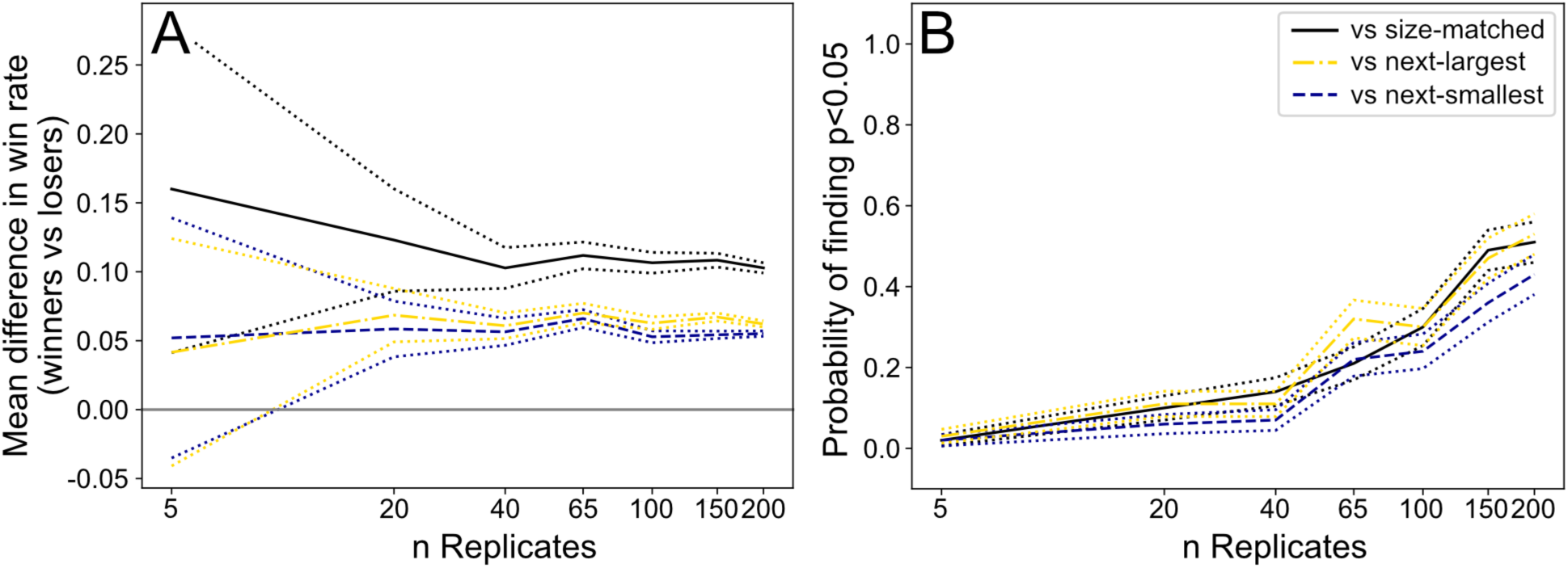
Power analysis of detecting persistence of winner-loser effects, based on a simulated experiment where individuals were removed from their group, given a forced-win or -loss against a “treatment” size-matched opponent, and returned to their group, after which winner-loser effects were assayed using a naive size-matched opponent. We simulated this experiment using varying numbers of replicates, shown on the x-axis. **A.** shows the mean difference in forced-winners or forced-losers, measured as the win rate against the size-matched assay opponent (black), the next-largest groupmate (gold), and the next-smallest groupmate (blue), with dotted lines showing the standard error across 100 iterations. **B.** shows the proportion of simulated experiments in which there was a significant difference between winners and losers across the 3 metrics used, with dotted lines against showing standard error.

#### 1.5 The amount of novel information determines the persistence of winner-loser effects

We see in figure B5 that the initial change in estimate attenuates over time. Figure B7 shows that the speed of this attenuation (i.e., the number of rounds required to recover) depends on the diversity of info, or more precisely, the frequency with which an agent receives information which conflicts with its current self-assessment. Thus, with a greater variety of opponent sizes and efforts, agents can more quickly return to near-accurate self-assessments. Similarly, larger mean-sized opponents will quickly erase a winner effect, but prolong a loser-effect (see green dash-dotted line in figure B7*A,C*). Put simply, repeatedly competing in uninformative contests limits the accuracy of individual self-assessments and thus can extend the duration of winner-loser effects.

**Figure B7.**
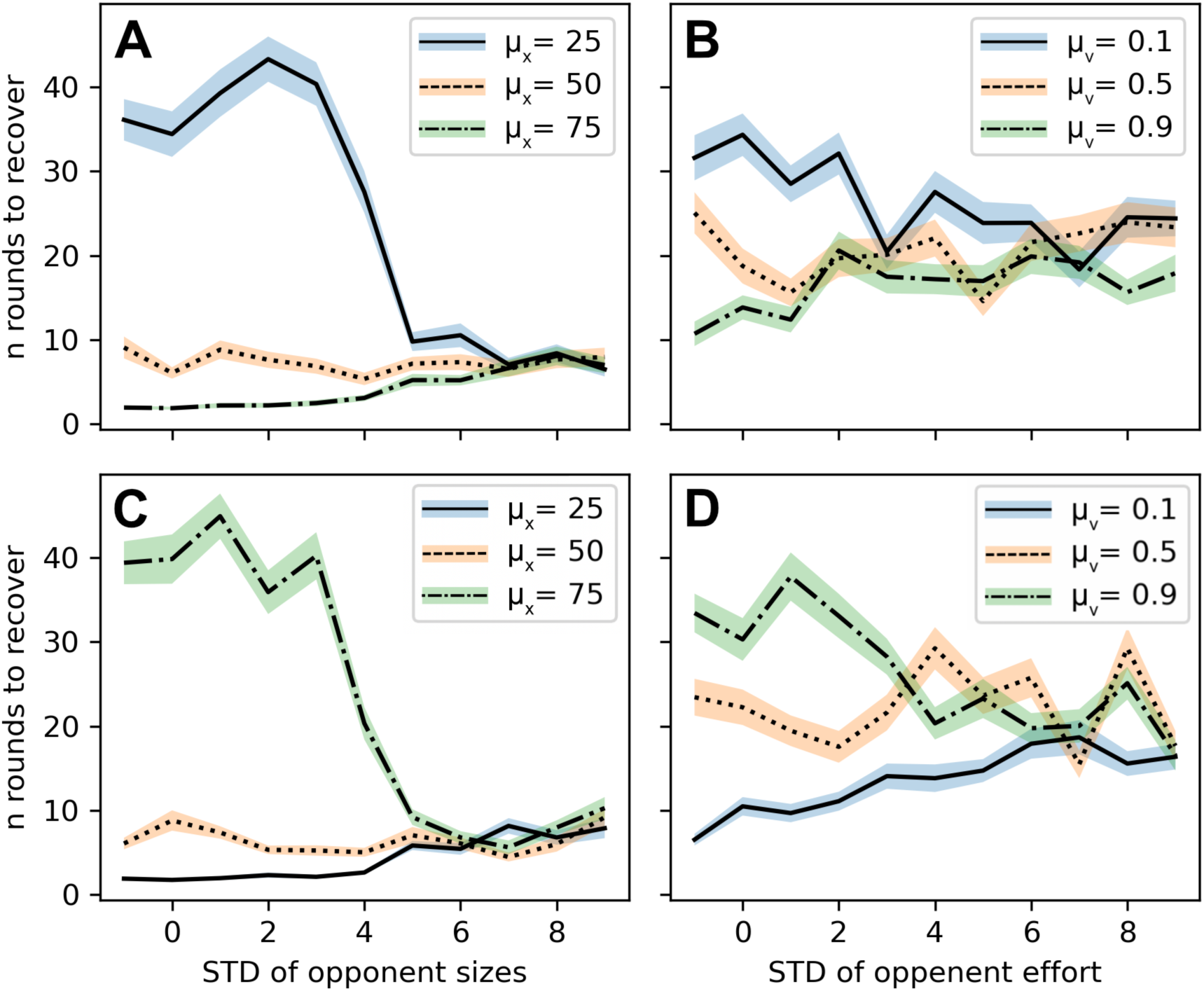
The top row (**A-B**) shows the time to recover from a win. Recovery is the number of rounds needed for an agents estimate to return to within 10 units of actual size (see figure B5). Here each round represents a new individual with randomly selected size and effort. (**A**) shows the number of rounds to recover with respect to the standard deviation of opponent sizes, plotted for three different mean sizes, while (**B**) shows this as a function of the standard deviation of opponent effort, again plotted over 3 different mean efforts. **C** and **D** show the same metrics after a staged loss.

#### 1.6 Bayesian updating stabilizes dominance hierarchies above and beyond intrinsic feedback

So far, we have discussed winner-loser effects in the context of isolated focal agents; however, in nature, winner-loser effects generally take place in networks of interacting individuals. As we see in figure B5, under Bayesian updating, agents tend to successfully approximate their true size—and thus behave accordingly. Because of this, in the absence of direct feedback on size, the long-term behaviour of dominance hierarchies under Bayesian updating is largely a function of the real intrinsic differences of the agents. In figure 3, we show that Bayesian updating increases the accuracy of estimates and lowers the intensity of contests compared to no updating. Here we further explore the effects of Bayesian updating on hierarchy structure, i.e., linearity and stability, in addition to accuracy and contest intensity, over a range of parameters, and observe how this compares to and interacts with direct feedback on intrinsic ability. As in other placed in this paper, we explore these metrics under the three updating schemes used in this paper: Bayesian updating, linear updating, and fixed estimates (no updating).

As in figure 3, we simulate groups of varying numbers of agents and allow them to interact over 10 rounds, 10 ∗ (*n* − 1) contests per individual, simulated over 100 iterations, from which we calculate accuracy, intensity, linearity, and stability. We describe how each of these are calculated below, before discussing how they vary with the parameters chosen.

To calculate accuracy, we measure the distance from the true agent size, *x*_*i*_ and the edges of the confidence interval, *x*^-^_*i*_ ± σ, such that

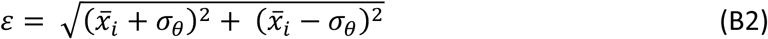

Under almost all parameters, estimate error decreases compared to initial estimates (the exception being the edge cases where size has no impact on contest outcome, i.e., *s* = 0 or *p* = ∞). Agents that do not perform updating obviously do not improve the accuracy of their estimates (figure B8, grey shaded line). Interestingly, linear updating leads to less accurate self-assessment, as individual estimates tend to stratify to be either extremely large or extremely small (depending on whether they win or lose more than half of their contests).

**Figure B8.**
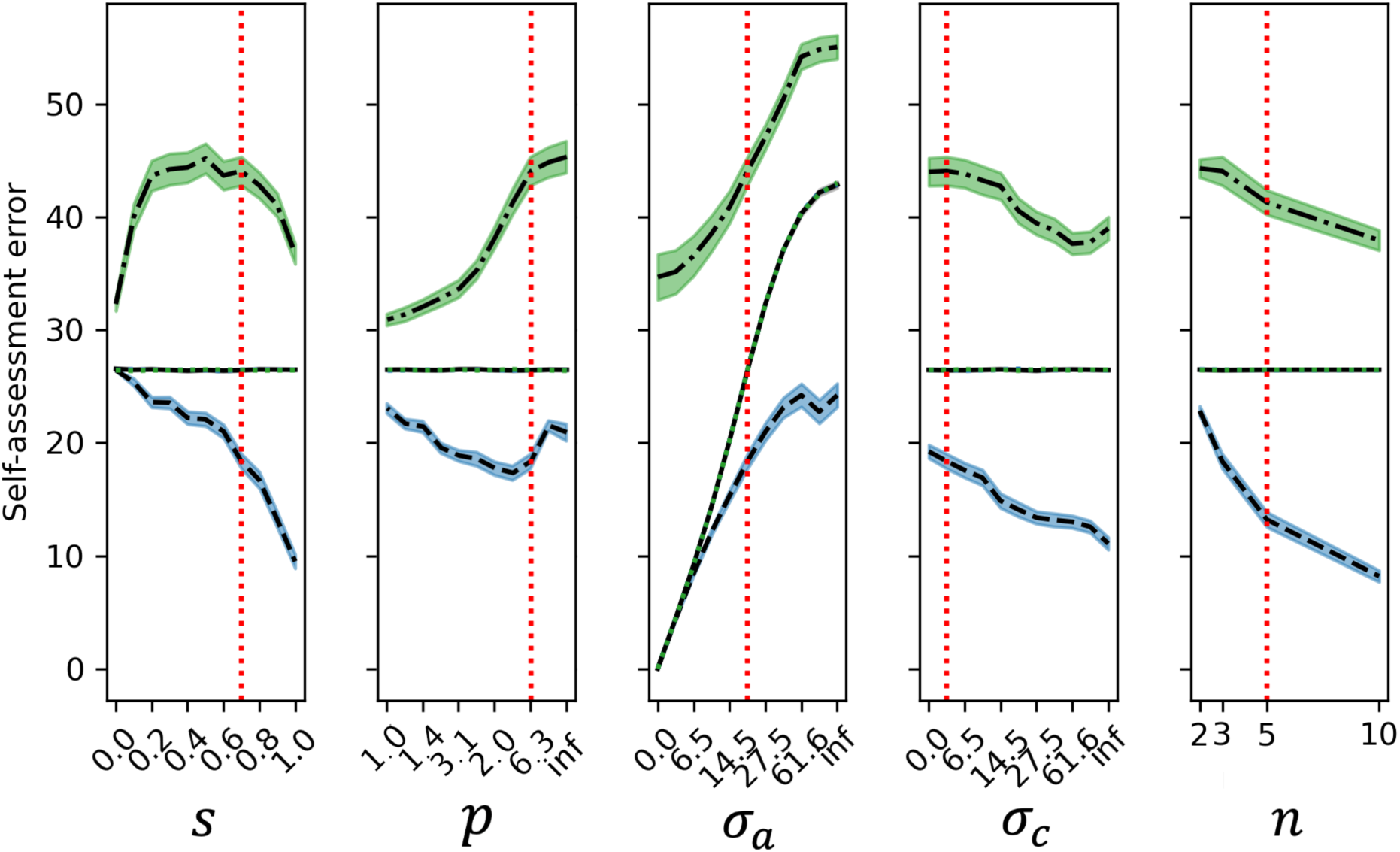
Estimate error plotted over a range of parameters. Groups of n agents are allowed to interact for 10 rounds, and lines represent the mean error (+/- SEM) over 100 iterations. groups of n agents competed over 10 rounds. Lines and shading represent the mean contest intensity (+/- SEM) over 100 iterations for groups using Bayesian updating (dashed, blue shading), linear updating (dot-dashed, green shading), or no updating (solid, grey shading). The dotted lines show the first-round error for agents using Bayesian updating (blue) or linear updating (green). In this case, first-round error precisely matches no-updating (black).

Estimate error is calculated based on squared difference between actual size and estimated size +/- prior standard deviation, as shown in equation (B2). The vertical red line shows the default parameters used for the main results (which are held constant in each panel while varying the relevant parameter)

Similarly, contest intensity, measured as the effort of the lower-investing agent, decreases over the course of the simulation. Although Bayesian updating reduces contest intensity, the effect of the Bayesian updating is much smaller than the effect of the simulation parameters themselves. For example, contest cost is very low when *l* is high, (representing strong bias against the ‘underdogs’), regardless of whether agents perform Bayesian updating. Where size is unimportant, or where initial assessment is high, we do not observe the emergence of low-cost behaviour that is typical of many empirical dominance hierarchies, suggesting that Bayesian updating provides the greatest benefits where naive self-assessment in poor and where size and effort both contribute to contest outcomes. Although mean intensity is lower for linear updating than for Bayesian updating, because linear updating leads to high self-assessment error, contests between two above-average individuals tend to be very intense, such that for linear updating 7% of contests are high intensity (i.e., loser effort > 0.8) vs 0.4% for Bayesian updating.

**Figure B9.**
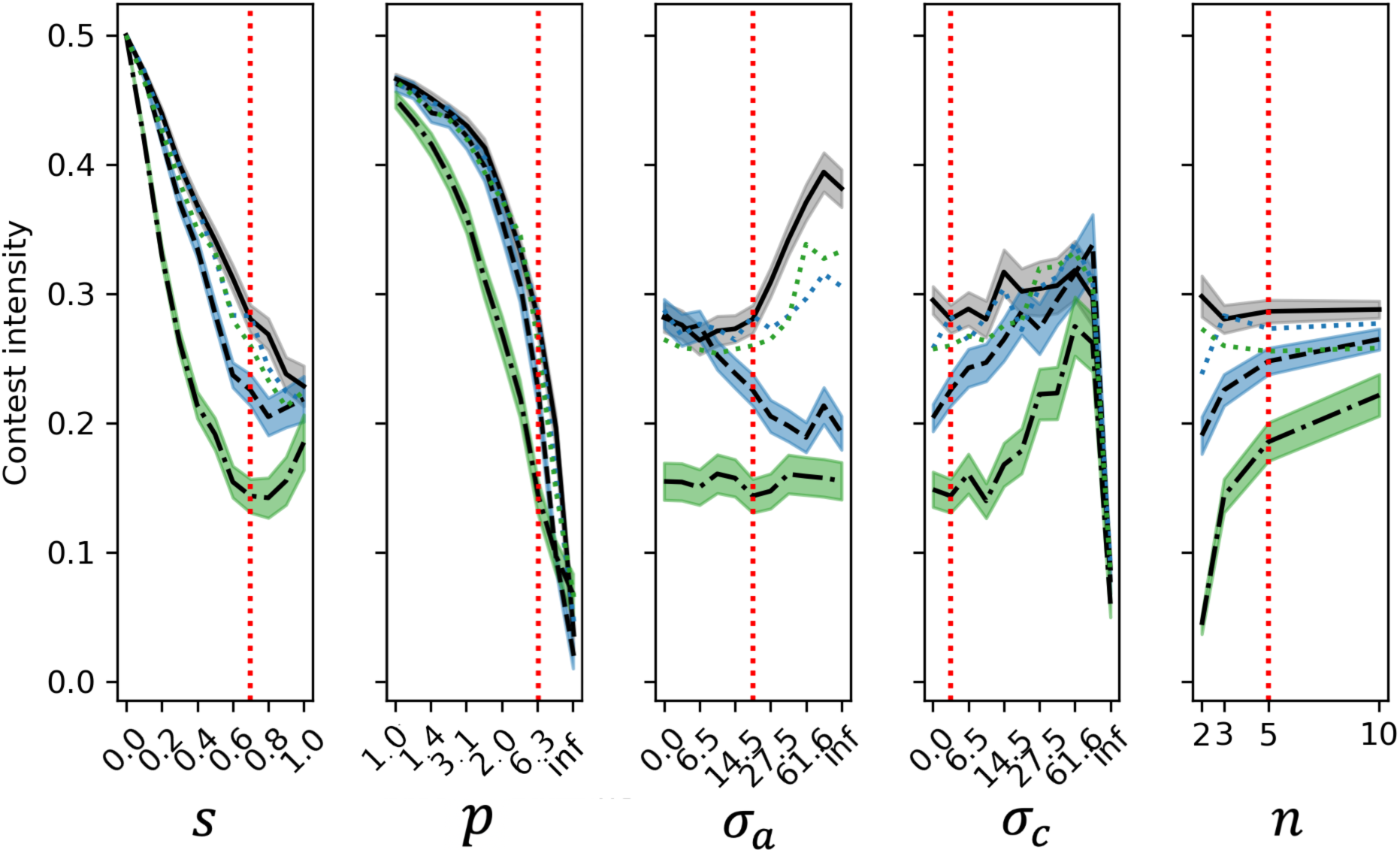
Contest intensity plotted over a range of parameters. As in figure B8, groups of n agents competed over 10 rounds. Lines and shading represent the mean contest intensity (+/- SEM) over 100 iterations for groups using Bayesian updating (dashed, blue shading), linear updating (dot-dashed, green shading), or no updating (solid, grey shading). The dotted lines show the mean first-round contest intensity for agents using Bayesian updating (blue) or linear updating (green). Contest cost is equal to the effort of the lower-investing individual. The vertical red line shows the default parameters used for the main results (which are held constant while other parameters vary).

Linearity and stability are hallmarks of social dominance hierarchies, and we find that under most conditions where contests are biased against the underdog (i.e., *p* > 6), Bayesian updating leads to a slight increase in both linearity (figure B10) and stability (figure B11) compared to no-updating. To measure the effect of Bayesian updating on the linearity of dominance hierarchies, we calculate linearity for the first and last rounds using Appleby’s approach (Appleby, 1983), measuring the proportion of triads which are transitive (if A dominates B, and B dominates C, A dominates C). Stability is calculated by composing per-round dominance hierarchies (a single round represents a series of contests in which every agent faces every other agent). We then bin the first 3 rounds, and the last 3 rounds, within which we measure the proportion of pairwise dominance relationships which are consistent for all rounds within that bin (for two agents, if A dominates B in round 1, round 2, and round 3, that is consistent, otherwise, it is inconsistent).

**Figure B10.**
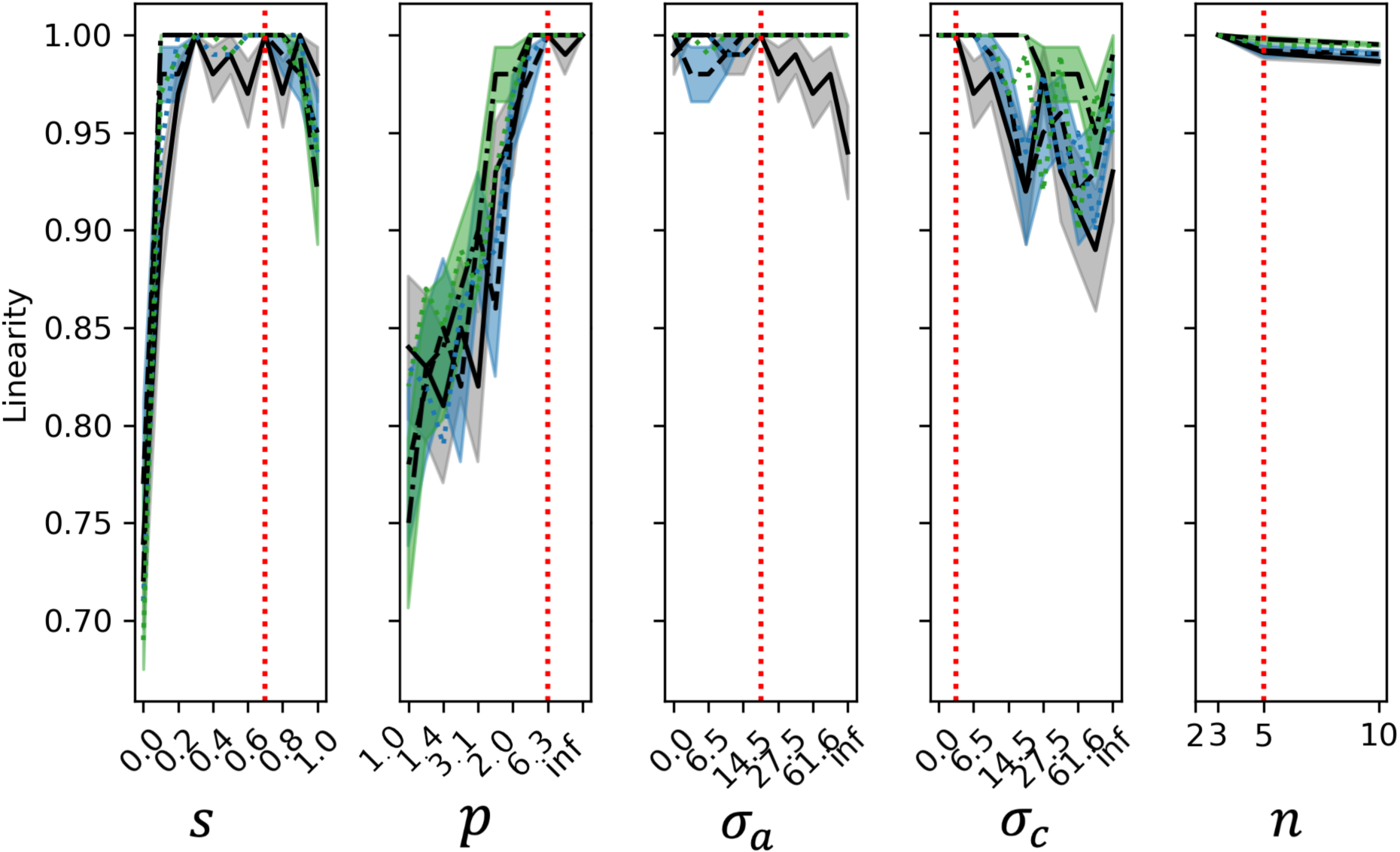
Network linearity plotted over a range of parameters. As in figure B8, groups of *n* agents compete over 10 rounds. Lines represent the mean, final-round linearity (+/- SEM) over 100 iterations. Lines and shading represent the mean contest intensity (+/- SEM) over 100 iterations for groups using Bayesian updating (dashed, blue shading), linear updating (dot-dashed, green shading), or no updating (solid, grey shading). The dotted lines show the mean first-round linearity for agents using Bayesian updating (blue) or linear updating (green). Linearity is measured as the proportion of triads which are transitive, following Appleby’s approach. The vertical red line shows the default parameters used for the main results (which are held constant for other plots while other parameters vary).

**Figure B11.**
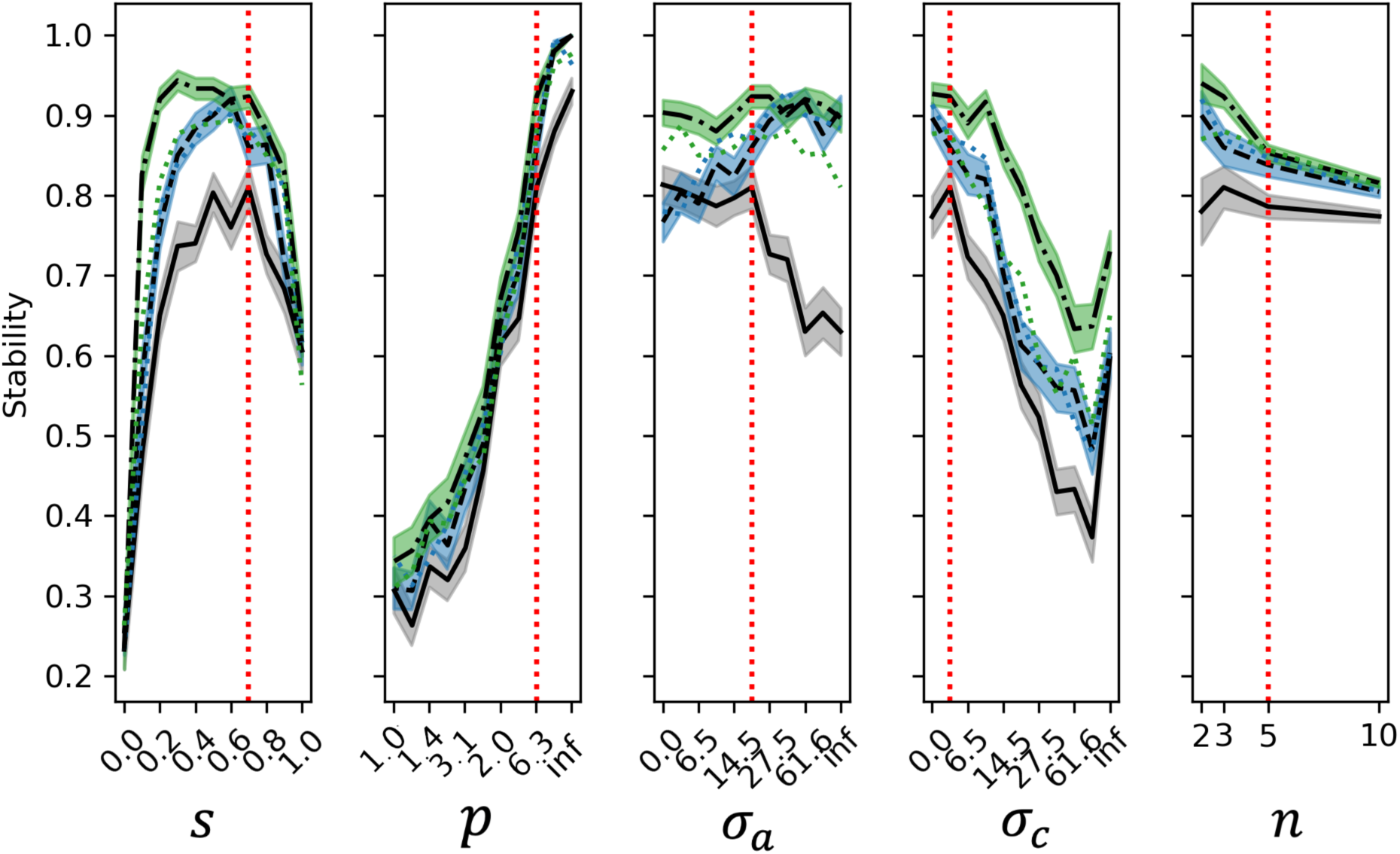
Network stability plotted over a range of parameters. As in figure B8, groups of *n* agents competed over 10 rounds. Lines and shading represent the mean contest intensity (+/- SEM) over 100 iterations for groups using Bayesian updating (dashed, blue shading), linear updating (dot-dashed, green shading), or no updating (solid, grey shading). The dotted lines show the mean first-bin stability for agents using Bayesian updating (blue) or linear updating (green). Stability is measured as the proportion of pairwise dominance relationships which are consistent across all three rounds. The vertical red line shows the default parameters used for the main results (which are held constant while other parameters vary).

As shown in figures B10 and B11, Bayesian updating produces some improvement in linearity and stability (compared to no updating, solid black line, or the first round, dotted blue line), particularly where initial self-assessment error is high. However, this depends on the existence of intrinsic differences in the individuals. To test whether Bayesian updating can produce linear hierarchies in identically sized individuals, we repeated the above simulation using groups of (*n* = 5) individuals of size, *x* = 50, and in general, under Bayesian updating, same-sized agents do not form linear hierarchies (figure B12*A*). While there is a temporary period of stratification, where early wins vs losses are important (figure B12*D*), with additional contests, agents recognize that they have equal likelihood of winning, and invest accordingly, leading to chaotic dominance networks. In other words, Bayesian updating, in isolation, does not generate strongly linear hierarchies, which is in contrast with prior models of winner-loser effects (e.g., Bonabeau et al. 1996; Dugatkin 1997). Linear updating (which is based directly on Dugatkin’s model), does produce linear hierarchies, as there is more opportunity for runaway winner-effects.

So far, this seems to suggest that Bayesian updating is a poor method for establishing linearity, but this is in this unique context of identically sized individuals. If we allow agent size (i.e., their ability to win contests) to vary as a function of wins and losses as in these previous models, our model recapitulates the self-organization described in empirical systems. To allow direct feedback on size following a win or loss, we set size post-contest size to be

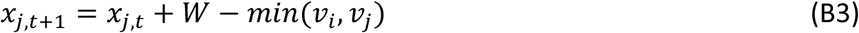

Where *W* = 1 for a win and *W* = 0 for a loss. Thus, contest cost is equal to the lower invested effort, *v*_*min*_, and contest benefit is 1, regardless of the circumstances. Simulating 5 individuals with identical starting size and priors, we see in figure B12*B* that after a few rounds of contests, agents have clear differences in both their sizes and estimates. Although we did not modify Bayesian updating to account for this change in size, we also see that agents’ estimates more-or-less match their actual (shifting) intrinsic ability.

Interestingly, for this feedback on intrinsic ability, Bayesian updating amplified the self-organization of networks, compared to changes in *size* alone (figure B12*E*). These feedback+Bayesian updating networks were as strongly linear as hierarchies where agent size was set to be highly different from the start (figure B12*C,F*). In short, under Bayesian updating the self-organization of dominance hierarchies must come through differences in intrinsic ability—without this, similarly sized individuals do not form stable, linear dominance hierarchies—but Bayesian updating can reinforce this effect to produce networks that are more stable and linear than are observed through feedback on intrinsic ability alone.

**Figure B12.**
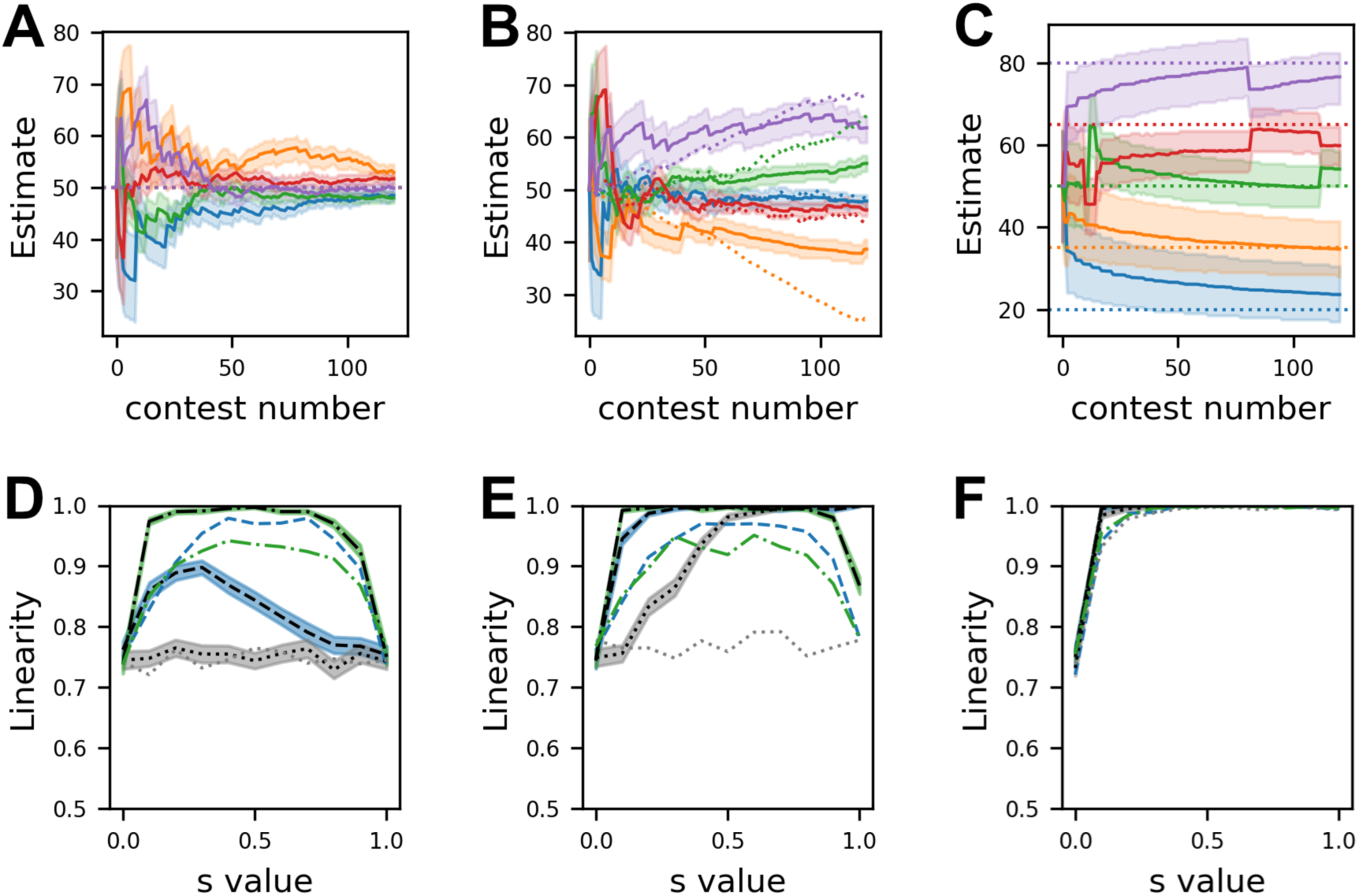
The linearity of dominance hierarchies is a function of intrinsic differences and updating method. **A-C** show the estimates of 5 agents using Bayesian updating during single simulations, where (**A**) has neither intrinsic differences in size, nor feedback on size; (**B**) has no initial intrinsic differences but direct feedback from contests with winners increasing size by 1, while losers decrease by 1 unit; (**C**) begins intrinsic differences in size, but without intrinsic feedback. **D-F** show the same scenarios as their corresponding panels above, repeated over 100 iterations. Blue, dashed lines show Bayesian updating; green, dash-dot lines show linear updating, while black dotted lines show no-updating. Lines without shading represent first-round linearity for each update method. Shading shows +/- one SEM.

## Summary of Part 1

So far, we have explored the observations in Part 1 of the main-body—namely that 1) Bayesian updating produces winner-loser effects; 2) that Bayesian updating has a modest bias towards the loser-effect; that under Bayesian-updating, winner-loser effects 3) show a recency bias and 4) attenuate quickly in the face of new information; and that 5) Bayesian updating helps to stabilize dominance hierarchies. In all of these cases, these qualitative phenomena were broadly observed throughout the parameter space, although it is important to note the edge-cases mentioned, and that winner effects generally disappear where effort is not a factor, or where contests are strongly dependent on chance.

### Part 2: The sensitivity of empirical predictions

Part 2 lays out specific predictions of the model that could be used to test for Bayesian updating in natural systems and distinguish it from alternative updating approaches. Here, we explore the sensitivity of these predictions across parameters.

**Table B1.** Summary of predictions for various models.

| Prediction | Continuous | Assessment: | Direct: Linear<sup>†</sup> | Binary Bayes | Reinforcement Learning |
| --- | --- | --- | --- | --- | --- |
|  | Bayes (ours) | Linear (ours) | (Dugatkin) | (Fawcett, Johnstone) | (Leimar) |
| Winner/Loser Effect | yes | yes | yes | yes | yes |
| Loser Effect Bias | yes | yes | **no** | yes (because of<br>max aggression) | yes |
| Winner/Loser effect bias<br>dependent on Strategy (i.e.<br>Aggression) | yes | yes | **n/a** | yes* | yes |
| Opponent Size determines<br>effect strength | yes | **no** | **no** | **no\*** | yes |
| Contest intensity determines<br>effect strength | yes | **no** | **no** | yes | yes |
| Recency Bias | yes | yes | **no\*** | **no\*** | yes* |
| Certainty Effect | yes | **mixed** | **no\*** | **mixed (loser effect**<br>**down, winner effect**<br>**up)** | **no\*** |
| Persistence depends on<br>encounter intensity | yes | **no** | **no** | **no\* (but only Hawk-**<br>**Hawk have effects)** | yes* |
| Linear Hierarchies | yes | yes | yes | **no** | yes |
| Linear Hierarchies w/o<br>intrinsic differences | no | **yes** | **yes** | no | **yes (due to bystander**<br>**learning)** |
| Juveniles/Naive hyper<br>aggressive | no | no | no | **yes** | **yes** |
| Losers become stuck | no | **yes** | **yes** | **yes** | No* |

\* Based on our understanding of the model; these predictions are not addressed in the corresponding paper
† As winner-loser effects are specified, for Dugatkin, all but the last 4 “predictions” are assumptions of the model.

#### 2.1 Individual experience influences winner-loser effects across a range of parameters

As we show in figure 4A, winner effects are subject to the amount experience before the treatment contest, and the same is true for loser effects (figure B15). We can measure this “certainty effect” by comparing naïve and experienced (in this case 10 previous contest) agents. As mentioned in the main results, when prior experience is random the strength of winner-loser effects also decreases under linear updating with experience as individual estimates stratify to extremes such that above average individuals assume they are extremely large and (almost) always win their assay contest, while losers always lose, regardless of whether they win or lose their treatment contest.

**Figure B13.**
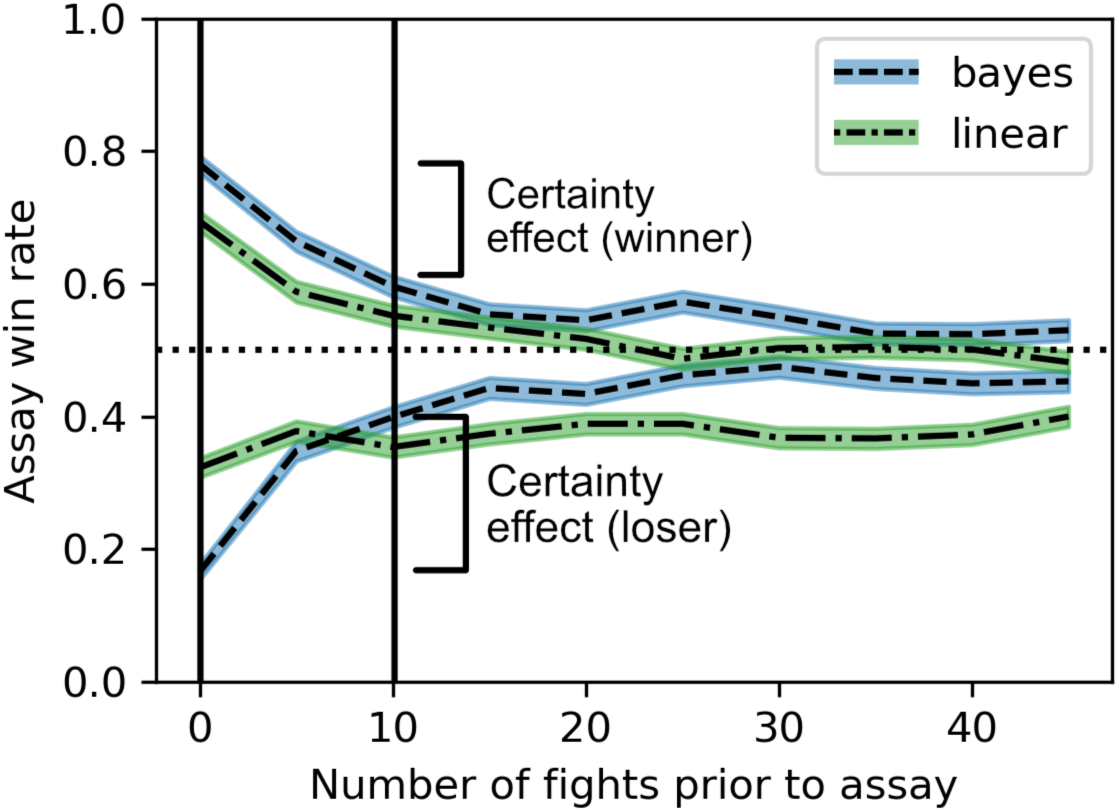
The certainty effect is measured as the difference between the probability of winning a size-matched ‘assay’ contest after a ‘treatment’ contest for a naïve individual vs an experienced individual. In this case, the only difference is the number of contests agents have experienced prior to their ‘treatment’ contest. For both focal winners and focal losers, Bayesian updating results in a weakened winner/loser effect as experience (and thus prior confidence) increases. In contrast to the main body, under this paradigm, where prior experience consists of uncontrolled fights against random opponents, linear updating also shows a clear weakening effect of experience for winners (and a slight experience-based decrease for losers).

To test how the certainty effect of Bayesian updating varies across parameters, we repeated this simulation, averaging across 1000 iterations, comparing the naïve and experienced (n=10 fights) agents—across all contest parameters—to observe where this certainty effect occurs. This phenomenon is observable under roughly the same parameters as winner-loser effects in figure B1, but with added noise introduced by the randomness of pre-treatment opponents.

**Figure B14.**
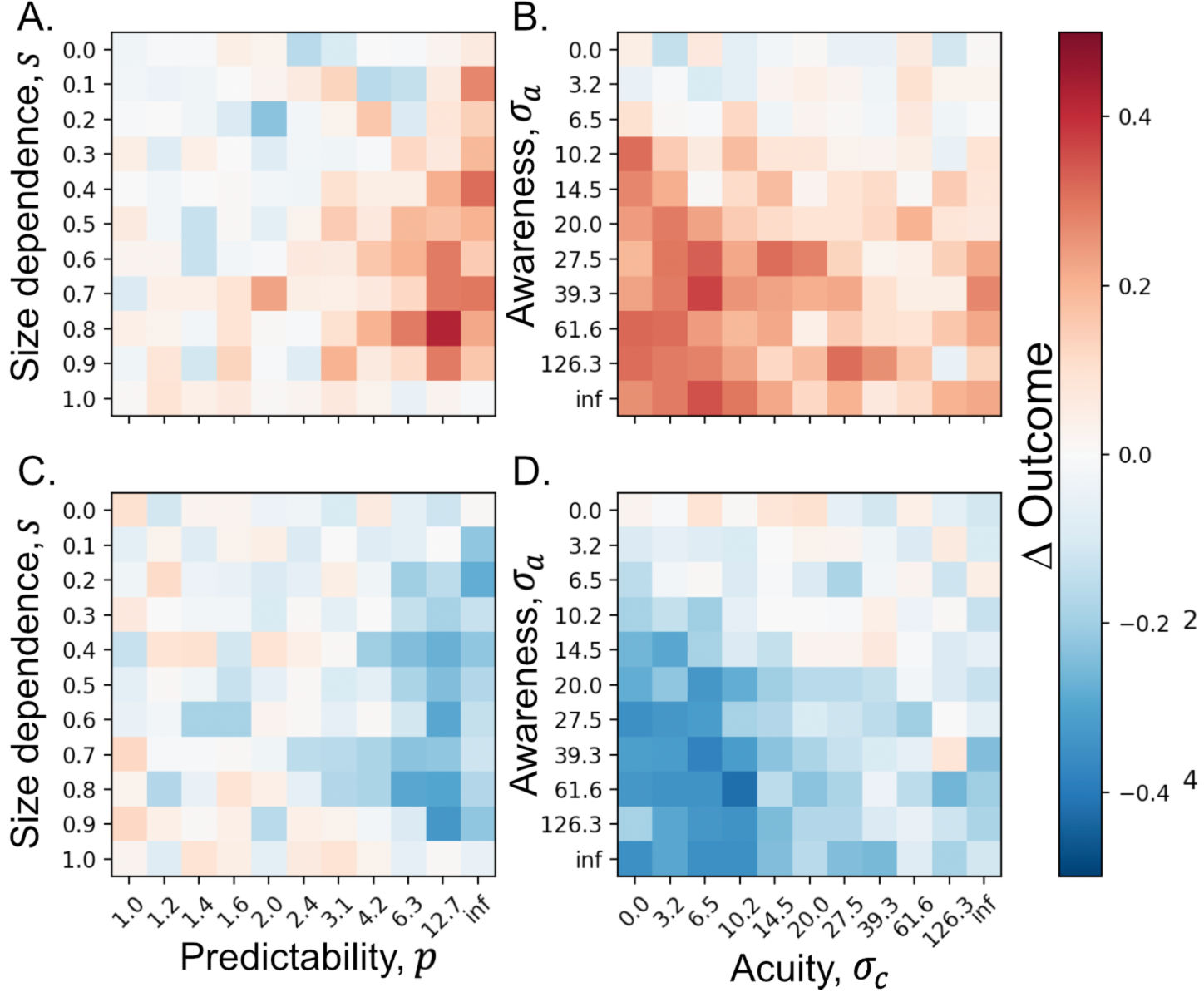
The certainty effect is broadly observed wherever the winner-loser effect exists. The top row shows the strength of the certainty effect (measured as the difference in assay-contest win probability between naïve and experienced individuals) for winners over contest parameters *s* and *p* (left column) and assessment parameters σ_*a*_ and σ_*c*_ (right column), while the bottom row shows the same for focal losers.

#### 2.2 Opponent size impacts winner-loser effects across a range of parameters

Similarly, in figure 4B, we observe a discrepancy effect for winners in Bayesian updating, but not other updating paradigms. As with the certainty effect, we repeat this simulation to test its sensitivity to different parameters, this time including both winners and losers, facing large or small treatment opponents (size 40,60 respectively, as shown on figure B15). The discrepancy effect is measured as the difference in win rate between focal agents who won (or lost) against a “treatment” opponent of size 40 and those that won (or lost) against one of size 60.

**Figure B15.**
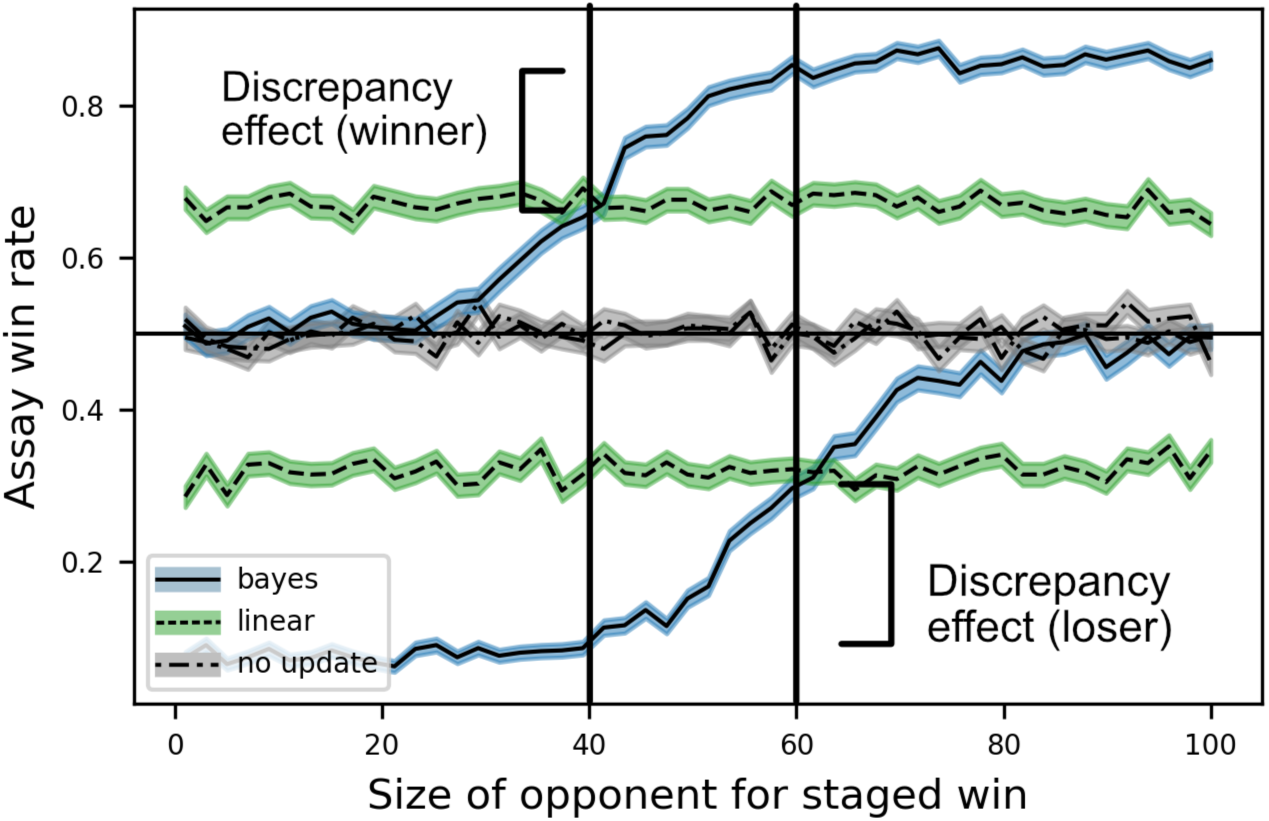
The discrepancy effect is measured as the difference between the winner/loser effect for a larger “treatment” opponent compared a smaller ‘treatment’ opponent. These lines show the probability of winning against a second, size-matched ‘assay’ opponent, for three different updating strategies. This phenomenon requires that opponent size is factored into the strength of the winner-loser effect, as in the Bayesian updating approach in blue. Note that for both winners and losers, competing against a larger “treatment” opponent result in a higher probability of winning the “assay” contest.

**Figure B16.**
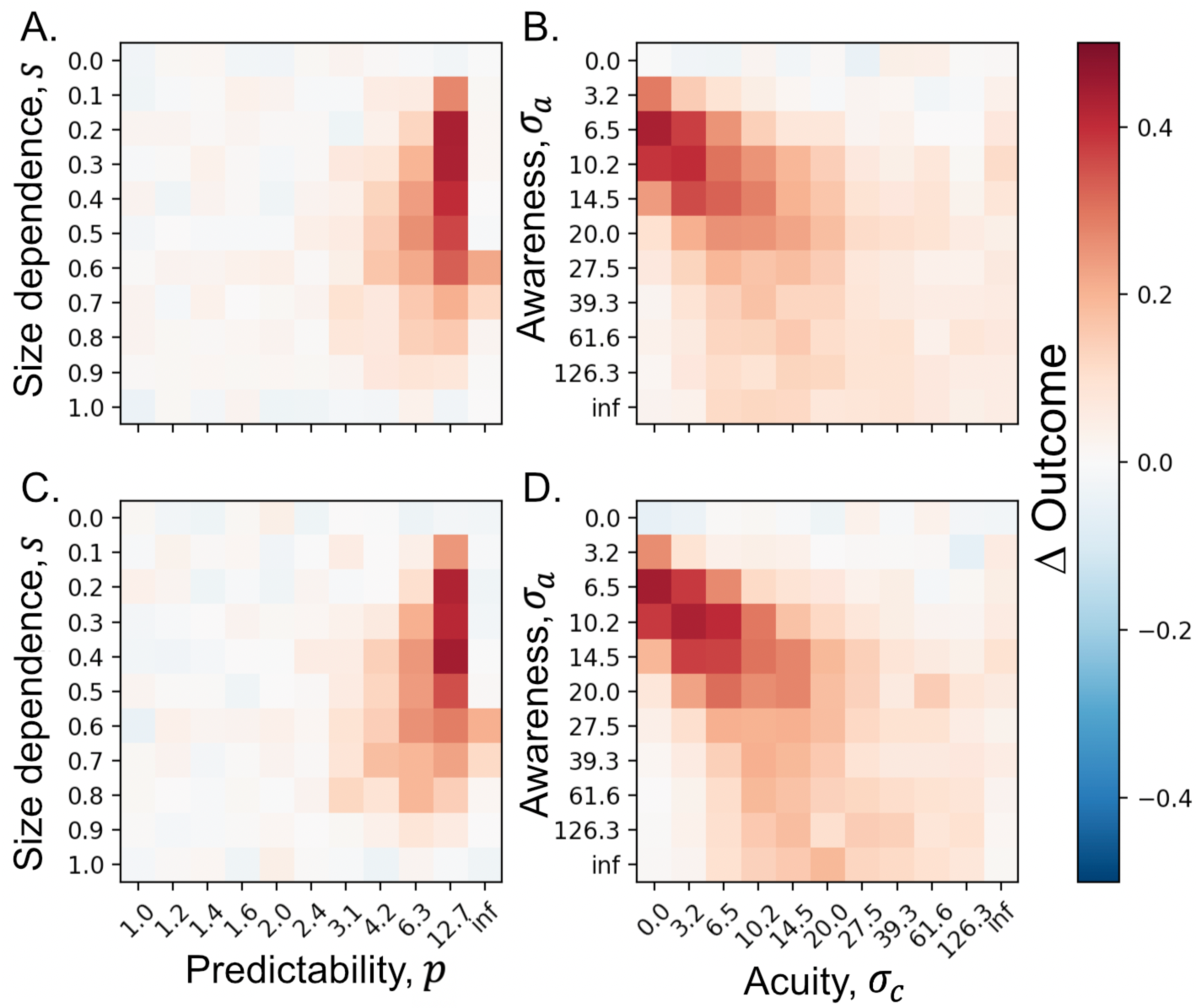
The top row shows the discrepancy effect, measured as the difference in the proportion of focal agents winning against size-matched assay opponents, after a win against a smaller or larger opponent. The bottom row shows the same information, but for focal losers. The discrepancy effect is broadly observed where contests are biased against the underdog but drops off quickly as *l* decreases. Like the winner-loser effect in general (see figure *B*1), the discrepancy effect is largely dependent on contests being strongly biased against the underdog (*l* > 4). Where the relationship between relative wager and winning is closer to linear, this effect is no longer observed.

Across parameters, we observe a discrepancy effect wherever size and effort can strongly predict outcome (figure B16). In other words, the discrepancy effect is only observed if both size and effort are factors, and where the value of *p* is quite large. On the other hand, where self-assessment error is 0, or opponent assessment error is lower than self-assessment error, we do not observe a clear discrepancy effect (figure B16, second column). This assessment constraint occurs because where opponent assessment is high, the opponent size-estimate in the size-matched ‘assay’ contest was largely random, meaning the focal agent effort was also very noisy. This behavioural noise masks weak winner/loser effects. Note that while we have demonstrated discrepancy effect using opponent size, which linear updating does not take into account, the discrepancy effect follows directly from Bayesian updating using the likelihood function, and the discrepancy effect should occur for any feature impacting the probability of winning (e.g., opponent effort, focal agent effort).

#### 2.3 Informed priors link size and effort wherever both influence contest outcome

Often when behaviourists test whether animals are using Bayesian strategies, they are asking specifically whether animals have “prior opinions” about the general state of the world which are then applied to each new context, for example the quality of a novel potential mate or some new foraging patch (reviewed in McNamara et al. 2006). In this sense, the question is not so much about how individuals update their self-assessment during the task, but whether animals come into the task having informed prior estimates which reflect their expected state of the world, either based on their own past experiences or evolved over generations (Stamps & Krishnan, 2014). With this in mind, we designed simulated experiments that would allow us to empirically distinguish whether individuals have informed priors, and how these priors form.

To explore this question, we generate agents that begin with either informative or uninformative priors. Priors are modelled as a probability distribution of size. For uninformative priors, we use either 1) a uniform distribution (“uninformed”) or 2) a normal distribution centred on a uniformly random variable (“misinformed”). The informative priors are similarly divided into 3) a normal distribution centred on the agent’s approximate size (“self-informed”) or 4) a distribution that reflects the distribution of nearby agents (“peer-informed”, for which we set peers to be either all large or all small). While it seems unlikely any animal would have a truly uniform distribution (i.e., without any lived or evolved sense of their respective size), there is known individual variation in behaviour that is independent of ability (Niemelä & Dingemanse, 2018), i.e., a “misinformed” prior. In the “informed priors”, individuals could have gained information either from some prior experience or internal sensing or based their size estimates on their observations of the size distribution of conspecifics (i.e. peers).

We conduct simulated experiments in which we measure contest intensity—which in an empirical context could be quantified as the time spent in aggressive behaviours (Francis, 1990; Schlinger et al., 1987)—among individuals using these different prior estimation methods. Each simulated agent is assayed 3 times, each time against a new, naïve opponent of size *x*_j_ = 50. Because we simulate 3 trials for each agent, we can measure both consistent individual differences and context-dependent variation aggressive behaviour (i.e., *effort*). As we are focusing on the influence of differences in prior information, we prevent agents from updating their self-assessments after each contest, though for the most part these results are qualitatively similar when updating is allowed (see figure B18).

In these simulated experiments, we find key differences in aggressive behaviour which successfully characterizes agents based on the source of their prior (figure B17). Specifically, in groups of agents using either self-informed or mis-informed priors, there are consistent individual differences in contest effort (i.e., across the repeated trials, some individuals consistently exhibit greater levels of ‘aggression’, or effort, compared to others). Both large peer-informed and small peer-informed agents do not show these consistent individual differences; that is, there are no average differences in aggressive behaviour among agents within these groups. Importantly, only self-informed agents displayed a correlation between individual variation in effort and size: agents that think they are bigger are consistently more aggressive than agents that think they are smaller, and this variation correlates with actual agent size, but only if agent priors are informed by their own size. By controlling for peer group and characterizing consistent individual variation in effort (and size), it is possible to identify whether agents form prior self-assessments and, if so, what information they use to inform these estimates.

**Figure B17.**
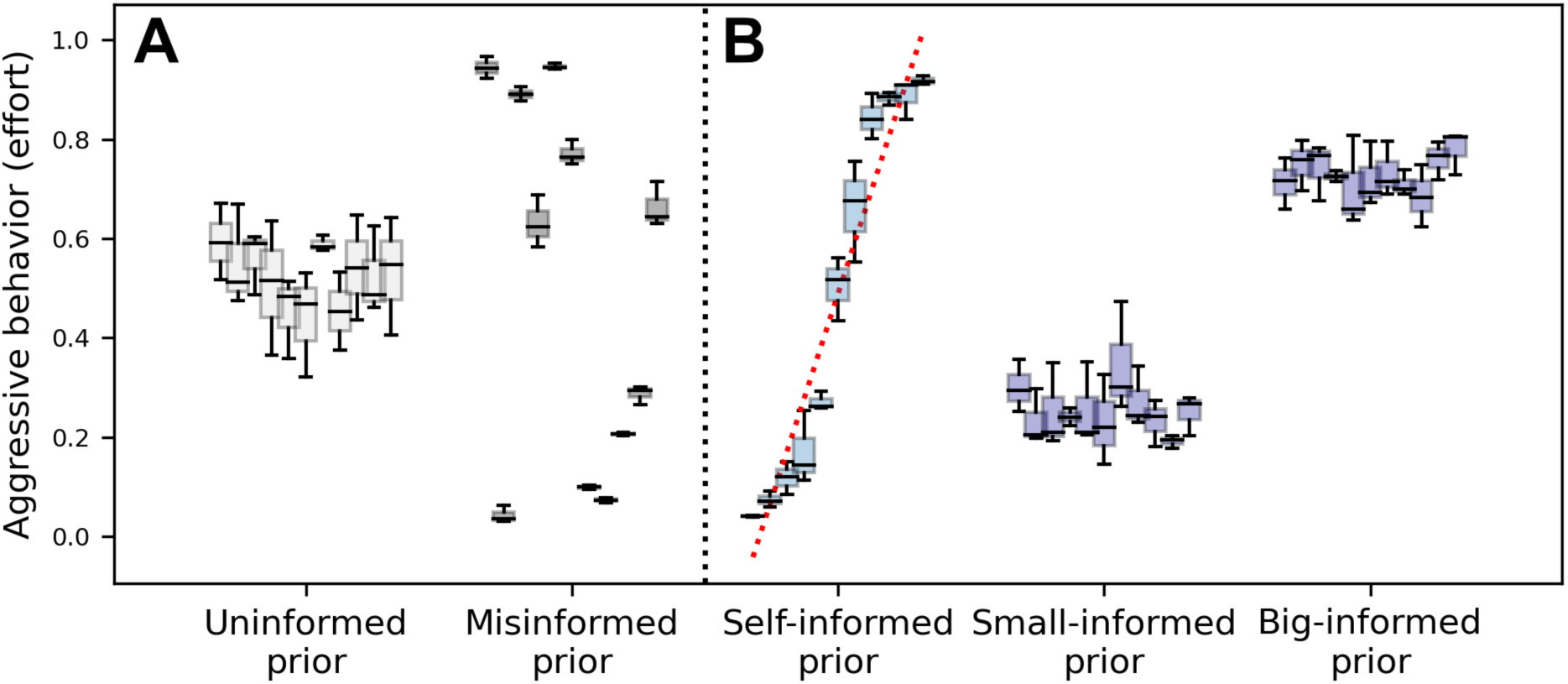
Aggressive behaviour (i.e., effort, *v*_*i*_) for 5 different types of simulated agents, each using a different starting prior. Each boxplot represents one agent’s behaviour across three trials; within each group, boxes are arranged from left to right according to agent size. **A,** the left-most two groups (light and dark grey) have non-informative priors. **B**, for the three groups on the right, individuals have “informed” priors. While significant individual differences in aggressive behaviour occur for both “misinformed” and “self-informed”, only self-informed priors lead to a correlation between size and effort (red dotted line).

To test the generality of this result, we repeat the simulation shown in figure B17 for every point in the parameter space, averaging the effect size across 1000 iterations (figure B18) and then tested to see whether we still see a positive correlation between size and effort in our “self-informed” agents. We observe a positive effect in all conditions, except in unlikely edge cases (e.g., where agent effort does not influence outcome).

**Figure B18.**
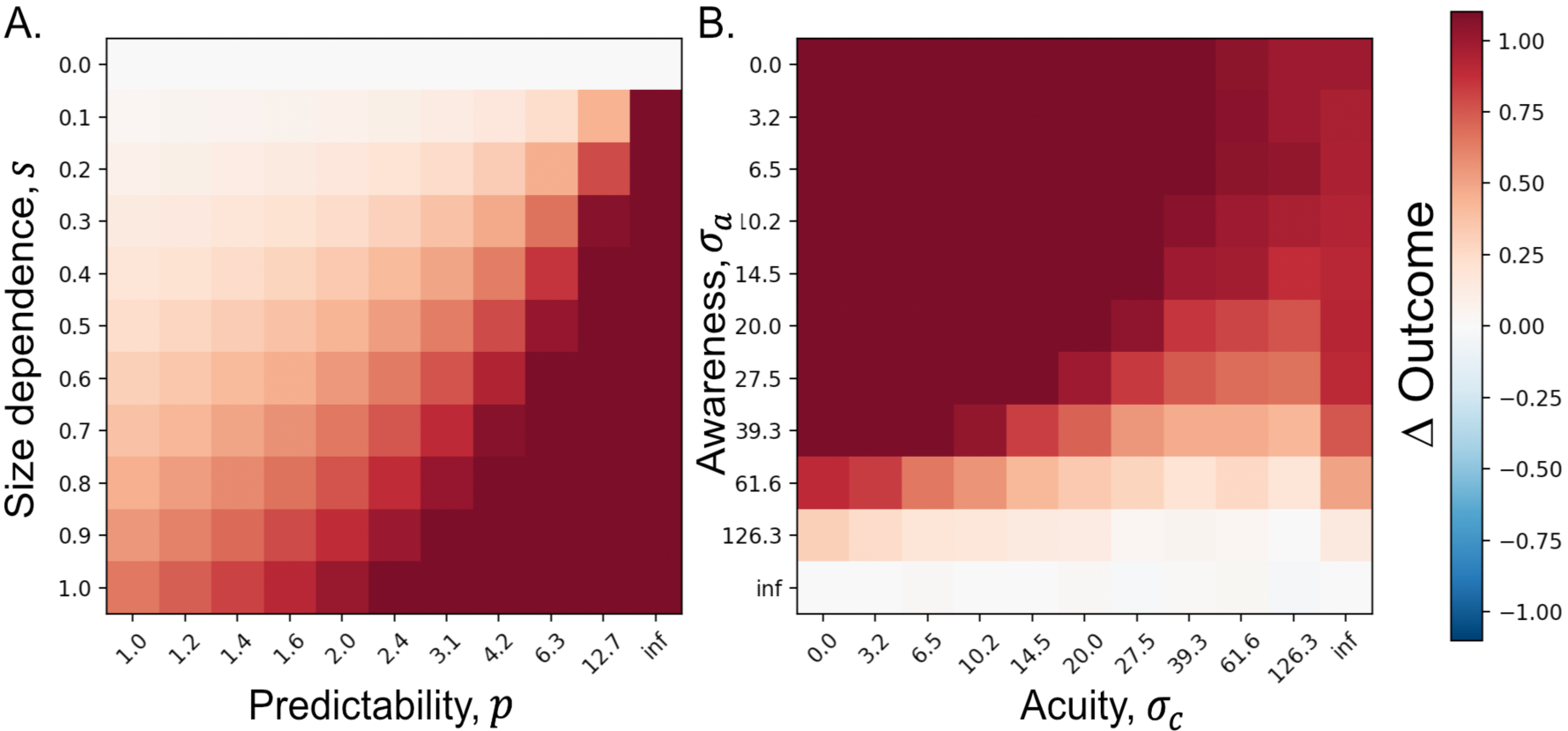
Size and effort correlates for self-informed agents across a range of parameter values. Here, the heatmap shows the strength of the effect (i.e., the slope of normalized size, 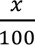, and effort, simulated as in figure 5, averaged across 1000 iterations. For greater resolution over the range of interest, the heatmap is clipped at 1.1, such that any slope higher than that is normalized. Over all values where size contributes, there is a significant effect between size and effort.

It is also important to note that while this effect is a consistent prediction of self-informed assessment; to measure this effect, it is necessary to isolate “effort” from possible confounds like energy budget or motivation. In our model, we typically think of effort, *v*_*i*_, as the amount of time an individual animal is willing to stay in the contest, but this will often be linked to their available energy, which is frequently a function of individual size. This experiment would be most useful where *size* referred to something more specific, such as weapon size (as in figure A1), so that experimenters could control for intrinsic differences in the ability to persist in a fight. Alternatively, *effort* could represent the latency to engage in a contest, or some other non-energy-based metric. Obviously, applying theoretical predictions to empirical systems requires care and system-specific understanding to design experiments appropriately.

In the above results, we did not allow agents to update their self-assessment between contests, in order to focus on the prior itself, although this may or may not be possible under empirical conditions. To further compare the self-informed priors to other approaches, we allow agents with different starting priors to modify their self-assessment in between contests using Bayesian updating in order to see how this influences the observed correlations between size and effort. Allowing updating does not meaningfully influence self-informed priors, but it does introduce weak to moderate correlations between size and effort (figure B19) in all other groups, where otherwise this correlation is not observed (figure B17). The correlation is strongest in the “uninformed prior”, since the others have all formed moderately confident estimates and are thus resilient to additional experiences. In an empirical context this confound could be avoided by using the initial contest to define the correlation between size and effort and using the repeated contests only to characterize consistent individual variation.

**Figure B19.**
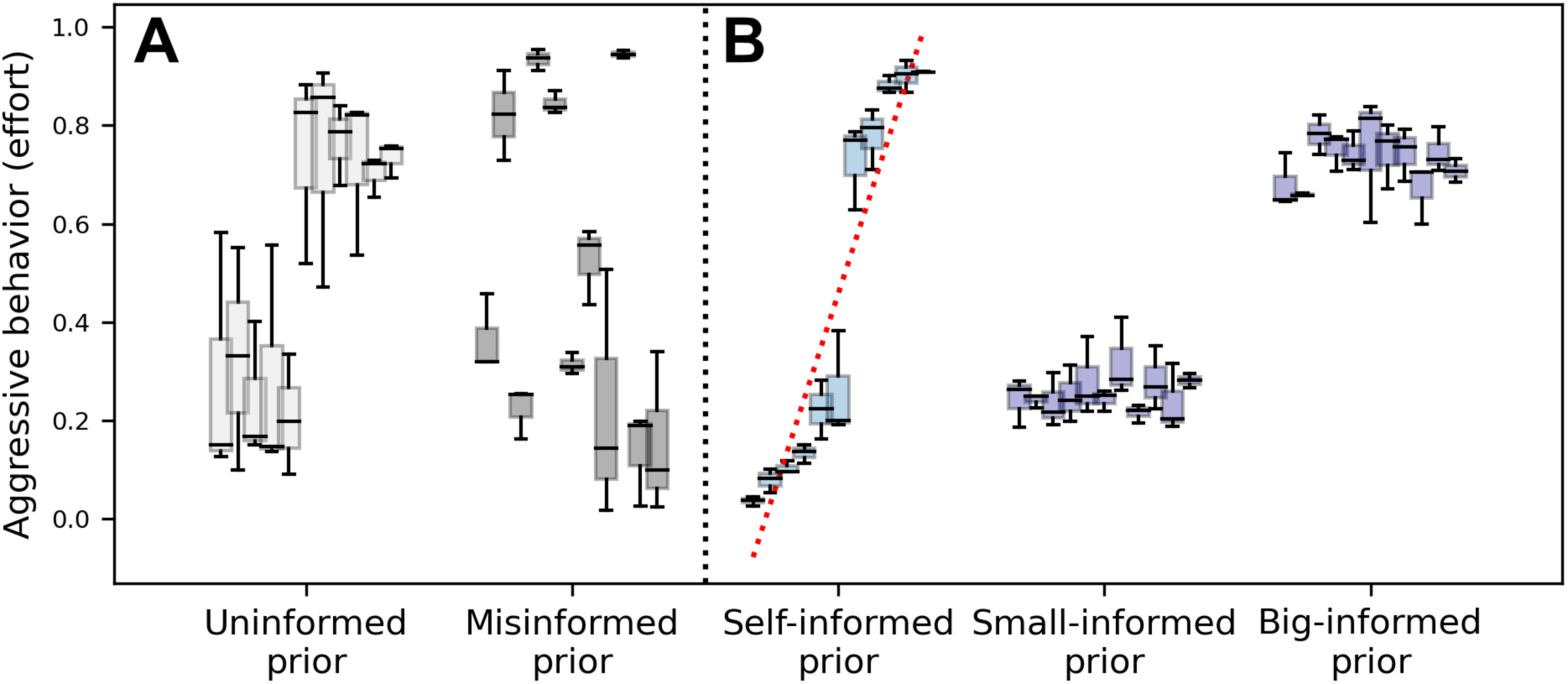
This modifies of the results figure B17 by allowing agents perform Bayesian updating after each repeated contest. Under these conditions, there are strong correlations for both uniform and self-informed priors, while random and externally-informed priors showed only weak correlations.

#### 3.1 A final note on effort

We show in figure B4 that the relative strength of winner versus loser effects depends on the effort strategy used. To check whether any other effects that we have observed are limited to our specific effort strategy, we replicate every analysis we have performed, using the default parameters but varying over the effort parameter h, which shifts the threshold for probability of winning above which individuals are willing to invest heavily in a contest, both for Bayesian updating, and linear updating. In general, while the specific values vary, the qualitative patterns are similar. It is notable that where h is close to 0, agents do not show a winner effect, and similarly the loser effect disappears when h is close to 1. As such, the recency effect (which as we measure it here depends on both winner and loser effects) only persists for moderate h values where winner and loser effects are present.

**Figure B20:**
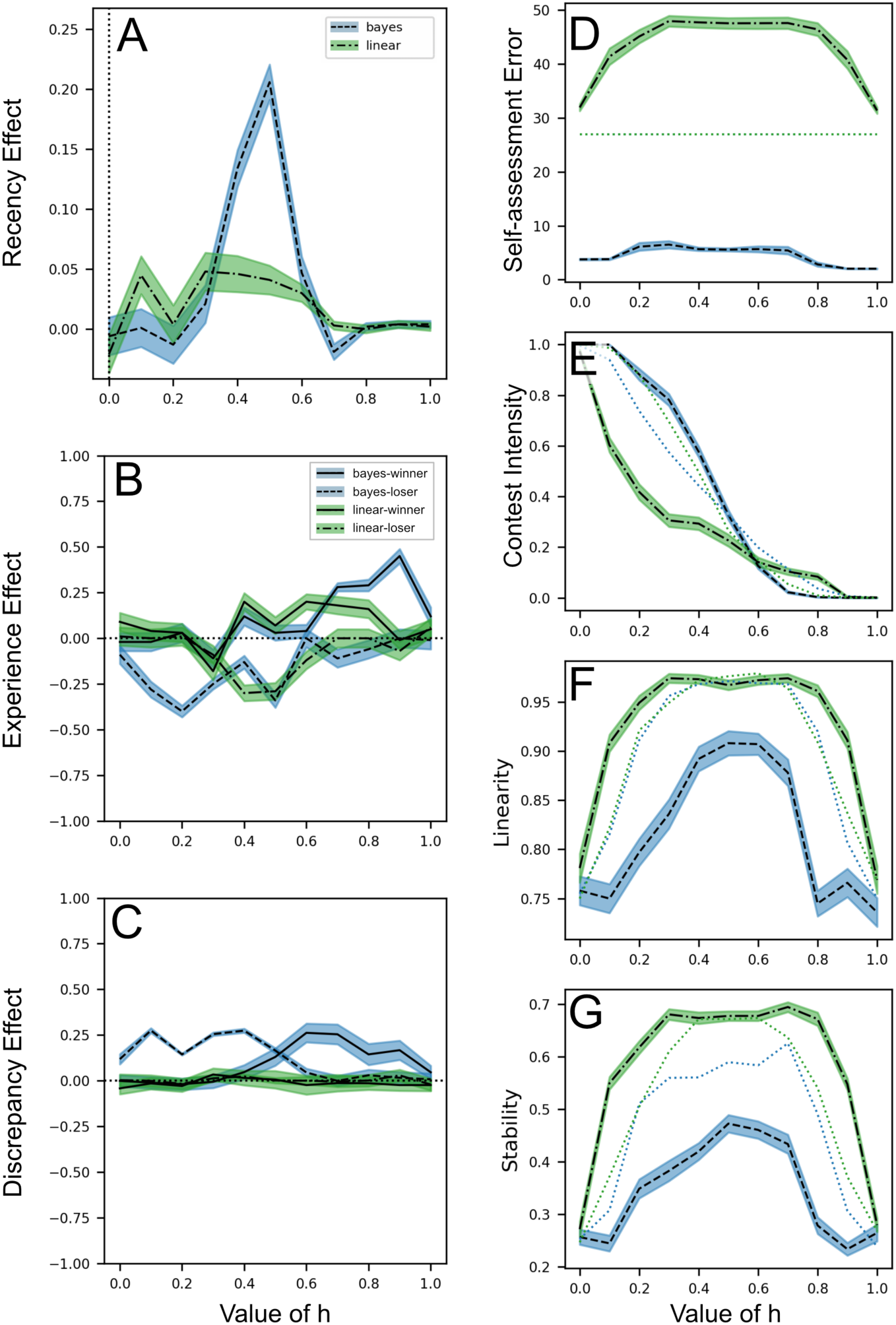
Exploration of the effect of effort strategy parameter on paper results. Each plot shows the impact of varying the parameter ℎ (see figure B3a), which could be conceived as setting the “hawk threshold” (the probability of winning above which should you invest highly). It also determines the strength of winner and loser effects, with high ℎ values corresponding to stronger winner effects, and low ℎ values corresponding to stronger loser effects (figure B3b). In each case, the measurements for the respective values are identical to their respective parameter explorations, i.e., the recency effect in panel A is measured as the difference in outcome between winning-then-losing, and losing-then-winning, as in figure B4e,f. C corresponds to figure B14, D corresponds to figure B16, and d-g correspond to figures B8-B11.

## Appendix C: Parameters and Equations

**Table C1.**
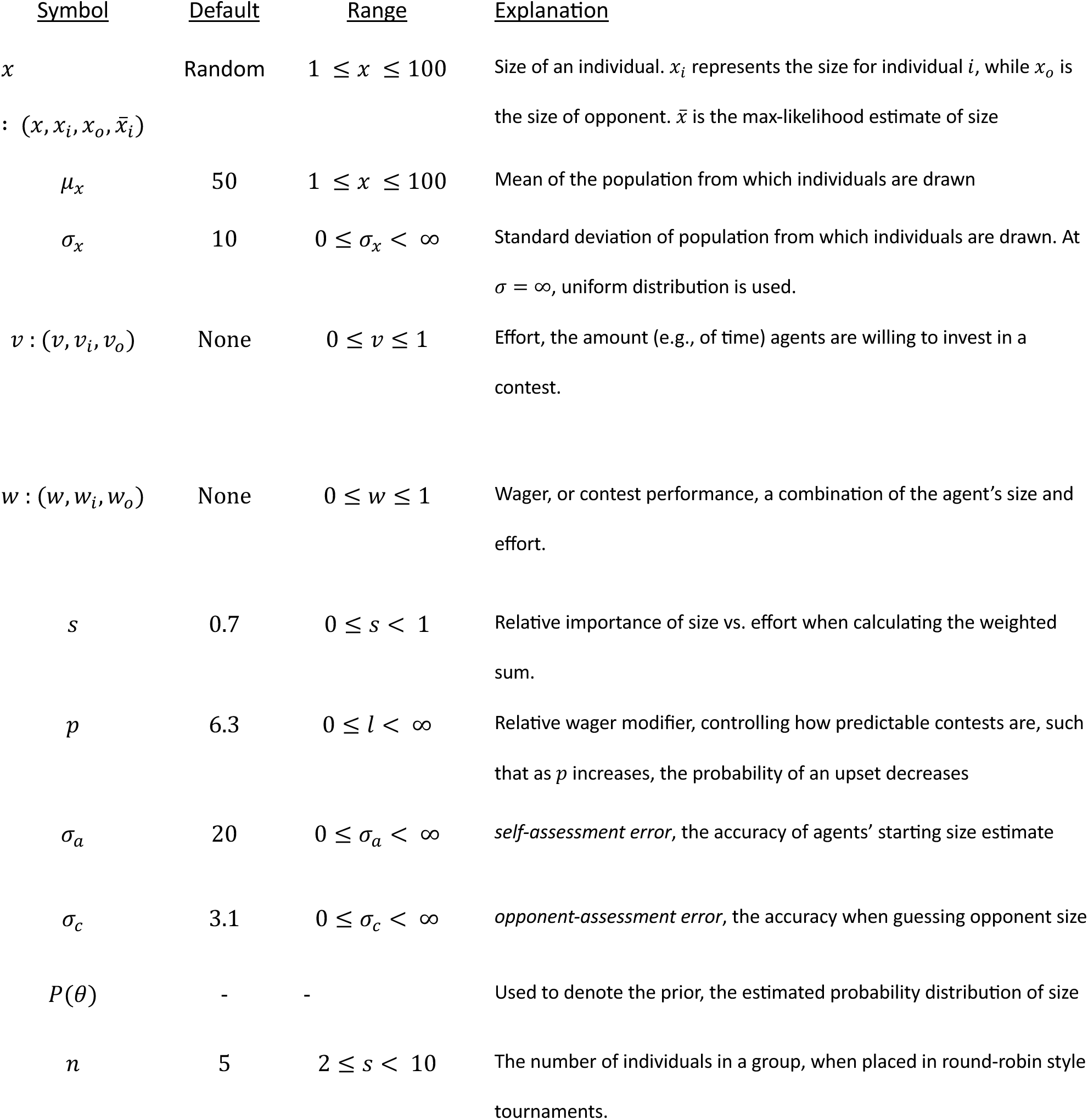
Overview of parameters and their default values sum.

| <u>Symbol</u> | <u>Default</u> | <u>Range</u> | <u>Explanation</u> |
| --- | --- | --- | --- |
| $x$<br>$: (x, x_i, x_o, \bar{x}_i)$ | Random | $1 \leq x \leq 100$ | Size of an individual. $x_i$ represents the size for individual $i$, while $x_o$ is the size of opponent. $\bar{x}$ is the max-likelihood estimate of size |
| $\mu_x$ | 50 | $1 \leq x \leq 100$ | Mean of the population from which individuals are drawn |
| $\sigma_x$ | 10 | $0 \leq \sigma_x < \infty$ | Standard deviation of population from which individuals are drawn. At $\sigma = \infty$, uniform distribution is used. |
| $v : (v, v_i, v_o)$ | None | $0 \leq v \leq 1$ | Effort, the amount (e.g., of time) agents are willing to invest in a contest. |
| $w : (w, w_i, w_o)$ | None | $0 \leq w \leq 1$ | Wager, or contest performance, a combination of the agent's size and effort. |
| $s$ | 0.7 | $0 \leq s < 1$ | Relative importance of size vs. effort when calculating the weighted sum. |
| $p$ | 6.3 | $0 \leq l < \infty$ | Relative wager modifier, controlling how predictable contests are, such that as $p$ increases, the probability of an upset decreases |
| $\sigma_a$ | 20 | $0 \leq \sigma_a < \infty$ | *self-assessment error*, the accuracy of agents' starting size estimate |
| $\sigma_c$ | 3.1 | $0 \leq \sigma_c < \infty$ | *opponent-assessment error*, the accuracy when guessing opponent size |
| $P(\theta)$ | - | - | Used to denote the prior, the estimated probability distribution of size |
| $n$ | 5 | $2 \leq s < 10$ | The number of individuals in a group, when placed in round-robin style tournaments. |

### Equations Sheet

Agent generation

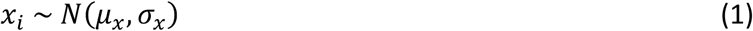

Agent estimate generation

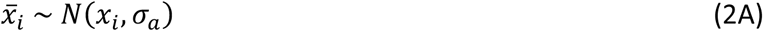

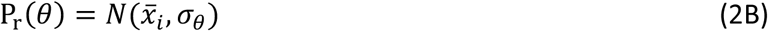

Investment strategy via opponent assessment

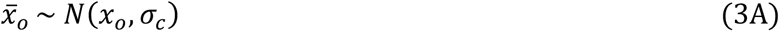

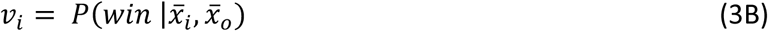

Calculating contest outcome as a function of relative wagers

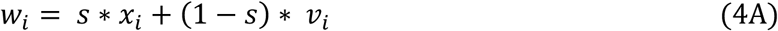

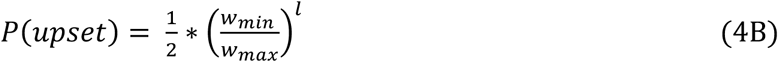

Linear updating strategy

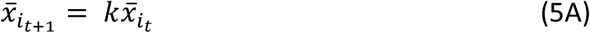

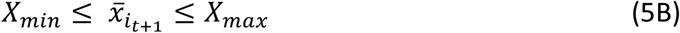

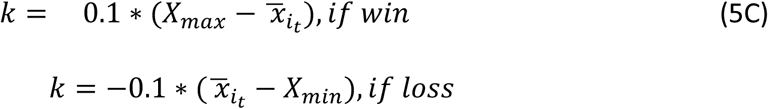

Bayesian updating of self-assessment

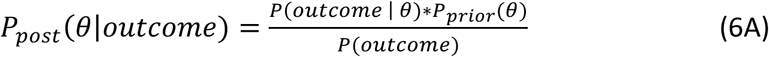

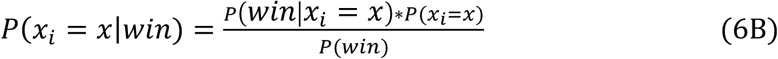

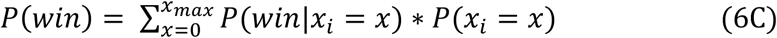

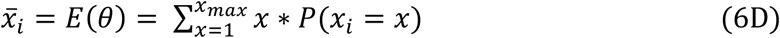

Full investment strategy

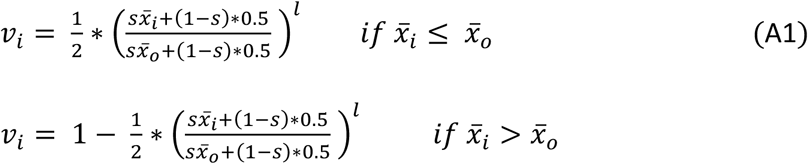

Full likelihood calculation

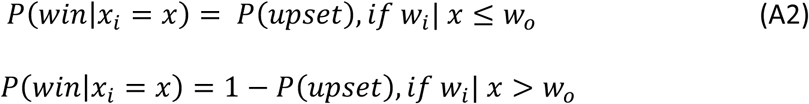

Scaling functions

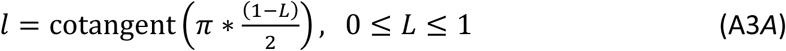

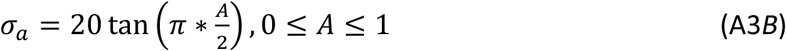

Hyperbolic tangent effort scaling

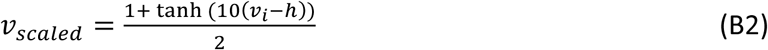

Accuracy Calculation

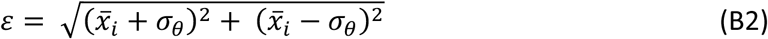

Post-contest feedback on intrinsic ability (*size*)

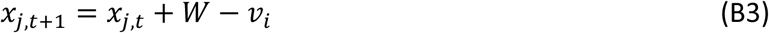

## References

Appleby, M. C. (1983). The probability of linearity in hierarchies. Animal Behaviour, 31(2), 600– 608. 10.1016/S0003-3472(83)80084-0

Arnott, G., & Elwood, R. W. (2008). Information gathering and decision making about resource value in animal contests. In Animal Behaviour (Vol. 76, Issue 3, pp. 529–542). 10.1016/j.anbehav.2008.04.019

Arnott, G., & Elwood, R. W. (2009). Assessment of fighting ability in animal contests. In Animal Behaviour (Vol. 77, Issue 5, pp. 991–1004). 10.1016/j.anbehav.2009.02.010

Ashwood, Z. C., Roy, N. A., Bak, J. H., & Pillow, J. W. (2020). Inferring learning rules from animal decision-making. Advances in Neural Information Processing Systems, 2020*-December*, 3442–3453.

Balph, M. H. (1979). Flock Stability in Relation to Social Dominance and Agonistic Behavior in Wintering Dark-Eyed Juncos. The Auk, 96(4), 714–722. 10.1093/auk/96.4.714

Baxter, C. M., & Dukas, R. (2017). Life history of aggression: effects of age and sexual experience on male aggression towards males and females. Animal Behaviour, 123, 11–20. 10.1016/j.anbehav.2016.10.022

Beacham, J. L. (1988). The relative importance of body size and aggressive experience as determinants of dominance in pumpkinseed sunfish, Lepomis gibbosus. Animal Behaviour, 36(2), 621–623. 10.1016/S0003-3472(88)80042-3

Benincasa, M. D., Earley, R. L., & Hamilton, I. M. (2023). Cumulative experience influences contest investment in a social fish. Behavioral Ecology, 34(6), 1076–1086. 10.1093/beheco/arad078

Bernstein, I., Williams, L., & Ramsay, M. (1983). The expression of aggression in old world monkeys. International Journal of Primatology, 4(2), 113–125. 10.1007/BF02743753

Bierbach, D., Klein, M., Sassmannshausen, V., Schlupp, I., Riesch, R., Parzefall, J., & Plath, M. (2012). Divergent Evolution of Male Aggressive Behaviour: Another Reproductive Isolation Barrier in Extremophile Poeciliid Fishes? International Journal of Evolutionary Biology, 2012, 148745. 10.1155/2012/148745

Bonabeau, E., Theraulaz, G., & Deneubourg, J. L. (1996). Mathematical model of self-organizing hierarchies in animal societies. Bulletin of Mathematical Biology, 58(4), 661–717. 10.1016/0092-8240(95)00364-9

Bonabeau, E., Theraulaz, G., & Deneubourg, J. L. (1999). Dominance orders in animal societies: The self-organization hypothesis revisited. Bulletin of Mathematical Biology, 61(4), 727– 757. 10.1006/bulm.1999.0108

Briffa, M., Sneddon, L. U., & Wilson, A. J. (2015). Animal personality as a cause and consequence of contest behaviour. Biology Letters, 11(3), 20141007. 10.1098/rsbl.2014.1007

Chase, I. D., Bartolomeo, C., & Dugatkin, L. A. (1994). Aggressive interactions and inter-contest interval: How long do winners keep winning? Animal Behaviour, 48(2), 393–400. 10.1006/anbe.1994.1253

Chase, I. D., Coelho, D., Lee, W., Mueller, K., & Curley, J. P. (2022). Networks never rest: An investigation of network evolution in three species of animals. Social Networks, 68, 356–373. 10.1016/j.socnet.2021.09.002

Chase, I. D., Tovey, C., Spangler-Martin, D., & Manfredonia, M. (2002). Individual differences versus social dynamics in the formation of animal dominance hierarchies. Proceedings of the National Academy of Sciences of the United States of America, 99(8), 5744–5749. 10.1073/pnas.082104199

Colombo, M., & Seriès, P. (2012). Bayes in the brain - On Bayesian modelling in neuroscience. British Journal for the Philosophy of Science, 63(3), 697–723. 10.1093/bjps/axr043

Courville, A. C., Daw, N. D., & Touretzky, D. S. (2006). Bayesian theories of conditioning in a changing world. Trends in Cognitive Sciences, 10(7), 294–300. 10.1016/j.tics.2006.05.004

Dayan, P., & Daw, N. D. (2008). Decision theory, reinforcement learning, and the brain. In *Cognitive*, Affective and Behavioral Neuroscience (Vol. 8, Issue 4, pp. 429–453). 10.3758/CABN.8.4.429

Dewsbury, D. A. (1988). Copulatory Behavior as Courtship Communication. Ethology, 79, 218–234. 10.1111/j.1439-0310.1988.tb00712.x

Dugatkin, L. A. (1997). Winner and loser effects and the structure of dominance hierarchies. Behavioral Ecology, 8(6), 583–587. 10.1093/beheco/8.6.583

Earley, R. L., & Dugatkin, L. A. (2002). Eavesdropping on visual cues in green swordtail (Xiphophorus helleri) fights: A case for networking. Proceedings of the Royal Society B: Biological Sciences, 269(1494), 943–952. 10.1098/rspb.2002.1973

Enquist, M., & Leimar, O. (1983). Evolution of fighting behaviour: Decision rules and assessment of relative strength. Journal of Theoretical Biology, 102(3), 387–410. 10.1016/0022-5193(83)90376-4

Enquist, M., Leimar, O., Ljungberg, T., Mallner, Y., & Segerdahl, N. (1990). A test of the sequential assessment game: fighting in the cichlid fish Nannacara anomala. Animal Behaviour, 40(1), 1–14. 10.1016/S0003-3472(05)80660-8

Favre, M., Martin, J. G. A., & Festa-Bianchet, M. (2008). Determinants and life-history consequences of social dominance in bighorn ewes. Animal Behaviour, 76(4), 1373–1380. 10.1016/j.anbehav.2008.07.003

Fawcett, T. W., & Johnstone, R. A. (2010). Learning your own strength: Winner and loser effects should change with age and experience. Proceedings of the Royal Society B: Biological Sciences, 277(1686), 1427–1434. 10.1098/rspb.2009.2088

Fortunato, J. A., & Earley, R. L. (2023). Age-dependent genetic variation in aggression. Biology Letters, 19(1), 20220456. 10.1098/rsbl.2022.0456

Francis, R. C. (1990). Temperament in a Fish: A Longitudinal Study of the Development of Individual Differences in Aggression and Social Rank in the Midas Cichlid. Ethology, 86(4), 311–325. 10.1111/j.1439-0310.1990.tb00439.x

Fuxjager, M. J., Montgomery, J. L., & Marler, C. A. (2011). Species differences in the winner effect disappear in response to post-victory testosterone manipulations. Proceedings of the Royal Society B: Biological Sciences, 278(1724), 3497–3503. 10.1098/rspb.2011.0301

Fuxjager, M. J., Oyegbile, T. O., & Marler, C. A. (2011). Independent and additive contributions of postvictory testosterone and social experience to the development of the winner effect. Endocrinology, 152(9), 3422–3429. 10.1210/en.2011-1099

Gherardi, F. (2006). Fighting behavior in hermit crabs: The combined effect of resource-holding potential and resource value in Pagurus longicarpus. Behavioral Ecology and Sociobiology, 59(4), 500–510. 10.1007/s00265-005-0074-z

Graham, Z. A., & Angilletta, M. J. (2020). Claw size predicts dominance within and between invasive species of crayfish. Animal Behaviour, 166, 153–161. 10.1016/j.anbehav.2020.06.021

Groves, S. (1978). Age-Related Differences in Ruddy Turnstone Foraging and Aggressive Behavior. The Auk, 95(1), 95–103. 10.2307/4085499

Guiaşu, R. C., & Dunham, D. W. (1997). Initiation and outcome of agonistic contests in male form I Cambarus robustus girard, 1852 crayfish (Decapoda, Cambaridae). Crustaceana, 70(4), 480–496. 10.1163/156854097X00069

Hemelrijk, C. K. (2000). Towards the integration of social dominance and spatial structure. Animal Behaviour, 59(5), 1035–1048. 10.1006/anbe.2000.1400

Hickey, J., & Davidsen, J. (2019). Self-organization and time-stability of social hierarchies. PLoS ONE, 14(1), e0211403. 10.1371/journal.pone.0211403

Higginson, A. D., Fawcett, T. W., Houston, A. I., & McNamara, J. M. (2018). Trust your gut: Using physiological states as a source of information is almost as effective as optimal bayesian learning. Proceedings of the Royal Society B: Biological Sciences, 285(1871), 20172411. 10.1098/rspb.2017.2411

Hock, K., & Huber, R. (2006). Modeling the acquisition of social rank in crayfish: Winner and loser effects and self-structuring properties. Behaviour, 143(3), 325–346. 10.1163/156853906775897914

Hotta, T., Awata, S., Jordan, L. A., & Kohda, M. (2021). Subordinate Fish Mediate Aggressiveness Using Recent Contest Information. Frontiers in Ecology and Evolution, 9, 685907. 10.3389/fevo.2021.685907

Hsu, Y., Earley, R. L., & Wolf, L. L. (2006). Modulation of aggressive behaviour by fighting experience: Mechanisms and contest outcomes. In Biological Reviews of the Cambridge Philosophical Society (Vol. 81, Issue 1, pp. 33–74). 10.1017/S146479310500686X

Hsu, Y., Lee, S. P., Chen, M. H., Yang, S. Y., & Cheng, K. C. (2008). Switching assessment strategy during a contest: fighting in killifish Kryptolebias marmoratus. Animal Behaviour, 75(5), 1641–1649. 10.1016/j.anbehav.2007.10.017

Hsu, Y., & Wolf, L. L. (1999). The winner and loser effect: Integrating multiple experiences. Animal Behaviour, 57(4), 903–910. 10.1006/anbe.1998.1049

Hsu, Y., & Wolf, L. L. (2001). The winner and loser effect: What fighting behaviours are influenced? Animal Behaviour, 61(4), 777–786. 10.1006/anbe.2000.1650

Huang, S. P., Yang, S. Y., & Hsu, Y. (2011). Persistence of Winner and Loser Effects Depends on the Behaviour Measured. Ethology, 117(2), 171–180. 10.1111/j.1439-0310.2010.01856.x

Jackson, W. M., & Winnegrad, R. L. (1988). Linearity in dominance hierarchies: a second look at the individual attributes model. Animal Behaviour, 36(4), 1237–1240. 10.1016/S0003-3472(88)80086-1

Kang, P., Tobler, P. N., & Dayan, P. (2024). Bayesian reinforcement learning: A basic overview. Neurobiology of Learning and Memory, 211, 107924. 10.1016/J.NLM.2024.107924

Kolter, J. Z., & Ng, A. Y. (2009). Near-Bayesian exploration in polynomial time. Proceedings of the 26th International Conference On Machine Learning, ICML 2009, 513–520.

Koops, M. A., & Grant, J. W. A. (1993). Weight asymmetry and sequential assessment in convict cichlid contests. Canadian Journal of Zoology, 71(3), 475–479. 10.1139/z93-068

Kumaran, D., Banino, A., Blundell, C., Hassabis, D., & Dayan, P. (2016). Computations Underlying Social Hierarchy Learning: Distinct Neural Mechanisms for Updating and Representing Self-Relevant Information. Neuron, 92(5), 1135–1147. 10.1016/j.neuron.2016.10.052

Kura, K., Broom, M., & Kandler, A. (2016). A Game-Theoretical Winner and Loser Model of Dominance Hierarchy Formation. Bulletin of Mathematical Biology, 78(6), 1259–1290. 10.1007/s11538-016-0186-9

Lan, Y. T., & Hsu, Y. (2011). Prior contest experience exerts a long-term influence on subsequent winner and loser effects. Frontiers in Zoology, 8, 1–12. 10.1186/1742-9994-8-28

Landau, H. G. (1951). On dominance relations and the structure of animal societies: I. Effect of inherent characteristics. The Bulletin of Mathematical Biophysics, 13(1), 1–19. 10.1007/BF02478336

Laskowski, K. L., Chang, C. C., Sheehy, K., & Aguiñaga, J. (2022). Consistent Individual Behavioral Variation: What Do We Know and Where Are We Going? In Annual Review of Ecology, Evolution, and Systematics (Vol. 53, pp. 161–182). 10.1146/annurev-ecolsys-102220-011451

Laskowski, K. L., Wolf, M., & Bierbach, D. (2016). The making of winners (And losers): How early dominance interactions determine adult social structure in a clonal fish. Proceedings of the Royal Society B: Biological Sciences, 283(1830), 20160183. 10.1098/rspb.2016.0183

Le Pelley, M. E. (2004). The role of associative history in models of associative learning: A selective review and a hybrid model. In Quarterly Journal of Experimental Psychology Section B: Comparative and Physiological Psychology (Vol. 57, Issue 3, pp. 193–243). 10.1080/02724990344000141

Leimar, O. (2021). The evolution of social dominance through reinforcement learning. American Naturalist, 197(5), 560–575. 10.1086/713758

Leimar, O., Austad, S., & Enquist, M. (1991). A test of the sequential assessment game: fighting in the bowl and doily spider Frontinella pyramitela. Evolution, 45(4), 862–874. 10.1111/j.1558-5646.1991.tb04355.x

Leimar, O., & Bshary, R. (2022). Reproductive skew, fighting costs and winner–loser effects in social dominance evolution. Journal of Animal Ecology, 91(5), 1036–1046. 10.1111/1365-2656.13691

Luttbeg, B. (1996). A comparative Bayes tactic for mate assessment and choice. Behavioral Ecology, 7(4), 451–460. 10.1093/beheco/7.4.451

Marden, J. H., & Waage, J. K. (1990). Escalated damselfly territorial contests are energetic wars of attrition. Animal Behaviour, 39(5), 954–959. 10.1016/S0003-3472(05)80960-1

Maynard Smith, J. (1974). The theory of games and the evolution of animal conflicts. Journal of Theoretical Biology, 47(1), 209–221. 10.1016/0022-5193(74)90110-6

Maynard Smith, J., & Parker, G. A. (1976). The logic of asymmetric contests. Animal Behaviour, 24(1), 159–175. 10.1016/S0003-3472(76)80110-8

McNamara, J. M., Green, R. F., & Olsson, O. (2006). Bayes’ theorem and its applications in animal behaviour. Oikos, 112(2), 243–251. 10.1111/j.0030-1299.2006.14228.x

McNamara, J. M., & Leimar, O. (2020). Game theory in biology: concepts and frontiers. Oxford University Press.

Mesterton-Gibbons, M. (1999). On the evolution of pure winner and loser effects: A game-theoretic model. Bulletin of Mathematical Biology, 61(6), 1151–1186. 10.1006/bulm.1999.0137

Mesterton-Gibbons, M., Dai, Y., & Goubault, M. (2016). Modeling the evolution of winner and loser effects: A survey and prospectus. In Mathematical Biosciences (Vol. 274, pp. 33–44). 10.1016/j.mbs.2016.02.002

Mesterton-Gibbons, M., Marden, J. H., & Dugatkin, L. A. (1996). On wars of attrition without assessment. Journal of Theoretical Biology, 181(1), 65–83. 10.1006/jtbi.1996.0115

Niemelä, P. T., & Dingemanse, N. J. (2018). Meta-analysis reveals weak associations between intrinsic state and personality. Proceedings of the Royal Society B: Biological Sciences, 285(1873), 20172823. 10.1098/rspb.2017.2823

Niv, Y. (2009). Reinforcement learning in the brain. Journal of Mathematical Psychology, 53(3), 139–154. 10.1016/J.JMP.2008.12.005

O’Connor, C. M., Reddon, A. R., Ligocki, I. Y., Hellmann, J. K., Garvy, K. A., Marsh-Rollo, S. E., Hamilton, I. M., & Balshine, S. (2015). Motivation but not body size influences territorial contest dynamics in a wild cichlid fish. Animal Behaviour, 107, 19–29. 10.1016/j.anbehav.2015.06.001

Okasha, S. (2013). The evolution of Bayesian updating. Philosophy of Science, 80(5), 745–757. 10.1086/674058

Olsson, O. (2006). Decisions under uncertainty: Bayesian foraging - A meeting held at Lund University in August 2003. Oikos, 112(2), 241–242. 10.1111/j.0030-1299.2006.14383.x

Parker, G. A. (1974). Assessment strategy and the evolution of animal conflicts. Journal of Theoretical Biology, 47(1), 223–243.

Pearce, J. M., & Hall, G. (1980). A model for Pavlovian learning: Variations in the effectiveness of conditioned but not of unconditioned stimuli. Psychological Review, 87(6), 532–552. 10.1037/0033-295X.87.6.532

Pouget, A., Beck, J. M., Ma, W. J., & Latham, P. E. (2013). Probabilistic brains: Knowns and unknowns. In Nature Neuroscience (Vol. 16, Issue 9, p. 11701178). 10.1038/nn.3495

Poupart, P., Vlassis, N., Hoey, J., & Regan, K. (2006). An analytic solution to discrete bayesian reinforcement learning. ACM International Conference Proceeding Series, 148, 697–704. 10.1145/1143844.1143932

Rescorla, R. A., & Wagner, A. R. (1972). A theory of Pavlovian conditioning: The effectiveness of reinforcement and non-reinforcement. In A. H. Black & W. F. Prokasy (Eds.), Classical Conditiong II: Current Theory and Research (Issue January 1972, pp. 64–69). Appleton-Century-Crofts.

Rowell, T. E. (1974). The concept of social dominance. Behavioral Biology, 11(2), 131–154. 10.1016/S0091-6773(74)90289-2

Rutte, C., Taborsky, M., & Brinkhof, M. W. G. (2006). What sets the odds of winning and losing? Trends in Ecology and Evolution, 21(1), 16–21. 10.1016/j.tree.2005.10.014

Samuels, A., Silk, J. B., & Altmann, J. (1987). Continuity and change in dominance relations among female baboons. Animal Behaviour, 35(3), 785–793. 10.1016/S0003-3472(87)80115-X

Schlinger, B. A., Palter, B., & Callard, G. V. (1987). A method to quantify aggressiveness in Japanese quail (Coturnix c. japonica). Physiology and Behavior, 40(3), 343–348. 10.1016/0031-9384(87)90057-6

Senar, J. C., Camerino, M., & Metcalfe, N. B. (1990). Familiarity Breeds Tolerance: the Development of Social Stability in Flocking Siskins (Carduelis spinus). Ethology, 85(1), 13–24. 10.1111/j.1439-0310.1990.tb00381.x

Simons, N. D., Michopoulos, V., Wilson, M., Barreiro, L. B., & Tung, J. (2022). Agonism and grooming behaviour explain social status effects on physiology and gene regulation in rhesus macaques. Philosophical Transactions of the Royal Society B: Biological Sciences, 377(1845), 20210132. 10.1098/rstb.2021.0132

Snyder-Mackler, N., Burger, J. R., Gaydosh, L., Belsky, D. W., Noppert, G. A., Campos, F. A., Bartolomucci, A., Yang, Y. C., Aiello, A. E., O’Rand, A., Harris, K. M., Shively, C. A., Alberts, S. C., & Tung, J. (2020). Social determinants of health and survival in humans and other animals. In Science (Vol. 368, Issue 6493, p. eaax9553). 10.1126/science.aax9553

Stamps, J. A., & Krishnan, V. V. (2014). Combining information from ancestors and personal experiences to predict individual differences in developmental trajectories. American Naturalist, 184(5), 647–657. 10.1086/678116

Thompson, K. V. (1998). Self assessment in juvenile play. In M. Bekoff & J. Byers (Eds.), Animal Play: Evolutionary, Comparative and Ecological Perspectives (pp. 183–204). Cambridge University Press. 10.1017/cbo9780511608575.010

Thor, D. H., & Holloway, W. R. (1984). Developmental analyses of social play behavior in juvenile rats. Bulletin of the Psychonomic Society, 22(6), 587–590. 10.3758/BF03333916

Tibbetts, E. A., Pardo-Sanchez, J., & Weise, C. (2022). The establishment and maintenance of dominance hierarchies. Philosophical Transactions of the Royal Society B: Biological Sciences, 377(1845), 20200450. 10.1098/rstb.2020.0450

Tibbetts, E. A., Wong, E., & Bonello, S. (2020). Wasps Use Social Eavesdropping to Learn about Individual Rivals. Current Biology, 30(15), 3007–3010.E2. 10.1016/j.cub.2020.05.053

Valone, T. J. (2006). Are animals capable of Bayesian updating? An empirical review. Oikos, 112(2), 252–259. 10.1111/j.0030-1299.2006.13465.x

Van Doorn, G. S., Hengeveld, G. M., & Weissing, F. J. (2003). The evolution of social dominance I: Two-player models. Behaviour, 140(10), 1305–1332. 10.1163/156853903771980602

Vlassis, N., Ghavamzadeh, M., Mannor, S., & Poupart, P. (2012). Bayesian Reinforcement Learning. In M. Wiering & M. van Otterlo (Eds.), Reinforcement Learning. Adaptation, Learning, and Optimization (Vol. 12, pp. 359–386). Springer. 10.1007/978-3-642-27645-3_11

Whitehouse, M. E. A. (1997). Experience influences male-male contests in the spider Argyrodes antipodiana (Theridiidae: Araneae). Animal Behaviour, 53(5), 913–923. 10.1006/anbe.1996.0313

Zhou, T., Sandi, C., & Hu, H. (2018). Advances in understanding neural mechanisms of social dominance. In Current Opinion in Neurobiology (Vol. 49, pp. 99–107). 10.1016/j.conb.2018.01.006

