## Supplemental Code for "Bayesian updating for self-assessment explains social dominance and winner-loser effects": fight.py

```
## Load required packages
import numpy as np

import itertools,random

from scipy.special import rel_entr, logit
from scipy.stats import mode
from scipy.stats import norm

from sklearn.metrics import auc

import matplotlib.pyplot as plt
from matplotlib import cm

from bayesbots import Fish
from bayesbots import Params

## Simple-ish object to keep track of matchups (and decide outcome if necessary)
class Fight():
    __slots__ = ('fish1', 'fish2', 'fishes', 'params', 'mechanism', 'level','efforts', 'outcome', 'outcome_params','_SE_func','_wager_func', 'scale', 'idx', 'food', 'p_win','f_min','min_normed', 'roll', 'winner', 'loser')
    def __init__(self,fish1,fish2,params=None,
                idx=0,outcome=None,level=None):
        self.fish1 = fish1
        self.fish2 = fish2
        self.fishes = [fish1,fish2]
        if params is None:
            params=Params()
        self.params = params.copy()
        if outcome is None:
            self.mechanism = params.f_outcome
        else:
            self.mechanism = outcome
        self.level = level
        self.outcome = '?'
        self.outcome_params = params.outcome_params
        self.scale= 0.1 # deprecated parameter used for hock.
        self.idx = idx
        self.food = params.food
        self.p_win = None
        self.f_min = None
        
        #self._SE_func = self._SE_product
        self._SE_func = self._SE_sum
        self._wager_func = self._wager_curve

    def run_outcome(self):
        if self.mechanism == 'math':
            self.outcome,self.level = self.mathy()
            
        else:
            real_outcome,real_level = self.mathy(self.outcome_params)
            self.outcome = self.mechanism
            if self.level is None:
                self.level = real_level
      
        self.winner = self.fishes[self.outcome]
        self.loser = self.fishes[1-self.outcome]
        return self.outcome

    def _wager_curve(self,w,L=None): #L=np.tan(np.pi/2 -  ((-0.9 + 1)/2)*np.pi*0.5)):
        return (w ** self.params.L) / 2

## Staty gave me this, but it's an L_B system apparently
    def _wager_curve_staty(self,w,L=None):
        p = self.params.m
        d = self.params.a
        B = (p/(1-p))**d
        X = (w/(1-w))**d
        return X / (X+B) /2

    def _SE_product(self,rel_size,effort):
        S = self.params.scaled_params[0]
        F = self.params.scaled_params[1]

        s = self.params.outcome_params[0]
        f = self.params.outcome_params[1]
## Now have to handle stupid edge cases here.
        if s == -1:
            if rel_size == 1:
                wager = 1
            else:
                wager = 0
            return wager
        elif f == -1:
            if effort == 1:
                wager = rel_size ** S
            elif effort == max([self.fish1.effort,self.fish2.effort]):
                wager = 1
            else:
                wager = 0
                
        else:
            wager = (rel_size ** S) * (effort ** F)
        #print('product:',S,F)
        return wager

    def _SE_sum(self,rel_size,effort):
        s = self.params.scaled_params[0]
        f = self.params.scaled_params[1]
        s = self.params.outcome_params[0]
        #s = 0.5
        f = 1-s
        wager = s * rel_size + f * effort
        #print(rel_size,effort,s,f)
        return wager

## Cleaning this up, hopefully it stil works
    def mathy(self,params=None):

        f1_wager,f2_wager = 0,0
        min_normed = 0
        if params is None:
            S,F,L = self.params.scaled_params
            s,f,l = self.params.outcome_params
        else:
            s,f,l = params
            scaled_params = (np.array(params) + 1) /2
            S,F,L = np.tan(np.pi/2 - scaled_params*np.pi/2)
        f1_size = self.fish1.size
        f2_size = self.fish2.size
        max_size = max([f1_size,f2_size])
        f1_rel_size = f1_size / max_size
        f2_rel_size = f2_size / max_size

        f1_effort = self.fish1._choose_effort(self.fish2,self)
        f2_effort = self.fish2._choose_effort(self.fish1,self)
        self.fish1.effort = f1_effort
        self.fish2.effort = f2_effort

        f1_rel_effort = f1_effort / max([f1_effort,f2_effort])
        f2_rel_effort = f2_effort / max([f1_effort,f2_effort])


        f1_wager = self._SE_func(f1_size / 100,f1_effort)
        f2_wager = self._SE_func(f2_size / 100,f2_effort)

        self.fish2.wager = f2_wager

        min_normed = min([f1_wager,f2_wager])/max([f1_wager,f2_wager])
        if np.isnan(min_normed):
            p_win = 0.5
            import pdb;pdb.set_trace()
        elif min_normed == 1:
            p_win = 0.5
        else:
            p_win = self._wager_func(min_normed,l)
        if p_win == 0.5:
            f_min = np.random.randint(2)
        else:
            f_min = 1 - np.argmax([f1_wager,f2_wager])
        self.p_win = p_win
        self.f_min = f_min
        self.min_normed = min_normed

        self.fish1.wager = f1_wager
        roll = random.random()
        
        if roll < self.p_win: ## probability that the "lower invested" fish wins
            winner = f_min
        else:
            winner = 1-f_min
        self.roll = roll
        loser = 1-winner
        level = min([f1_wager,f2_wager])
        self.efforts = [self.fish1.effort,self.fish2.effort]

        return winner,level

    def summary(self):
        sum_str =  ' '.join([str(self.fish1.idx),'vs',str(self.fish2.idx),str(self.outcome),': Fitness =',str(not self.food),': So,',str(self.winner.idx),'won, prob of upset was:',str(self.p_win)])
        strat_str = ' '.join(['Fish1:',str(self.fish1.params.effort_method),'Fish2:',str(self.fish2.params.effort_method),str(self.fish1.params.mutant),str(self.fish2.params.mutant)])
        header_str = 'Fish : Size : Effort : Estimate : Guess'
        effort_str1 = ' '.join(['Fish',str(self.fish1.idx),str(self.fish1.size),str(self.fish1.effort),str(self.fish1.estimate),str(self.fish1.guess)])
        effort_str2 = ' '.join(['Fish',str(self.fish2.idx),str(self.fish2.size),str(self.fish2.effort),str(self.fish2.estimate),str(self.fish2.guess)])
        #header_str = 'Fish : Size Own_estimate Opp_estimate Effort'
        #effort_str1 = ' '.join(['Fish1:',str(self.fish1.size),str(self.fish1.estimate),str(self.fish1.opp_estimate),str(self.fish1.effort)])
        #effort_str2 = ' '.join(['Fish2:',str(self.fish2.size),str(self.fish2.estimate),str(self.fish2.opp_estimate),str(self.fish2.effort)])
        sum_str = '\n'.join([sum_str,strat_str,header_str,effort_str1,effort_str2])
        return sum_str

if __name__ == "__main__":
    f1 = Fish()
    f2 = Fish()
    f = Fight(f1,f2)
    f.run_outcome()
    print(f.outcome,f.winner.idx,f.level)
```
