## Supplemental Code for "Bayesian updating for self-assessment explains social dominance and winner-loser effects": fish.py

```
## Load required packages
import numpy as np

import itertools,random

from scipy.special import rel_entr, logit
from scipy.stats import mode
from scipy.stats import norm
from scipy.stats import truncnorm

from sklearn.metrics import auc

import matplotlib.pyplot as plt
from matplotlib import cm

from bayesbots import Params

## Define a fish object

## Fish object with internal rules and estimates
# Reminder: r_rhp is the rate that rhp increases when they eat/win
#           a_growth is whether the growth rate should be asymptotic 
#           Prior is either None, in which case it uses the baseline, 
#               True: in which is uses a true mean with mean/5 as the std
#               an int: in which case it uses the int as the std (10 would be average)
class Fish:
    __slots__ = ('idx', 'name', 'params', 'prior', 'likelihood', 'likelihood_dict', 'age', 'xs', 'r_rhp', 'a_growth', 'c_aversion', 'acuity', 'awareness', 'insight', 'size', 'estimate', 'default_prior', 'cdf_prior', 'estimate_', 'prior_mean', 'prior_std', 'win_record', 'est_record', 'est_record_', 'sdest_record', 'range_record', 'effort', 'decay', 'decay_all', 'discrete', 'update', '_choose_effort', 'update_method', 'wager', 'boost', 'energy', 'max_energy', 'energy_cost', 'size_record', 'energy_record', 'fitness_record', 'alive', 's_max', 'naive_params', 'naive_prior', 'naive_estimate', 'naive_likelihood', 'guess', 'correction','trunced','_scale_func')
    def __init__(self,idx=0,params=None,
                 age=None,size=None,
                 prior=None,likelihood=None,likelihood_dict=None,
                 ):
        self.idx = idx
        self.name = idx

        if params is None:
            params = Params()
        self.params = params.copy()
        if prior is None:
            self.prior = prior
        else:
            self.prior = params.prior
        if likelihood is None:
            self.likelihood = likelihood
        else:
            self.likelihood = params.likelihood
        if likelihood_dict is None:
            self.likelihood_dict = likelihood_dict
        else:
            self.likelihood_dict = params.likelihood_dict

        if age is not None:
            self.age = age
            self.params.age = self.age
        else:
            self.age = params.age
        self.xs = params.xs
        self.r_rhp = params.r_rhp
        self.a_growth = params.a_growth
        self.c_aversion = params.c_aversion
        self.acuity = params.acuity
        self.awareness = params.awareness
        self.insight = params.insight

        if size is not None:
            self.size = size
            self.params.size = self.size
        elif params.size is not None:
            self.size = params.size
        else:
            self.size = np.random.normal(self.params.mean_size,self.params.sd_size)
        self.size = np.clip(self.size,self.params.min_size,self.params.max_size)
        self.params.size = self.size
        self.trunced = None

        if np.isinf(self.params.A):
            self.prior = np.ones_like(self.xs) / len(self.xs)
            self.estimate = 0.5
        elif self.params.A == 0:
            self.prior = np.zeros_like(self.xs)
            self.prior[np.argmax(self.xs >= self.size)] = 1
        elif params.prior is True:
            self.prior = norm.pdf(self.xs,self.size,self.params.A)
            self.prior = self.prior / np.sum(self.prior)
        elif isinstance(params.prior,int):
            self.estimate = np.clip(np.random.normal(params.prior,self.params.A),self.params.min_size,self.params.max_size)
            if params.prior == -1:
                self.prior = np.ones_like(self.xs) / len(self.xs)
            else:
                self.prior = norm.pdf(self.xs,self.estimate,self.params.A)
                self.prior = self.prior / np.sum(self.prior)
        elif params.prior is not None:
            self.prior = params.prior
        else: ## e.g. prior is a string
            
            if self.params.awareness == 0:
                self.estimate = self.size
                self.prior = np.zeros_like(self.xs)
                self.prior[np.argmax(self.xs >= self.estimate)] = 1
            elif self.params.awareness == 1:
                self.estimate = (self.params.max_size + self.params.min_size ) / 2
                self.prior = np.ones_like(self.xs) / len(self.xs)
            else:
                mu,scale = self.size,self.params.A
                a_min,b_max = self.params.min_size,self.params.max_size
                a,b = (a_min - mu) / scale,(b_max - mu) / scale
                trunc_prior = truncnorm(a,b,loc=mu,scale=scale)

                self.estimate = trunc_prior.rvs()
                prior = trunc_prior.pdf(self.xs)
                self.prior = prior / np.sum(prior)
        self.default_prior = norm.pdf(self.xs,50,self.params.A)
        self.default_prior = self.prior / np.sum(self.prior)

        ## Define the rules one when to escalate, based on confidence
        #self.escalation_thresholds = escalation
        self.cdf_prior = self._get_cdf_prior(self.prior)
        est_5,est_25,est_75,est_95 = self._get_range_prior(self.cdf_prior)
        self.estimate_ = self.xs[np.argmax(self.prior)]
        prior_mean,prior_std = self.get_stats()
        self.estimate = prior_mean

        self.win_record = []
        self.est_record = [self.estimate]
        self.est_record_ = [self.estimate_]
        self.sdest_record = [prior_std]
        self.range_record = [[est_5,est_25,est_75,est_95]]
        self.effort = None

        update_method = params.update_method
        effort_method = params.effort_method
        if update_method == 'bayes':
            #print('using bayes')
            self.update = self.update_prior
        elif update_method == 'linear':
            self.update = self.update_linear
        elif update_method == 'dugatkin':
            self.update = self.update_dugatkin
        elif update_method == 'size_boost':
            self.update = self._size_boost
        elif update_method == 'blind':
            self.update = self.update_blind
        else:
            self.update = self.no_update

        #print('setting effort...',effort_method)
        if effort_method[0] is None:
            if effort_method[1] is None:
                print('leroy!')
                self._choose_effort = self.leroy_jenkins
            elif effort_method[1] == 0.5: 
                #print('half leroy!')
                self._choose_effort = self.half_jenkins
            elif effort_method[1] == '?':
                self._choose_effort = self.random_effort
            elif effort_method[1] == '!':
                self._choose_effort = self.explore_effort
                self.params.effort_method = [1,1]
            else: ## if you give anything else, it uses baseline effort
                if self.params.baseline_effort ==0 or self.params.baseline_effort is None:
                    self.params.baseline_effort = np.random.random()
                self.effort = params.max_energy * self.params.baseline_effort
                self._choose_effort = self.float_jenkins
        elif effort_method == 'PerfectNudge':
            self._choose_effort = self.nudge_effort
        elif effort_method == 'PerfectPoly':
            #self._choose_effort = self.poly_effort
            self._choose_effort = self.poly_effort_combo
            self.acuity = 0
            self.awareness = 0
            self.estimate = self.size
        elif effort_method == 'EstimatePoly':
            self._choose_effort = self.poly_effort
        elif effort_method == 'SmoothPoly':
            #self._choose_effort = self.poly_effort_prob
            self._choose_effort = self.poly_effort_combo
            self._scale_func = lambda effort: effort
        elif effort_method == 'ScaledPoly':
            self._choose_effort = self.poly_effort_combo
            self._scale_func = self.scale_effort
        elif effort_method == 'LuckyPoly':
            self._choose_effort = self.poly_effort_lucky
        elif effort_method == 'ExplorePoly':
            self._choose_effort = self.poly_explore
        else:
            print('Warning!! Effort strategy not found')
            print('Defaulting to choose_effort_energy')

            self._choose_effort = self.choose_effort_energy

        self.update_method = update_method
        self.params.effort_method = params.effort_method
        #self.effort = 0
        self.wager = 0
        self.boost = 0 ## Initial boost, needs to be 0, will change with winner/loser effect
        self.energy = params.start_energy ## add some cost to competing
        self.max_energy = params.max_energy
        self.energy_cost = params.energy_cost

## Initialize size and energy records
        self.size_record = [self.size]
        self.energy_record = [self.energy]
        self.fitness_record = [0]

        self.alive = True
        self.s_max = 100
        ## Define naive prior/likelihood for 'SA'
        self.naive_params = params.outcome_params
## This should be an average fish.
        #naive_prior = self._prior_size(self.age,xs=self.xs)
        self.naive_prior = np.array(self.prior)
        #self.naive_estimate = np.sum(self.prior * self.xs / np.sum(self.prior))
        self.naive_estimate = self.estimate
        if params.likelihood is not None:
            #print('using existing likelihood')
            self.naive_likelihood = params.likelihood
        elif self.params.effort_method[1] == 0 or update_method == 'blind':
            self.naive_likelihood = self._define_naive_likelihood()
        else:
            self.naive_likelihood = None
        self.likelihood_dict = params.likelihood_dict
## Apply winner/loser effect. This could be more nuanced, eventually should be parameterized.
        
    def build_trunc(self,size,C):
        mu = size
        a = (self.params.min_size - size) / C
        b = (self.params.max_size - size) / C
        X = truncnorm(a,b,loc=mu,scale=C)
        return X

    def assess_me(self,opp):
        if opp.params.C != self.params.C:
            temp_trunc = self.build_trunc(self.size,opp.params.C)
        elif self.trunced == None:
            self.trunced = self.build_trunc(self.size,self.params.C)
            temp_trunc = self.trunced
        else:
            temp_trunc = self.trunced
        return temp_trunc.rvs()

    def coin(self):
        if random.random() < 0.5:
            return 1
        else:
            return -1

    def mutate(self,step=0.01,jump=False): ## function to mutate both baseline effort, and poly params
        if random.random() < 0.2 or jump:
            shift = (np.random.random() - 0.5) * step * 10
            #shift = np.random.random() - 1
        else:
            shift = step
        self.params.poly_param_a = self.params.poly_param_a + shift * self.coin()
        self.params.poly_param_b = self.params.poly_param_b + shift * self.coin()

        if self.params.effort_method[0] == None:
            self.params.baseline_effort = self.params.baseline_effort + self.coin() * step 
            self.params.baseline_effort = np.clip(self.params.baseline_effort,0,1)
            self.effort = self.params.baseline_effort

    def _set_boost(self,win,fight):
        if win:
            other_fish = fight.loser
            self.boost = np.clip(self.boost + .1,-.5,.5)
        else:
            other_fish = fight.winner
            self.boost = np.clip(self.boost - .1,-.5,.5)
        self.win_record.append([other_fish.size,win,self.effort])
        
    def _size_boost(self,win,fight,shift=2):
        if win:
            other_fish = fight.loser
            self.estimate += shift
            self.estimate_ += shift
        else:
            other_fish = fight.winner
            self.estimate -= shift
            self.estimate_ -= shift
        self.estimate = np.clip(self.estimate,self.params.min_size,self.params.max_size)
        self.win_record.append([other_fish.size,win,self.effort])
        self.est_record.append(self.estimate)

    def _get_cdf_prior(self,prior):
        normed_prior = self.prior / np.sum(self.prior)
        cdf_prior = np.cumsum(normed_prior)
        return cdf_prior
    
    def _get_range_prior(self,cdf_prior):
        cdf_5 = np.argmax(cdf_prior > 0.05)
        cdf_25 = np.argmax(cdf_prior > 0.25)
        cdf_75 = np.argmax(cdf_prior > 0.75)
        cdf_95 = np.argmax(cdf_prior >= 0.95)
        return self.xs[cdf_5],self.xs[cdf_25],self.xs[cdf_75],self.xs[cdf_95]

    def _update(self,prior,likelihood,xs = None):
        post = prior * likelihood
        post_sum = np.sum(post)
        if post_sum == 0: ## handles impossible cases
            post = np.ones_like(post) / len(post)
        else:
            post = post / post_sum
        return post
    
    def _decay_flat(self,prior,xs = None,decay=.001):
        post = prior + np.ones_like(prior) * decay
        post = post / np.sum(post)
        return post

    def _decay_norm(self,prior,xs = None,decay = 2):
        post = prior + (self.naive_prior ** decay)
        post = post / np.sum(post)
        return post

    def _prior_size(self,t,xs = None):
        if xs is None:
            xs = self.xs
        s_mean = self._growth_func(t)
        sd = s_mean / 5
        prior = norm.pdf(xs,s_mean,sd)
        return prior / np.sum(prior)

    def _growth_func(self,t):
        size = 7 + self.params.max_size - self.params.max_size/np.exp(t/100)
        return size
    
    def _win_by_ratio(self,r, k=0.1,m=0):
        # sigmoid function
        # use k to adjust the slope
        p = 1 / (1 + np.exp(-(r-m) / k))
        return p

## This is a little clunky to have to define this here also, be sure to know if it matches the fight._wager_curve
## Technically, fight could inherit this from here, but this opens the possibility of having different fish with different LF's. 
    def _wager_curve_old(self,w,l=.25):
        a = logit(1-l)
        prob_win = w ** (float(np.abs(a))**np.sign(a)) / 2
        return prob_win

    def _wager_curve(self,w,l=.05):
        if l == -1:
            prob_win = 0
        if l == 1:
            prob_win = 0.5
        else:
            #L = np.tan((np.pi - l)/2)
            #L = np.tan(np.pi/2 - l*np.pi/2) 
            L = self.params.L
            prob_win = (w**L) / 2
        return prob_win

## As below, but instead it's based on the wager (assuming opponent size and effort are unknown)
## Assumes opponent size is your own estimate. It's all a mess. 

## I don't think I use this anymore
## NOTE: Come back
    def _likelihood_function_wager(self,x,opp_size=None,opp_effort=None,opp_wager=None,outcome_params = [.3,.3,.3]):
        s,e,l = outcome_params
        shifted_params = (np.array(outcome_params) + 1) / 2
        #S,F,L = np.tan(np.pi/2 - np.array(outcome_params)*np.pi/2)
        S,F,L = np.tan(np.pi/2 - shifted_params*np.pi/2)
        if opp_size is None:
            opp_size = self.est_record[0] ## Assume all opponents are average size
        opp_rel_size = opp_size / max([opp_size,x])
        if opp_effort is None:
            if opp_wager is not None:
                opp_effort = (opp_wager ** (1/e)) / (opp_rel_size ** S) ## This should work, but check it
            else:
                opp_effort = opp_size / 100
        if opp_wager is None:
            opp_wager = (opp_rel_size ** S) * (opp_effort ** F)
        rel_size = x / max([opp_size,x])
        my_wager = (rel_size ** S) * (self.effort ** F)
        if my_wager > opp_wager:
            rel_wager = opp_wager / my_wager
            p_win = 1-self._wager_curve(rel_wager,l)
        elif my_wager == opp_wager:
            p_win = 0.5
        else:
            rel_wager = my_wager / opp_wager
            p_win = self._wager_curve(rel_wager,l)
        return p_win

## It would be nice to just update all this to include fight info
    def _define_likelihood_w(self,outcome_params = [.3,.3,.3],win=True):
        if xs is None:
            xs = self.xs
        likelihood = np.zeros(len(xs))
        if win:
            for s in range(len(xs)):
                likelihood[s] = self._likelihood_function_wager(xs[s],outcome_params=outcome_params)
        elif not win:
            for s in range(len(xs)):
                likelihood[s] = 1-self._likelihood_function_wager(xs[s],outcome_params=outcome_params)
        return likelihood

    def _use_simple_likelihood(self,fight,win):
        if False in (fight.outcome_params == self.naive_params):
            #print('rebuilding likelihood')
            self.naive_likelihood = self._define_naive_likelihood(fight)
            self.naive_params = fight.outcome_params
        if win:
            likelihood = self.naive_likelihood
        else:
            likelihood = 1 - self.naive_likelihood
        return likelihood

## This is very slow, slow enough that for simulations, I should do it only once
    def _define_naive_likelihood(self,fight=None):
        #print('initializing likelihood')
        if fight is None:
            s,e,l = self.naive_params
        else:
            s,e,l = fight.outcome_params
        shifted_params = (np.array([s,e,l]) + 1) / 2
        S,F,L = np.tan(np.pi/2 - shifted_params*np.pi/2)
        likelihood = np.zeros(len(self.xs))
## This assumes that all fish are the same age as you
## Should assume they are average? 
        for i_ in range(len(self.xs)):
            i = self.xs[i_]
            prob_ij = 0
            for j_ in range(len(self.xs)):
                j = self.xs[j_]
                prob_j = self.default_prior[j_]
                if prob_j != 0:
                    if i > j:
                        i_wager = (i/100)**F
                        j_wager = (j / i)**S * (j/100)**F
                        rel_wager = j_wager / i_wager
                        likelihood_j = 1 - self._wager_curve(rel_wager,l)
                    elif i <= j:
                        i_wager = (i/j)**S * (i/100)**F
                        j_wager = (j/100)**F
                        rel_wager = i_wager / j_wager
                        likelihood_j = self._wager_curve(rel_wager,l)
                    #print(rel_wager)
                    #print(j,likelihood_j,prob_j)
                    prob_ij += likelihood_j * prob_j
                    #print(i,j,rel_wager)
                else:
                    continue
            #print(prob_ij)
            likelihood[i_] = prob_ij
        return likelihood

## Uses fight as input, runs solo, should be one likelihood to rule them all
    def _define_likelihood_solo(self,fight,win):
        likelihood = np.zeros(len(self.xs))
        s,e,l = fight.outcome_params
        shifted_params = (np.array(fight.outcome_params) + 1) / 2
        S,F,L = np.tan(np.pi/2 - shifted_params*np.pi/2)
        if win:
            e_self = fight.winner.effort
            w_opp = fight.loser.wager
## Assume fixed, avg effort for all opponents
## NOTE: Somehow you need to transform size to relative size, without knowing opponent size.
            xs_rel = self.xs / self.estimate 
            for s in range(len(self.xs)):
                likelihood[s] = self._likelihood_function_wager(self.xs[s],outcome_params=fight.outcome_params)
        elif not win:
            #print('they lost!')
            e_self = fight.loser.effort
            w_opp = fight.winner.wager
            xs_rel = self.xs / max([fight.loser.size,fight.winner.wager / (self.effort**e)])
            for s in range(len(self.xs)):
                likelihood[s] = 1 - self._likelihood_function_wager(self.xs[s],outcome_params=fight.outcome_params)
        return likelihood

## Assumes equal effort, which probably isn't quite right. I could assume accurate effort
    def _likelihood_function_size(self,x,x_opp=50,effort=False):
        if x >=x_opp:
            r_diff = (x - x_opp)/x # Will be positive
        elif x_opp > x:
            r_diff = (x - x_opp)/x_opp # Will be negative
        p_win = self._win_by_ratio(r_diff)
        return p_win

## This is closer to the true likelihood, although maybe it should infer relative effort
    def _likelihood_function_se(self,x_size,other_fish,x_eff = None,o_eff = None,fight=None):
        print('using likelihood function se')
## Check fight params
        if fight is None:
            s,e,l = self.naive_params
        else:
            s,e,l = fight.outcome_params
        shifted_params = (np.array([s,e,l]) + 1) / 2
        S,F,L = np.tan(np.pi/2 - shifted_params*np.pi/2)
        if x_eff is None:
            x_eff = 1
        if o_eff is None:
            if self.params.post_acuity is True:
                print('using known effort')
                o_eff = other_fish.effort
            elif fight.level is not None:
                o_eff = fight.level
            else:
                o_eff = 1
## Get relative sizes and wagers
        o_size = other_fish.size
        if x_size >= o_size:
            x_wag = x_eff**e
            o_wag = (o_size/x_size)**s * (o_eff**e)
        else:
            x_wag = (x_size/o_size)**s * (x_eff**e)
            o_wag = o_eff ** e
## Get relative wagers, and plug them into the wager_curve
        if x_wag > o_wag:
            rel_wag = o_wag / x_wag
            p_win = 1-self._wager_curve(rel_wag,l) ## because opponent is the underdog, 1-upset is x's p(win)
        elif x_wag == o_wag:
            return 0.5
        else: ## x_wager is smaller, so they're the underdog
            rel_wag = x_wag / o_wag
            p_win = self._wager_curve(rel_wag,l)
        print('rel_wager',rel_wag)
        if False and fight.p_win is not None:
            if fight.fish1.size == o_size:
                other_fish = fight.fish1
                me = fight.fish2
            elif fight.fish2.size == o_size:
                other_fish = fight.fish2
                me = fight.fish1
            else:
                print('\nummmm.....')
            
            print('my size,opp size,my effo, opp effort, my wager, opp wager')
            print(x_size,o_size,x_eff,o_eff,x_wag,o_wag)
            print('perceived p_win',p_win)
            print('actual p_win for smaller investor',fight_.p_win)
            print('actual efforts:',me.effort,other_fish.effort)
        return p_win

    def _use_mutual_likelihood(self,fight,win=True):
        #import pdb; pdb.set_trace()
        if self.likelihood_dict is None:
            #likelihood = self._define_likelihood_mutual(fight,win) 
            likelihood = self._define_likelihood_mut_array(fight,win)
        elif True or (fight.winner.idx,fight.loser.idx) not in self.likelihood_dict.keys():
            #likelihood = self._define_likelihood_mutual(fight,win) 
            likelihood = self._define_likelihood_mut_array(fight,win)

        else: ## The dict is fast, but it doesn't work, I would need every possible effort in there too...
            likelihood = self.likelihood_dict[fight.winner.idx,fight.loser.idx]
            if not win:
                likelihood = 1-likelihood
        return likelihood

    def _define_likelihood_mutual(self,fight,win=True):
        xs = self.xs
        likelihood = np.zeros(len(xs))
        if win:
            other_fish = fight.loser
        else:
            other_fish = fight.winner
        x_opp = other_fish.size
        if self.effort == 0:
            likelihood = np.ones(len(xs))
        if win:
            for s in range(len(xs)):
                if False:
                    likelihood[s] = self._likelihood_function_size(xs[s],x_opp)
                else:
                    likelihood[s] = self._likelihood_function_se(xs[s],other_fish,x_eff=self.effort,fight=fight)
        elif not win:
            for s in range(len(xs)):
                if False:
                    likelihood[s] = 1-self._likelihood_function_size(xs[s],x_opp)
                else:
                    likelihood[s] = 1- self._likelihood_function_se(xs[s],other_fish,x_eff=self.effort,fight=fight)
        return likelihood
   
## Wager function optimized for array multiplication
    def _wager_curve_smart(self,w,L=None): #L=np.tan((np.pi/2 - np.pi*-0.9)/2)):
        return (w ** self.params.L) / 2

## Updated likelihood function that *should* be faster
    def _define_likelihood_mut_array(self,fight,win=True):

        if fight is None:
            s,e,l = self.naive_params
        else:
            s,e,l = fight.outcome_params
        #shifted_params = (np.array([s,e,l]) + 1) / 2
        #S,F,L = np.tan(np.pi/2 - shifted_params*np.pi/2)
        S,F,L = fight.params.scaled_params
        #print(S,F,L)
        if l == -1: ## several weird edge cases here where you can't learn anything
            pass 
            #import pdb;pdb.set_trace()
            #return np.ones_like(self.xs)
        if s == 0:
            return np.ones_like(self.xs)
        #elif e == -1:
        #    return np.ones_like(self.xs)
        if self.effort == 0:
            pass
            x_eff = self.effort
            #return np.ones_like(self.xs)
        if self.effort == None:
            x_eff = 1
        else:
            x_eff = self.effort

## Define other fish size and effort
## I define this based on the win argument, so make sure that is correct
        if win:
            o_eff = fight.level 
        else:
            o_eff = 1 ## this is an assumption of how much the other fish would have fought.
        if self == fight.fish1:
            other_fish = fight.fish2
        else:
            other_fish = fight.fish1
        if self.insight == True:
            o_eff = other_fish.effort
        xs = self.xs ## for simplicity, although sometimes I forget it exists
        likelihood = np.empty_like(xs)

## Build arrays of rel sizes of each fish as focal fish increases size
## NOTE: This enforces post-acuity. How do I handle cases where this is a guess, or something?
        size_index = np.argmax(self.xs >= other_fish.size)
        rel_xs = np.ones_like(xs)
        rel_os = np.ones_like(xs)
        rel_xs[:size_index] = self.xs[:size_index] / other_fish.size
        rel_os[size_index:] = other_fish.size / self.xs[size_index:] 
## Get wager arrays based on relative sizes
        #x_wager = rel_xs ** S * x_eff ** F ## Wager of focal fish
        #o_wager = rel_os ** S * o_eff ** F ## Wager of opponent

        x_eff_rel = x_eff / max([x_eff,o_eff])
        o_eff_rel = o_eff / max([x_eff,o_eff])

        #print('in fish:',x_eff_rel,o_eff_rel)
        #import pdb;pdb.set_trace()
        #x_wager = fight._SE_func(rel_xs,x_eff)
        #o_wager = fight._SE_func(rel_os,o_eff)
        #x_wager = fight._SE_func(rel_xs,x_eff_rel)
        #o_wager = fight._SE_func(rel_os,o_eff_rel)
        x_wager = fight._SE_func(self.xs / 100,x_eff)
        o_wager = fight._SE_func(other_fish.size / 100,o_eff)
        #import pdb;pdb.set_trace()
## Build relative wager array
        wager_array = np.empty_like(xs)
        if x_wager[-1] <= o_wager:
            wager_index = len(x_wager) ## this deals with the case where x_wager is never bigger
## NOTE: 
            wager_array = x_wager / o_wager
        else:
            wager_index = np.argmax(x_wager > o_wager) ## the point x_wager becomes bigger
            #wager_array[:wager_index] = x_wager[:wager_index] / o_wager[:wager_index]
            #wager_array[wager_index:] = o_wager[wager_index:] / x_wager[wager_index:]
        if True:
            wager_array[:wager_index] = x_wager[:wager_index] / o_wager
            wager_array[wager_index:] = o_wager / x_wager[wager_index:]
            #wager_array[:size_index] = x_wager[:size_index]
            #wager_array[size_index:] = o_wager[size_index:]
        #L = np.tan((np.pi - l)/2) ## calculate this once to speed things up
        L = self.params.L
        #likelihood = fight._wager_curve_smart(wager_array,L)
        likelihood = fight._wager_func(wager_array,L)
        if np.isnan(likelihood[wager_index-1]):
            #import pdb;pdb.set_trace()
            likelihood[wager_index-1] = 0.5            

        #likelihood = self._wager_curve_smart(wager_array,L)
        if win: ## since likelihood is the probability of what happened, and wager_array was p(upset)
#NOTE::
            likelihood[wager_index:] = 1 - likelihood[wager_index:]
            #likelihood[size_index:] = 1 - likelihood[size_index:]
             
        else:
            likelihood[:wager_index] = 1 - likelihood[:wager_index]
            #likelihood[:size_index] = 1 - likelihood[:size_index]
## PDB

        return likelihood

## This also includes opponent assesment...need to fix that
    def update_prior_old(self,win,x_opp=False,xs=None,w_opp=None,e_self=None,outcome_params=None):
        if xs is None:
            xs = self.xs

## Get likelihood function (there are a couple possible methods)
        if x_opp == False:
            likelihood = self._define_likelihood_w(outcome_params,win)
        else:
            likelihood = self._define_likelihood(fight,win)
        #self.win_record.append([x_opp,win])
        self.win_record.append([x_opp,win,self.effort])
        self.prior = self._update(self.prior,likelihood,xs)
        self.cdf_prior = self._get_cdf_prior(self.prior)

        if False: ## Need to decide which of these to use...
            estimate = self.xs[np.argmax(self.prior)] ## This is easy, but should it be the mean?
        else:
            estimate = np.sum(self.prior * self.xs / np.sum(self.prior))
        self.estimate_ = np.sum(self.prior * self.xs / np.sum(self.prior))
        
        prior_mean,prior_std = self.get_stats()
        self.est_record_.append(prior_mean)
        self.sdest_record.append(prior_std)
        
        self.estimate = estimate
        self.est_record.append(estimate)
        
        return self.prior,self.estimate
   
    def calculate_cost(self,win,fight,other_fish):
        if not other_fish.alive:
            cost = 0
        elif not win:
            cost = self.effort
        elif self.size >= other_fish.size: 
            #cost = min([fight.fish1.wager,fight.fish2.wager])
            cost = min([self.effort,other_fish.effort]) 
        else: ## if a smaller fish wins, should we scale up effort?
            if self.effort == 0:
                cost = 0
            else:
                #cost = np.nanmin([self.effort,other_fish.effort * (other_fish.wager / self.wager)])
                cost = min([self.effort,other_fish.effort])
        return cost

    def update_energy(self,win,fight):
        #print('in update energy:',fight)
        if win:
            other_fish = fight.loser
            cost = self.calculate_cost(win,fight,other_fish)
            if fight.food is not None:
                if fight.food > 0:
                    #print('pre energy:',self.energy,'cost:',cost)
                    #print('food:',fight.food,'free food',self.params.free_food)
                    self.energy = np.round(self.energy - cost + fight.food + self.params.free_food,2)
                    #print('post energy:',self.energy)
                    self.energy = np.clip(self.energy,0,self.max_energy)
                    self.size = self.size + self.r_rhp * (self.s_max - self.size) ** self.a_growth
                    #self.fitness_record.append(0)
                else:
                    self.energy = np.round(self.energy - cost + self.params.free_food,2)
                    self.energy = np.clip(self.energy,0,self.max_energy)
                    self.fitness_record.append(1)
        else:
            other_fish = fight.winner
            cost = self.effort
            if fight.food is not None:
                self.energy = np.round(self.energy - cost + self.params.free_food,2)
                self.energy = np.clip(self.energy,0,self.max_energy)
                if fight.food <= 0:
                    self.fitness_record.append(0)
        if self.energy_cost is False:
            print('#### RESTORING ENERGY #####')
            self.energy = self.max_energy 
        if self.energy <= 0:
            #print('I am dying!',fight.level,cost,self.energy,self.effort,self.idx,self.size,other_fish.size,self.params.mutant)
            self.energy = 0
            self.alive = False
            #print(self.energy_record)
        self.size_record.append(self.size)
        self.energy_record.append(self.energy)

        return other_fish,cost

## For testing what happens if you just don't update ever
    def no_update(self,win,fight):

        if False and self.params.effort_method == 'Perfect':
            if win:
                print('I won!')
            else:
                print('I lost??')
                print(fight.fish1.size,fight.fish2.size)
                print(fight.fish1.effort,fight.fish2.effort)
                print(fight.fish1.wager,fight.fish2.wager)
                print('calculated p_win:',fight.p_win,'\n')
        if self.energy_cost:
            pass
            #self.update_energy(win,fight)
        if win:
            other_fish = fight.loser
        else:
            other_fish = fight.winner
        cost = self.calculate_cost(win,fight,other_fish)
        self.win_record.append([other_fish.size,win,self.effort,cost])
        return self.prior,self.estimate

# Updates the prior and handles food and cost. Should probably be rebuilt eventually
    def update_prior(self,win,fight):
        size_idx = np.argmax(self.xs >= self.size)
        size_possible_pre = self.prior[size_idx] > 0
## Establish fishes and impose costs and benefits
        #if self.params.energy_cost:
        #    other_fish,cost = self.update_energy(win,fight) # This currently updates win record, not ideal
        if win:
            other_fish = fight.loser
        else:
            other_fish = fight.winner

        cost = self.calculate_cost(win,fight,other_fish)
## Get likelihood function
        #import pdb;pdb.set_trace()
        self.win_record.append([other_fish.size,win,self.effort,cost])
        if self.effort > other_fish.effort:
            pass
        if self.params.effort_method[1] == 0:
            likelihood = self._use_simple_likelihood(fight,win)
            i_estimate = np.argmax(self.xs > self.estimate)
        else:
            likelihood = self._use_mutual_likelihood(fight,win)
            #likelihood = self._define_likelihood_mutual(fight,win)
        self.likelihood = likelihood
        #print(likelihood[size_idx],fight.p_win)
        pre_prior = np.array(self.prior)
        likelihood_sum = np.sum(likelihood)
        if likelihood_sum == 0 or likelihood_sum == len(likelihood):
            ## If there's no information, don't update
            pass
        else:
            self.prior = self._update(self.prior,likelihood,self.xs) ## this just multiplies prior*likelihood

        self.cdf_prior = self._get_cdf_prior(self.prior)
        pre_estimate = self.estimate
        estimate_ = self.xs[np.argmax(self.prior)]
        prior_mean,prior_std = self.get_stats()
        estimate = prior_mean
        post_estimate = estimate

        self.estimate =  estimate
        self.est_record.append(estimate)
        if np.isnan(self.estimate):
            import pdb;pdb.set_trace()
        if True: ## most of the time I shouldn't need this stuff
            #self.est_record_.append(estimate_)

            self.sdest_record.append(prior_std)

            est_5,est_25,est_75,est_95 = self._get_range_prior(self.cdf_prior) 
            self.range_record.append([est_5,est_25,est_75,est_95])
            
            self.estimate_ = estimate_
            self.est_record_.append(estimate_)

        size_possible_post = self.prior[size_idx] > 0
        if self.idx == 0 and False:
            #plt.cla()
            fig,ax = plt.subplots()
            ax.plot(self.xs,pre_prior*100,color='blue')
            ax.plot(self.xs,self.likelihood,color='red')
            ax.plot(self.xs,self.prior*100,color='green')
            ax.axvline(self.estimate,color='green')
            ax.axvline(self.size,color='blue')
            ax.axvline(self.guess,color='black',linestyle=':')
            ax.axvline(other_fish.size,color='black')
            print(self.effort,other_fish.effort)
            print(self.size,other_fish.size)
            print(self.wager,other_fish.wager)
            print(fight.p_win)
            plt.show()
        return self.prior,self.estimate

    def update_blind(self,win,fight):
        size_idx = np.argmax(self.xs >= self.size)
        size_possible_pre = self.prior[size_idx] > 0
## Establish fishes and impose costs and benefits
        #if self.params.energy_cost:
        #    other_fish,cost = self.update_energy(win,fight) # This currently updates win record, not ideal
        if win:
            other_fish = fight.loser
        else:
            other_fish = fight.winner

        cost = self.calculate_cost(win,fight,other_fish)
## Get likelihood function
        #import pdb;pdb.set_trace()
        self.win_record.append([other_fish.size,win,self.effort,cost])

        likelihood = self._use_simple_likelihood(fight,win)
        i_estimate = np.argmax(self.xs > self.estimate)

        self.likelihood = likelihood

        pre_prior = np.array(self.prior)
        likelihood_sum = np.sum(likelihood)
        if likelihood_sum == 0 or likelihood_sum == len(likelihood):
            ## If there's no information, don't update
            pass
        else:
            self.prior = self._update(self.prior,likelihood,self.xs) ## this just multiplies prior*likelihood

        self.cdf_prior = self._get_cdf_prior(self.prior)
        pre_estimate = self.estimate
        estimate_ = self.xs[np.argmax(self.prior)]
        prior_mean,prior_std = self.get_stats()
        estimate = prior_mean
        post_estimate = estimate

        self.estimate =  estimate
        self.est_record.append(estimate)
        if np.isnan(self.estimate):
            import pdb;pdb.set_trace()
        if True: ## most of the time I shouldn't need this stuff
            #self.est_record_.append(estimate_)

            self.sdest_record.append(prior_std)

            est_5,est_25,est_75,est_95 = self._get_range_prior(self.cdf_prior) 
            self.range_record.append([est_5,est_25,est_75,est_95])
            
            estimate_ = self.xs[np.argmax(self.prior)]
            self.estimate_ = estimate_
            self.est_record_.append(estimate_)

        size_possible_post = self.prior[size_idx] > 0
        if self.idx == 0 and False:
            #plt.cla()
            fig,ax = plt.subplots()
            ax.plot(self.xs,pre_prior*100,color='blue')
            ax.plot(self.xs,self.likelihood,color='red')
            ax.plot(self.xs,self.prior*100,color='green')
            ax.axvline(self.estimate,color='green')
            ax.axvline(self.size,color='blue')
            ax.axvline(self.guess,color='black',linestyle=':')
            ax.axvline(other_fish.size,color='black')
            print(self.effort,other_fish.effort)
            print(self.size,other_fish.size)
            print(self.wager,other_fish.wager)
            print(fight.p_win)
            plt.show()
    
        return self.prior,self.estimate

## No fancy ratio of distance shift. Modeled more directly on Dugatkin. 
    def update_dugatkin(self,win,fight):
        m = self.params.poly_param_m
        b = self.params.poly_param_b

        if win:
            other_fish = fight.loser
            from_max = self.params.max_size - self.estimate
            new_estimate = self.estimate*(1+m) + b
            shift = new_estimate - self.estimate
        else:
            other_fish = fight.winner
            from_min = self.estimate - self.params.min_size
            shift = -1 * from_min*m - b
            new_estimate = self.estimate*(1-m) + b
            shift = new_estimate - self.estimate
        #print(win,shift)
        #new_estimate = self.estimate + shift
        new_estimate = np.clip(new_estimate,self.params.min_size,self.params.max_size)
## bit of a hack
        prior = norm.pdf(self.xs,new_estimate,self.params.A)
        self.estimate = new_estimate
        self.prior = prior / np.sum(prior)
        self.est_record.append(self.estimate)
        cost = self.calculate_cost(win,fight,other_fish)
        #cost = None
        self.win_record.append([other_fish.size,win,self.effort,cost])
        if True: ## most of the time I shouldn't need this stuff
            #self.est_record_.append(estimate_)

            prior_mean,prior_std = self.get_stats()
            self.sdest_record.append(prior_std)

            self.cdf_prior = self._get_cdf_prior(self.prior)
            est_5,est_25,est_75,est_95 = self._get_range_prior(self.cdf_prior) 
            self.range_record.append([est_5,est_25,est_75,est_95])
            
            estimate_ = self.xs[np.argmax(self.prior)]
            self.estimate_ = estimate_
            self.est_record_.append(estimate_)

        return self.prior,self.estimate

    def update_linear(self,win,fight):
        m = self.params.poly_param_m
        b = self.params.poly_param_b
        #b = 0
        #m = 0.1 

        if win:
            other_fish = fight.loser
            from_max = self.params.max_size - self.estimate
            shift = from_max*m + b
        else:
            other_fish = fight.winner
            from_min = self.estimate - self.params.min_size
            shift = -1 * from_min*m - b
        #print(win,shift)
        new_estimate = self.estimate + shift
        new_estimate = np.clip(new_estimate,self.params.min_size,self.params.max_size)
## bit of a hack
        prior = norm.pdf(self.xs,new_estimate,self.params.A)
        self.estimate = new_estimate
        self.prior = prior / np.sum(prior)
        self.est_record.append(self.estimate)
        cost = self.calculate_cost(win,fight,other_fish)
        #cost = None
        self.win_record.append([other_fish.size,win,self.effort,cost])

        if True: ## most of the time I shouldn't need this stuff
            #self.est_record_.append(estimate_)

            prior_mean,prior_std = self.get_stats()
            self.sdest_record.append(prior_std)

            self.cdf_prior = self._get_cdf_prior(self.prior)
            est_5,est_25,est_75,est_95 = self._get_range_prior(self.cdf_prior) 
            self.range_record.append([est_5,est_25,est_75,est_95])
            
            estimate_ = self.xs[np.argmax(self.prior)]
            self.estimate_ = estimate_
            self.est_record_.append(estimate_)
        return self.prior,self.estimate
  
    def plot_prior(self,ax=None):
        if ax is None:
            fig,ax = plt.subplots()
        ax.plot(self.xs,self.prior)
        if ax is None:
            fig.show()

    def summary(self,print_me): # print off a summary of the fish
        if print_me:
            print('My number is:',self.idx)
            print('Actual size:',self.size)
            print('Size estimate:',self.estimate)
            print('Win record:',self.win_record)
        return self.name,self.size,self.estimate,self.win_record

    def scale_effort(self,x,a=15,b=1.0,h=None):
        h = self.params.outcome_params[1]  
        eff = b * (np.tanh(a*(x-h))/2 + 1/2)
        eff = np.clip(eff,0,1)
        return eff

    def poly_effort_combo(self,f_opp,fight):
        #order = 1
        #print(f_opp.size,self.params.C,self.params.min_size,self.params.max_size)
        #import pdb;pdb.set_trace()
        if self.params.acuity == -1 or self.params.acuity == 0:
            if self.params.acuity == 0:
                opp_size_guess = f_opp.size
            else:
                opp_size_guess = np.random.random() * 100
        else:
            #opp_size_guess = np.clip(np.random.normal(f_opp.size,self.params.C),self.params.min_size,self.params.max_size)
            opp_size_guess = f_opp.assess_me(self)

            #mu,scale = f_opp.size,self.params.C
            #a_min,b_max = self.params.min_size,self.params.max_size
            #a,b = (a_min - mu) / scale,(b_max - mu) / scale
            #opp_size_guess = truncnorm(a,b,loc=mu,scale=scale).rvs()

            #opp_size_guess = f_opp.trunced.rvs()

        opp_size_guess = np.clip(opp_size_guess,self.params.min_size,self.params.max_size)
        self.guess = opp_size_guess
        s,e,l = self.params.outcome_params
        #shifted_params = (np.array([s,e,l]) + 1) / 2
        #S,F,L = np.tan(np.pi/2 - shifted_params*np.pi/2)
        S,F = self.params.S,self.params.F
        L = self.params.L

        #import pdb;pdb.set_trace()
        if opp_size_guess > self.estimate:  ## if you guess you are SMALLER
            est_ratio = self.estimate / opp_size_guess

            #rough_wager = est_ratio ** S * self.energy ** F
            #rough_wager = fight._SE_func(est_ratio,self.energy)
            rough_wager = fight._SE_func(self.estimate / 100,self.energy) / fight._SE_func(opp_size_guess / 100,self.energy)
            #print(self.estimate,opp_size_guess,rough_wager)
            #effort = fight._wager_curve(rough_wager)
            if rough_wager == 1: ## this is relative wager, if 1, effort = 0.5
                effort = 0.5
            else:
                effort = fight._wager_func(rough_wager)
            #effort = (rough_wager ** self.params.poly_param_a)/2
        else: ## if you think you're bigger
            est_ratio = opp_size_guess / self.estimate

            #rough_wager = est_ratio ** S * self.energy ** F
            #rough_wager = fight._SE_func(est_ratio,self.energy)
            rough_wager = fight._SE_func(opp_size_guess / 100,self.energy) / fight._SE_func(self.estimate/100,self.energy)
            #effort = 1 - (rough_wager ** self.params.poly_param_a)/2
            #effort = 1- fight._wager_curve(rough_wager)
            if rough_wager >= 1:
                effort = 0.5
            else:
                effort = 1 - fight._wager_func(rough_wager)
        effort = self._scale_func(effort)

        scaled_effort = effort 
        #scaled_effort = (effort * confidence_correction) ** self.params.B
        if np.isnan(effort):
            print('probably breaking:')
            print(effort)
            print(rough_wager)
            print(opp_size_guess,self.energy,self.estimate)
            print('going to just pretend effort was 0')
            #import pdb;pdb.set_trace()
            scaled_effort = 0

        scaled_effort = np.clip(scaled_effort,0,1)
        if self.params.print_me:
            print('###',self.idx,self.params.effort_method)
            print(self.params.poly_param_a,rough_wager)
            print(effort,scaled_effort,confidence_correction)
            print(self.size,self.estimate,self.awareness,self.params.A)
            print(f_opp.size,self.guess,self.acuity,self.params.C)
        return scaled_effort * self.energy

## The only difference is that here they happen to guess the correct size of their opponent
    def poly_effort_lucky(self,f_opp,fight):
        opp_size_guess = f_opp.size
        self.guess = opp_size_guess
        s,e,l = self.params.outcome_params
        S,F = self.params.S,self.params.F

        if opp_size_guess > self.estimate:  ## if you guess you are SMALLER
            est_ratio = self.estimate / opp_size_guess

            rough_wager = est_ratio ** S * self.energy ** F
            effort = fight._wager_curve(rough_wager)
        else: ## if you think you're bigger
            est_ratio = opp_size_guess / self.estimate
            rough_wager = est_ratio ** S * self.energy ** F
            effort = 1- fight._wager_curve(rough_wager)

        if self.params.C == 0 and self.prior_std == 0:
            confidence_correction = 1
        else:
## ideally this would be probabiliy of being wrong, but error will have to do.
            confidence_correction = 1 - norm.cdf(-0.1 * self.guess,self.estimate - self.guess,self.params.C + self.prior_std)
        self.correction = confidence_correction

        scaled_effort = (effort * confidence_correction) ** self.params.B
        scaled_effort = np.clip(scaled_effort,0,1)
        if self.params.print_me:
            print('###',self.idx,self.params.effort_method)
            print(self.params.poly_param_a,rough_wager)
            print(effort,scaled_effort,confidence_correction)
            print(self.size,self.estimate,self.awareness,self.params.A)
            print(f_opp.size,self.guess,self.acuity,self.params.C)
        return scaled_effort * self.energy

    def choose_effort_energy(self,f_opp,fight,strategy=None):
        effort = self.choose_effort(f_opp,fight,strategy)
## A slightly more careful strategy, invests a proportion of available energy, should generally avoid death
        if True:
            effort = self.energy * (effort ** self.c_aversion)
            effort = np.clip(effort,0,self.energy)
        return effort

    def leroy_jenkins(self,f_opp,fight):
        effort= self.energy
        return effort

    def half_jenkins(self,f_opp,fight):
        effort = self.energy * .5
        return effort

    def float_jenkins(self,f_opp,fight):
        #import pdb;pdb.set_trace()
        effort = self.params.baseline_effort
        if not fight.food:
            #print('FOR QBERT!!')
            effort = np.cbrt(effort)
            pass
        return effort

    def random_effort(self,f_opp,fight):
        effort = np.random.random() * self.energy
        return effort

    ## Function to estimate opponent size, should return a prior distribution of opponent size like own prior_cdf
    def estimate_opponent(self,f_opp,fight):
        return f_opp.cdf_prior

    # Check whether you believe you are bigger with some required confidence
    def probably_bigger(self,x_opp,conf=.5):
        cutoff = self.xs[np.argmax(self.cdf_prior > (1-conf))] ## Calculate cutoff for a given confidence
        if x_opp < cutoff:
            return True
        else:
            return False

    def get_stats(self):
        if self.awareness == 0:
            prior_mean = self.size
            prior_std = 0
            self.prior_mean = prior_mean
            self.prior_std = prior_std
            return prior_mean,prior_std

        prior_mean = np.sum(self.prior * self.xs / np.sum(self.prior))
        prior_std = np.sum((self.prior * (self.xs - prior_mean)**2)) ## I need this for confidence correction
        prior_std = np.sqrt(prior_std)
        self.prior_mean = prior_mean
        self.prior_std = prior_std
        
        return prior_mean,prior_std

    def copy(self):
        self.params.prior = np.array(self.prior)
        f = Fish(self.idx,self.params)
        return f

## Simplified fish object to decrease overhead when desired
class FishNPC(Fish):
    def __init__(self,idx=0,params=None,prior=None,likelihood=None,likelihhood_dict=None):
        self.idx = idx
        self.name = 'NPC '+str(idx) 
        self.alive = True
        self.trunced = None

        if params is not None:
            self.params = params.copy()
        else:
            self.params = Params()

        if self.params.size is not None:
            if self.params.size == 0:   
                self.size = self._growth_func(self.params.age)
            else:
                self.size = self.params.size
        else:
            self.size = np.random.normal(self.params.mean_size,self.params.sd_size)
        self.size = np.clip(self.size,self.params.min_size,self.params.max_size)
        self.params.size = self.size

        if self.params.baseline_effort == 0 or self.params.baseline_effort is None:
            self.params.baseline_effort = np.random.random()
        self.effort = self.params.baseline_effort

        self.energy = 1
        self.max_energy = 1
        self._choose_effort = self.float_jenkins
        self.update = self.empty_func
        self.no_update = self.empty_func
    
    def build_trunc(self,size,C):
        mu = size
        a = (self.params.min_size - size) / C
        b = (self.params.max_size - size) / C
        X = truncnorm(a,b,loc=mu,scale=C)
        return X

    def assess_me(self,opp):
        if opp.params.C != self.params.C:
            temp_trunc = self.build_trunc(self.size,opp.params.C)
        elif self.trunced == None:
            self.trunced = self.build_trunc(self.size,self.params.C)
            temp_trunc = self.trunced
        else:
            temp_trunc = self.trunced
        return temp_trunc.rvs()

    def empty_func(self,*args):
        return None

## This doesn't work great, something about the module setup.
if __name__ == '__main__':
    from fight import Fight

    f1 = Fish() 
    f2 = FishNPC()
    fight = Fight(f1,f2)
    fight.run_outcome()

    print(f1.idx,f1.size,len(f1.prior))
    print(f2.idx,f2.size)
    print(f2._choose_effort(f1,fight))
    print(f2.update(True,fight))
    print(f1._choose_effort(f1,fight))
    f1.update(True,fight)
    print(f1.estimate)
```
