## Supplemental Code for "Bayesian updating for self-assessment explains social dominance and winner-loser effects": params.py

```
## Load required packages
import numpy as np
import copy

class Params():
    __slots__ = ('n_iterations', 'iterations', 'print_me', 'mutant_effort', 'mutant_update', 'mutant_prior', 'n_fish', 'n_npcs', 'n_rounds', 'f_method', 'mean_age', 'min_size', 'max_size', 'mean_size', 'sd_size', 'energy_refill', 'energy_cost', 'n_fights', 'fitness_ratio', 'death', 'food', 'free_food', 'f_outcome', 'outcome_params', 'scaled_params', 'S', 'F', 'L', 'L_set', 'outcome', 'effort_method', 'effort_exploration', 'baseline_effort', 'update_method', 'age', 'size', 'prior', 'likelihood', 'likelihood_dict', 'xs', 'r_rhp', 'a_growth', 'c_aversion', 'max_energy', 'start_energy', 'acuity', 'pre_acuity', 'post_acuity', 'awareness', 'insight', 'poly_param_a', 'poly_param_b', 'poly_param_c','boldness','A','B','C','poly_param_m', 'poly_step', 'verbose', 'mutant','S_set','F_set','A_set','B_set','C_set','sigmoid_params','K','k','m','a','fig_font')
    def __init__(self,iterations=1000,print_me=False,     ## Sim Params
                n_fish=5,n_rounds=20,f_method='shuffled',    ## Tank params
                energy_refill=0.5,energy_cost=False,n_fights=10,
                fitness_ratio=None,death=False,food=0.5,free_food=0,
                mean_age=None,mean_size=None,sd_size=None,
                min_size = 1,max_size = 100,
                f_outcome='math',outcome_params=[0.7,0.5,-0.8], ## Fight Params
                sigmoid_params = [10,4,0.9], #k,a,m
                effort_method='SmoothPoly',baseline_effort=.5,update_method='bayes',  ## Fish Params
                age=50,size=None,prior=None,likelihood=None,likelihood_dict=None,
                xs=np.linspace(1,100,199),r_rhp=0,a_growth=True,c_aversion=1,
                max_energy=1,start_energy=1,effort_exploration=0.1,
                acuity=.1,pre_acuity=.1,post_acuity=True,awareness=.5,insight=True,
                boldness=0,
                #poly_param_a = 3,poly_param_b=-2.4,poly_param_c=0.1,
                poly_param_a = 5,poly_param_b=0,poly_param_c=0.3,poly_param_m=0.1,
                mutant_effort=[1,1],mutant_update='bayes',mutant_prior=None,
                fig_font = 10,
                verbose=False):                             ## Other params
## Sim params
        self.n_iterations = iterations ## Deprecated
        self.iterations = iterations
        self.print_me = print_me
        self.mutant_effort = mutant_effort
        self.mutant_update = mutant_update
        self.mutant_prior = mutant_prior
## Tank Params
        self.n_fish = n_fish
        self.n_npcs = 0
        self.n_rounds = n_rounds
        self.f_method = f_method ## this defines how much fish can pick their opponents
        self.mean_age = mean_age
        self.min_size = min_size 
        self.max_size = max_size
        if mean_size == None:
            if self.mean_age is not None:
                mean_size = self._growth_func(self.mean_age)
            else:
                mean_size = (self.max_size + self.min_size) / 2
        if sd_size == None:
            sd_size = mean_size/5
        self.mean_size,self.sd_size = mean_size,sd_size

        self.energy_refill=0.5
        self.energy_cost=energy_cost
        self.n_fights = n_fights
        self.fitness_ratio=fitness_ratio
        self.death=death
        self.food=food
        self.free_food=free_food

## Fight Params
        self.f_outcome = f_outcome ## This defines how fights are determined.
        self.outcome_params = np.array(outcome_params)   ## This determines how fights are settled, skill,effort,luck

        shifted_params = (self.outcome_params +1) / 2
        self.scaled_params = np.tan(np.array(np.pi/2 - shifted_params*np.pi/2))
        self.scaled_params = np.round(self.scaled_params,4)
        self.S,self.F,self.L = self.scaled_params
        #self.L = np.tan(np.pi/2 - outcome_params[2]*np.pi/2)
        self.L_set = False
        self.outcome = None

        self.sigmoid_params = sigmoid_params
        self.k,self.a,self.m = sigmoid_params
        self.K = 1 + np.exp(self.k*(self.m-1))

## Fish Params
        self.effort_method = effort_method     ## self-assessment vs opponent assessment, [1,1] is MA
        self.effort_exploration = effort_exploration
        self.baseline_effort = baseline_effort
        self.update_method = update_method     ## how individuals update their self assessment
        self.age = age
        self.size = size
        self.prior = prior
        self.likelihood=likelihood
        self.likelihood_dict=likelihood_dict
        self.xs=xs
        self.r_rhp = r_rhp
        self.a_growth=a_growth
        self.c_aversion=c_aversion
        self.max_energy = max_energy
        self.start_energy=start_energy
        self.acuity=acuity
        self.pre_acuity=pre_acuity
        self.post_acuity=post_acuity
        self.awareness=awareness
        self.insight=insight
        self.boldness = boldness

        self.A = np.tan(self.awareness*np.pi/2) * 20 ## this defines std at a = 0.5
        self.C = np.tan(self.acuity*np.pi/2) * 20
        shifted_boldness = (boldness + 1) / 2
        if self.boldness == 0:
            self.B = 1
        else:
            self.B = np.tan(np.pi/2 - shifted_boldness*np.pi/2) 

        self.poly_param_a = poly_param_a
        self.poly_param_b = poly_param_b
        self.poly_param_c = poly_param_c ## Currently scales up effort (i.e. boldness)
        self.poly_param_m = poly_param_m
        self.poly_step = 0.1

## General Params
        self.verbose=verbose
        self.mutant=False
        self.fig_font = fig_font 
    def copy(self):
        return copy.copy(self)

    def se_wager_function(self,rel_size,effort):    
        wager = rel_size ** self.S * effort ** self.F
        return wager

    def wager_function(self,rel_wager):
        p_win = (rel_wager ** self.L) /2
        return p_win

## (deprecated)
    def get_L(self):
        print('get_L is deprecated, use set_params')
        #self.L = np.tan((np.pi - np.pi*self.outcome_params[2])/2)
        self.set_params()

    def set_L(self):
        #self.L = np.tan((np.pi - np.pi*self.outcome_params[2])/2)
        #self.L_set = True
        print('set_L is deprecated, use set_params')
        self.set_params()

    def set_params(self):
        self.outcome_params = np.array(self.outcome_params)
        shifted_params = (self.outcome_params +1) / 2
        self.scaled_params = np.tan(np.array(np.pi/2 - shifted_params*np.pi/2))
        self.scaled_params = np.round(self.scaled_params,4)
        self.S,self.F,self.L = self.scaled_params
        self.S_set = True
        self.F_set = True
        self.L_set = True


        if self.boldness == 0:
            self.B = 1
        else:
            shifted_boldness = (self.boldness + 1) / 2
            self.B = np.tan(np.pi/2 - shifted_boldness*np.pi/2) 
        self.A = np.tan(self.awareness*np.pi/2) * 20
        self.C = np.tan(self.acuity*np.pi/2) * 20

        self.A_set = True
        self.B_set = True
        self.C_set = True

    def _mutate(self):
        self.effort_method = self.mutant_effort
        self.update_method = self.mutant_update
        self.prior = self.mutant_prior
        self.mutant = True

    def _growth_func(self,t):
        size = 1 + self.max_size - self.max_size/np.exp(t/100)
        return size
   
    def summary(self):
        print('Number iterations:',self.n_iterations)
        print('Number of Fish:',self.n_fish)
        print('Number of rounds:',self.n_rounds)
        print('Fight Selection Method:',self.f_method)
        print('Fight Outcome:',self.outcome_params)
        print('Effort Method:',self.effort_method)
        print('Update Method:',self.update_method)

if __name__ == "__main__":
    params = Params()
    print(params.outcome_params)
    print(params.S,params.F,params.L)
    s = 0
    e = 1
    l = 0
    params.outcome_params = [s,e,l]
    params.set_params()
    print(params.L)
    params.effort_method = [1,1]
    params.n_fights = 10*50
    params.n_iterations = 50
    params.n_fish = 7
    params.f_method = 'random' 
    
    params.summary()
```
