## Supplemental Code for "Bayesian updating for self-assessment explains social dominance and winner-loser effects": tank.py

```
## Load required packages
import numpy as np

import itertools,random

from scipy.special import rel_entr
from scipy.stats import mode
from scipy.stats import norm
from scipy.ndimage import gaussian_filter1d 
from sklearn.metrics import auc

import matplotlib.pyplot as plt
from matplotlib import cm

from tqdm import tqdm

from bayesbots import Fish,FishNPC
from bayesbots import Fight
from bayesbots import Params

class Tank():
    def __init__(self,fishes,params=None,fight_list = None,npc_params=None
                 ):
        if params is None:
            params = Params()
        self.params = params.copy()
        self.npc_params = npc_params
        self.fishes = fishes
        self.n_fish = len(fishes)
        self.sizes = [f.size for f in fishes]
        self.f_method = params.f_method
        self.f_outcome = params.f_outcome
        self.f_params = params.outcome_params
        self.u_method = params.update_method
        self.win_record = np.zeros([len(fishes),len(fishes)])
        self.fitness_ratio=params.fitness_ratio
        self.death = params.death
        self.food = params.food
        if fight_list is not None:
            self.fight_list = fight_list
        elif npc_params is not None:
            self.npc_params = npc_params
            self.fight_list = self.get_npc_fights(npc_params.n_npcs,params.n_fights)
        else:
            self.fight_list = self.get_matchups(self.f_method,self.f_outcome,params.n_rounds)

            self.n_fights = len(self.fight_list)
        if self.f_method == 'balanced' or self.f_method == 'shuffled':
            self.n_rounds = self.fight_list[-1].idx+1
        else:
            self.n_rounds = len(self.fight_list)
        self.history = np.zeros([self.n_rounds,len(fishes),len(fishes)])

## Frustratingly long table to calculate linearity
        self._applebys = {
            2:{0:1}, ## This is so that everything works
            3:{0:0.750},
            4:{0:0.375},
            5:{0:0.117},
            6:{0:0.022,
                1:0.051,
                2:0.120},
            7:{1:0.006,
                2:0.017,
                3:0.033,
                4:0.069,
                5:0.112},
            8:{4:0.006,
                5:0.011,
                6:0.023,
                7:0.037,
                8:0.063,
                9:0.094,
                10:0.153
            },
            9:{9:0.007,
                10:0.012,
                11:0.019,
                12:0.030,
                13:0.045,
                14:0.067,
                15:0.095,
                16:0.138
            },
            10:{16:0.008,
                17:0.012,
                18:0.018,
                19:0.026,
                20:0.038,
                21:0.052,
                22:0.073,
                23:0.097,
                24:0.131
            }
            }

    def get_npc_fights(self,n_npcs=0,n_fights=10):
        fight_list = []

        if n_npcs == 0:
            n_npcs = len(self.fishes)
        if self.npc_params.real_fish:
            self.npcs = [Fish(n,self.npc_params) for n in range(n_npcs)]
        else:
            self.npcs = [FishNPC(n,self.npc_params) for n in range(n_npcs)]
        npc_sizes = [n.size for n in self.npcs]
        fish_sizes = [f.size for f in self.fishes]
        #print('NPC vs Fish',np.mean(npc_sizes),np.mean(fish_sizes))
        for i in range(n_fights):
            for n in self.npcs:
                for f in self.fishes:
                    fight_list.append(Fight(f,n,self.params,idx=i))
        random.shuffle(fight_list)
        if self.fitness_ratio is not None:
            for i in range(0,len(fight_list),int(1/self.fitness_ratio)):
                fight_list[i].food = False
        return fight_list

    def get_matchups(self,f_method='shuffled',f_outcome='chance',n_fights=10):
## N fights for balanced and shuffled is actually the number of rounds
        fight_list = []
        #import pdb;pdb.set_trace()
        if f_method == 'balanced' or f_method == 'shuffled':
            short_list = []
            for i in range(n_fights):
                combs = list(itertools.combinations(self.fishes,2))
                if f_method == 'shuffled':
                    random.shuffle(combs)
                for f1,f2 in combs:
                    fight_list.append(Fight(f1,f2,self.params,idx=i)) ## So balanced is organized as rounds
        if f_method == 'random':
            combs = list(itertools.combinations(self.fishes,2))
            for i in range(n_fights):
                f1,f2 = random.choice(combs)
                fight_list.append(Fight(f1,f2,self.params,idx=i))
        if self.fitness_ratio is not None:
            for i in range(0,len(fight_list),int(1/self.fitness_ratio)):
                fight_list[i].food = False
        #print('n_fights:',len(fight_list))
        return fight_list

    def process_fight(self,fight): ## This works without returning because of how objects work in python
        fight.run_outcome()
        fight.winner.update(True,fight)
        fight.loser.update(False,fight)
        
        self.update_size(fight.winner,True)
        self.update_size(fight.loser,False)

        self.win_record[fight.winner.idx,fight.loser.idx] += 1
        self.history[fight.idx,fight.winner.idx,fight.loser.idx] = 1 ## Note, this works a bit different for 'random' and 'balanced'
        return 0

    def update_size(self,fish,win):
        if self.params.r_rhp == 0 and self.params.energy_cost == False:
            return 0
        size = fish.size
        if win:
            size = size + self.params.r_rhp
        if self.params.energy_cost is True:
            size = size - fish.effort
        elif self.params.energy_cost != False:
            fish = fish - self.params.energy_cost
        fish.size = np.clip(size,1,100)
        fish.size_record.append(fish.size)

    def print_status(self):
        for f in self.fishes:
            print(f.name,':','size=',np.round(f.size,3),'estimate=',np.round(f.estimate,3))

    def run_all(self,progress=True,print_me=False,plot_stuff=False,cutoff=None):
        if cutoff is None:
            cutoff = len(self.fight_list)
        if plot_stuff:
            fig,ax = plt.subplots()
        if print_me:
            print("Starting fights!")
            print("Initial status_____")
            self.print_status()
        if self.u_method == 'hock':
            process = self.process_hock
        else:
            process = self.process_fight
        if progress:
            iterator = tqdm(range(cutoff))
        else:
            iterator = range(cutoff)
        for i in iterator:
            c = self.fight_list[i]
            process(c)
            if plot_stuff:
                if c.fish1.idx == 0:
                    ax.plot(c.fish1.naive_likelihood,alpha=.2)

            if print_me:
                print('UPDATE:_____')
                print(c.summary())
                #self.print_status()
        if plot_stuff:
            return fig,ax

    def plot_estimates(self,fish_list=None,food=False,legend=False):
        fig,ax = plt.subplots()
        if fish_list is None:
            fish_list = self.fishes
        for i in range(len(fish_list)):
            if len(fish_list) > 10:
                color = cm.viridis(i/len(fish_list))
            else:
                color = cm.tab10(i)
            f = fish_list[i]
            ax.plot(f.est_record, color=color,label=str(i))
            if food:
                ax.plot(f.size_record,color=color,linestyle=':')
            else:
                ax.axhline(f.size,color=color,linestyle=':')
            if self.u_method == 'bayes':
                f.range_record = np.array(f.range_record)
                ax.fill_between(np.arange(len(f.range_record)),f.range_record[:,1],f.range_record[:,2],color=color,alpha=0.2)
            ax.set_ylabel('Estimate')
        ax.set_xlabel('contest number')
        if legend:
            ax.legend()
        fig.show()
        return fig,ax

## Similar to above, but here simply plot effort
    def plot_effort(self,fish_list=None):
        fig,ax = plt.subplots()
        if fish_list is None:
            fish_list = self.fishes
        for i in range(len(fish_list)):
            if len(fish_list) > 10:
                color= cm.viridis(i/len(fish_list))
            else:
                color=cm.tab10(i)
            f = fish_list[i]
            effort_record = np.array(f.win_record)[:,-1] ## note that there are fish win records and tank win records. yikes.
            smooth_effort = effort_record
            #smooth_effort = gaussian_filter1d(effort_record,3)
            ax.plot(smooth_effort,color=color,alpha=.5,label=str(i))
        ax.legend()
        fig.show()
        return fig,ax

    def _calc_linearity(self,idx=None): ## idx is a slice object
        n_fish = len(self.fishes)
        h_matrix = np.zeros([n_fish,n_fish])
        if idx is not None:
            win_record = np.nansum(self.history[idx],axis=0)
        else:
            win_record = self.win_record
        win_record_dif = win_record - np.transpose(win_record)
        h_matrix[win_record_dif > 0] = 1
        self.h_matrix = h_matrix
        ## DO THE MATHY THING HERE
        N = n_fish

        D = N * (N-1) * (N-2) / 6 ## Total number of possible triads
## Calculate the number of triads: 
        d = N * (N-1) * (2*N-1) / 12 - 1/2 * np.sum(np.sum(h_matrix,1) ** 2) ## From Appleby, 1983
        if N <= 10:
            if d in self._applebys[N].keys():
                p = self._applebys[N][round(d)]
            elif d < min(self._applebys[N].keys()):
                p = min(self._applebys[N].values())
            else:
                p = max(self._applebys[N].values())
        else:
            df = N*(N-1)*(N-2)/(N-4)**2
            chi_stat = (8/(N-4)) * ((N*(N-1)*(N-2)/24) - d + 0.5) + df
            p = 1 - chi2.cdf(chi_stat,df)
        linearity = 1 - (d / D) ## Percentage of non triadic interactions
        #import pdb;pdb.set_trace()
        return linearity,[d,p]
         
## Stability the proportion of interactions consistent with the overall mean. 
    def _calc_stability(self,idx = None):
## A nicer metric would be the proportion of bins where mean heirarchy == overall hierarchy, 
        #import pdb;pdb.set_trace()
        if self.f_method == 'balanced' or self.f_method == 'shuffled':
            binned_history = self.history
        else: ## Something feels wrong here... 
            ## First calculate a sliding window bigger than 2*n^2. We're going to have some missing values
            min_slide = 2*self.n_fish*(self.n_fish-1)
            n_points = len(self.history)
            stagger = 2 # determines the degree to which windows overlap
            n_bins = int(n_points / min_slide * stagger)
            win_size = int(n_points / n_bins)
            binned_history = np.zeros([n_bins,self.n_fish,self.n_fish])
## There might be a more efficient way to do this, but this shoudl work.
            for w in np.arange(n_bins):
                h0 = w*win_size
                h1 = (w+1)*win_size
                binned_history[w] = np.sum(self.history[h0:h1],0)

# Instead, calculate the proportion of binned interactions that = overall interactions
        if idx is not None:
            binned_history = binned_history[idx]
            mean_history = np.mean(self.history[idx],0)
        else:
            mean_history = np.mean(self.history,0)
        binary_bins = np.sign(binned_history - np.transpose(binned_history,axes=[0,2,1]))
        binary_final = np.sign(mean_history - np.transpose(mean_history))
        binary_mean = np.mean(binary_bins,0)
        proportion_consistent = np.sum(np.abs(binary_mean) == 1) / (self.n_fish * (self.n_fish - 1))
## Use nCr formulat to get the total number of possible interactions
        total_interactions = len(binary_bins) * self.n_fish * (self.n_fish-1) 
        binary_difference = np.abs(binary_bins - binary_final) == 2
        number_consistent = total_interactions - np.sum(binary_difference)
        return proportion_consistent, binary_final
    
    def _calc_error(self):
        n_fishes = len(self.fishes)
        n_fights = len(self.fishes[0].range_record)
        e_array = np.empty([n_fishes,n_fights])
        for f_ in range(n_fishes):
            fish_f = self.fishes[f_]
            for r_ in range(n_fights):
                fight_r = fish_f.range_record[r_]
                e1,e2 = fight_r[1:3]
                f_error = (e1 - fish_f.size)**2 + (e2 - fish_f.size)**2
                e_array[f_,r_] = f_error
        return e_array

    def __getitem__(self,idx):
        return self.fishes[idx]
    

if __name__ == '__main__':
    fishes = [Fish(),Fish()]
    t = Tank(fishes)
    t.run_all()
    print(np.shape(t.win_record))
```
