## Supplemental Code for "Bayesian updating for self-assessment explains social dominance and winner-loser effects": figure2a_compareStrength.py

```
#! /usr/bin/env python

## Code for fig 3b, showing the loser effect is stronger

import numpy as np
from matplotlib import pyplot as plt

from bayesbots import Fish,FishNPC
from bayesbots import Fight
from bayesbots import Params

from tqdm import tqdm


iterations = 1000
params = Params()

plt.rcParams.update({'font.size': params.fig_font})

#params.outcome_params = [0.7,0,-0.8]
#params.outcome_params = [0.9,0,-0.8]
#params.awareness =  0.8

#params.prior = True
#params.awareness = 0.5
#params.acuity = 0.0
print(params.awareness)
params.set_params()
print(params.outcome_params)
params.size = 50

#assay_fish = Fish(1,params)


assay_params = params.copy()
assay_params.baseline_effort = 0.535
assay_params.prior = True

outcome_array = np.empty([iterations,2])
outcome_array.fill(np.nan)

win_info_array = np.array(outcome_array)
loss_info_array = np.array(outcome_array)
#iterations = 1000
w_probs,l_probs = [],[]

f_sizes = []

for i in tqdm(range(iterations)):
    focal_winner = Fish(i+2,params)
    f_sizes.append(focal_winner.size)
    focal_loser = focal_winner.copy() 
## Stage a bunch of wins and losses against size-matched fish

    #assay_params.size = focal_winner.size
    staged_opp = FishNPC(0,assay_params)
    staged_opp.size = focal_winner.size
    
    staged_win = Fight(staged_opp,focal_winner,params,outcome=1)
    staged_win.run_outcome()
    focal_winner.update(True,staged_win)

    staged_loss = Fight(staged_opp,focal_loser,params,outcome=0)
    staged_loss.run_outcome()
    focal_loser.update(False,staged_loss)
    #print(focal_winner.effort,focal_loser.effort)
    #print(focal_winner.estimate,focal_loser.estimate)
    #import pdb;pdb.set_trace()
    #if focal_loser.estimate > 50:
    #    import pdb;pdb.set_trace()
    #print('## staged opp size:',staged_opp.size)

## Assay against size matched fish
    assay_params.size = focal_winner.size
    assay_fish = Fish(1,assay_params)

    assay_winner = Fight(assay_fish,focal_winner,params)
    winner_output = assay_winner.run_outcome()
    outcome_array[i,1] = winner_output
    
    if assay_fish.wager > focal_winner.wager:
        w_probs.append(assay_winner.p_win)
    else:
        w_probs.append(1-assay_winner.p_win)
    #print('## Assay fish size:',assay_fish.size,assay_fish.estimate)
    #print('win:',assay_fish.effort,focal_winner.estimate,focal_winner.effort,assay_winner.p_win,winner_output) 
    assay_loser = Fight(assay_fish,focal_loser,params)
    loser_output = assay_loser.run_outcome()

    outcome_array[i,0] = loser_output
    win_info_array[i] = focal_winner.effort,focal_winner.estimate 
    loss_info_array[i] = focal_loser.effort,focal_loser.estimate 
    #print('loss:',assay_fish.effort,focal_loser.estimate,focal_loser.effort,assay_loser.p_win,loser_output)
    if assay_fish.wager > focal_loser.wager:
        l_probs.append(assay_loser.p_win)
    else:
        l_probs.append(1-assay_loser.p_win)
    #break
print(np.mean(outcome_array,axis=0))
print(np.std(outcome_array,axis=0))
print('probs:',np.mean(w_probs),np.mean(l_probs))
means = np.mean(outcome_array,axis=0)-0.5
means_ = -1 * means
sems = np.std(outcome_array,axis=0) / np.sqrt(iterations)

fig,ax = plt.subplots()

ax.bar([0,1],means,yerr = sems,bottom = 0.5,color = ['darkblue','gold'])
#ax.bar([0,1],means_,yerr = sems,bottom = 0.5,color = ['darkblue','gold'],alpha=0.1)
ax.axhline(0.5,color='black',linestyle=':')

ax.set_xticks([0,1])
ax.set_xticklabels(['Post-loss','Post-win'])
ax.set_xlabel('_')
#ax.set_ylabel('Probability of win\n vs size-matched fish')
ax.set_ylabel('Assay win rate')
ax.set_ylim([0,1])
plt.yticks(fontsize=int(params.fig_font * 3/4))

#fig.savefig('./figures/fig3b_winvloss.png',dpi=300)
print('win effort, win estimate')
print(np.mean(win_info_array,axis=0))
print(np.mean(loss_info_array,axis=0))
fig.set_size_inches(2.2,2.2)
fig.tight_layout()

fig.savefig('./figures/fig2a_WinVLoss.png',dpi=300)
fig.savefig('./figures/fig2a_WinVLoss.svg')
if False:
    plt.show()
```
