## Supplemental Code for "Bayesian updating for self-assessment explains social dominance and winner-loser effects": figure2b_pathDependence.py

```
#! /usr/bin/env python

## Script to show that under bayes', the duration of the winner effect depends on the about of new info 

from bayesbots import Fish
from bayesbots import Fight
from bayesbots import Tank
from bayesbots import Params

import numpy as np
#from scipy.stats import binom_test

from matplotlib import pyplot as plt
from matplotlib import cm

from scipy.ndimage import gaussian_filter1d
from tqdm import tqdm

import random, copy
import itertools


#s,e,l = .6,.3,.1

#s,e,l = -0.5,0.5,-0.9
s= 0.5
e= 0.7
l= -0.8

params = Params()
plt.rcParams.update({'font.size': params.fig_font})

params.acuity = 0
#params.boldness = 0.7
if True:
    params.effort_method = 'SmoothPoly'
else:
    params.effort_method = [0,.5]

#params.poly_param_a = 2
#params.n_fights = 50 
params.energy_cost = False
params.awareness = 0.5
params.prior=True
#params.start_energy = 1

params.iterations = 2000
params.n_rounds = 6


params.n_fish = 5
#params.f_method = 'balanced'
#params.u_method = 'bayes'
params.f_outcome = 'math'
params.outcome_params = [s,e,l]
params.set_params()

winners,losers = [],[]
PLOT = 'estimate'

## Have a fish win-lose, or lose-win, then measure the winner effect for that fish
scale = 2

upset_count = 0
win_count1,win_count2 = 0,0
win_shifts,loss_shifts = [],[]
## Lets assume their estimate is correct.

params.age = 51
params.size = 50
## Some helpful functions:
## Build copies of f0 with naive priors
def build_fish(idx,f0):
    return Fish(idx,f0.params)

def check_success(f,f_match,params):
    fight = Fight(f,f_match,params)
    fight.run_outcome()
    return fight,fight.outcome

## Tidy function to build a dict for storing everything
def build_results(n_matches=1):
    match_results = {}
    for d in range(n_matches):
        combos = [''.join(r) for r in itertools.product(['w','l'],repeat=d)]
        for c in combos:
            match_results[c] = []
    return match_results

## Dict to convert from outcome to letters
conversion_dict = {0:'w',1:'l'}

f0 = Fish(1,params)
fishes = []
match_results = build_results(params.n_rounds)
for i in range(params.iterations):
#for i in tqdm(range(iterations)):
    results_str = ''
    f = build_fish(1,f0)
    f_match = build_fish(0,f0)

    ## Run all matches
    for m in range(params.n_rounds):
        match,outcome =check_success(f,f_match,params)
        f.update(1-outcome,match)
        match_results[results_str].append(1-outcome)
        results_str += conversion_dict[outcome]
    fishes.append(f)
print('round one average:',np.mean(match_results['']))
print('round two average:')
print('winners:',np.mean(match_results['w']))
print('losers:',np.mean(match_results['l']))
print('...')
fig,ax = plt.subplots()

loser_estimates, winner_estimates = [],[]
final_estimates = []

for f in fishes:
    jitter = np.random.randn() * 1
    jitter = 0
    ax.plot(np.array(f.est_record)[:] + jitter,alpha=.02,color='black',linewidth=1)
    if False:
        if f.win_record[0][1] == 1: ## If you won the first fight
            if f.win_record[1][1] == 0: # but lost the second fight
                winner_estimates.append(f.est_record[2])
        elif f.win_record[1][1] == 1: # vs if you lost the first fight and won the second fight
            loser_estimates.append(f.est_record[2])
    if True: ## Get estimates after the first fight
        if f.win_record[0][1] == 1: ## if you won the first fight:
            winner_estimates.append(f.est_record[1])
        else:
            loser_estimates.append(f.est_record[1])
        final_estimates.append(f.est_record[-1])
[ax.axvline(f,color='black',linewidth=1,linestyle=':') for f in range(params.n_rounds)]

if False:
    y_max = max(winner_estimates)
    y_min = min(loser_estimates)
else:
    y_max = max(final_estimates)
    y_min = min(final_estimates)

for r in range(params.n_rounds):
    win_key = 'w' * r
    r_winners = match_results[win_key]
    win_str = str(np.round(np.mean(r_winners),2)) + ' +/- ' + str(np.round(np.std(r_winners) / np.sqrt(len(r_winners)),3))

    loss_key = 'l' * r
    r_losers = match_results[loss_key]
    loss_str = str(np.round(np.mean(r_losers),2)) + ' +/- ' + str(np.round(np.std(r_losers) / np.sqrt(len(r_winners)),3))

    #ax.text(r+0.01,y_max+1,win_str)
    #ax.text(r+0.01,y_min-1,loss_str)

ax.set_xticks(list(range(params.n_rounds)))
ax.set_xlim([-0.1,params.n_rounds - 0.5])
ax.set_ylabel('Estimate')
ax.set_xlabel('Contest round')

#ax.set_ylim([y_min - 1.5,y_max+1.5])
ax.set_ylim([y_min - 1.5,y_max+1.5])
#ax.set_title('Fish estimates are path dependent and have recency bias')
plt.xticks(fontsize=int(params.fig_font * 3/4))
plt.yticks(fontsize=int(params.fig_font * 3/4))

fig.set_size_inches(2.2,2.2)
fig.tight_layout()
fig.savefig('./figures/fig2b_pathDep.png',dpi=300)
fig.savefig('./figures/fig2b_pathDep.svg')
if False:
    fig.show()
    plt.show()
```
