## Supplemental Code for "Bayesian updating for self-assessment explains social dominance and winner-loser effects": figure3_costAccuracy.py

```
#! /usr/bin/env python

## Trying something new:
""" 
This script contains all the code for plotting figure 1 of Ammon's BayesBattleBots paper
For questions, contact Ammon Perkes
"""

import numpy as np
from matplotlib import pyplot as plt
import copy
from tqdm import tqdm

from bayesbots import Fish
from bayesbots import Fight
from bayesbots import Tank
from bayesbots import Params

from matplotlib import cm


## Define some global variables to determine if you will plot and save figures.
#PLOT = True

params = Params()
plt.rcParams.update({'font.size': params.fig_font})

params.set_params()

params.n_fish = 5
params.n_rounds = 30
params.iterations = 1000
#params.n_rounds = params.n_fights * (params.n_fish-1)

## Set up a tank
cost_array = np.empty([params.iterations,params.n_fish,params.n_rounds * (params.n_fish-1)])
error_array = np.empty([params.iterations,params.n_fish,params.n_rounds * (params.n_fish - 1) + 1])

cost_array_linear = np.empty_like(cost_array)
error_array_linear = np.empty_like(error_array)

cost_array_null = np.empty_like(cost_array)
error_array_null = np.empty_like(error_array)

linear_params = copy.deepcopy(params)
linear_params.update_method = 'linear'

null_params = copy.deepcopy(params)
null_params.update_method = None

for i in tqdm(range(params.iterations)):
    fishes = [Fish(f,params) for f in range(5)]
    tank = Tank(fishes,params)
    tank.run_all(progress=False)

    null_fishes = [Fish(f,null_params) for f in range(5)]
    null_tank = Tank(null_fishes,null_params)
    null_tank.run_all(progress=False)

    linear_fishes = [Fish(f,linear_params) for f in range(5)]
    linear_tank = Tank(linear_fishes,linear_params)
    linear_tank.run_all(progress=False)

    error_array[i] = tank._calc_error()
    error_array_null[i] = null_tank._calc_error()
    error_array_linear[i] = linear_tank._calc_error()
    for f in range(len(fishes)):
        cost_array[i,f] = np.array(tank.fishes[f].win_record)[:,3]
        cost_array_null[i,f] = np.array(null_tank.fishes[f].win_record)[:,3]
        cost_array_linear[i,f] = np.array(linear_tank.fishes[f].win_record)[:,3]
    
print('mean cost for bayes vs linear, +- SEM')
print(np.mean(cost_array > 0.8),np.std(cost_array > 0.8)/np.sqrt(params.iterations))
print(np.mean(cost_array_linear > 0.8),np.std(cost_array_linear > 0.8)/np.sqrt(params.iterations))

fig,(ax,ax1) = plt.subplots(1,2)

mean_int = np.nanmean(cost_array,axis=(0,1))
sem_int = np.nanstd(cost_array,axis=(0,1)) / np.sqrt(params.iterations)

mean_int_null = np.nanmean(cost_array_null,axis=(0,1))
sem_int_null = np.nanstd(cost_array_null,axis=(0,1)) / np.sqrt(params.iterations)

mean_int_linear = np.nanmean(cost_array_linear,axis=(0,1))
sem_int_linear = np.nanstd(cost_array_linear,axis=(0,1)) / np.sqrt(params.iterations)

ax.plot(mean_int,color='black',linewidth=1,linestyle='dashed')
ax.fill_between(range(len(mean_int)),mean_int-sem_int,mean_int+sem_int,color='royalblue',alpha=0.5)

ax.plot(mean_int_null,color='black',linewidth=1,linestyle='dotted')
ax.fill_between(range(len(mean_int_null)),mean_int_null-sem_int_null,mean_int_null+sem_int_null,color='gray',alpha=0.5)

ax.plot(mean_int_linear,color='black',linewidth=1,linestyle='dashdot')
ax.fill_between(range(len(mean_int_linear)),mean_int_linear-sem_int_linear,mean_int_linear+sem_int_linear,color='green',alpha=0.5)

ax.set_xlabel('Contest number')
ax.set_ylabel('Contest intensity') #\n(+/- SEM of iterations')

## Plot error
mean_err = np.mean(error_array,axis=(0,1))
sem_err = np.std(error_array,axis=(0,1)) / np.sqrt(params.iterations)

mean_err_null = np.mean(error_array_null,axis=(0,1))
sem_err_null = np.std(error_array_null,axis=(0,1)) / np.sqrt(params.iterations)

mean_err_linear = np.mean(error_array_linear,axis=(0,1))
sem_err_linear = np.std(error_array_linear,axis=(0,1)) / np.sqrt(params.iterations)

tick_size = int(params.fig_font * 3/4)

ax.tick_params(axis='both', which='major', labelsize=tick_size)


ax1.plot(mean_err,color='black',linewidth=1,linestyle='dashed',label='bayes')
ax1.fill_between(range(len(mean_err)),mean_err-sem_err,mean_err+sem_err,color='royalblue',alpha=0.5)

ax1.plot(mean_err_null,color='black',linewidth=1,label='no update',linestyle='dotted')
ax1.fill_between(range(len(mean_err_null)),mean_err_null-sem_err_null,mean_err_null+sem_err_null,color='gray',alpha=0.5)

ax1.plot(mean_err_linear,color='black',linewidth=1,label='linear',linestyle='dashdot')
ax1.fill_between(range(len(mean_err_linear)),mean_err_linear-sem_err_linear,mean_err_linear+sem_err_linear,color='green',alpha=0.5)

ax1.set_ylim([min(mean_err-sem_err) - 10,max(mean_err_linear+sem_err_linear) + 50])
ax1.set_xlabel('Contest number')
ax1.set_ylabel('Estimate error')# \n(+/- SEM of iterations')


ax1.tick_params(axis='both', which='major', labelsize=tick_size)

ax1.legend(loc='upper right',fontsize=tick_size)


fig.set_size_inches(6.5,3)
fig.tight_layout()
fig.savefig('./figures/fig3_IntenseAccuracy_linear.png',dpi=300)
fig.savefig('./figures/fig3_IntenseAccuracy_linear.svg')

if False:
    plt.show()
```
