## Supplemental Code for "Bayesian updating for self-assessment explains social dominance and winner-loser effects": figure4b_experience.py

```
#! /usr/bin/env python

## Code for fig 6a, showing how size of opponent impacts the strength of winner effect on different individuals


import numpy as np
from matplotlib import pyplot as plt

from bayesbots import Params
from bayesbots import Fish,FishNPC
from bayesbots import Fight

from tqdm import tqdm
import copy

from joblib import Parallel, delayed

plt.rcParams.update({'font.size': 10})

n_fight_list = np.arange(0,50,5)

# It turns out the exact extent of the decay for linear depends on the size distribution of the generated opponents, which I only do one time to save time.
## I could force the size distribution to be balanced, but it's easier to just roll the dice.
# And it works most of the time, but I wanted the figure to be consistent.
np.random.seed(3)

params = Params()
opp_params = Params()
#opp_params.baseline_effort = 0.5

#params.prior = True
params.size = 50
#params.poly_param_m = 0.2
## Make lots of naive fish of random sizes
npcs = [Fish(s,opp_params) for s in range(int(max(n_fight_list) * 4))]
npc_sizes = [npc.size for npc in npcs]
print('Median npc size:',np.mean(npc_sizes))

staged_opp = FishNPC(0,params) ## NPCs by default always invest 0.5 effort
staged_opp.size = 50
iterations = 1000
strategies = ['bayes','linear','no update']
## Make lots of focal fish with different strategies
outcome_array = np.empty([len(n_fight_list),len(strategies),iterations])
outcome_array.fill(np.nan)

def run_simulation(n_fights,print_me=False):
    outcome_array_n = np.empty([len(strategies),iterations])
    staged_opp_ = copy.deepcopy(staged_opp)
    for s_ in range(len(strategies)):
        #f_params = strat_params[s_]
        f_params = params.copy()
        f_params.update_method = strategies[s_]
        if print_me:
            print(n_fights,strategies[s_])
        for i in range(iterations):
            #npcs = copy.deepcopy(naive_npcs)
            opps = np.random.choice(npcs,n_fights)
            focal_fish = Fish(0,f_params)
## Provide background experiecne
            for o in opps:
                pre_fight = Fight(o,focal_fish,f_params)
                outcome = pre_fight.run_outcome()
                focal_fish.update(outcome,pre_fight)
                #print(i,len(focal_fish.est_record),o.effort,focal_fish.effort)
## Run a staged fight
            pre_est = focal_fish.estimate
            staged_opp_.size = focal_fish.size
            staged_opp_.params.size = focal_fish.size
            staged_fight = Fight(staged_opp_,focal_fish,outcome=1)
            staged_fight.run_outcome()
            focal_fish.update(True,staged_fight)
            #print(n_,s_,i,'pre,post:',pre_est,focal_fish.estimate)
## ASsay against naive size matched fish (with accuractely centered prior)
            assay_opp = Fish(1,f_params) 
            assay_opp.size = focal_fish.size
            assay_fight = Fight(assay_opp,focal_fish,f_params)
            outcome = assay_fight.run_outcome()
            outcome_array_n[s_,i] = outcome
    return outcome_array_n
   
print('running simulation in parallel...')
outcome_arrays = Parallel(n_jobs=10)(delayed(run_simulation)(n_,False) for n_ in n_fight_list)

outcome_array = np.array(outcome_arrays)
fig,ax = plt.subplots()

cors = ['tab:blue','tab:green','grey']
styles = ['dashed','dashdot','solid']

for s_ in range(len(strategies)):
    mean_outcome = np.mean(outcome_array[:,s_],axis=1)
    sem_outcome = np.std(outcome_array[:,s_],axis=1) / np.sqrt(iterations)
    ax.plot(n_fight_list,mean_outcome,label=strategies[s_],color='black',linestyle=styles[s_],linewidth=1)
    ax.fill_between(n_fight_list,mean_outcome - sem_outcome,mean_outcome+sem_outcome,alpha=0.5,color=cors[s_])

ax.set_ylim([0.45,1.0])
ax.axhline(0.5,color='black',linestyle=':')

ax.set_xlabel('Number of fights prior to assay')
ax.set_ylabel('Assay win rate')

tick_size = int(params.fig_font * 3/4)

plt.xticks(fontsize=tick_size)
plt.yticks(fontsize=tick_size)

ax.legend(fontsize=tick_size)

fig.set_size_inches(3.25,3)
fig.tight_layout()

fig.savefig('./figures/fig4b_experience.png',dpi=300)
fig.savefig('./figures/fig4b_experience.svg')
print('Done!')
#plt.show()
## Check against a fair fish
## Run them against a naive, size-matched fish to test for winner effect  
# (do I do win percent, or just contrast winner vs loser)
```
