## Supplemental Code for "Bayesian updating for self-assessment explains social dominance and winner-loser effects": figure5_priors.py

```
#! /usr/bin/env python

## Script to show that under bayes', the duration of the winner effect depends on the about of new info 

from bayesbots import Fish
from bayesbots import Fight
from bayesbots import Tank
from bayesbots import Params

import numpy as np
from scipy.stats import binom_test,norm
from scipy.stats import f_oneway,pearsonr

from matplotlib import pyplot as plt
from matplotlib import cm

from scipy.ndimage import gaussian_filter1d
from tqdm import tqdm

import random, copy
import itertools


#s,e,l = .6,.3,.1
UPDATE = True
params = Params()
plt.rcParams.update({'font.size': params.fig_font})
#if True:
    #params.effort_method = 'SmoothPoly'

#params.poly_param_a = 2
#params.poly_param_c = 0.1
#params.n_fights = 50 
#params.size = 50
#params.energy_cost = False
#params.acuity = 0.1
#params.awareness = 0.2
#params.start_energy = 1

params.iterations = 11
params.n_fish = 5
params.n_fights = 3
#params.f_method = 'balanced'
params.update_method = 'bayes'
params.f_outcome = 'math'
params.set_L()


## Have a fish win-lose, or lose-win, then measure the winner effect for that fish

## Let fish duke it out, then pull a fish out, let it win, and put it back in with the rest.
scale = 2

upset_count = 0
win_count1,win_count2 = 0,0
win_shifts,loss_shifts = [],[]
## Lets assume their estimate is correct.

params.age = 51
#params.size = 50 
#sizes = [10,20,30,40,50,60,70,80,90,100]
sizes = np.linspace(10,90,params.iterations)

## Set up the various fish conditions
#size_params = params.copy() ## Fish determine prior from their size
self_params = params.copy() ## fish determine prior from their size/age
#self_params.awareness = 0.2
self_params.set_params()

big_params = params.copy() ## Fish determine prior from surround fish (controlled to be big)
big_prior = norm.pdf(params.xs,60,5)
big_params.prior = big_prior / sum(big_prior)

small_params = params.copy() ## Fish determine prior from surrounding fish (controlled to be small)
small_prior = norm.pdf(params.xs,40,5)
small_params.prior = small_prior / sum(small_prior)

null_params = params.copy() ## Fish have null prior
guess_params = params.copy() ## Fish have random prior (but confident)

#guess_params.awareness = 20
#guess_params.awareness = 0.5

opp_params = params.copy()
opp_params.prior = True
opp_params.size = 50

## Uniform prior
null_params.prior = np.ones_like(null_params.xs) / len(null_params.xs)

self_fishes = [Fish(n+1,self_params) for n in range(params.iterations)]
null_fishes = [Fish(n+1,null_params) for n in range(params.iterations)]
big_fishes = [Fish(n+1,big_params) for n in range(params.iterations)]
small_fishes = [Fish(n+1,small_params) for n in range(params.iterations)]
guess_fishes = []
self_fishes,null_fishes,big_fishes,small_fishes = [],[],[],[]
## Make sloppy fish confident

for n in range(params.iterations):
    s = sizes[n]
    guess_params.size = s
    self_params.size = s
    null_params.size = s
    small_params.size = s
    big_params.size = s

    guess_prior = norm.pdf(params.xs,np.random.random() * 99 + 1,10)
    guess_params.prior = guess_prior / sum(guess_prior)
    f = Fish(n+1,guess_params)
    #prior = f.prior ** 10
    #f.prior = prior / np.sum(prior)
    f.get_stats()
    guess_fishes.append(f)

    self_fishes.append(Fish(n+1,self_params))
    big_fishes.append(Fish(n+1,big_params))
    small_fishes.append(Fish(n+1,small_params))

    null_fishes.append(Fish(n+1,null_params))
    null_fishes[n].get_stats()

for n in tqdm(range(params.iterations)):
    f = Fish(n+1,guess_params)
    #print(null_fishes[n].estimate,null_fishes[n].size)
opp_fish = Fish(0,opp_params)

#print(opp_fish.estimate)
#print(prior_fishes[0].estimate,null_fishes[0].estimate,guess_fishes[0].estimate)
p_efforts,n_efforts,g_efforts = [],[],[] # Prior(self), uNiform, random (Guess)
b_efforts,s_efforts = [],[] ## big adjacent, small adjactent

effort_array = np.empty([params.iterations,5,3])


for n in tqdm(range(params.iterations)):
    p,o = self_fishes[n],null_fishes[n] ## not sure why I use o vs n for null here
    g = guess_fishes[n]
    b,s = big_fishes[n],small_fishes[n]

    self_fight = Fight(opp_fish,self_fishes[n],params)
    null_fight = Fight(opp_fish,null_fishes[n],params)
    guess_fight = Fight(opp_fish,guess_fishes[n],params)
    big_fight = Fight(opp_fish,big_fishes[n],params)
    small_fight = Fight(opp_fish,small_fishes[n],params)

    for m in range(params.n_fights):
        p_out = self_fight.run_outcome()
        o_out = null_fight.run_outcome()
        g_out = guess_fight.run_outcome()
        b_out = big_fight.run_outcome()
        s_out = small_fight.run_outcome()

        if UPDATE:
            p.update(p_out,self_fight)
            s.update(s_out,small_fight)
            b.update(b_out,big_fight)

            if True:
                o.update(o_out,null_fight)
                g.update(g_out,guess_fight)

        effort_array[n,0,m] = p.effort
        effort_array[n,1,m] = o.effort
        effort_array[n,2,m] = g.effort
        effort_array[n,3,m] = b.effort
        effort_array[n,4,m] = s.effort
    p_efforts.append(p.effort)
    n_efforts.append(o.effort)
    g_efforts.append(g.effort)
    b_efforts.append(b.effort)
    s_efforts.append(s.effort)
    #print(o.size,o.effort)
p_mean,p_std = np.mean(p_efforts),np.std(p_efforts)
n_mean,n_std = np.mean(n_efforts),np.std(n_efforts)
g_mean,g_std = np.mean(g_efforts),np.std(g_efforts)
print(np.mean(p_efforts),np.mean(n_efforts),np.mean(g_efforts))
print(np.std(p_efforts),np.std(n_efforts),np.std(g_efforts))

print('self-informed mean:',np.mean(effort_array[:,0,:],axis=0))

#fig,(ax,ax2,ax3) = plt.subplots(3)

print('Uniform ONE WAY ANOVA:')
print(f_oneway(*effort_array[:,1]))


print('Random ONE WAY ANOVA:')
print(f_oneway(*effort_array[:,2]))

print('SELF ONE WAY ANOVA:')
print(f_oneway(*effort_array[:,0]))
print('Small ONE WAY ANOVA:')
print(f_oneway(*effort_array[:,4]))
print('bigger ONE WAY ANOVA:')
print(f_oneway(*effort_array[:,3]))

fish_groups = [self_fishes,null_fishes,guess_fishes,big_fishes,small_fishes]
strats = ['Self-Informed','Uniform','Random','Big-informed','Small-informed']

for strat in range(5):
    ys = np.mean(effort_array[:,strat],axis=1)
    fishes = fish_groups[strat]
    xs = [fishes[f].size for f in range(params.iterations)]
    print(strats[strat],pearsonr(xs,ys))

fig,ax = plt.subplots()

#ax.bar([1,2,3],[p_mean,n_mean,g_mean],yerr=[p_std,n_std,g_std],alpha=.1)
p_norms = np.mean(effort_array[:,0],axis=1)
n_norms = np.mean(effort_array[:,1],axis=1)
g_norms = np.mean(effort_array[:,2],axis=1)
b_norms = np.mean(effort_array[:,3],axis=1)
s_norms = np.mean(effort_array[:,4],axis=1)

offsets = [0.0,1.25,2.5,3.75,5]

p_sizes = [self_fishes[f].size/100+offsets[2]-0.5 for f in range(params.iterations)]
n_sizes = [null_fishes[f].size/100+offsets[0]-0.5 for f in range(params.iterations)]
g_sizes = [guess_fishes[f].size/100+offsets[1]-0.5 for f in range(params.iterations)]
b_sizes = [big_fishes[f].size/100+offsets[4]-0.5 for f in range(params.iterations)]
s_sizes = [small_fishes[f].size/100+offsets[3]-0.5 for f in range(params.iterations)]

all_sizes = [p_sizes,n_sizes,g_sizes,b_sizes,s_sizes]
cors = ['tab:blue','lightgray','black','darkblue','darkblue']
#for f in range(params.iterations):
for s in range(len(all_sizes)):
    xs = all_sizes[s]
    vals = effort_array[:,s].T
    cor = cors[s]
    #import pdb;pdb.set_trace()
    ax.boxplot(vals,positions=xs,widths=0.1,patch_artist=True,
        boxprops=dict(facecolor=cor,color='black',alpha=0.3),medianprops=dict(color='black'))

print(effort_array[:,0].shape)
print(pearsonr(np.mean(effort_array[:,0],axis=1),sizes))

#for i in [0]:
for i in [0,1,2,3,4]:
    fit_line = np.poly1d(np.polyfit(sizes,np.mean(effort_array[:,i],axis=1),1))
    if i == 0:
        color='red'
        ax.plot(all_sizes[i],fit_line(np.array(sizes)),color=color,linestyle=':')
    else:
        color = 'red'
        #ax.plot(all_sizes[i],fit_line(np.array(sizes)),color=color,linestyle=':')
#ax.boxplot([n_norms,g_norms,p_norms,s_norms,b_norms],positions=[0.5,1.5,2.5,3.25,3.72],widths=0.1)


#ax.scatter(p_sizes,p_norms,alpha=0.3,color=cors[2])
#ax.scatter(n_sizes,n_norms,alpha=0.3,color=cors[0])
#ax.scatter(g_sizes,g_norms,alpha=0.3,color=cors[1])
#ax.scatter(b_sizes,b_norms,alpha=0.3,color=cors[4])
#ax.scatter(s_sizes,s_norms,alpha=0.3,color=cors[3])

#ax.scatter([self_fishes[f].size/100+0.5 for f in range(params.iterations)],np.mean(effort_array[:,1],axis=1))

ax.set_xticks(offsets)
ax.set_xticklabels(['Uninformed\nprior','Misinformed\nprior','Self-informed\nprior','Small-informed\nprior','Big-informed\nprior']) #,rotation=45)
ax.axvline((offsets[1] + offsets[2]) / 2,linestyle=':',color='black')
#ax2.scatter([f.guess for f in prior_fishes],[f.effort for f in prior_fishes])
#ax3.scatter([f.estimate for f in prior_fishes],[f.effort for f in prior_fishes])
ax.set_ylabel('Aggressive behavior (effort)')

tick_size = int(params.fig_font * 3/4)

#plt.xticks(fontsize=tick_size)
plt.yticks(fontsize=tick_size)

fig.set_size_inches(6.5,3)
fig.tight_layout()
if UPDATE is False:
    fig.savefig('./figures/fig5_priors.png',dpi=300)
    fig.savefig('./figures/fig5_priors.svg')
else:
    fig.savefig('./figures/figB17_shiftingPriors.png',dpi=300)
    fig.savefig('./figures/figB17_shiftingPrior.svg')
#plt.show()
```
