## Supplemental Code for "Bayesian updating for self-assessment explains social dominance and winner-loser effects": README.md

# BayesBattleBots
Simulation of the winner effect using Bayes Theorem

This repo contains all the code that accompanies the paper: Bayesian updating for self-assessment explains social dominance and winner-loser effects.

The directory bayesbots contains the structure of the simulation: 

fish.py defines the individual agents
fight.py governs the interaction rules for a single contest
tank.py holds groups of agents and runs the contests beetween them
params.py contains all of the parameters for everything

Each script in the parent directory creates a figure (more/less), and are useful examples of different standard uses of the model.

I perform some figure formatting using Affinity Designer (combinging panels, adding letters, etc). 

See paper for full details: https://doi.org/10.1101/2023.05.10.540269
