## Supplemental Code for "Bayesian updating for self-assessment explains social dominance and winner-loser effects": SfigureA02.py

```
import matplotlib.pyplot as plt
from matplotlib import cm

import numpy as np
from bayesbots import Fight
from bayesbots import Fish

F = Fight(Fish(),Fish())

## set K at the top of the params, and store it there.
def bar(x):
    X = k * (m-x)
    return K * (1 / (1 + np.exp(X)) - S)

def foo(x):
    X = k * (m-x)
    return K_ / (1 + np.exp(X))

def tanthing(x):
    a = 10
    k = -1 / (2 * np.arctan(-a/(2*np.pi)))
    j = np.arctan(a*(x-m)/np.pi)
    y = k*j + m 
    return y

def ellie(x=0.7,a=2):
    X = x**a
    return X

## My original function
def art(x=0.7,a=1):
    X = k * (m-x**a)
    return K_ / (1 + np.exp(X)) / 2

## forget where this came from
def bob(x,B=2,r=2):
    y = (x**r)**B / ((x**r)**B + (1-x**r)**B)
    return y

## H function from staty on twitter
def hugh(x=0.7,p=0.35,d=6):
    if x == 1:
        H = 0.5
    H = 1/(1 + (p/(1-p))**d / (x/(1-x))**d)
    return H

p = 0.35
d=6
A = (p/(1-p))**-d
## G function from staty on twitter
def greg(x=0.7,p=0.35,d=6):
    return A / (A + (x/(1-x))**-d)

B = (p/(1-p))**d
## F function from staty on twitter
def frank(x=0.8,p=0.35,d=6):
    X = (x/(1-x))**d
    return X/ (X+B)

from scipy import stats

## CDF of difference between two distributions
def cory(x=0.7,s=50):
## Here, it's the CDF of being bigger, for some std
    x_abs = x*50 - 50
    #var = 1*x*50
    var = s
    y = 1 - stats.norm.cdf(0,x_abs,var)
    return y

from scipy.special import logit
## Logit-sigmoid
def perrin(x=0.7):
    x_ = logit(x)
    return 1 / (1 + np.exp(-x_))

import math
## Hyperbolic tangent tanh
def taco(x=0.7):
    a = 5
    h = 0.7
    #math.tanh(a*(x-h))/2 + 1/2    
    y = np.tanh(a*(x-h))/2 + 1/2
    return y

m = 0.5
k= 10
K_ = (1 + np.exp(k*(m-1)))
a = 2

S = 1 / (1 + np.exp(k*m))
K = 1 / (1/(1 + np.exp(-k*(1-m))) - S)
K_ = (1 + np.exp(k*(m-1)))

import timeit

print('sigmoidal',timeit.Timer(art).timeit(number=10000))
print('H cleverness',timeit.Timer(hugh).timeit(number=10000))
print('G more cleverness',timeit.Timer(greg).timeit(number=10000))
print('F more cleverness',timeit.Timer(frank).timeit(number=10000))
print('logit-sigmoid:',timeit.Timer(perrin).timeit(number=10000))
print('cdf',timeit.Timer(cory).timeit(number=10000))
print('tanh',timeit.Timer(taco).timeit(number=10000))
print('exp',timeit.Timer(ellie).timeit(number=10000))
#funk = F._wager_curve_sig
funk = cory
#print(foo(0),foo(m),foo(1))
print(funk(0.001),funk(m),funk(.99))
print(hugh(0.001),hugh(0.5),hugh(.99))
#xs = np.linspace(-3,3,100)
xs = np.linspace(0.01,1,100)
ys = funk(xs)

plt.rcParams.update({'font.size': F.params.fig_font})
tick_size = int(F.params.fig_font * 3/4)

fig,ax = plt.subplots()
a_list = [1,2,4,8,16]

if True:
    #for a in [0,0.25,0.5,1,2,4,np.inf]:
    for a_ in range(len(a_list)):
        a = a_list[a_]
        F.params.a = a
        ys = funk(xs,a)

        cmap = cm.get_cmap('viridis')
        a_cor = cmap((len(a_list) - a_) / len(a_list))

        ax.plot(xs,ys,color=a_cor,label='SD = ' + str(a),linewidth=1)
    ax.set_xlabel('Relative wager')
    ax.set_ylabel('Probability of upset')
    ax.legend(loc='upper left',fontsize=tick_size)

#plt.legend()
#plt.plot(xs,ys)
#plt.axvline(m)

ax.tick_params(axis='both', which='major', labelsize=tick_size)

fig.set_size_inches(2.1,2.1)
fig.tight_layout()

fig2,ax2 = plt.subplots()

xs = np.linspace(0,100.1,101)
d1 = stats.norm.pdf(xs,50,8)
d2 = stats.norm.pdf(xs,45,8)

d3 = stats.norm.pdf(xs,50,4)
d4 = stats.norm.pdf(xs,45,4)

ax2.plot(xs,d1,color='gold',label='Larger')
ax2.plot(xs,d2,color='royalblue',label='Smaller')
ax2.set_ylabel('Probability density')
ax2.set_xlabel('Wager')
ax2.tick_params(axis='both', which='major', labelsize=tick_size)
#ax2.legend(loc='upper left')
#ax2.plot(xs,d3,color='tab:blue',label='std=4')
#ax2.plot(xs,d4,color='tab:blue')

fig2.set_size_inches(2.1,2.1)
fig2.tight_layout()

rel_xs = np.linspace(0,1,100)
fig3,ax3 = plt.subplots()

l_list = np.linspace(-1,0,11)
L_sublist = ['1.0','4.2','6.3','12.7','inf']
shifted_l = (l_list + 1)/2
l_list = np.round(np.tan(np.array(np.pi/2 - shifted_l*np.pi/2)),1)

for l_ in range(len(l_list)):
    l = l_list[::-1][l_]
    if l > 10000:
        l_label = 'inf'
    else:
        l_label = str(l)
    cmap = cm.get_cmap('viridis')
    l_cor = cmap(l_ / len(l_list))
    if l_label in L_sublist:
        ax3.plot(rel_xs,ellie(rel_xs,l),label="l = " + l_label,color=l_cor)

ax3.set_xlabel('Relative wager')
ax3.set_ylabel('Probability of upset')

ax3.tick_params(axis='both', which='major', labelsize=tick_size)
ax3.legend(fontsize=tick_size)
fig3.set_size_inches(2.1,2.1)
fig3.tight_layout()

fig.savefig('./figures/figA2a_cdf.png',dpi=300)
fig.savefig('./figures/figA2a_cdf.svg')

fig2.savefig('./figures/figA2b_dist.png',dpi=300)
fig2.savefig('./figures/figA2b_dist.svg')

fig3.savefig('./figures/figA2c_exp.png',dpi=300)
fig3.savefig('./figures/figA2c_exp.svg')

#plt.show()
```
