## Supplemental Code for "Bayesian updating for self-assessment explains social dominance and winner-loser effects": SfigureB01.py

```
import numpy as np
from matplotlib import pyplot as plt
import matplotlib

from bayesbots import Fish,FishNPC
from bayesbots import Fight
from bayesbots import Params

from tqdm import tqdm

## A couple of helper functions to keep things tidy

def mean_sem(a):
    mean_a = np.nanmean(a)
    sem_a = np.nanstd(a) / np.sqrt(len(a))
    return mean_a,sem_a

def plot_fill(xs,a,fig=None,ax=None,color='grey',alpha=0.5):
    if ax is None:
        fig,ax = plt.subplots()
    ax.plot(xs,a[:,0],color='black')
    ax.fill_between(xs,a[:,0] - a[:,1],a[:,0] + a[:,1],color=color,alpha=alpha)
    return fig,ax

def run_sim(params):
    iterations = params.iterations

    outcome_array = np.empty([iterations,2])
    outcome_array.fill(np.nan)

    win_info_array = np.array(outcome_array)
    loss_info_array = np.array(outcome_array)

    for i in tqdm(range(iterations)):
        focal_winner = Fish(i+2,params)
        focal_loser = focal_winner.copy() 

        f_sizes.append(focal_winner.size)
## Stage a bunch of wins and losses against size-matched fish

        staged_opp = Fish(0,params)
        
        staged_win = Fight(staged_opp,focal_winner,params,outcome=1)
        staged_win.run_outcome()
        focal_winner.update(True,staged_win)

        staged_loss = Fight(staged_opp,focal_loser,params,outcome=0)
        staged_loss.run_outcome()
        focal_loser.update(False,staged_loss)

## Assay against size matched fish
        assay_fish = Fish(1,assay_params)

        assay_winner = Fight(assay_fish,focal_winner,params)
        winner_output = assay_winner.run_outcome()
        outcome_array[i,1] = winner_output

        assay_loser = Fight(assay_fish,focal_loser,params)
        loser_output = assay_loser.run_outcome()
        outcome_array[i,0] = loser_output

        win_info_array[i] = focal_winner.effort,focal_winner.estimate 
        loss_info_array[i] = focal_loser.effort,focal_loser.estimate 

        #if assay_fish.wager > focal_winner.wager:
        #    w_probs.append(assay_winner.p_win)
        #else:
        #    w_probs.append(1-assay_winner.p_win)

        #if assay_fish.wager > focal_loser.wager:
        #    l_probs.append(assay_loser.p_win)
        #else:
        #    l_probs.append(1-assay_loser.p_win)


    return outcome_array,win_info_array,loss_info_array


iterations = 1000
params = Params()

plt.rcParams.update({'font.size': params.fig_font})
plt.rcParams.update({'lines.linewidth': 1})

tick_size = (params.fig_font * 2/3)

matplotlib.rc('xtick', labelsize=tick_size) 
matplotlib.rc('ytick', labelsize=tick_size) 

params.iterations = iterations

params.size = 50

assay_params = params.copy()

print(params.awareness)
print(params.outcome_params)

#assay_params.baseline_effort = 0.535
#assay_params.prior = True

outcome_array = np.empty([iterations,2])
outcome_array.fill(np.nan)

win_info_array = np.array(outcome_array)
loss_info_array = np.array(outcome_array)
w_probs,l_probs = [],[]

f_sizes = []

s_res = 10+1
s_params = np.linspace(0,1.00,s_res)

win_estimates = np.empty([4,s_res,2])
win_efforts = np.empty_like(win_estimates)
win_outputs = np.empty_like(win_estimates)

loss_estimates = np.empty_like(win_estimates)
loss_efforts = np.empty_like(win_estimates)
loss_outputs = np.empty_like(win_estimates)

def set_custom_params(params,name,value):
    if name == 's':
        params.outcome_params[0] = value
    if name == 'l':
        params.outcome_params[2] = value
    if name == 'Sa':
        params.awareness = value
    if name == 'Sc':
        params.acuity = value
    params.set_params()
    return params

param_list = ['s','l','Sa','Sc']

l_res = s_res
a_res = s_res

param_set = {'s':np.linspace(0,1,s_res),
             'l':np.linspace(-1,0,l_res),
             'Sa':np.linspace(0,1,a_res),
             'Sc':np.linspace(0,1,a_res)
}


## Build labels for l and a/c sets
np.set_printoptions(formatter={'all':lambda x: str(x)})
shifted_l = (param_set['l'] + 1)/2
l_labels = np.round(np.tan(np.array(np.pi/2 - shifted_l*np.pi/2)),1).astype('str')
l_labels[0] = 'inf' 

a_labels = np.round(np.tan(np.array(param_set['Sa'])*np.pi/2) * 20,1).astype('str')
a_labels[-1] = 'inf'

default_params = [params.outcome_params[0],params.outcome_params[2],params.awareness,params.acuity]

for p_ in range(len(param_list)):
    params = Params()
    params.iterations = iterations
    params.size = 50
    assay_params = params.copy()

    p = param_list[p_]
    param_space = param_set[p]
    for i_ in range(len(s_params)):
        i = param_space[i_]
        print(p,i)
        params = set_custom_params(params,p,i)
        print(params.outcome_params[0],params.L,params.A,params.C)
        outcome_array,win_info_array,loss_info_array = run_sim(params)
## get win stats, using little helper function
        win_outputs[p_,i_] = mean_sem(outcome_array[:,1])
        win_estimates[p_,i_] = mean_sem(win_info_array[:,1])
        win_efforts[p_,i_] = mean_sem(win_info_array[:,0])

        loss_outputs[p_,i_] = mean_sem(outcome_array[:,0])
        loss_estimates[p_,i_] = mean_sem(loss_info_array[:,1])
        loss_efforts[p_,i_] = mean_sem(loss_info_array[:,0])

fig,axes = plt.subplots(3,4,sharex='col',sharey='row')

for p_ in range(len(param_list)):
    p = param_list[p_]
    xs_params = param_set[p]
    print(p,xs_params)
    plot_fill(xs_params,win_estimates[p_],ax=axes[0,p_],color='gold')
    plot_fill(xs_params,loss_estimates[p_],ax=axes[0,p_],color='darkblue')

    plot_fill(xs_params,win_efforts[p_],ax=axes[1,p_],color='gold')
    plot_fill(xs_params,loss_efforts[p_],ax=axes[1,p_],color='darkblue')

    plot_fill(xs_params,win_outputs[p_],ax=axes[2,p_],color='gold')
    plot_fill(xs_params,loss_outputs[p_],ax=axes[2,p_],color='darkblue')

axes[2,0].set_xlabel('s value')
axes[2,0].set_xticks(param_set['s'][1::2])
axes[2,0].set_xticklabels(np.round(param_set['s'],1)[1::2],rotation='45')

axes[2,1].set_xlabel('l value')
axes[2,1].set_xticks(param_set['l'][::2])
axes[2,1].set_xticklabels(l_labels[::2],rotation='45')
axes[2,1].invert_xaxis()

axes[2,2].set_xlabel('Sigma_a value')
axes[2,2].set_xticks(param_set['Sa'][1::2])
axes[2,2].set_xticklabels(a_labels[1::2],rotation='45')

axes[2,3].set_xlabel('Sigma_c value')
axes[2,3].set_xticks(param_set['Sa'][1::2])
axes[2,3].set_xticklabels(a_labels[1::2],rotation='45')

axes[0,0].set_ylabel('Estimate')
axes[1,0].set_ylabel('Assay effort')
axes[2,0].set_ylabel('Assay win rate')

axes[0,0].set_ylim([20,80])
axes[0,0].set_yticks([25,50,75])


axes[1,0].set_ylim([-0.1,1.1])
axes[1,0].set_yticks([0,0.5,1.0])

axes[2,0].set_ylim([-0.1,1.1])
axes[2,0].set_yticks([0,0.5,1.0])


fig.set_size_inches(6.5,4)
fig.tight_layout()

for c_ in range(4):
    for r_ in range(3):
        ax = axes[r_,c_]
        ax.axvline(default_params[c_],color='red',linestyle=':')

fig.savefig('./figures/figB01_WinLoss.png',dpi=300)
fig.savefig('./figures/figB01_WinLoss.svg')

#plt.show()
```
