## Supplemental Code for "Bayesian updating for self-assessment explains social dominance and winner-loser effects": SfigureB02.py

```
import numpy as np
from matplotlib import pyplot as plt
import matplotlib

from bayesbots import Fish,FishNPC
from bayesbots import Fight
from bayesbots import Params

from tqdm import tqdm

## A couple of helper functions to keep things tidy

def mean_sem(a):
    mean_a = np.nanmean(a)
    sem_a = np.nanstd(a) / np.sqrt(len(a))
    return mean_a,sem_a

## Stupid little function to extract blocks of true values from an array
def get_chunks(x_bool):
    start = None 
    i_list = []
    all_is = []
    for i_ in range(len(x_bool)):
        if x_bool[i_] == False:
            if start is not None:
                all_is.append(i_list)
                i_list = []
                start = None
        else:
            if start is None:
                start = i_
            i_list.append(i_)
    if start is not None:
        all_is.append(i_list)
    return all_is

def plot_fill(xs,a,fig=None,ax=None,color='grey',alpha=0.5):
    if ax is None:
        fig,ax = plt.subplots()
    ax.plot(xs,a[:,0],color='black')
    top_line = a[:,0] + a[:,1]
    bottom_line = a[:,0] - a[:,1]
    if 'STAT' in color:
        #import pdb;pdb.set_trace()
        gold_points = bottom_line > 0
        blue_points = top_line < 0

        new_xs = np.linspace(min(xs),max(xs),len(xs) * 20)
        new_bottom = np.interp(new_xs,xs,bottom_line)
        new_top = np.interp(new_xs,xs,top_line)

        xs = new_xs
        if color == 'STAT':

            gray_bool = (new_bottom <= 0) & (new_top >= 0)
            gold_bits = get_chunks(new_bottom > 0)
            blue_bits = get_chunks(new_top < 0)
            gray_bits = get_chunks(gray_bool)
            for c in gray_bits:
                ax.fill_between(xs[c],new_bottom[c],new_top[c],color='gray',alpha=alpha)
            for c in gold_bits:
                ax.fill_between(xs[c],new_bottom[c],new_top[c],color='gold',alpha=alpha)
            for c in blue_bits:
                ax.fill_between(xs[c],new_bottom[c],new_top[c],color='darkblue',alpha=alpha)
        elif color == 'STAT_SIMPLE':
            gold_bits = get_chunks(new_top > 0)
            blue_bits = get_chunks(new_bottom < 0)
            for c in gold_bits:
                ax.fill_between(xs[c],np.clip(new_bottom[c],0,a_max=None),new_top[c],color='gold',alpha=alpha)
            for c in blue_bits:
                ax.fill_between(xs[c],new_bottom[c],np.clip(new_top[c],None,0),color='darkblue',alpha=alpha)
    else:
        ax.fill_between(xs,a[:,0] - a[:,1],a[:,0] + a[:,1],facecolor=color,alpha=alpha)
    y_max = max([np.max(np.abs(a[:,0] - a[:,1])),np.max(np.abs(a[:,0] + a[:,1]))])
    ax.set_ylim([y_max*-1.1,y_max*1.1])
   
    return fig,ax

def run_sim(params):
    iterations = params.iterations

    outcome_array = np.empty([iterations,2])
    outcome_array.fill(np.nan)

    win_info_array = np.array(outcome_array)
    loss_info_array = np.array(outcome_array)

    for i in tqdm(range(iterations)):
        focal_winner = Fish(i+2,params)
        focal_loser = focal_winner.copy() 

        f_sizes.append(focal_winner.size)
## Stage a bunch of wins and losses against size-matched fish

        staged_opp = Fish(0,params)
        
        staged_win = Fight(staged_opp,focal_winner,params,outcome=1)
        staged_win.run_outcome()
        focal_winner.update(True,staged_win)

        staged_loss = Fight(staged_opp,focal_loser,params,outcome=0)
        staged_loss.run_outcome()
        focal_loser.update(False,staged_loss)

## Assay against size matched fish
        assay_fish = Fish(1,assay_params)

        assay_winner = Fight(assay_fish,focal_winner,params)
        winner_output = assay_winner.run_outcome()
        outcome_array[i,1] = winner_output

        assay_loser = Fight(assay_fish,focal_loser,params)
        loser_output = assay_loser.run_outcome()
        outcome_array[i,0] = loser_output

        win_info_array[i] = focal_winner.effort,focal_winner.estimate 
        loss_info_array[i] = focal_loser.effort,focal_loser.estimate 

        #if assay_fish.wager > focal_winner.wager:
        #    w_probs.append(assay_winner.p_win)
        #else:
        #    w_probs.append(1-assay_winner.p_win)

        #if assay_fish.wager > focal_loser.wager:
        #    l_probs.append(assay_loser.p_win)
        #else:
        #    l_probs.append(1-assay_loser.p_win)


    return outcome_array,win_info_array,loss_info_array


iterations = 1000 
params = Params()
plt.rcParams.update({'font.size': params.fig_font})
plt.rcParams.update({'lines.linewidth': 1})

tick_size = (params.fig_font * 2/3)

matplotlib.rc('xtick', labelsize=tick_size) 
matplotlib.rc('ytick', labelsize=tick_size) 

params.iterations = iterations

params.size = 50

assay_params = params.copy()

print(params.awareness)
print(params.outcome_params)

#assay_params.baseline_effort = 0.535
#assay_params.prior = True

outcome_array = np.empty([iterations,2])
outcome_array.fill(np.nan)

win_info_array = np.array(outcome_array)
loss_info_array = np.array(outcome_array)
w_probs,l_probs = [],[]

f_sizes = []

s_res = 10+1
s_params = np.linspace(0,1.00,s_res)

win_estimates = np.empty([4,s_res,2])
win_efforts = np.empty_like(win_estimates)
win_outputs = np.empty_like(win_estimates)

loss_estimates = np.empty_like(win_estimates)
loss_efforts = np.empty_like(win_estimates)
loss_outputs = np.empty_like(win_estimates)

def set_custom_params(params,name,value):
    if name == 's':
        params.outcome_params[0] = value
    if name == 'l':
        params.outcome_params[2] = value
    if name == 'Sa':
        params.awareness = value
    if name == 'Sc':
        params.acuity = value
    params.set_params()
    return params

param_list = ['s','l','Sa','Sc']

l_res = s_res
a_res = s_res

param_set = {'s':np.linspace(0,1,s_res),
             'l':np.linspace(-1,0,l_res),
             'Sa':np.linspace(0,1,a_res),
             'Sc':np.linspace(0,1,a_res)
}

print(param_set['l'])
print(param_set['Sa'])

np.set_printoptions(formatter={'all':lambda x: str(x)})
shifted_l = (param_set['l'] + 1)/2
l_labels = np.round(np.tan(np.array(np.pi/2 - shifted_l*np.pi/2)),1).astype('str')
l_labels[0] = 'inf' 

a_labels = np.round(np.tan(np.array(param_set['Sa'])*np.pi/2) * 20,1).astype('str')
a_labels[-1] = 'inf'
print(l_labels,a_labels)

default_params = [params.outcome_params[0],params.outcome_params[2],params.awareness,params.acuity]

for p_ in range(len(param_list)):
    params = Params()
    params.iterations = iterations
    params.size = 50
    assay_params = params.copy()

    p = param_list[p_]
    param_space = param_set[p]
    for i_ in range(len(s_params)):
        i = param_space[i_]
        print(p,i)
        params = set_custom_params(params,p,i)
        print(params.outcome_params[0],params.L,params.A,params.C)
        outcome_array,win_info_array,loss_info_array = run_sim(params)
## get win stats, using little helper function
        win_outputs[p_,i_] = mean_sem(outcome_array[:,1])
        win_estimates[p_,i_] = mean_sem(win_info_array[:,1])
        win_efforts[p_,i_] = mean_sem(win_info_array[:,0])

        loss_outputs[p_,i_] = mean_sem(outcome_array[:,0])
        loss_estimates[p_,i_] = mean_sem(loss_info_array[:,1])
        loss_efforts[p_,i_] = mean_sem(loss_info_array[:,0])

fig,axes = plt.subplots(3,4,sharex='col',sharey='row')

for p_ in range(len(param_list)):
    p = param_list[p_]
    xs_params = param_set[p]
    est_array = np.empty_like(win_estimates[p_])
    effort_array =np.empty_like(est_array)
    output_array = np.empty_like (est_array)

    est_array[:,0] = (win_estimates[p_,:,0] - 50) - (50-loss_estimates[p_,:,0])
    effort_array[:,0] = (win_efforts[p_,:,0] - 0.5) - (0.5 - loss_efforts[p_,:,0])
    output_array[:,0] = (win_outputs[p_,:,0] - 0.5) - (0.5 - loss_outputs[p_,:,0])

    est_array[:,1] = win_estimates[p_,:,1] + loss_estimates[p_,:,1]
    effort_array[:,1] = win_efforts[p_,:,1] + loss_efforts[p_,:,1]
    output_array[:,1] = win_outputs[p_,:,1] + loss_outputs[p_,:,1]

    #plot_fill(xs_params,est_array,ax=axes[0,p_],color='STAT')
    plot_fill(xs_params,est_array,ax=axes[0,p_],color='STAT_SIMPLE')
    plot_fill(xs_params,effort_array,ax=axes[1,p_],color='STAT_SIMPLE')
    plot_fill(xs_params,output_array,ax=axes[2,p_],color='STAT_SIMPLE')
    #plot_fill(xs_params,output_array,ax=axes[2,p_],color='gray')

    '''
    plot_fill(xs_params[est_array > 0],est_array[est_array > 0],ax=axes[0,p_],color='gold')
    plot_fill(xs_params[est_array <= 0],est_array[est_array <= 0],ax=axes[0,p_],color='darkblue')

    plot_fill(xs_params[effort_array > 0],effort_array[effort_array > 0],ax=axes[1,p_],color='gold')
    plot_fill(xs_params[effort_array <= 0],effort_array[effort_array <= 0],ax=axes[1,p_],color='darkblue')

    plot_fill(xs_params[output_array > 0],output_array[output_array > 0],ax=axes[2,p_],color='gold')
    plot_fill(xs_params[output_array <= 0],output_array[output_array <= 0],ax=axes[2,p_],color='blue')
    '''
axes[2,0].set_xlabel('s value')
axes[2,0].set_xticks(param_set['s'][1::2])
axes[2,0].set_xticklabels(np.round(param_set['s'],1)[1::2],rotation='45')

axes[2,1].set_xlabel('l value')
axes[2,1].set_xticks(param_set['l'][::2])
axes[2,1].set_xticklabels(l_labels[::2],rotation='45')
axes[2,1].invert_xaxis()

axes[2,2].set_xlabel('Sigma_a value')
axes[2,2].set_xticks(param_set['Sa'][1::2])
axes[2,2].set_xticklabels(a_labels[1::2],rotation='45')

axes[2,3].set_xlabel('Sigma_c value')
axes[2,3].set_xticks(param_set['Sa'][1::2])
axes[2,3].set_xticklabels(a_labels[1::2],rotation='45')

axes[0,0].set_ylabel('Estimate')
axes[1,0].set_ylabel('Assay effort')
axes[2,0].set_ylabel('Assay win rate')

axes[0,0].set_ylim([-2.1,2.1])
axes[0,0].set_yticks([-2.00,-1.00,0.00,1.00,2.00])
#axes[0,0].set_yticklabels(['-2.00','-1.00','0.00','1.00','2.00'])
#axes[0,0].set_yticklabels(np.round(np.arange(-2,2.1),4))

axes[1,0].set_ylim([-0.061,0.061])
axes[1,0].set_yticks([-0.06,-0.03,0,0.03,0.06])

axes[2,0].set_ylim([-0.11,.11])
axes[2,0].set_yticks([-0.10,-0.05,0,0.05,0.10])


fig.set_size_inches(6.5,4)
fig.tight_layout()

for c_ in range(4):
    for r_ in range(3):
        ax = axes[r_,c_]
        y_max = 0
        ax.axvline(default_params[c_],color='red',linestyle=':')
        ax.axhline(0,color='black',alpha=0.2,linestyle=':')


fig.savefig('./figures/figB02_WinvLoss.png',dpi=300)
fig.savefig('./figures/figB02_WinvLoss.svg')
#plt.show()
```
