## Supplemental Code for "Bayesian updating for self-assessment explains social dominance and winner-loser effects": SfigureB03.py

```
#! /usr/bin/env python

## Code for fig 3b, showing the loser effect is stronger

import numpy as np
from matplotlib import pyplot as plt
import matplotlib

from bayesbots import Fish,FishNPC
from bayesbots import Fight
from bayesbots import Params

from tqdm import tqdm

iterations = 1000
params = Params()

plt.rcParams.update({'font.size': params.fig_font})
plt.rcParams.update({'lines.linewidth': 1})

tick_size = int(params.fig_font * 3/4)
matplotlib.rc('xtick', labelsize=tick_size) 
matplotlib.rc('ytick', labelsize=tick_size) 
params.effort_method='ScaledPoly'

e_res = 11
e_list = np.linspace(0,1,e_res)

params.size = 50

results_array = np.empty([e_res,2,2])

for e_ in range(e_res):
    e = e_list[e_]

    params.outcome_params = [0.7,e,-0.8]
    params.set_params()

    assay_params = params.copy()
    assay_params.baseline_effort = 0.535
    assay_params.prior = True

    outcome_array = np.empty([iterations,2])
    outcome_array.fill(np.nan)

    win_info_array = np.array(outcome_array)
    loss_info_array = np.array(outcome_array)
#iterations = 1000
    w_probs,l_probs = [],[]

    f_sizes = []

    for i in tqdm(range(iterations)):
        focal_winner = Fish(i+2,params)
        f_sizes.append(focal_winner.size)
        focal_loser = focal_winner.copy() 
## Stage a bunch of wins and losses against size-matched fish

        #assay_params.size = focal_winner.size
        staged_opp = FishNPC(0,assay_params)
        staged_opp.size = focal_winner.size
        
        staged_win = Fight(staged_opp,focal_winner,params,outcome=1)
        staged_win.run_outcome()
        focal_winner.update(True,staged_win)

        staged_loss = Fight(staged_opp,focal_loser,params,outcome=0)
        staged_loss.run_outcome()
        focal_loser.update(False,staged_loss)
        #print(focal_winner.effort,focal_loser.effort)
        #print(focal_winner.estimate,focal_loser.estimate)
        #import pdb;pdb.set_trace()
        #if focal_loser.estimate > 50:
        #    import pdb;pdb.set_trace()
        #print('## staged opp size:',staged_opp.size)

## Assay against size matched fish
        assay_params.size = focal_winner.size
        assay_fish = Fish(1,assay_params)

        assay_winner = Fight(assay_fish,focal_winner,params)
        winner_output = assay_winner.run_outcome()
        outcome_array[i,1] = winner_output
        
        if assay_fish.wager > focal_winner.wager:
            w_probs.append(assay_winner.p_win)
        else:
            w_probs.append(1-assay_winner.p_win)
        #print('## Assay fish size:',assay_fish.size,assay_fish.estimate)
        #print('win:',assay_fish.effort,focal_winner.estimate,focal_winner.effort,assay_winner.p_win,winner_output) 
        assay_loser = Fight(assay_fish,focal_loser,params)
        loser_output = assay_loser.run_outcome()

        outcome_array[i,0] = loser_output
        win_info_array[i] = focal_winner.effort,focal_winner.estimate 
        loss_info_array[i] = focal_loser.effort,focal_loser.estimate 
        #print('loss:',assay_fish.effort,focal_loser.estimate,focal_loser.effort,assay_loser.p_win,loser_output)
        if assay_fish.wager > focal_loser.wager:
            l_probs.append(assay_loser.p_win)
        else:
            l_probs.append(1-assay_loser.p_win)
        #break
    results_array[e_,:,0] = np.mean(outcome_array,axis=0)
    results_array[e_,:,1] = np.std(outcome_array,axis=0) / np.sqrt(iterations)

fig,(ax0,ax) = plt.subplots(1,2)

low_params = params.copy()
low_params.outcome_params[1] = 0.3
low_params.set_params()

high_params = params.copy()
high_params.outcome_params[1] = 0.7
high_params.set_params()

xs = np.linspace(0,1,50)
low_fish = Fish(0,low_params)
high_fish = Fish(1,high_params)

low_e = low_fish._scale_func(xs)
high_e = high_fish._scale_func(xs)

ax0.plot(xs,xs,color='black',linestyle=':')
ax0.plot(xs,high_e,color='red',label='h=0.7')
ax0.plot(xs,low_e,color='orange',linestyle='dashed',label='h=0.3')
ax0.set_xlabel('Estimated probabilty of winning')
ax0.set_ylabel('Effort')
ax0.legend(fontsize=tick_size)

winner_mean = results_array[:,1,0]
winner_sem = results_array[:,1,1]

loser_mean = results_array[:,0,0]
loser_sem = results_array[:,0,1]

ax.plot(e_list,winner_mean,color='black',label='Post-win')
ax.fill_between(e_list,winner_mean - winner_sem,winner_mean + winner_sem,color='gold',alpha=0.5)

ax.plot(e_list,loser_mean,color='black',linestyle='dashed',label='Post-loss')
ax.fill_between(e_list,loser_mean - loser_sem,loser_mean + loser_sem,color='darkblue',alpha=0.5)

ax.axhline(0.5,color='gray',linestyle=':')
ax.legend(fontsize=tick_size)
ax.set_xlabel('Value of h')
ax.set_ylabel('Assay win rate')

fig.set_size_inches(6.5,3)
fig.tight_layout()

fig.savefig('./figures/figB03_lossBias.png',dpi=300)
fig.savefig('./figures/figB03_lossBias.svg')

#plt.show()
```
