## Supplemental Code for "Bayesian updating for self-assessment explains social dominance and winner-loser effects": SfigureB04.py

```
import numpy as np
from matplotlib import pyplot as plt
import matplotlib

from bayesbots import Fish,FishNPC
from bayesbots import Fight
from bayesbots import Params

from tqdm import tqdm
from joblib import Parallel, delayed

## A couple of helper functions to keep things tidy

def mean_sem(a):
    mean_a = np.nanmean(a)
    sem_a = np.nanstd(a) / np.sqrt(len(a))
    return mean_a,sem_a

def plot_fill(xs,a,fig=None,ax=None,color='grey',alpha=0.5):
    if ax is None:
        fig,ax = plt.subplots()
    ax.plot(xs,a[:,0],color='black')
    ax.fill_between(xs,a[:,0] - a[:,1],a[:,0] + a[:,1],color=color,alpha=alpha)
    return fig,ax

def run_sim(params,assay_params):
    iterations = params.iterations

    outcome_array = np.empty([iterations,2])
    outcome_array.fill(np.nan)

    WL_info_array = np.array(outcome_array)
    LW_info_array = np.array(outcome_array)

    for i in range(iterations):
## Fish WL, LW
        focal_WL = Fish(i+2,params)
        focal_LW = focal_WL.copy() 

        #f_sizes.append(focal_winner.size)
## Stage a bunch of wins and losses against size-matched fish

        staged_opp = Fish(0,params)
        
        WL_w = Fight(staged_opp,focal_WL,params,outcome=1)
        WL_w.run_outcome()
        focal_WL.update(True,WL_w)
        
        WL_l = Fight(staged_opp,focal_WL,params,outcome=0) 
        WL_l.run_outcome()
        focal_WL.update(False,WL_l)

        LW_l = Fight(staged_opp,focal_LW,params,outcome=0) 
        LW_l.run_outcome()
        focal_LW.update(False,LW_l)

        LW_w = Fight(staged_opp,focal_LW,params,outcome=1) 
        LW_w.run_outcome()
        focal_LW.update(True,LW_w)

## Assay against size matched fish
        assay_fish = Fish(1,assay_params)

        assay_WL = Fight(assay_fish,focal_WL,params)
        WL_output = assay_WL.run_outcome()
        outcome_array[i,1] = WL_output

        assay_LW = Fight(assay_fish,focal_LW,params)
        LW_output = assay_LW.run_outcome()
        outcome_array[i,0] = LW_output

        WL_info_array[i] = focal_WL.effort,focal_WL.estimate 
        LW_info_array[i] = focal_LW.effort,focal_LW.estimate 
    return outcome_array,WL_info_array,LW_info_array


iterations = 1000
params = Params()
plt.rcParams.update({'font.size': params.fig_font})
plt.rcParams.update({'font.size': params.fig_font})
plt.rcParams.update({'lines.linewidth': 1})

tick_size = (params.fig_font * 2/3)

matplotlib.rc('xtick', labelsize=tick_size) 
matplotlib.rc('ytick', labelsize=tick_size) 

params.iterations = iterations

params.size = 50

assay_params = params.copy()

s_res = 10+1
a_res = s_res

s_params = np.linspace(0,1.00,s_res)

wl_estimates = np.empty([s_res,s_res,a_res,a_res,2])
wl_efforts = np.empty_like(wl_estimates)
wl_outputs = np.empty_like(wl_estimates)

lw_estimates = np.empty_like(wl_estimates)
lw_efforts = np.empty_like(wl_estimates)
lw_outputs = np.empty_like(wl_estimates)

l_res = s_res
a_res = s_res

s_set = np.linspace(0,1,s_res)
l_set = np.linspace(-1,0,l_res)
a_set = np.linspace(0,1,a_res)
c_set = np.linspace(0,1,a_res)


np.set_printoptions(formatter={'all':lambda x: str(x)})
shifted_l = (l_set + 1)/2
l_labels = np.round(np.tan(np.array(np.pi/2 - shifted_l*np.pi/2)),1).astype('str')
l_labels[0] = 'inf' 

a_labels = np.round(np.tan(np.array(a_set)*np.pi/2) * 20,1).astype('str')
a_labels[-1] = 'inf'


default_params = [params.outcome_params[0],params.outcome_params[2],params.awareness,params.acuity]

def run_many_sims(s):
    wl_estimates_s = np.empty([s_res,a_res,a_res,2])
    wl_efforts_s = np.empty_like(wl_estimates_s)
    wl_outputs_s = np.empty_like(wl_estimates_s)

    lw_estimates_s = np.empty_like(wl_estimates_s)
    lw_efforts_s = np.empty_like(wl_estimates_s)
    lw_outputs_s = np.empty_like(wl_estimates_s)


    for l_ in tqdm(range(l_res)):
        l = l_set[l_]
        for a_ in range(a_res):
            awareness = a_set[a_]
            for c_ in range(a_res):
                acuity = c_set[c_]
                params = Params()
                params.acuity = acuity
                params.awareness = awareness
                params.outcome_params = [s,0.5,l]
                params.set_params()

                params.iterations = iterations
                params.size = 50
                assay_params = params.copy()

## I may need to run this in parallel
                outcome_array,wl_info_array,lw_info_array = run_sim(params,assay_params)
## get win stats, using little helper function
                wl_outputs_s[l_,a_,c_] = mean_sem(outcome_array[:,1])
                wl_estimates_s[l_,a_,c_] = mean_sem(wl_info_array[:,1])
                wl_efforts_s[l_,a_,c_] = mean_sem(wl_info_array[:,0])

                lw_outputs_s[l_,a_,c_] = mean_sem(outcome_array[:,0])
                lw_estimates_s[l_,a_,c_] = mean_sem(lw_info_array[:,1])
                lw_efforts_s[l_,a_,c_] = mean_sem(lw_info_array[:,0])

   
    return wl_outputs_s,wl_estimates_s,wl_efforts_s,lw_outputs_s,lw_estimates_s,lw_efforts_s

if __name__ == '__main__':
    if False: ## allows for easy debugging without parallel weirdness.
        s_outputs = np.empty([s_res,6,l_res,a_res,a_res,2])
        for s_ in range(s_res):
            s = s_set[s_]
            s_outputs[s_] = np.array(run_many_sims(s))
    else:
        s_outputs = Parallel(n_jobs=11)(delayed(run_many_sims)(s) for s in s_set)

    s_outputs = np.array(s_outputs)
    wl_outputs = s_outputs[:,0]
    wl_estimates = s_outputs[:,1]
    wl_efforts = s_outputs[:,2]
    lw_outputs = s_outputs[:,3]
    lw_estimates = s_outputs[:,4]
    lw_efforts = s_outputs[:,5]

    est_diff = lw_estimates[:,:,:,:,0] - wl_estimates[:,:,:,:,0]
    eff_diff = lw_efforts[:,:,:,:,0] - wl_efforts[:,:,:,:,0]
    out_diff = lw_outputs[:,:,:,:,0] - wl_outputs[:,:,:,:,0]

    est_diff_sl = est_diff[:,:,5,1]
    est_diff_ac = est_diff[7,1,:,:]

    eff_diff_sl = eff_diff[:,:,5,1]
    eff_diff_ac = eff_diff[7,1,:,:]

    out_diff_sl = out_diff[:,:,5,1]
    out_diff_ac = out_diff[7,1,:,:]

    es_sl_max = np.max(np.abs(est_diff_sl))
    ef_sl_max = np.max(np.abs(eff_diff_sl))
    ot_sl_max = np.max(np.abs(out_diff_sl))

    es_ac_max = np.max(np.abs(est_diff_ac))
    ef_ac_max = np.max(np.abs(eff_diff_ac))
    ot_ac_max = np.max(np.abs(out_diff_ac))

    est_max = max(es_sl_max,es_ac_max)
    eff_max = max(ef_sl_max,ef_ac_max)
    out_max = max(ot_sl_max,ot_ac_max)

    fig,axes = plt.subplots(3,2,sharex='col',sharey='col')

    est_slim = axes[0,0].imshow(est_diff_sl,vmin = -1*es_sl_max,vmax=es_sl_max,cmap='RdBu_r')
    est_acim = axes[0,1].imshow(est_diff_ac,vmin = -1*es_ac_max,vmax=es_ac_max,cmap='RdBu_r')
#plt.colorbar(ax=axes[:,1])

    eff_slim = axes[1,0].imshow(eff_diff_sl,vmin = -1*ef_sl_max,vmax=ef_sl_max,cmap='RdBu_r')
    eff_acim = axes[1,1].imshow(eff_diff_ac,vmin = -1*ef_ac_max,vmax=ef_ac_max,cmap='RdBu_r')

    out_slim = axes[2,0].imshow(out_diff_sl,vmin = -1*ot_sl_max,vmax=ot_sl_max,cmap='RdBu_r')
    out_acim = axes[2,1].imshow(out_diff_ac,vmin = -1*ot_ac_max,vmax=ot_ac_max,cmap='RdBu_r')

    fig.colorbar(est_slim,ax=axes[0,0])
    fig.colorbar(eff_slim,ax=axes[1,0])
    fig.colorbar(out_slim,ax=axes[2,0])

    fig.colorbar(est_acim,ax=axes[0,1])
    fig.colorbar(eff_acim,ax=axes[1,1])
    fig.colorbar(out_acim,ax=axes[2,1])

    axes[0,0].set_yticks(range(len(l_set)))
    axes[2,0].set_xticks(range(len(s_set)))
    axes[0,1].set_yticks(range(len(c_set)))
    axes[2,1].set_xticks(range(len(a_set)))

    if False:
        axes[0,0].set_yticklabels(np.round(s_set,2))
        axes[2,0].set_xticklabels(np.round(l_set,2),rotation=45)
        axes[0,1].set_yticklabels(np.round(a_set,2))
        axes[2,1].set_xticklabels(np.round(c_set,2),rotation=45)
    else:
        axes[0,0].set_yticklabels(np.round(s_set,2))
        axes[2,0].set_xticklabels(l_labels,rotation=45)
        axes[2,0].invert_xaxis()

        axes[0,1].set_yticklabels(a_labels)
        axes[2,1].set_xticklabels(a_labels,rotation=45)

    axes[0,0].set_ylabel('s value')
    axes[1,0].set_ylabel('s value')
    axes[2,0].set_ylabel('s value')
    axes[2,0].set_xlabel('l value')

    axes[0,1].set_ylabel('awareness value')
    axes[1,1].set_ylabel('awareness value')
    axes[2,1].set_ylabel('awareness value')
    axes[2,1].set_xlabel('acuity value')

    fig.set_size_inches(6.5,6.9)
    fig.tight_layout()

    fig.savefig('./figures/figB04_recency.png',dpi=300)
    fig.savefig('./figures/figB04_recency.svg')
#plt.show()

    print('Done!')
```
