## Supplemental Code for "Bayesian updating for self-assessment explains social dominance and winner-loser effects": SfigureB05.py

```
#! /usr/bin/env python

## Script to show that under bayes', the duration of the winner effect depends on the about of new info 

from bayesbots import Fish,FishNPC
from bayesbots import Fight
from bayesbots import Tank
from bayesbots import Params

import numpy as np
from matplotlib import pyplot as plt
from matplotlib import cm
from scipy.stats import ttest_ind

from tqdm import tqdm

import copy

#s,e,l = -.8,-.6,-0.99
#s,e,l=-0.9,-0.5,-0.7

params = Params()

plt.rcParams.update({'font.size': params.fig_font})

params.mean_size = 50
params.sd_size = 10
#params.poly_param_c = 0
#params.effort_method = [1,1]
params.f_method = 'shuffled'
params.n_rounds = 25
params.n_fish = 5
params.acuity = 0.1 
params.awareness = 0.5
#params.outcome_params = [s,e,l]
params.set_params()
#params.baseline_effort = 0.3

replicates = 200

opp_sizes = [40,50,60] ## size of staged fight
tank_sizes = [35,45,65,75] ## size of other fish

params_f0 = params.copy()
params_f0.prior = True
params_f0.size = 50
#f0 = Fish(0,params_f0)

n_points = params.n_rounds * (params.n_fish - 1) + 2
est_array_w = np.zeros([len(opp_sizes),replicates,n_points])
est_array_l = np.zeros_like(est_array_w)
other_fishes = [Fish(f,params) for f in range(len(tank_sizes))]

for f_ in range(len(other_fishes)):
    other_fishes[f_].size = tank_sizes[f_]

win_cases = [True,False]
est_arrays = [est_array_w,est_array_l]
for w_ in range(2):
    w = win_cases[w_]
    est_array = est_arrays[w_]
    for s_ in range(len(opp_sizes)):
        for i in tqdm(range(replicates)):
            opp_size = opp_sizes[s_]
            fishes = copy.deepcopy(other_fishes)
            f0 = Fish(0,params_f0)
            fishes.append(f0)
            tank = Tank(fishes,params)

            ## Before running the tank, simulate a win: 
            focal_fish = fishes[-1]
            f_opp = FishNPC(0,focal_fish.params)
            f_opp.size = opp_size
            focal_fight = Fight(f_opp,focal_fish,params,outcome=int(w))
            focal_fight.run_outcome()
            focal_fish.update(w,focal_fight)

## Run tank and save est_record
            tank.run_all(progress=False)
            est_array[s_,i] = np.array(f0.est_record)

fig,ax = plt.subplots()
styles = ['solid','dashed','dashdot']

for w_ in range(2):
    est_array = est_arrays[w_]
    for s_ in range(len(opp_sizes)):
        est_mean = np.mean(est_array[s_],axis=0)
        est_sem = np.std(est_array[s_],axis=0) / np.sqrt(replicates)
        xs = np.arange(len(est_mean))
        cor = cm.viridis((s_+1) / len(opp_sizes))
        ax.plot(xs,est_mean,color='black',linestyle=styles[s_]) #,label=opp_sizes[s_])
        if w_ == 1:
            ax.fill_between(xs,est_mean-est_sem,est_mean+est_sem,alpha=0.5,color=cor,label='x = ' + str(opp_sizes[s_]))
        else:
            ax.fill_between(xs,est_mean-est_sem,est_mean+est_sem,alpha=0.5,color=cor)

ax.axhline(65,color='gray',linestyle=':')
ax.axhline(75,color='gray',linestyle=':')
ax.axhline(50,color='black',linestyle=':')
ax.axhline(45,color='gray',linestyle=':')

ax.set_ylabel('Estimate')
ax.set_xlabel('Contest round')

ax.legend()

fig.set_size_inches(4,3)
fig.tight_layout()


fig.savefig('./figures/figB05_PeristEx.png',dpi=300)
fig.savefig('./figures/figB05_PersistEx.svg')
#plt.show()
```
