## Supplemental Code for "Bayesian updating for self-assessment explains social dominance and winner-loser effects": SfigureB05b.py

```
#! /usr/bin/env python

## Script to show that under bayes', the duration of the winner effect depends on the about of new info 

from bayesbots import Fish
from bayesbots import Fight
from bayesbots import Tank
from bayesbots import Params

import numpy as np
from matplotlib import pyplot as plt
import matplotlib
from scipy.stats import ttest_ind

from tqdm import tqdm

import copy
import cProfile,pstats

from joblib import Parallel, delayed

## for debugging runtime warnings:
#np.seterr(all='raise')

#profile = cProfile.Profile()
#profile.enable()

def run_experiment(params):
    fishes1 = [Fish(f,params) for f in range(params.n_fish)]
    fishes2 = [Fish(f,params) for f in range(params.n_fish)]
    tank1 = Tank(fishes1,params)
    tank2 = Tank(fishes2,params)

    tank1.run_all(False)
    tank2.run_all(False)

    sizes = [f.size for f in fishes1]
    mid1 = np.argsort(sizes)[2]
    mid2 = np.argsort([f.size for f in fishes2])[2]
    f1 = tank1.fishes[mid1]
    f2 = tank2.fishes[mid2]
    opp1 = Fish(size = f1.size*scale)
    opp2 = Fish(size = f2.size/scale)
    f_win = Fight(f1,opp1,params,outcome=0)
    f_loss = Fight(f2,opp2,params,outcome=1) 
## We skip the actual fight to save time
    f_win.winner = f1
    f_win.loser = opp1
    f_loss.winner = opp2
    f_loss.loser = f2

    ## Because this fight is never actually run, it never asks about effort, so you have to provide it
    f_win.winner.effort = 0.5
    f_win.loser.effort = 0.5
    f_loss.loser.effort = 0.5
    f_loss.winner.effort = 0.5

    f1.update(True,f_win)    
    f2.update(False,f_loss)
## Continue tanks after win/loss
    tank1_ = Tank(fishes1,params)
    tank2_ = Tank(fishes2,params)
    """
    fishes2 = [fishy.copy() for fishy in fishes]
    fishes2[0] = f
    fishes[mid1]
    tank2 = Tank(fishes2,params)

    fishes3 = [fishy.copy() for fishy in fishes]
    fishes3[1] = f2
    tank3 = Tank(fishes3,params)
    """
    tank1_.run_all(False)
    tank2_.run_all(False)

    winner_r = f1
    size_order1 = np.argsort(np.array(tank1_.sizes))
    size_order2 = np.argsort(np.array(tank2_.sizes))

    s_ranks1 = np.empty_like(size_order1)
    s_ranks2 = np.empty_like(size_order2)
    s_ranks1[size_order1] = np.arange(tank1.n_fish) ## This took me an embarrasingly long time to get right. 
    s_ranks2[size_order2] = np.arange(tank2.n_fish) ## This took me an embarrasingly long time to get right. 
    w_rank = s_ranks1[f1.idx]
    l_rank = s_ranks2[f2.idx]

    #winner_record_r = np.sum(tank1_.history[-1,f1.idx]) / (params.n_fish - 1)
    #loser_record_r = np.sum(tank2_.history[-1,f2.idx]) / (params.n_fish - 1)

### Check success against bigger and smaller
    idx_bigger_l = np.arange(tank2_.n_fish)[s_ranks2 == (l_rank + 1)][0]
    loser_v_bigger_r = tank2_.win_record[f2.idx,idx_bigger_l]/params.n_rounds
    idx_smaller_w = np.arange(tank1_.n_fish)[s_ranks1 == (w_rank - 1)][0]
    winner_v_smaller_r = tank1_.win_record[f1.idx,idx_smaller_w]/params.n_rounds
    
    idx_bigger_w = np.arange(tank1.n_fish)[s_ranks1 == (w_rank + 1)][0]
    winner_v_bigger_r = tank1_.win_record[f1.idx,idx_bigger_w]/params.n_rounds ## should this be rounds or fights.


    #biggers_r = tank.fishes1[idx_bigger]

    idx_smaller_l = np.arange(tank2.n_fish)[s_ranks2 == (l_rank - 1)][0]
    loser_v_smaller_r = tank2_.win_record[f2.idx,idx_smaller_l]/params.n_rounds

    #smaller_r = tank2_.fishes[idx_smaller]
    loser_r = f2

    #import pdb;pdb.set_trace()
    results = [winner_v_bigger_r, loser_v_bigger_r,
                winner_v_smaller_r, loser_v_smaller_r,
                winner_r, loser_r]
    return results

def run_simulation(params):
    winners = [np.nan] * replicates
    losers = [np.nan] * replicates
    biggers = [np.nan] * replicates
    smallers = [np.nan] * replicates

## So for each replicate, we fill in 
    winner_v_bigger = np.empty(replicates)
    loser_v_bigger = np.empty_like(winner_v_bigger)
    winner_v_smaller = np.empty_like(winner_v_bigger)
    loser_v_smaller = np.empty_like(winner_v_bigger)

    winner_record = np.empty_like(winner_v_bigger)
    loser_record = np.empty_like(winner_v_bigger)

    winner_v_bigger.fill(np.nan)
    winner_v_smaller.fill(np.nan)
    loser_v_bigger.fill(np.nan)
    loser_v_smaller.fill(np.nan)
    winner_record.fill(np.nan)
    loser_record.fill(np.nan)

    for r in range(replicates):
        results = run_experiment(params)

        winner_v_bigger[r],loser_v_bigger[r] = results[0:2]
        winner_v_smaller[r],loser_v_smaller[r] = results[2:4]
        winners[r],losers[r] = results[4:]

    print('winner_v_bigger:',winner_v_bigger)
    print('loser_v_bigger:',loser_v_bigger)
    print('winner_v_smaller:',winner_v_smaller)
    print('loser_v_smaller:',loser_v_smaller)

    win_outcomes = np.empty(replicates)
    loss_outcomes = np.empty_like(win_outcomes) 

    opp_measures1 = []
    opp_measures2 = []
    for f_i in range(len(winners)):
        f = winners[f_i]
        check_fish = Fish(0,params,size=f.size)
        check_fight = Fight(check_fish,f,params)
        outcome = check_fight.run_outcome()
        win_outcomes[f_i] = outcome
        opp_measures1.append([check_fish.size,check_fish.effort]) 

    for f_i in range(len(losers)):
        f = losers[f_i]

### Check loser against a size matched fish
        check_fish = Fish(0,params,size=f.size)
        check_fight = Fight(check_fish,f,params)
        outcome = check_fight.run_outcome()
        loss_outcomes[f_i] = outcome
        opp_measures2.append([check_fish.size,check_fish.effort]) 
       
    print('winners opps:',opp_measures1)
    print('losers opps:',opp_measures2)

    import pdb;pdb.set_trace()
    cleaned_wvs = winner_v_smaller[~np.isnan(winner_v_smaller)]
    cleaned_lvs = loser_v_smaller[~np.isnan(loser_v_smaller)]
    cleaned_wvb = winner_v_bigger[~np.isnan(winner_v_bigger)]
    cleaned_lvb = loser_v_bigger[~np.isnan(loser_v_bigger)]

    t_s,p_s = ttest_ind(cleaned_wvs,cleaned_lvs)
    t_b,p_b = ttest_ind(cleaned_wvb,cleaned_lvb)
    t_m,p_m = ttest_ind(win_outcomes,loss_outcomes)

    smaller_diff = winner_v_smaller - loser_v_smaller
    s_diff = np.nanmean(smaller_diff)
    s_diff_std = np.nanstd(smaller_diff)

    bigger_diff = winner_v_bigger - loser_v_bigger
    b_diff = np.nanmean(bigger_diff)
    b_diff_std = np.nanstd(bigger_diff)

    m_diff_array = win_outcomes - loss_outcomes
    m_diff = np.nanmean(m_diff_array)
    m_diff_std = np.nanstd(m_diff_array)
    
    #import pdb;pdb.set_trace()
    #ps_array[rep,i] = p_s
    #pb_array[rep,i] = p_b
    #pm_array[rep,i] = p_m

    #smean_array[rep,i] = s_diff
    #bmean_array[rep,i] = b_diff
    #mmean_array[rep,i] = m_diff

    #mstd_array[rep,i] = m_diff_std
    #sstd_array[rep,i] = s_diff_std
    #bstd_array[rep,i] = b_diff_std 

    if b_diff > 1 or s_diff > 1 or m_diff > 1:
        print('uh oh.')

    m_results = [p_m,m_diff,m_diff_std] ## this is matched opponen, winner vs loser, t-test, mean difference, std difference  (in win rate)
    s_results = [p_s,s_diff,s_diff_std]  ## this is within the tank!
    b_results = [p_b,b_diff,b_diff_std]

    tri_results = [m_results,s_results,b_results]
    return tri_results

#s,e,l = -.8,-.6,-0.99
#s,e,l=-0.9,-0.5,-0.7

## just use default params first
params = Params()
#params.xs = np.linspace(1,100,100)
#params.mean_size = 50
#params.sd_size = 20
#params.poly_param_c = 0
#params.effort_method = [1,1]
params.n_fights = 5 
params.n_fish = 5
##params.acuity = 0.1
#params.f_method = 'balanced'
#params.u_method = 'bayes'
#params.f_outcome = 'math'
#params.outcome_params = [s,e,l]
params.set_params()
## Let fish duke it out, then pull a fish out, let it win, and put it back in with the rest.
#replicates = 10
scale = 1

#plt.rcParams.update({'font.size': 18})
#plt.rcParams.update({'lines.linewidth':5})
#plt.rcParams.update({'axes.linewidth':2})

INIT = False

#ew_pairs = []

iterations = 100
min_reps = 5
max_reps = 80 
rep_step = 10

## Something is weird in parallel, it freees after around 300
#rep_array = np.array([5,20,40,65,100,150,200,300,500])
rep_array = np.array([5,20,40,65,100,150,200])
#rep_array = np.arange(min_reps,max_reps,rep_step)
r_bins = len(rep_array)

ps_array = np.empty([r_bins,iterations])
pb_array = np.empty_like(ps_array)
pm_array = np.empty_like(ps_array)

smean_array = np.empty_like(ps_array)
bmean_array = np.empty_like(ps_array)
mmean_array = np.empty_like(ps_array)

mstd_array = np.empty_like(ps_array)
sstd_array = np.empty_like(ps_array)
bstd_array = np.empty_like(ps_array)

print('n iterations:',iterations)


for rep in range(r_bins):
    replicates = rep_array[rep]
## test sim:
    #import pdb;pdb.set_trace()
    foo = run_simulation(params)
    print('running with',rep_array[rep],'replicates')
## This code was way to slow. This speeds it up by around 10x
    import pdb;pdb.set_trace()
    all_iterations = Parallel(n_jobs=20)(delayed(run_simulation)(params) for _ in tqdm(range(iterations)))
    for i in range(iterations):
        #tri_results = run_simulation(params)
        tri_results = all_iterations[i]
        m_results,s_results,b_results = tri_results
        pm_array[rep,i],mmean_array[rep,i],mstd_array[rep,i] = m_results
        ps_array[rep,i],smean_array[rep,i],sstd_array[rep,i] = s_results
        pb_array[rep,i],bmean_array[rep,i],bstd_array[rep,i] = b_results

## Things I needj
xs = rep_array
log_xs = np.log(rep_array)
log_xs = np.emath.logn(5, rep_array)

## Probabilities for size matched
prob_sig_m = pm_array < 0.05
mean_prob_m = np.nanmean(prob_sig_m,1)
std_prob_m = np.nanstd(prob_sig_m,1)

## Probabilities for smaller
prob_sig_s = ps_array < 0.05
mean_prob_s = np.nanmean(prob_sig_s,1)
std_prob_s = np.nanstd(prob_sig_s,1)

## Probabilities for bigger
prob_sig_b = pb_array < 0.05
mean_prob_b = np.nanmean(prob_sig_b,1)
std_prob_b = np.nanstd(prob_sig_b,1)

## Means (and std) for size matched, smallers, and biggers
mean_diff_m = np.nanmean(mmean_array,1)
sem_diff_m = np.nanmean(mstd_array,1) / np.sqrt(rep_array)

mean_diff_s = np.nanmean(smean_array,1)
#sem_diff_s = np.nanstd(smean_array,1) / np.sqrt(rep_array)
sem_diff_s = np.nanmean(sstd_array,1) / np.sqrt(rep_array)

mean_diff_b = np.nanmean(bmean_array,1)
#sem_diff_b = np.nanstd(bmean_array,1) / np.sqrt(rep_array)
sem_diff_b = np.nanmean(bstd_array,1) / np.sqrt(rep_array)


m_color = 'black'
s_color = 'darkblue'
b_color = 'gold'
## Plot the probability of detecting a significant difference (with std across 1000 iterations)

fig,ax = plt.subplots()
ax.plot(log_xs,mean_prob_m,color=m_color,label='Size matched')
ax.plot(log_xs,np.clip(mean_prob_m - 2*std_prob_m,0,1),linestyle=':',color=m_color)
ax.plot(log_xs,np.clip(mean_prob_m + 2*std_prob_m,0,1),linestyle=':',color=m_color)

ax.plot(log_xs,mean_prob_s,color=s_color,label='Vs. smaller',linestyle='dashed')
ax.plot(log_xs,np.clip(mean_prob_s - 2*std_prob_s,0,1),linestyle=':',color=s_color)
ax.plot(log_xs,np.clip(mean_prob_s + 2*std_prob_s,0,1),linestyle=':',color=s_color)

ax.plot(log_xs,mean_prob_b,color=b_color,label='Vs. bigger',linestyle='dashdot')
ax.plot(log_xs,np.clip(mean_prob_b - 2*std_prob_b,0,1),linestyle=':',color=b_color)
ax.plot(log_xs,np.clip(mean_prob_b + 2*std_prob_b,0,1),linestyle=':',color=b_color)

ax.set_xticks(log_xs)
ax.set_xticklabels(xs)
ax.set_xlabel('n Replicates')
ax.set_ylabel('P(getting significant result)')
ax.set_ylim([-0.1,1.1])

#ax.fill_between(xs,mean_prob_m - std_prob_m,mean_prob_m + std_prob_m)

## Plot the mean effect strength (with avg SEM for r replicates)
fig1,ax1 = plt.subplots()
#print(log_xs,mean_diff_m)
#print(mean_diff_s,mean_diff_b)
ax1.plot(log_xs,mean_diff_m,color=m_color,label='Size matched')
ax1.plot(log_xs,mean_diff_m - sem_diff_m,linestyle=':',color=m_color)
ax1.plot(log_xs,mean_diff_m + sem_diff_m,linestyle=':',color=m_color)

ax1.plot(log_xs,mean_diff_s,color=s_color,label='vs. smaller',linestyle='dashed')
ax1.plot(log_xs,mean_diff_s - sem_diff_s,linestyle=':',color=s_color)
ax1.plot(log_xs,mean_diff_s + sem_diff_s,linestyle=':',color=s_color)

ax1.plot(log_xs,mean_diff_b,color=b_color,label='vs. bigger',linestyle='dashdot')
ax1.plot(log_xs,mean_diff_b - sem_diff_b,linestyle=':',color=b_color)
ax1.plot(log_xs,mean_diff_b + sem_diff_b,linestyle=':',color=b_color)

ax1.axhline(0,color='grey')
ax1.set_xticks(log_xs)
ax1.set_xticklabels(xs)
ax1.set_xlabel('n Replicates')
ax1.set_ylabel('Mean difference observed (winner vs loser)')

ax.legend()

if True:
    fig1.savefig('./imgs/fig5bS_diff.svg')
    fig.savefig('./imgs/fig5bS_prob.svg')
if True:
    plt.show()

if np.nanmax(mean_diff_b) > 1:
    import pdb;pdb.set_trace()
if np.nanmax(mean_diff_s) > 1:
    import pdb;pdb.set_trace()
if np.nanmax(mean_diff_m) > 1:
    import pdb;pdb.set_trace()


print('Done!')
```
