## Supplemental Code for "Bayesian updating for self-assessment explains social dominance and winner-loser effects": SfigureB06.py

```
import numpy as np
from matplotlib import pyplot as plt
import matplotlib

from scipy import stats

from bayesbots import Fish,FishNPC
from bayesbots import Fight
from bayesbots import Params

from tqdm import tqdm
import copy
from joblib import Parallel, delayed

## A couple of helper functions to keep things tidy

def mean_sem(a):
    mean_a = np.nanmean(a)
    sem_a = np.nanstd(a) / np.sqrt(len(a))
    return mean_a,sem_a

def plot_fill(xs,a,fig=None,ax=None,color='grey',alpha=0.5):
    if ax is None:
        fig,ax = plt.subplots()
    ax.plot(xs,a[:,0],color='black')
    ax.fill_between(xs,a[:,0] - a[:,1],a[:,0] + a[:,1],color=color,alpha=alpha)
    return fig,ax

def run_sim(params,opp_params,npc_size_list = [50],npc_effort_list = [0.5]):
    iterations = params.iterations

## Fish WL, LW
    focal_winner = Fish(1,params)
    focal_loser = Fish(2,params)

## Cant pre run the first two fights against NPC if I want acuity
    staged_opp = FishNPC(0,opp_params)
    max_rounds = 100

    random_opp = FishNPC(1,opp_params)
    results = np.empty([iterations,2])
    for i in range(iterations):
        focal_W = copy.deepcopy(focal_winner)
        focal_L = copy.deepcopy(focal_loser)

        staged_win = Fight(staged_opp,focal_W,params,outcome=1)
        staged_win.run_outcome()
        focal_W.update(True,staged_win)

        staged_loss = Fight(staged_opp,focal_L,params,outcome=0)
        staged_loss.run_outcome()
        focal_L.update(False,staged_loss)

        #f_sizes.append(focal_winner.size)
## Stage a bunch of wins and losses against size-matched fish
        W_thresh = 60
        L_thresh = 40
        w_rounds,l_rounds = np.nan,np.nan
        npc_sizes = np.random.choice(npc_size_list,max_rounds)
        npc_efforts = np.random.choice(npc_effort_list,max_rounds)
        for n in range(max_rounds):
            if ~np.isnan(w_rounds) and ~np.isnan(l_rounds):
                break
            npc = random_opp
            npc.params.baseline_effort = npc_efforts[n]
            npc.size = npc_sizes[n]
            if np.isnan(w_rounds):
                if focal_W.estimate <= W_thresh:
                    w_rounds = n
                else:
## Run a random fight
                    random_fight = Fight(npc,focal_W,params) 
                    outcome = random_fight.run_outcome()
                    focal_W.update(outcome,random_fight)
            if np.isnan(l_rounds):
                if focal_L.estimate >= L_thresh:
                    l_rounds = n
                else:
                    random_fight = Fight(npc,focal_L,params)
                    outcome = random_fight.run_outcome()
                    focal_L.update(outcome,random_fight)
        #import pdb;pdb.set_trace()
        results[i] = w_rounds,l_rounds
    return results

#iterations = 2
iterations = 100
params = Params()
plt.rcParams.update({'font.size': params.fig_font})
plt.rcParams.update({'lines.linewidth': 1})

tick_size = (params.fig_font * 2/3)

matplotlib.rc('xtick', labelsize=tick_size) 
matplotlib.rc('ytick', labelsize=tick_size) 


params.iterations = iterations

params.size = 50
params.acuity = 0
params.prior = True

opp_params = params.copy()

s_res = 10+1
e_res = s_res

s_set = np.linspace(0,100,s_res)
e_set = np.linspace(0,1,e_res)
e_set[0] = 0.01
s_set[0] = 1

default_params = [params.outcome_params[0],params.outcome_params[2],params.awareness,params.acuity]

all_results = np.empty([s_res,e_res,iterations,2])

def build_trunc(mu,std,lower,upper):
    X = stats.truncnorm((lower-mu)/std,(upper-mu)/std,loc=mu,scale=std)
    return X

mu_s,std_s = 50,10
lower_s,upper_s = 1,100

mu_e,std_e = 0.5,0.1
lower_e,upper_e = 0.001,1

## Using the truncated distribution prevents weird bunching up at edge
#X_size = build_trunc(mu_s,std_s,lower_s,upper_s)
#X_eff = build_trunc(mu_e,std_e,lower_e,upper_e)
#npc_size_list = X_size.rvs(100)
#npc_effort_list = X_eff.rvs(100)

mu_slist = [25,50,75]
mu_elist = [0.1,0.5,0.9]
std_elist  = np.linspace(0.01,1,11)

def run_many_sims(std_s):
    #some_results = np.empty([e_res,iterations,2])
    opp_params = params.copy()
    some_results = np.empty([3,3,e_res,iterations,2])
## Naming convention is a bit crap here, sorry, but us_ is the mu_slist index, etc
    for se_ in tqdm(range(len(std_elist))):
        std_e = std_elist[se_]
        for ue_ in range(len(mu_elist)):
            mu_e = mu_elist[ue_]
            X_eff = build_trunc(mu_e,std_e,lower_e,upper_e)
            npc_effort_list = X_eff.rvs(100)
            for us_ in range(len(mu_slist)):
                mu_s = mu_slist[us_]
                X_size = build_trunc(mu_s,std_s,lower_s,upper_s)
                npc_size_list = X_size.rvs(100)
                some_results[us_,ue_,se_] = run_sim(params,opp_params,npc_size_list,npc_effort_list)
    return some_results

stdx_set = [2**n for n in np.arange(-1,10).astype(float)]
std_xs = [n for n in np.arange(-1,10).astype(int)]

all_results = Parallel(n_jobs=11)(delayed(run_many_sims)(std_x) for std_x in stdx_set)
all_results = np.array(all_results)

mean_results = np.nanmean(all_results,axis=4)
sem_results = np.nanstd(all_results,axis=4) / np.sqrt(iterations)

mean_winners = mean_results[:,:,:,:,0]
sem_winners = sem_results[:,:,:,:,0]

mean_losers = mean_results[:,:,:,:,1]
sem_losers = sem_results[:,:,:,:,1]

#import pdb;pdb.set_trace()

## Plot stuff

#fig,ax = plt.subplots(1,3,sharey=True,sharex=True)
fig,axes = plt.subplots(2,2,sharey=True,sharex=True)

styles = ['solid','dotted','dashdot']

for i_ in range(len(mu_slist)):
    mu_s = mu_slist[i_]
    mu_e = mu_elist[i_]
    style = styles[i_]

    mean_00 = mean_winners[:,i_,0,5]
    sem_00 = sem_winners[:,i_,0,5]
    up_00,down_00 = mean_00 + sem_00, mean_00 - sem_00

    mean_10 = mean_losers[:,i_,2,5]
    sem_10 = sem_losers[:,i_,2,5]
    up_10,down_10 = mean_10 + sem_10, mean_10 - sem_10

    mean_01 = mean_winners[5,0,i_,:]
    sem_01 = sem_winners[5,0,i_,:]
    up_01,down_01 = mean_01 + sem_01, mean_01 - sem_01

    mean_11 = mean_losers[5,2,i_,:]
    sem_11 = sem_losers[5,2,i_,:]
    up_11,down_11 = mean_11 + sem_11, mean_11 - sem_11

    axes[0,0].plot(std_xs,mean_winners[:,i_,0,5],label='u = ' + str(mu_s),linestyle=style,color='black')
    axes[0,0].fill_between(std_xs,down_00,up_00,alpha=0.3)

    axes[1,0].plot(std_xs,mean_losers[:,i_,2,5],label='u = ' + str(mu_s),linestyle=style,color='black')
    axes[1,0].fill_between(std_xs,down_10,up_10,alpha=0.3)

    axes[0,1].plot(std_xs,mean_winners[5,0,i_,:],label='u = ' + str(mu_e),linestyle=style,color='black')
    axes[0,1].fill_between(std_xs,down_01,up_01,alpha=0.3)

    axes[1,1].plot(std_xs,mean_losers[5,2,i_,:],label='u = ' + str(mu_e),linestyle=style,color='black')
    axes[1,1].fill_between(std_xs,down_11,up_11,alpha=0.3)

axes[0,0].legend(fontsize=tick_size)
axes[0,1].legend(fontsize=tick_size)

axes[0,0].set_ylabel('n rounds to recover')
axes[1,0].set_ylabel('n rounds to recover')
axes[1,0].set_xlabel('STD of opponent sizes')
axes[1,1].set_xlabel('STD of oppenent effort')

fig.set_size_inches(6,5)
fig.tight_layout()

fig.savefig('./figures/figB06_recovery.png',dpi=300)
fig.savefig('./figures/figB06_recovery.svg')
#plt.show()

print('done!')
```
