## Supplemental Code for "Bayesian updating for self-assessment explains social dominance and winner-loser effects": SfigureB08_B11.py

```
from bayesbots import Fish
from bayesbots import Tank
from bayesbots import Params

import numpy as np
from matplotlib import pyplot as plt
import copy
from tqdm import tqdm

from joblib import Parallel, delayed
## Build repeated sim: 

DEBUG = 0
def mean_sem(a):
    mean_a = np.nanmean(a)
    sem_a = np.nanstd(a) / np.sqrt(len(a))
    return mean_a,sem_a

def run_sim(params):
## Initialize a tank

    f_errors = np.empty([params.n_iterations,params.n_fish])
    f_initial = np.empty([params.n_iterations,params.n_fish])
    end_costs = np.empty_like(f_initial)
    init_costs = np.empty_like(f_initial)

    end_lin = np.empty_like(f_initial)
    init_lin = np.empty_like(f_initial)

    end_stab = np.empty_like(f_initial)
    init_stab = np.empty_like(f_initial)
    for i in range(params.n_iterations):
        fishes = [Fish(f,params) for f in range(params.n_fish)]

## Run the tank
        tank = Tank(fishes,params)

        tank.run_all(progress=False)

## Check accuracy (How) 
        for f_ in range(params.n_fish):
            f = tank.fishes[f_]

            last_rounds = [f.win_record[r_][3] for r_ in range(-params.n_fish + 1,0)]
            first_rounds = [f.win_record[r_][3] for r_ in range(0,params.n_fish)]
            initial_cost = np.mean(first_rounds)
            last_cost = np.mean(last_rounds)

            #print(f.size,f.estimate,f.sdest_record[-1])
            f_estimate = f.estimate
            f_estimate_0 = f.est_record[0]
            f_std_0 = f.sdest_record[0]

            f_std = f.sdest_record[-1]
            f_error = np.sqrt((f_estimate + f_std - f.size)**2 + (f_estimate - f_std - f.size)**2)
            f_error_0 = np.sqrt((f_estimate_0 + f_std_0 - f.size)**2 + (f_estimate_0 - f_std_0 - f.size)**2)
            #f_error = (f.size - f.estimate)**2 /(f.sdest_record[-1])
            init_idx = slice(0,3)
            final_idx = slice(-3,None)
            initial_linearity,_ = tank._calc_linearity(init_idx)
            last_linearity,_ = tank._calc_linearity(final_idx)

            initial_stability,_ = tank._calc_stability(init_idx)
            last_stability,_ = tank._calc_stability(final_idx)

            #f_errors[i,f_] = f.sdest_record[-1]
            f_errors[i,f_] = f_error
            f_initial[i,f_] = f_error_0

            end_costs[i,f_] = last_cost
            init_costs[i,f_] = initial_cost


            end_lin[i,f_] = last_linearity
            init_lin[i,f_] = initial_linearity

            end_stab[i,f_] = last_stability
            init_stab[i,f_] = initial_stability
            #f_initial[i,f_] = (f.size - f.est_record[0])**2 / f.est_record[0]
    out_list = [f_errors,f_initial,
                end_costs,init_costs,    
                end_lin,init_lin,
                end_stab,init_stab]
    return out_list

## should I do the whole range? 

s_res = 11
l_res = s_res
a_res = s_res

s_list = np.linspace(0,1,s_res)
l_list = np.linspace(-1,0,l_res)
a_list = np.linspace(0,1,a_res)
c_list = np.linspace(0,1,a_res)
n_list = [2,3,5,10]
n_res = len(n_list)

np.set_printoptions(formatter={'all':lambda x: str(x)})
shifted_l = (l_list + 1)/2
l_labels = np.round(np.tan(np.array(np.pi/2 - shifted_l*np.pi/2)),1).astype('str')
l_labels[0] = 'inf' 

a_labels = np.round(np.tan(np.array(a_list)*np.pi/2) * 20,1).astype('str')
a_labels[-1] = 'inf'

params = Params()
plt.rcParams.update({'font.size': params.fig_font})

params.n_rounds = 10
params.n_iterations = 100
#params.n_iterations = 3

def run_many_sims(s,params):
    params = copy.deepcopy(params)
    some_errors = np.empty([n_res,l_res,a_res,a_res,16])

    for l_ in tqdm(range(l_res)):
        l = l_list[l_]
        params.outcome_params = [s,0.5,l]
        for n_ in range(n_res):
            n = n_list[n_]
            params.n_fish = n
            for a_ in range(a_res):
                a = a_list[a_]
                params.awareness = a
                for c_ in range(a_res):
                    c = c_list[c_]
                    params.acuity = c                
                    params.set_params()
                    sim_output = run_sim(params)
                    f_errors,f_initial = sim_output[:2]
                    end_costs,init_costs = sim_output[2:4]
                    end_lin,init_lin = sim_output[4:6]
                    end_stab,init_stab = sim_output[6:8]

                    #f_errors,f_initial,end_costs,init_costs = run_sim(params)
                    some_errors[n_,l_,a_,c_,:2] = mean_sem(f_errors)
                    some_errors[n_,l_,a_,c_,2:4] = mean_sem(f_initial)
                    some_errors[n_,l_,a_,c_,4:6] = mean_sem(end_costs)
                    some_errors[n_,l_,a_,c_,6:8] = mean_sem(init_costs)
                    some_errors[n_,l_,a_,c_,8:10] = mean_sem(end_lin)
                    some_errors[n_,l_,a_,c_,10:12] = mean_sem(init_lin)
                    some_errors[n_,l_,a_,c_,12:14] = mean_sem(end_stab)
                    some_errors[n_,l_,a_,c_,14:16] = mean_sem(init_stab)
    return some_errors

def parse_results(all_results):
    all_errors = all_results[:,:,:,:,:,0:4]
    all_costs = all_results[:,:,:,:,:,4:8]
    all_lins = all_results[:,:,:,:,:,8:12]
    all_stabs = all_results[:,:,:,:,:,12:16]

    mean_errors = all_errors[:,:,:,:,:,0]
    sem_errors = all_errors[:,:,:,:,:,1]
    mean_init = all_errors[:,:,:,:,:,2]
#sem_init = all_errors[:,:,:,:,3]

    mean_costs = all_costs[:,:,:,:,:,0]
    sem_costs = all_costs[:,:,:,:,:,1]
    init_costs = all_costs[:,:,:,:,:,2]

    mean_lins = all_lins[:,:,:,:,:,0]
    sem_lins = all_lins[:,:,:,:,:,1]
    init_lins = all_lins[:,:,:,:,:,2]

    mean_stabs = all_stabs[:,:,:,:,:,0]
    sem_stabs = all_stabs[:,:,:,:,:,1]
    init_stabs = all_stabs[:,:,:,:,:,2]
    return [mean_errors,sem_errors,mean_init],[mean_costs,sem_costs,init_costs],[mean_lins,sem_lins,init_lins],[mean_stabs,sem_stabs,init_stabs]


def build_inputs(mean_array,sem_array,init_array):
    n_mean = mean_array[7,:,2,5,1]
    n_sem = sem_array[7,:,2,5,1]
    n_init = init_array[7,:,2,5,1]
    n_data = [n_mean,n_sem,n_init]

    s_mean = mean_array[:,1,2,5,1]
    s_sem = sem_array[:,1,2,5,1]
    s_init = init_array[:,1,2,5,1]
    s_data = [s_mean,s_sem,s_init]

    l_mean = mean_array[7,1,:,5,1]
    l_sem = sem_array[7,1,:,5,1]
    l_init = init_array[7,1,:,5,1]
    l_data = [l_mean,l_sem,l_init]

    a_mean = mean_array[7,1,2,:,1]
    a_sem = sem_array[7,1,2,:,1]
    a_init = init_array[7,1,2,:,1]
    a_data = [a_mean,a_sem,a_init]

    c_mean = mean_array[7,1,2,5,:]
    c_sem = sem_array[7,1,2,5,:]
    c_init = init_array[7,1,2,5,:]
    c_data = [c_mean,c_sem,c_init]
    return n_data,s_data,l_data,a_data,c_data

def make_plots(n_data,s_data,l_data,a_data,c_data,ylabel='None',fill_color='gray',l_style=None,l_label=None,fax=None):
    if fax is None:
        fig,axes = plt.subplots(1,5,sharey=True)
        red_label = None
    else:
        fig,axes = fax
        red_label = 'default value'
    n_mean,n_sem,n_init = n_data
    s_mean,s_sem,s_init = s_data
    l_mean,l_sem,l_init = l_data
    a_mean,a_sem,a_init = a_data
    c_mean,c_sem,c_init = c_data

    axes[0].plot(s_list,s_mean,color='black',linestyle=l_style)
    if l_label == 'Bayes updating' or l_label == 'Linear updating':
        axes[0].plot(s_list,s_init,linestyle=':',color=fill_color)

    axes[0].fill_between(s_list,s_mean-s_sem,s_mean+s_sem,alpha=0.5,color=fill_color)
    axes[0].set_xlabel('s value')
    axes[0].set_ylabel(ylabel)
    axes[0].axvline(0.7,color='red',linestyle=':')
    axes[0].set_xticks(s_list[::2])
    axes[0].set_xticklabels(np.round(s_list,2)[::2],rotation=45)

    axes[1].plot(l_list,l_mean,color='black',linestyle=l_style)
    if l_label == 'Bayes updating'  or l_label == 'Linear updating':
        axes[1].plot(l_list,l_init,linestyle=':',color=fill_color)
        axes[1].invert_xaxis()
    axes[1].fill_between(l_list,l_mean-l_sem,l_mean+l_sem,alpha=0.5,color=fill_color)
    axes[1].set_xlabel('predictability')
    axes[1].axvline(-0.8,color='red',linestyle=':')

    axes[1].set_xticks(l_list[::2])
    axes[1].set_xticklabels(l_labels[::2],rotation=45)

    axes[2].plot(a_list,a_mean,color='black',linestyle=l_style)
    if l_label == 'Bayes updating' or l_label == 'Linear updating':
        axes[2].plot(a_list,a_init,linestyle=':',color=fill_color)
    axes[2].fill_between(a_list,a_mean-a_sem,a_mean+a_sem,alpha=0.5,color=fill_color)
    axes[2].axvline(0.5,color='red',linestyle=':')

    axes[2].set_xlabel('self assessment error')
    axes[2].set_xticks(a_list[::2])
    axes[2].set_xticklabels(a_labels[::2],rotation=45)

    axes[3].plot(c_list,c_mean,color='black',linestyle=l_style,label=l_label)
    if l_label == 'Bayes updating' or l_label == 'Linear updating':
        axes[3].plot(c_list,c_init,linestyle=':',color=fill_color)
    axes[3].fill_between(c_list,c_mean-c_sem,c_mean+c_sem,alpha=0.5,color=fill_color)
    
    axes[3].axvline(0.1,color='red',linestyle=':',label=red_label)

    axes[3].set_xlabel('opp assessment error')
    axes[3].set_xticks(a_list[::2])
    axes[3].set_xticklabels(a_labels[::2],rotation=45)

    axes[4].plot(n_list,n_mean,color='black',linestyle=l_style,label=l_label)
    if l_label == 'Bayes updating' or l_label == 'Linear updating':
        axes[4].plot(n_list,n_init,linestyle=':',color=fill_color)
    axes[4].fill_between(n_list,n_mean-n_sem,n_mean+n_sem,alpha=0.5,color=fill_color)
    
    axes[4].axvline(5,color='red',linestyle=':',label=red_label)

    axes[4].set_xlabel('n agents')
    axes[4].set_xticks(n_list)
    axes[4].set_xticklabels(n_list)

    return fig,axes


if __name__ == "__main__":
    print('Running bayesian simulation')
    if DEBUG: ## Handy for debug
        all_results = np.empty([s_res,n_res,l_res,a_res,a_res,16])
        for s_ in tqdm(range(s_res)):
            s = s_list[s_]
            all_results[s_] = run_many_sims(s,params)

    else:
        all_results = Parallel(n_jobs=11)(delayed(run_many_sims)(s,params) for s in s_list)
        all_results = np.array(all_results)


    print('Running no-update simulation')
    null_params = params.copy()
    null_params.update_method = None

    if DEBUG: ## Handy for debug
        null_results = np.empty([s_res,l_res,a_res,a_res,16])
        for s_ in tqdm(range(s_res)):
            s = s_list[s_]
            null_results[s_] = run_many_sims(s,null_params)

    else:
        null_results = Parallel(n_jobs=11)(delayed(run_many_sims)(s,null_params) for s in s_list)
        null_results = np.array(null_results)

    print('Running Linear simulation')

    linear_params = params.copy()
    linear_params.update_method = 'linear'

    if DEBUG: ## Handy for debug
        linear_results = np.empty([s_res,n_res,l_res,a_res,a_res,16])
        for s_ in tqdm(range(s_res)):
            s = s_list[s_]
            linear_results[s_] = run_many_sims(s,linear_params)

    else:
        linear_results = Parallel(n_jobs=11)(delayed(run_many_sims)(s,linear_params) for s in s_list)
        linear_results = np.array(linear_results)

    error_info,cost_info,lin_info,stab_info = parse_results(all_results)
    null_error_info,null_cost_info,null_lin_info,null_stab_info = parse_results(null_results)
    linear_error_info,linear_cost_info,linear_lin_info,linear_stab_info = parse_results(linear_results)

    error_inputs = build_inputs(*error_info)
    cost_inputs = build_inputs(*cost_info)
    lin_inputs = build_inputs(*lin_info)
    stab_inputs = build_inputs(*stab_info)

    null_error_inputs = build_inputs(*null_error_info)
    null_cost_inputs = build_inputs(*null_cost_info)
    null_lin_inputs = build_inputs(*null_lin_info)
    null_stab_inputs = build_inputs(*null_stab_info)

    linear_error_inputs = build_inputs(*linear_error_info)
    linear_cost_inputs = build_inputs(*linear_cost_info)
    linear_lin_inputs = build_inputs(*linear_lin_info)
    linear_stab_inputs = build_inputs(*linear_stab_info)

#error_inputs = build_inputs(mean_errors,sem_errors,mean_init)
#cost_inputs = build_inputs(mean_costs,sem_costs,init_costs)
#lin_inputs = build_inputs(mean_lins,sem_lins,init_lins)
#stab_inputs = build_inputs(mean_stabs,sem_stabs,init_stabs)

    fig1,axes1 = make_plots(*null_error_inputs,fill_color='gray',l_style=None,ylabel='Self-assessment error',l_label = 'Fixed estimate')

    fig2,axes2 = make_plots(*null_cost_inputs,fill_color='gray',l_style=None,ylabel='Contest intensity',l_label='Fixed estimate')
    fig3,axes3 = make_plots(*null_lin_inputs,fill_color='gray',l_style=None,ylabel='Linearity',l_label='Fixed estimate')
    fig4,axes4 = make_plots(*null_stab_inputs,fill_color='gray',l_style=None,ylabel='Stability',l_label='Fixed estimate')


    fig1,axes1 = make_plots(*error_inputs,fax=(fig1,axes1),ylabel='Self-assessment error',fill_color='tab:blue',l_style='dashed',l_label='Bayes updating')
    fig2,axes2 = make_plots(*cost_inputs,fax=(fig2,axes2),ylabel='Contest intensity',fill_color='tab:blue',l_style='dashed',l_label='Bayes updating')
    fig3,axes3 = make_plots(*lin_inputs,fax=(fig3,axes3),ylabel='Linearity',fill_color='tab:blue',l_style='dashed',l_label='Bayes updating')
    fig4,axes4 = make_plots(*stab_inputs,fax=(fig4,axes4),ylabel='Stability',fill_color='tab:blue',l_style='dashed',l_label='Bayes updating')

    fig1,axes1 = make_plots(*linear_error_inputs,fax=(fig1,axes1),ylabel='Self-assessment error',fill_color='tab:green',l_style='dashdot',l_label='Linear updating')
    fig2,axes2 = make_plots(*linear_cost_inputs,fax=(fig2,axes2),ylabel='Contest intensity',fill_color='tab:green',l_style='dashdot',l_label='Linear updating')
    fig3,axes3 = make_plots(*linear_lin_inputs,fax=(fig3,axes3),ylabel='Linearity',fill_color='tab:green',l_style='dashdot',l_label='Linear updating')
    fig4,axes4 = make_plots(*linear_stab_inputs,fax=(fig4,axes4),ylabel='Stability',fill_color='tab:green',l_style='dashdot',l_label='Linear updating')


#fig1.legend()
#fig2.legend()
#fig3.legend()
#fig4.legend()

    fig1.set_size_inches(6.5,4)
    fig1.tight_layout()

    fig2.set_size_inches(6.5,4)
    fig2.tight_layout()

    fig3.set_size_inches(6.5,4)
    fig3.tight_layout()

    fig4.set_size_inches(6.5,4)
    fig4.tight_layout()

    fig1.savefig('./figures/figB08_error.png',dpi=300)
    fig1.savefig('./figures/figB08_error.svg')

    fig2.savefig('./figures/figB09_intensity.png',dpi=300)
    fig2.savefig('./figures/figB09_intensity.svg')
    fig3.savefig('./figures/figB10_linearity.png',dpi=300)
    fig3.savefig('./figures/figB10_linearity.svg')
    fig4.savefig('./figures/figB11_stability.png',dpi=300)
    fig4.savefig('./figures/figB11_stability.svg')

#plt.show()
```
