## Supplemental Code for "Bayesian updating for self-assessment explains social dominance and winner-loser effects": SfigureB12.py

```
#! /usr/bin/env python

## Code for fig 6a, showing how size of opponent impacts the strength of winner effect on different individuals


import numpy as np
from matplotlib import pyplot as plt
import matplotlib

from bayesbots import Params
from bayesbots import Fish,FishNPC
from bayesbots import Fight
from tqdm import tqdm

sizes = np.linspace(1,100)

params = Params()
plt.rcParams.update({'font.size': params.fig_font})
plt.rcParams.update({'lines.linewidth': 1})

tick_size = (params.fig_font * 2/3)

matplotlib.rc('xtick', labelsize=tick_size) 
matplotlib.rc('ytick', labelsize=tick_size) 

params.prior = True
params.size = 50

## Make lots of NPC fish of different size
npcs = [FishNPC() for s in sizes]
for n_ in range(len(npcs)):
    npcs[n_].size = sizes[n_]

iterations = 1000
strategies = ['bayes','linear','no update']
## Make lots of focal fish with different strategies

outcome_array = np.empty([len(sizes),len(strategies),iterations,2])
outcome_array.fill(np.nan)

for o_ in tqdm(range(len(sizes))):
    opp = npcs[o_]
    for s_ in range(len(strategies)):
        f_params = params.copy()
        f_params.update_method = strategies[s_]
        for i in range(iterations):
## Stage a fight
            focal_fish = Fish(0,f_params)
            focal_loser = Fish(1,f_params)
            stage_fight = Fight(opp,focal_fish,f_params,outcome=1)
            stage_fight.run_outcome()
            focal_fish.update(True,stage_fight)

            stage_loss = Fight(opp,focal_loser,f_params,outcome=0)
            stage_loss.run_outcome()
            focal_loser.update(False,stage_loss)

## ASsay against naive size matched fish (with accuractely centered prior)
            assay_opp = Fish(1,f_params) 
            assay_fight = Fight(assay_opp,focal_fish,f_params)
            outcome = assay_fight.run_outcome()
            outcome_array[o_,s_,i,1] = outcome

            assay_loser = Fight(assay_opp,focal_loser,f_params)
            outcome_l = assay_loser.run_outcome()
            outcome_array[o_,s_,i,0] = outcome_l

fig,ax = plt.subplots()

cors = ['tab:blue','tab:green','grey']
styles = ['solid','dashed','dashdot']

for win in [0,1]:
    for s_ in range(len(strategies)):
        mean_outcome = np.mean(outcome_array[:,s_,:,win],axis=1)
        sem_outcome = np.std(outcome_array[:,s_,:,win],axis=1) / np.sqrt(iterations)
        if win == 0:
            label = strategies[s_]
        else:
            label = None
        ax.plot(sizes,mean_outcome,label=label,color='black',linestyle=styles[s_])            
        ax.fill_between(sizes,mean_outcome - sem_outcome,mean_outcome+sem_outcome,alpha=0.5,color=cors[s_])

ax.set_xlabel('Size of opponent for staged win')
ax.set_ylabel('Probability of winning vs. size-matched opponent')
ax.set_ylabel('Assay win rate')
ax.axhline(0.5,color='black')

ax.legend(fontsize=tick_size,loc='lower left')
fig.set_size_inches(4.5,3)
fig.tight_layout()

fig.savefig('./figures/figB12_discrepencyEx.png',dpi=300)
fig.savefig('./figures/figB12_discrepencyEx.svg')

#fig.savefig('./figures/fig6a_discrepency.png',dpi=300)
#plt.show()
## Check against a fair fish
## Run them against a naive, size-matched fish to test for winner effect  
# (do I do win percent, or just contrast winner vs loser)
```
