## Supplemental Code for "Bayesian updating for self-assessment explains social dominance and winner-loser effects": SfigureB13.py

```
#! /usr/bin/env python

## Code for fig 6a, showing how size of opponent impacts the strength of winner effect on different individuals


import numpy as np
from matplotlib import pyplot as plt

from bayesbots import Params
from bayesbots import Fish,FishNPC
from bayesbots import Fight
import copy

from tqdm import tqdm
from joblib import Parallel, delayed

#sizes = np.linspace(1,100)
sizes = [40,60]

params = Params()
plt.rcParams.update({'font.size': params.fig_font})

params.prior = True
params.size = 50

## Make lots of NPC fish of different size
npcs = [FishNPC() for s in sizes]
for n_ in range(len(npcs)):
    npcs[n_].size = sizes[n_]

iterations = 1000
#strategies = ['bayes','linear','no update']

def mean_sem(a):
    mean_a = np.nanmean(a,axis=0)
    sem_a = np.nanmean(a,axis=0) / np.sqrt(len(a))
    return mean_a,sem_a

def run_sim(params):
    f_params = params
    outcome_array = np.empty([iterations,2,2]) ## iterations x w/l x sizes
    outcome_array.fill(np.nan)
    platos_winner = Fish(0,f_params)
    platos_loser = Fish(0,f_params)
    start_prior = np.array(platos_winner.prior)
    start_estimate = platos_winner.estimate
    assay_opp = Fish(1,f_params) 
    for o_ in range(len(sizes)):
        opp = npcs[o_]
        for i in range(iterations):
## Stage a fight
            #focal_fish = Fish(0,f_params)
            #focal_loser = Fish(1,f_params)
            #focal_winner = copy.deepcopy(platos_winner)
            #focal_loser = copy.deepcopy(platos_loser)
            #print(params.S,params.L,params.awareness,params.acuity)
            focal_fish = platos_winner
            focal_loser = platos_loser
            focal_fish.estimate = start_estimate
            focal_fish.prior = start_prior
            focal_loser.estimate = start_estimate
            focal_loser.prior = start_prior

            stage_fight = Fight(opp,focal_fish,f_params,outcome=1)
            stage_fight.run_outcome()
            focal_fish.update(True,stage_fight)

            stage_loss = Fight(opp,focal_loser,f_params,outcome=0)
            stage_loss.run_outcome()
            focal_loser.update(False,stage_loss)

## ASsay against naive size matched fish (with accuractely centered prior)
            assay_fight = Fight(assay_opp,focal_fish,f_params)
            outcome = assay_fight.run_outcome()
            outcome_array[i,1,o_] = outcome

            assay_loser = Fight(assay_opp,focal_loser,f_params)
            outcome_l = assay_loser.run_outcome()
            outcome_array[i,0,o_] = outcome_l
    return outcome_array

s_res = 11
l_res = s_res
a_res = s_res
c_res = a_res

s_list = np.linspace(0,1,s_res)
l_list = np.linspace(-1,0,l_res)
a_list = np.linspace(0,1,a_res)
c_list = np.linspace(0,1,c_res)

np.set_printoptions(formatter={'all':lambda x: str(x)})
shifted_l = (l_list + 1)/2
l_labels = np.round(np.tan(np.array(np.pi/2 - shifted_l*np.pi/2)),1).astype('str')
l_labels[0] = 'inf' 

a_labels = np.round(np.tan(np.array(a_list)*np.pi/2) * 20,1).astype('str')
a_labels[-1] = 'inf'

## The last two are winners/losers x mean/sem
all_results = np.empty([s_res,l_res,a_res,c_res,2,2])
def run_many_sims(s):
    many_results = np.empty([l_res,a_res,c_res,2,2])
    params = Params()
    params.size = 50
    params.prior = True
    for l_ in tqdm(range(l_res)):
        l = l_list[l_]
        for a_ in range(a_res):
            a = a_list[a_]
            a_list[a_]
            for c_ in range(c_res):
                c = c_list[c_]
                params.outcome_params = [s,0.5,l]
                params.awareness = a
                params.acuity = c    
                params.set_params()
                some_results = run_sim(params)
                #if l_ == 1 and s_ == 7:
                #   import pdb;pdb.set_trace()
                #some_diff = some_results[:,:,0] - some_results[:,:,1]
                mean_results,sem_results = mean_sem(some_results)
                mean_diff = mean_results[:,1] - mean_results[:,0]
                sem_diff = np.sum(sem_results,axis=1)
                #many_results[l_,a_,c_] = mean_sem(some_diff)
                many_results[l_,a_,c_,:,0] = mean_diff
                many_results[l_,a_,c_,:,1] = sem_diff
    return many_results

if __name__ == '__main__':
    if False:
        for s_ in tqdm(range(s_res)):
            s = s_list[s_]
            all_results[s_] = run_many_sims(s)
    else:
        all_results = Parallel(n_jobs=11)(delayed(run_many_sims)(s) for s in s_list)
        all_results = np.array(all_results)
#import pdb;pdb.set_trace()

    mean_winners = all_results[:,:,:,:,1,0]
    sem_winners = all_results[:,:,:,:,1,1]
    mean_losers = all_results[:,:,:,:,0,0]
    sem_losers = all_results[:,:,:,:,0,1]

    fig,axes = plt.subplots(2,2,sharey = 'col',sharex = 'col')
    vmin = -0.5
    vmax = 0.5
    cmap = 'RdBu_r'
    print('default params:',s_list[7],l_list[1],a_list[5],c_list[1])

    im = axes[0,0].imshow(mean_winners[:,:,5,1],vmin=vmin,vmax=vmax,cmap=cmap)
    axes[0,1].imshow(mean_winners[7,1,:,:],vmin=vmin,vmax=vmax,cmap=cmap)
    axes[1,0].imshow(mean_losers[:,:,5,1],vmin=vmin,vmax=vmax,cmap=cmap)
    axes[1,1].imshow(mean_losers[7,1,:,:],vmin=vmin,vmax=vmax,cmap=cmap)

    axes[1,0].set_xticks(range(l_res)) 
    axes[1,0].set_xticklabels(l_labels,rotation=45)
    axes[1,0].invert_xaxis()

    axes[0,0].set_yticks(range(s_res))
    axes[0,0].set_yticklabels(np.round(s_list,2))

    axes[0,0].set_ylabel('s value')
    axes[1,0].set_ylabel('s value')
    axes[1,0].set_xlabel('l value')

    axes[1,1].set_xticks(range(c_res))
    axes[1,1].set_xticklabels(a_labels,rotation=45)

    axes[1,1].set_yticks(range(a_res))
    axes[1,1].set_yticklabels(a_labels)

    axes[0,1].set_ylabel('a value')
    axes[1,1].set_ylabel('a value')
    axes[1,1].set_xlabel('c value')

    fig.colorbar(im,ax=axes)
    fig.set_size_inches(6,4.8)
#fig.tight_layout()

    fig.savefig('./figures/figB13_discrepency.png',dpi=300)
    fig.savefig('./figures/figB13_discrepency.svg')
#plt.show()
```
