## Supplemental Code for "Bayesian updating for self-assessment explains social dominance and winner-loser effects": SfigureB14.py

```
#! /usr/bin/env python

## Code for fig 6a, showing how size of opponent impacts the strength of winner effect on different individuals


import numpy as np
from matplotlib import pyplot as plt

from bayesbots import Params
from bayesbots import Fish,FishNPC
from bayesbots import Fight

from tqdm import tqdm
import copy

from joblib import Parallel, delayed

n_fight_list = np.arange(0,50,5)

params = Params()
plt.rcParams.update({'font.size': params.fig_font})

opp_params = Params()
#opp_params.baseline_effort = 0.5

params.prior = True
params.size = 50
#params.poly_param_m = 0.2
## Make lots of naive fish of random sizes
npcs = [Fish(s,opp_params) for s in range(int(max(n_fight_list) * 10))]
npc_sizes = [npc.size for npc in npcs]
print('Median npc size:',np.mean(npc_sizes))

staged_opp = FishNPC(0,params) ## NPCs by default always invest 0.5 effort
staged_opp.size = 50
iterations = 1000
#strategies = ['bayes','linear','no update']
strategies = ['bayes','linear']
## Make lots of focal fish with different strategies
"""
b_params = params

l_params = params.copy()
l_params.update_method = 'linear'

s_params = params.copy()
s_params.update_method = 'no update'

strat_params = [b_params,l_params,s_params]
"""

outcome_array = np.empty([len(n_fight_list),len(strategies),iterations,2])
outcome_array.fill(np.nan)

def run_simulation(n_fights,print_me=False):
    outcome_array_n = np.empty([len(strategies),iterations,2])
    staged_opp_ = copy.deepcopy(staged_opp)
    for s_ in range(len(strategies)):
        #f_params = strat_params[s_]
        f_params = params.copy()
        f_params.update_method = strategies[s_]
        if print_me:
            print(n_fights,strategies[s_])
        for i in range(iterations):
            #npcs = copy.deepcopy(naive_npcs)
            opps = np.random.choice(npcs,n_fights)
            focal_fish = Fish(0,f_params)
## Provide background experiecne
            for o in opps:
                pre_fight = Fight(o,focal_fish,f_params)
                outcome = pre_fight.run_outcome()
                focal_fish.update(outcome,pre_fight)
                #print(i,len(focal_fish.est_record),o.effort,focal_fish.effort)
## Run a staged fight
            pre_est = focal_fish.estimate
            staged_opp_.size = focal_fish.size
            staged_opp_.params.size = focal_fish.size
            focal_loser = copy.deepcopy(focal_fish)
            staged_fight = Fight(staged_opp_,focal_fish,outcome=1)
            staged_loss = Fight(staged_opp_,focal_loser,outcome=0)

            staged_fight.run_outcome()
            focal_fish.update(True,staged_fight)

            staged_loss.run_outcome()
            focal_loser.update(False,staged_loss)
            #print(n_,s_,i,'pre,post:',pre_est,focal_fish.estimate)
## ASsay against naive size matched fish (with accuractely centered prior)
            assay_opp = Fish(1,f_params) 
            assay_opp.size = focal_fish.size
            assay_fight = Fight(assay_opp,focal_fish,f_params)
            outcome = assay_fight.run_outcome()
            outcome_array_n[s_,i,1] = outcome

            assay_fight_ = Fight(assay_opp,focal_loser,f_params)
            outcome_ = assay_fight_.run_outcome()
            outcome_array_n[s_,i,0] = outcome_
    return outcome_array_n

if __name__ == "__main__":   
    print('running simulation in parallel...')
    outcome_arrays = Parallel(n_jobs=10)(delayed(run_simulation)(n_,False) for n_ in n_fight_list)

#import pdb;pdb.set_trace()
    outcome_array = np.array(outcome_arrays)

    fig,ax = plt.subplots()

    cors = ['tab:blue','tab:green','grey']
    styles = ['dashed','dashdot']

    for w_ in range(2):
        for s_ in range(len(strategies)):
            mean_outcome = np.mean(outcome_array[:,s_,:,w_],axis=1)
            sem_outcome = np.std(outcome_array[:,s_,:,w_],axis=1) / np.sqrt(iterations)
            if w_ == 0:
                label = strategies[s_]
            else:
                label = None
            ax.plot(n_fight_list,mean_outcome,label=label,color='black',linestyle=styles[s_])            
            ax.fill_between(n_fight_list,mean_outcome - sem_outcome,mean_outcome+sem_outcome,alpha=0.5,color=cors[s_])

    ax.set_ylim([0,1.0])
    ax.axhline(0.5,color='black',linestyle=':')

    ax.set_xlabel('Number of fights prior to assay')
    ax.set_ylabel('Assay win rate')
    ax.legend()

    fig.set_size_inches(4,3)
    fig.tight_layout()

    fig.savefig('./figures/figB14_experienceEx_linear.png',dpi=300)
    fig.savefig('./figures/figB14_experienceEx_linear.svg')
#fig.savefig('./figures/fig6b_experience.png',dpi=300)
#plt.show()
## Check against a fair fish
## Run them against a naive, size-matched fish to test for winner effect  
# (do I do win percent, or just contrast winner vs loser)
```
