## Supplemental Code for "Bayesian updating for self-assessment explains social dominance and winner-loser effects": SfigureB15.py

```
#! /usr/bin/env python

## Code for fig 6a, showing how size of opponent impacts the strength of winner effect on different individuals


import numpy as np
from matplotlib import pyplot as plt

from bayesbots import Params
from bayesbots import Fish,FishNPC
from bayesbots import Fight

from tqdm import tqdm
import copy

from joblib import Parallel, delayed

#n_fight_list = np.arange(0,50,5)
n_fight_list = [0,10]

params = Params()
plt.rcParams.update({'font.size': params.fig_font})

opp_params = Params()
#opp_params.baseline_effort = 0.5

params.prior = True
params.size = 50
#params.poly_param_m = 0.2
## Make lots of naive fish of random sizes
npcs = [Fish(s,opp_params) for s in range(int(max(n_fight_list) * 10))]
npc_sizes = [npc.size for npc in npcs]
print('Median npc size:',np.mean(npc_sizes))

staged_opp = FishNPC(0,params) ## NPCs by default always invest 0.5 effort
staged_opp.size = 50
params.iterations = 100
iterations = params.iterations
## Make lots of focal fish with different strategies

s_res = 11
l_res = s_res
a_res = s_res
c_res = s_res

s_list = np.linspace(0,1,s_res)
l_list = np.linspace(-1,0,l_res)
a_list = np.linspace(0,1,a_res)
c_list = np.linspace(0,1,c_res)

np.set_printoptions(formatter={'all':lambda x: str(x)})
shifted_l = (l_list + 1)/2
l_labels = np.round(np.tan(np.array(np.pi/2 - shifted_l*np.pi/2)),1).astype('str')
l_labels[0] = 'inf' 

a_labels = np.round(np.tan(np.array(a_list)*np.pi/2) * 20,1).astype('str')
a_labels[-1] = 'inf'

def run_many_sims(s):
    some_outputs = np.empty([l_res,a_res,c_res,2,iterations,2])
    params = Params()
    params.prior = True
    params.size = 50
    for l_ in tqdm(range(l_res)):
        l = l_list[l_]
        for a_ in range(a_res):
            a = a_list[a_]
            for c_ in range(c_res):
                c = c_list[c_]
                params.outcome_params = [s,0.5,l]
                params.awareness = a
                params.acuity = c
                params.set_params()
                for n_ in range(len(n_fight_list)):
                    n_fights = n_fight_list[n_]
                    some_outputs[l_,a_,c_,n_] = run_simulation(n_fights,params)
    return some_outputs

def run_simulation(n_fights,params):
    outcome_array_n = np.empty([params.iterations,2])
    staged_opp_ = copy.deepcopy(staged_opp)
    f_params = params.copy()
    focal_fish = Fish(0,f_params)
    start_prior = focal_fish.prior
    start_estimate = focal_fish.estimate

    for i in range(params.iterations):
        focal_fish.prior = start_prior
        focal_fish.estimate = start_estimate
        opps = np.random.choice(npcs,n_fights)
## Provide background experiecne
        for o in opps:
            pre_fight = Fight(o,focal_fish,f_params)
            outcome = pre_fight.run_outcome()
            focal_fish.update(outcome,pre_fight)
            #print(i,len(focal_fish.est_record),o.effort,focal_fish.effort)
## Run a staged fight
        pre_est = focal_fish.estimate
        staged_opp_.size = focal_fish.size
        staged_opp_.params.size = focal_fish.size
        focal_loser = copy.deepcopy(focal_fish)
        staged_fight = Fight(staged_opp_,focal_fish,outcome=1)
        staged_loss = Fight(staged_opp_,focal_loser,outcome=0)

        staged_fight.run_outcome()
        focal_fish.update(True,staged_fight)

        staged_loss.run_outcome()
        focal_loser.update(False,staged_loss)
        #print(n_,s_,i,'pre,post:',pre_est,focal_fish.estimate)
## ASsay against naive size matched fish (with accuractely centered prior)
        assay_opp = Fish(1,f_params) 
        assay_opp.size = focal_fish.size
        assay_fight = Fight(assay_opp,focal_fish,f_params)
        outcome = assay_fight.run_outcome()
        outcome_array_n[i,1] = outcome

        assay_fight_ = Fight(assay_opp,focal_loser,f_params)
        outcome_ = assay_fight_.run_outcome()
        outcome_array_n[i,0] = outcome_
    return outcome_array_n

if __name__ == '__main__': 
    if False:
        outcome_arrays = np.empty([s_res,l_res,a_res,c_res,2,params.iterations,2])
        for s_ in range(s_res):
            s = s_list[s_]
            outcome_arrays[s_] = run_many_sims(s) 
    else:
        print('running simulation in parallel...')
        outcome_arrays = Parallel(n_jobs=11)(delayed(run_many_sims)(s) for s in s_list)
        outcome_arrays = np.array(outcome_arrays)

    print(outcome_arrays.shape)

    mean_winners = np.nanmean(outcome_arrays[:,:,:,:,:,:,1],axis=5)
    mean_losers = np.nanmean(outcome_arrays[:,:,:,:,:,:,0],axis=5)

    winner_diff = mean_winners[:,:,:,:,0] - mean_winners[:,:,:,:,1]
    loser_diff = mean_losers[:,:,:,:,0] - mean_losers[:,:,:,:,1]

    fig,axes = plt.subplots(2,2,sharex='col',sharey='col')

    vmax = 0.5
    vmin = -0.5
    cmap = 'RdBu_r'

    im = axes[0,0].imshow(winner_diff[:,:,5,1],vmin=vmin,vmax=vmax,cmap=cmap)
    axes[0,1].imshow(winner_diff[7,1,:,:],vmin=vmin,vmax=vmax,cmap=cmap)
    axes[1,0].imshow(loser_diff[:,:,5,1],vmin=vmin,vmax=vmax,cmap=cmap)
    axes[1,1].imshow(loser_diff[7,1,:,:],vmin=vmin,vmax=vmax,cmap=cmap)

    axes[1,0].set_xticks(range(l_res))
    axes[1,0].set_xticklabels(l_labels,rotation=45)
    axes[1,0].invert_xaxis()

    axes[0,0].set_yticks(range(s_res))
    axes[0,0].set_yticklabels(np.round(s_list,2))

    axes[0,0].set_ylabel('s value')
    axes[1,0].set_ylabel('s value')
    axes[1,0].set_xlabel('l value')

    axes[1,1].set_xticks(range(c_res))
    axes[1,1].set_xticklabels(a_labels,rotation=45)

    axes[1,1].set_yticks(range(a_res))
    axes[1,1].set_yticklabels(a_labels)

    axes[0,1].set_ylabel('a value')
    axes[1,1].set_ylabel('a value')
    axes[1,1].set_xlabel('c value')

    fig.colorbar(im,ax=axes)
    fig.set_size_inches(6,5)
#fig.tight_layout()

    fig.savefig('./figures/figB15_experience.png',dpi=300)
    fig.savefig('./figures/figB15_experience.svg')

#plt.show()
```
