## Supplemental Code for "Bayesian updating for self-assessment explains social dominance and winner-loser effects": SfigureB16.py

```
#! /usr/bin/env python

## Script to show that under bayes', the duration of the winner effect depends on the about of new info 

from bayesbots import Fish
from bayesbots import Fight
from bayesbots import Tank
from bayesbots import Params

import numpy as np
from scipy.stats import binom_test,norm
from scipy.stats import f_oneway,pearsonr

from matplotlib import pyplot as plt
from matplotlib import cm

from scipy.ndimage import gaussian_filter1d
from tqdm import tqdm

import random, copy
import itertools

from joblib import Parallel, delayed

params = Params()
plt.rcParams.update({'font.size': params.fig_font})

params.iterations = 100
#params.iterations = 3
params.n_fights = 3

size_res = 11
sizes = np.linspace(10,90,size_res)

## Set up the various fish conditions
opp_params = params.copy()
opp_params.size = 50

s_res = 11
l_res = s_res
a_res = s_res
c_res = s_res

s_list = np.linspace(0,1,s_res)
l_list = np.linspace(-1,0,l_res)
a_list = np.linspace(0,1,a_res)
c_list = np.linspace(0,1,c_res)

np.set_printoptions(formatter={'all':lambda x: str(x)})
shifted_l = (l_list + 1)/2
l_labels = np.round(np.tan(np.array(np.pi/2 - shifted_l*np.pi/2)),1).astype('str')
l_labels[0] = 'inf' 

a_labels = np.round(np.tan(np.array(a_list)*np.pi/2) * 20,1).astype('str')
a_labels[-1] = 'inf'

def run_sim(self_params):
    opp_fish = Fish(0,opp_params)
    r_array = np.empty(params.iterations)

    for i in range(params.iterations):
        effort_array = np.empty([size_res,params.n_fights])
        for n in range(size_res):
            self_params.size = sizes[n]
            p = Fish(n+1,self_params)
            self_fight = Fight(p,opp_fish,self_params)
            for m in range(params.n_fights):
                self_fight.run_outcome()
                effort_array[n,m] = p.effort

        p_norms = np.mean(effort_array,axis=1)

        #r,p = pearsonr(sizes/100,p_norms)
        slope,_ = np.polyfit(sizes/100,p_norms,1)
        #r_array[i] = r
        r_array[i] = slope
    return r_array

def run_many_sims(s):
    some_arrays = np.empty([l_res,c_res,a_res,params.iterations])
    s_params = params.copy()

    for l_ in tqdm(range(l_res)):
        l = l_list[l_]
        for a_ in range(a_res):
            a = a_list[a_]
            for c_ in range(c_res):
                c = c_list[c_]
                s_params.outcome_params = [s,0.5,l]
                s_params.awareness = a
                s_params.acuity = c
                s_params.set_params()

                some_arrays[l_,a_,c_] = run_sim(s_params)
    return some_arrays

if False:
    all_arrays = np.empty([s_res,l_res,c_res,a_res,params.iterations])

    for s_ in range(s_res):
        s = s_list[s_]
        all_arrays[s_] = run_many_sims(s)
else:
    all_arrays = Parallel(n_jobs=11)(delayed(run_many_sims)(s) for s in s_list)
    all_arrays = np.array(all_arrays)


fig,axes = plt.subplots(1,2)

mean_r = np.nanmean(all_arrays,axis=4)

#vmax = np.nanmax(mean_r)
vmax = 1.1
vmin = -1.1
cmap = 'RdBu_r'

axes[0].imshow(mean_r[:,:,5,1],vmin=vmin,vmax=vmax,cmap=cmap)
im = axes[1].imshow(mean_r[7,1,:,:],vmin=vmin,vmax=vmax,cmap=cmap)

axes[0].set_xticks(range(l_res))
axes[0].set_xticklabels(l_labels,rotation=45)
axes[0].invert_xaxis()

axes[0].set_yticks(range(s_res))
axes[0].set_yticklabels(np.round(s_list,2))

axes[0].set_ylabel('s value')
axes[0].set_xlabel('l value')

axes[1].set_xticks(range(c_res))
axes[1].set_xticklabels(a_labels,rotation=45)

axes[1].set_yticks(range(a_res))
axes[1].set_yticklabels(a_labels)

axes[1].set_ylabel('a value')
axes[1].set_xlabel('c value')

fig.colorbar(im,ax=axes)

fig.set_size_inches(6.5,3)
#fig.tight_layout()

fig.savefig('./figures/figB16_effCorr.png',dpi=300)
fig.savefig('./figures/figB16_effCorr.svg')
#plt.show()

#import pdb;pdb.set_trace()
```
