## Supplemental Code for "Bayesian updating for self-assessment explains social dominance and winner-loser effects": SfigureB20.py

```
import numpy as np
from matplotlib import pyplot as plt
import matplotlib

from bayesbots import Fish,FishNPC
from bayesbots import Fight
from bayesbots import Params

from tqdm import tqdm

from SfigureB04 import run_sim as check_recency
from SfigureB04 import mean_sem

from SfigureB15 import run_simulation as check_certainty
from SfigureB13 import run_sim as check_discrepency
from SfigureB13 import mean_sem as mean_sem0

from SfigureB08_B11 import run_sim as check_groups

params = Params()
params.effort_method='ScaledPoly'
params.iterations = 1000
params.n_iterations = params.iterations
assay_params = params.copy()
params.size = 50
params.n_fish = 5
e_res = 11
e_list = np.linspace(0,1,e_res)

def recency_sims(base_params=None,assay_params=None,fax=None,label=None,linestyle=None,color=None,save=True):
    wl_estimates = np.empty([e_res,2])
    wl_efforts = np.empty_like(wl_estimates)
    wl_outputs = np.empty_like(wl_estimates)

    lw_estimates = np.empty_like(wl_estimates)
    lw_efforts = np.empty_like(wl_estimates)
    lw_outputs = np.empty_like(wl_estimates)


    for e_ in tqdm(range(e_res)):
        e = e_list[e_]
        if base_params is None:
            params = Params()
        else:
            params = base_params.copy()
        params.outcome_params = [0.7,e,-0.8]
        params.set_params()
        if assay_params is None:
            assay_params = params.copy()

        outcome_array,wl_info_array,lw_info_array = check_recency(params,assay_params)
        wl_outputs[e_] = mean_sem(outcome_array[:,1])
        wl_estimates[e_] = mean_sem(wl_info_array[:,1])
        wl_efforts[e_] = mean_sem(wl_info_array[:,0])

        lw_outputs[e_] = mean_sem(outcome_array[:,0])
        lw_estimates[e_] = mean_sem(lw_info_array[:,1])
        lw_efforts[e_] = mean_sem(lw_info_array[:,0])

    eff_diff = lw_outputs[:,0] - wl_outputs[:,0]
    est_diff = lw_estimates[:,0] - wl_estimates[:,0] 
    out_diff = lw_outputs[:,0] - wl_outputs[:,0]

    mean_out = out_diff
    sem_out = (lw_outputs[:,1] + wl_outputs[:,1]) / 2
    if fax is None:
        fig,ax = plt.subplots()
    else:
        fig,ax = fax
    ax.plot(e_list,mean_out,color='black',label=label,linestyle=linestyle)
    ax.fill_between(e_list,mean_out - sem_out,mean_out + sem_out,alpha=0.5,color=color) 
    ymin,ymax = ax.get_ylim()
    new_ymin = min(-0.025,np.nanmin(mean_out))
    new_ymax = np.nanmax(mean_out)*1.3
    ymin,ymax = min([ymin,new_ymin]),max([ymax,new_ymax])

    #ax.set_ylim([min(-0.025,np.nanmin(mean_out)),np.nanmax(mean_out)*1.3])
    ax.set_ylim([ymin,ymax])
    ax.axvline(0,color='black',linestyle=':')

    if save:
        fig.set_size_inches(3,3)
        fig.tight_layout()
        fig.savefig('./figures/Sfigure20a.png',dpi=300)
        fig.savefig('./figures/Sfigure20a.svg')
    return fig,ax

def certainty_sims(base_params=None,fax=None,label=None,linestyle=None,color=None,save=True):
    n_fight_list = [0,10]
    if base_params is None:
        base_params = Params()
    outcome_arrays = np.empty([e_res,2,base_params.iterations,2])
    for e_ in tqdm(range(e_res)):
        for n_ in range(len(n_fight_list)):
            e = e_list[e_]
            params = base_params.copy()
            params.outcome_params = [0.7,e,-0.8]
            params.set_params()
            n_fights = n_fight_list[n_]

            outcome_arrays[e_,n_] = check_certainty(n_fights,params)
    mean_winners = np.nanmean(outcome_arrays[:,:,:,1],axis=2)
    sem_winners = np.nanstd(outcome_arrays[:,:,:,1],axis=2) / np.sqrt(params.iterations)

    mean_losers = np.nanmean(outcome_arrays[:,:,:,0],axis=2)
    sem_losers = np.nanstd(outcome_arrays[:,:,:,0],axis=2) / np.sqrt(params.iterations)

    winner_diff = mean_winners[:,0] - mean_winners[:,1]
    winner_sem = (sem_winners[:,0] + sem_winners[:,1]) / 2
    loser_diff = mean_losers[:,0] - mean_losers[:,1]
    loser_sem = (sem_losers[:,0] + sem_losers[:,1]) / 2

    if fax is None:
        fig,ax = plt.subplots()
    else:
        fig,ax = fax
    ax.plot(e_list,winner_diff,color='black',label=label,linestyle='solid')
    ax.fill_between(e_list,winner_diff - winner_sem,winner_diff + winner_sem,alpha=0.5,color=color) 

    ax.plot(e_list,loser_diff,color='black',label=label,linestyle=linestyle)
    ax.fill_between(e_list,loser_diff - loser_sem,loser_diff + loser_sem,alpha=0.5,color=color) 

    ax.set_ylim(-1,1)
    ax.axhline(0,color='black',linestyle=':')

    if save:
        fig.set_size_inches(3,3)
        fig.tight_layout()
        fig.savefig('./figures/Sfigure20b.png',dpi=300)
        fig.savefig('./figures/Sfigure20b.svg')
    return fig,ax

def discrepency_sims(base_params=None,fax=None,label=None,linestyle=None,color=None,save=True):
    n_fight_list = [0,10]
    if base_params is None:
        base_params = Params()
    outcome_array = np.empty([e_res,2,2])
    for e_ in tqdm(range(e_res)):
        e = e_list[e_]
        params = base_params.copy()
        params.outcome_params = [0.7,e,-0.8]
        params.set_params()
        some_results = check_discrepency(params)
        mean_results,sem_results = mean_sem0(some_results)
        mean_diff = mean_results[:,1] - mean_results[:,0]
        sem_diff = np.sum(sem_results,axis=1)
        outcome_array[e_,:,0] = mean_diff
        outcome_array[e_,:,1] = sem_diff
            
    mean_winners = outcome_array[:,1,0]
    sem_winners = outcome_array[:,1,1]
    mean_losers = outcome_array[:,0,0]
    sem_losers = outcome_array[:,0,1]
  
 
    if fax is None:
        fig,ax = plt.subplots()
    else:
        fig,ax = fax
    ax.plot(e_list,mean_winners,color='black',label=label,linestyle='solid')
    ax.fill_between(e_list,mean_winners - sem_winners,mean_winners + sem_winners,alpha=0.5,color=color) 

    ax.plot(e_list,mean_losers,color='black',label=label,linestyle=linestyle)
    ax.fill_between(e_list,mean_losers - sem_losers,mean_losers + sem_losers,alpha=0.5,color=color) 

    ax.set_ylim(-1,1)
    ax.axhline(0,color='black',linestyle=':')

    if save:
        fig.set_size_inches(3,3)
        fig.tight_layout()
        fig.savefig('./figures/Sfigure20c.png',dpi=300)
        fig.savefig('./figures/Sfigure20c.svg')
    return fig,ax


def parse_sim(sim_output):
    f_errors,f_initial = sim_output[:2]
    end_costs,init_costs = sim_output[2:4]
    end_lin,init_lin = sim_output[4:6]
    end_stab,init_stab = sim_output[6:8]
    output_array = np.empty(16)

    output_array[:2] = mean_sem(f_errors)
    output_array[2:4] = mean_sem(f_initial)
    output_array[4:6] = mean_sem(end_costs)
    output_array[6:8] = mean_sem(init_costs)
    output_array[8:10] = mean_sem(end_lin)
    output_array[10:12] = mean_sem(init_lin)
    output_array[12:14] = mean_sem(end_stab)
    output_array[14:16] = mean_sem(init_stab)
    return output_array

def parse_results(all_results):
    all_errors = all_results[:,0:4]
    all_costs = all_results[:,4:8]
    all_lins = all_results[:,8:12]
    all_stabs = all_results[:,12:16]

    mean_errors = all_errors[:,0]
    sem_errors = all_errors[:,1]
    mean_init = all_errors[:,2]
#sem_init = all_errors[:,:,:,:,3]

    mean_costs = all_costs[:,0]
    sem_costs = all_costs[:,1]
    init_costs = all_costs[:,2]

    mean_lins = all_lins[:,0]
    sem_lins = all_lins[:,1]
    init_lins = all_lins[:,2]

    mean_stabs = all_stabs[:,0]
    sem_stabs = all_stabs[:,1]
    init_stabs = all_stabs[:,2]
    return [mean_errors,sem_errors,mean_init],[mean_costs,sem_costs,init_costs],[mean_lins,sem_lins,init_lins],[mean_stabs,sem_stabs,init_stabs]


def group_sims(base_params=None,fax=None,label=None,linestyle=None,color=None,save=True):
    if base_params is None:
        base_params = Params()
    outcome_array = np.empty([e_res,16])
    
## I could parallelize this...
    for e_ in tqdm(range(e_res)):
        e = e_list[e_]
        params = base_params.copy()
        params.outcome_params = [0.7,e,-0.8]
        params.set_params()
 
        sim_output = check_groups(params)
        outcome_array[e_] = parse_sim(sim_output)
        error_info,cost_info,lin_info,stab_info = parse_results(outcome_array)

    if fax is None:
        fig,axes = plt.subplots(4)
    else:
        fig,axes = fax
    axes[0].plot(e_list,error_info[0],color='black',label=label,linestyle=linestyle)
    axes[0].fill_between(e_list,error_info[0] - error_info[1],error_info[0] + error_info[1],alpha=0.5,color=color) 
    axes[0].plot(e_list,error_info[2],color=color,label=label,linestyle='dotted')
    axes[0].set_ylabel('Error')

    axes[1].plot(e_list,cost_info[0],color='black',label=label,linestyle=linestyle)
    axes[1].fill_between(e_list,cost_info[0] - cost_info[1],cost_info[0] + cost_info[1],alpha=0.5,color=color) 
    axes[1].plot(e_list,cost_info[2],color=color,label=label,linestyle='dotted')
    axes[1].set_ylabel('Cost')
    
    axes[2].plot(e_list,lin_info[0],color='black',label=label,linestyle=linestyle)
    axes[2].fill_between(e_list,lin_info[0] - lin_info[1],lin_info[0] + lin_info[1],alpha=0.5,color=color) 
    axes[2].plot(e_list,lin_info[2],color=color,label=label,linestyle='dotted')

    axes[2].set_ylabel('Linearity')


    axes[3].plot(e_list,stab_info[0],color='black',label=label,linestyle=linestyle)
    axes[3].fill_between(e_list,stab_info[0] - stab_info[1],stab_info[0] + stab_info[1],alpha=0.5,color=color) 
    axes[3].plot(e_list,stab_info[2],color=color,label=label,linestyle='dotted')
    axes[3].set_ylabel('Stability')
    if save:
        fig.set_size_inches(3,9)
        fig.tight_layout()
        fig.savefig('./figures/Sfigure20d.png',dpi=300)
        fig.savefig('./figures/Sfigure20d.svg')
    return fig,axes

if False:
    print('running recency:')
    fig,ax = recency_sims(base_params = params,save=False,label='Bayesian updating',color='tab:blue',linestyle='dashed')
    linear_params = params.copy()
    linear_params.update_method = 'linear'
    fig,ax = recency_sims(base_params = linear_params,fax=(fig,ax),save=True,label='Linear updating',color='tab:green',linestyle='dashdot')
    print('done')    

if False:
    params.update_method = 'bayes'
    print('running discrepency:')
    fig,ax = discrepency_sims(base_params = params,save=False,label='Bayesian updating',color='tab:blue',linestyle='dashed')

    linear_params = params.copy()
    linear_params.update_method = 'linear'
    fig,ax = discrepency_sims(base_params = linear_params,fax=(fig,ax),save=True,label='Linear updating',color='tab:green',linestyle='dashdot')
    print('done')

if False:
    params.update_method = 'bayes'
    params.iterations = 100
    params.n_iterations = params.iterations 
    print('running certainty:')
    fig,ax = certainty_sims(base_params = params,save=False,label='Bayesian updating',color='tab:blue',linestyle='dashed')

    linear_params = params.copy()
    linear_params.update_method = 'linear'
    fig,ax = certainty_sims(base_params = linear_params,fax=(fig,ax),save=True,label='Linear updating',color='tab:green',linestyle='dashdot')
    print('done')

if True:
    params.update_method = 'bayes'
    params.iterations = 100
    params.n_iterations = params.iterations
    print(params.iterations)
    print('running group measures:')
    fig,axes = group_sims(base_params = params,save=False,label='Bayesian updating',color='tab:blue',linestyle='dashed')

    linear_params = params.copy()
    linear_params.update_method = 'linear'
    fig,ax = group_sims(base_params = linear_params,fax=(fig,axes),save=True,label='Linear updating',color='tab:green',linestyle='dashdot')
    print('done')
```
